# Pancreatic β-cell specific loss of *E2f1* impairs insulin secretion and β-cell identity through the epigenetic repression of non β-cell programs

**DOI:** 10.1101/2020.10.14.339929

**Authors:** Frédérik Oger, Cyril Bourouh, Xavier Gromada, Maeva Moreno, Charlène Carney, Emilie Courty, Nabil Rabhi, Emmanuelle Durand, Souhila Amanzougarene, Lionel Berberian, Mehdi Derhourhi, Laure Rolland, Sarah Anissa Hannou, Pierre-Damien Denechaud, Zohra Benfodda, Patrick Meffre, Lluis Fajas, Julie Kerr-Conte, François Pattou, Philippe Froguel, Amélie Bonnefond, Jean-Sébastien Annicotte

## Abstract

The loss of pancreatic β-cell identity emerges as an important feature of type 2 diabetes development, but the molecular mechanisms are still elusive. Here, we explore the cell-autonomous role of the cell cycle regulator and transcription factor E2F1 in the maintenance of β-cell identity and insulin secretion. We show that the β-cell-specific loss of *E2f1* function in mice triggers glucose intolerance associated with defective insulin secretion, an altered α-to-β-cell ratio, a downregulation of many β-cell genes and a concomitant increase of non-β-cell markers. Mechanistically, the epigenomic profiling of non-beta cell upregulated gene promoters identified an enrichment of bivalent H3K4me3/H3K27me3 or H3K27me3 marks. Conversely, downregulated genes were enriched in active chromatin H3K4me3 and H3K27ac histone marks. We find that histone deacetylase inhibitors modulate E2F1 transcriptional and epigenomic signatures associated with these β-cell dysfunctions. Finally, the pharmacological inhibition of E2F transcriptional activity in human islets also impairs insulin secretion and the expression of β-cell identity genes. Our data suggest that E2F1 is critical for maintaining β-cell identity through a sustained repression of non β-cell transcriptional programs.

## Introduction

Type 2 Diabetes (T2D) is a progressive metabolic disorder characterized by permanent high blood glucose levels due to inadequate pancreatic β-cell response to peripheral insulin resistance. Normally, β-cells respond to obesity and aging associated insulin loss of sensibility by increasing insulin secretion to avoid rising glycemia. However, in case of imbalance between insulin secretion and action, the emerging chronic hyperglycemic state progressively leads to massive β-cell dysfunction and decreased mass (Weir & Bonner-Weir, 2004). Importantly, single cell transcriptomic analysis of human pancreatic islet cells from diabetic and healthy individuals allowed the community to define specific gene signatures of endocrine cell types and revealed a complex multi cellular lineage identity of T2D (Segerstolpe *et al*, 2016; Wang *et al*, 2016; Xin *et al*, 2016). In addition, recent studies demonstrated a loss of transcriptional maturity (Avrahami *et al*, 2020) and cell-type-specific regulatory profiles underlying T2D (Rai *et al*, 2020). Moreover, histology studies of diabetic human pancreata found a significant increase in bi-hormonal Insulin+ (Ins+)/Glucagon+ (Glu+) cells suggesting an altered β-cell identity (Dor & Glaser, 2013; Spijker *et al*, 2015; White *et al*, 2013). Accordingly, recent studies using β-cell lineage tracing murine models have demonstrated that islet cells have the capacity to directly trans-differentiate to another islet cell fate and/or de-differentiate to a progenitor-like cell (Brereton *et al*, 2014; Spijker *et al*, 2013; Spijker *et al*., 2015; Talchai *et al*, 2012). In this context, both genetic and epigenetic mechanisms are important to maintain adult β-cell fate and function in mouse and human (Bramswig *et al*, 2013; Chakravarthy *et al*, 2017; Collombat *et al*, 2009; Ediger *et al*, 2017; Gutierrez *et al*, 2017; Lawlor & Stitzel, 2019; Lu *et al*, 2018; Schaffer *et al*, 2013; Thorel *et al*, 2010; Yang *et al*, 2011). Yet, the molecular actors controlling β-cell mass, identity maintenance and cellular plasticity remain poorly understood.

Gene transcription and chromatin states are tightly regulated to ensure the appropriate transcriptome for a specific cell type. Many transcription factors have been identified as key regulators of β-cell identity and function, including Pdx1 (Gao *et al*, 2014), Pax6 (Swisa *et al*, 2017), Nkx6.1 (Schaffer *et al*., 2013) and Nkx2.2 (Gutierrez *et al*., 2017). Interestingly, pleiotropic transcription factors are also involved in the control of β-cell functions and glucose homeostasis, suggesting their important roles in activating or repressing gene transcription. Amongst those, members of E2F transcription factors (E2F1 to E2F8) family play critical roles in cell survival and proliferation, by regulating the gene expression of several proteins involved in cell-cycle progression (Kent & Leone, 2019; Poppy Roworth *et al*, 2015). The transcriptional activity of E2F1, the founder member of the family, is regulated by several protein complexes including the retinoblastoma tumor suppressor family (pRB, p107, p130), cyclin-dependent kinases (such as CDK4) and their regulatory partner cyclins (Ccn), as well as the family of the cdk inhibitors (CDKi, such as p16^Ink4A^ encoded by the *Cdkn2a* locus) (Poppy Roworth *et al*., 2015). Interestingly, the role of the cell cycle machinery goes beyond the unique regulation of cell proliferation. Indeed, modulating the expression levels of these cell cycle regulators revealed an important role for these proteins in glucose homeostasis (Denechaud *et al*, 2017) and diabetes development through the control of β-cell mass (Salas *et al*, 2014; Wang *et al*, 2015). Although *E2F1* gene expression is decreased in human T2D islets (Lupi *et al*, 2008), the causal effect of E2F1 deficiency on impaired β-cell mass, function and T2D development is not elucidated. In particular, the cellular and molecular mechanisms underlying the contribution of E2F1 as a transcription factor to β-cell identity and/or plasticity in mice and humans remain unknown. We recently demonstrated that the germline deletion of *E2f1* in the obese and diabetic *db/db* mouse model, despite lowering liver steatosis, does not protect against diabetes or obesity (Denechaud *et al*, 2016). Interestingly, we observed decreased plasma insulin levels, increased plasma glucose and glucose intolerance in *db/db::E2f1^-/-^* mice compared to *db/db::E2f1^+/+^* controls (Denechaud *et al*., 2016). These metabolic alterations in a diabetic background raises the possibility that E2f1 may contribute to islet morphology, cell identity and function in a cell-autonomous manner. To test this hypothesis, we generated mice lacking *E2f1* in β cells and identified that E2F1 is necessary for maintaining β-cell identity gene expression in both mouse models and human islets. By combining cellular and mouse models to pharmacological approaches, our results identify E2F1 as a critical transcription factor necessary to maintain proper β-cell gene expression and function, while repressing non β-cell transcriptional programs.

## Material and methods

### Materials and Oligonucleotides

Chemicals, unless stated otherwise, were purchased from Sigma-Aldrich. Anti-insulin (ab7842), anti-glucagon (ab11909) and IgG (ab37415 ChIP grade) antibodies for ChIP-seq experiments were from Abcam ; anti-E2F1 (sc-193 for immunofluorescence on mouse tissues) antibody was from Santa Cruz Biotechnology. H3K4me3 (#61379), H3K27ac (#39685) and H3K27me3 (#61017) antibodies were from Active motif. The HLM006474 compound (Ma *et al*, 2008) was synthetized as previously described (Rosales-Hurtado *et al*, 2019).

### Animal Experiments

Mice were maintained according to European Union guidelines for the use of laboratory animals. In vivo experiments were performed in compliance with the French ethical guidelines for studies on experimental animals (animal house agreement no. A 59-35015, Authorization for Animal Experimentation no.59-350294, project approval by our local ethical committee no. CEEA 482012, APAFIS#2915-201511300923025v4). All experiments were performed with male mice. Mice were housed under a 12-hr light/dark cycle and given a regular chow (A04;Safe).

*E2f1-/-* (B6; 129S4-*E2f1*tm1Meg/J) mice and *db/+* mice (Janvier Labs) were crossed to obtain *db/db::E2f1^+/+^* and *db/db::E2f1^-/-^* mice and were previously described in (Denechaud *et al*., 2016). *CMV-CDK4^R24C^* (Sotillo *et al*, 2001) and *Cdkn2a*-/- (Serrano *et al*, 1996) were described elsewhere and were crossed with *E2f1-/-* mice to obtain *E2f1-/- ::*CMV-CDK4^R24C^ and *E2f1-/-*::*Cdkn2a*-/-. *E2f1* floxed (*E2f1^flox/flox^,* Taconic Biosciences, NY, USA) mice were previously described (Denechaud *et al*., 2016; Giralt *et al*, 2018). The congenic mice carrying the floxed *E2f1* allele were thereafter mated with rat insulin II promoter (RIP)-Cre mice (Herrera, 2000) and then further intercrossed to generate pure mutant RIPcre^Tg/+^::*E2f1^flox/flox^* mice. A PCR genotyping strategy was subsequently used to identify RIPcre^+/+^::*E2f1^flox/flox^* (*E2f1^flox/flox^*), RIPcre^Tg/+^::*E2f1^+/+^* (RIPCre/+), and RIPcre^Tg/+^::*E2f1^flox/flox^* (*E2f1^β-/-^*) mice.

Metabolic phenotyping experiments were performed according to the EMPRESS protocols. Intraperitoneal glucose and insulin tolerance tests (ipGTT and ITT, respectively) were performed as previously described (Annicotte *et al*, 2009; Rabhi *et al*, 2016) on 16-hr-fasted animals for ipGTT and 5-hr-fasted animals for ITT. Glycemia was measured using the Accu-Check Performa (Roche Diagnostics). Circulating insulin levels were measured using the mouse Insulin ELISA kit (Mercodia).

### Immunofluorescence, Immunohistochemistry and Morphometry

Immunofluorescence and immunohistochemistry were performed exactly as described previously (Annicotte *et al*., 2009; Rabhi *et al*., 2016). Pancreatic tissues were fixed in 10% formalin, embedded in paraffin and sectioned at 5 µm. For immunofluorescence microscopy analyses, after antigen retrieval using citrate buffer, 5-µm formalin-fixed paraffin embedded (FFPE) pancreatic sections were incubated with the indicated antibodies. Immunofluorescence staining was revealed by using a fluorescein-isothiocyanate-conjugated anti-rabbit (for E2f1; Santa Cruz), Alexa-conjugated anti-mouse (for glucagon co-staining with E2f1) or anti-guinea pig (for insulin co-staining with E2f1) secondary antibodies. Nuclei were stained with Hoechst. For morphometric analysis, three to ten animals from each genotype were analyzed, and images were processed and quantified using ImageJ software by an observer blinded to experimental groups.

### Pancreatic Islet Studies

For mouse islet studies, pancreata were digested by type V collagenase (C9263; 1.5 mg/ml) for 10 min at 37°C as described previously (Rabhi *et al*., 2016). Briefly, after digestion and separation in a density gradient medium, islets were purified by handpicking under a macroscope and cultured during 16 hours before subsequent analysis. For glucose-stimulated insulin secretion (GSIS) tests, approximately twenty islets were exposed to either 2.8 mM or 20 mM glucose in Krebs-Ringer bicarbonate HEPES buffer containing 0.5% fatty-acid-free Bovine Serum Albumin (BSA). Insulin released in the medium was measured 1 hr later using the mouse insulin ELISA kit (Mercodia). For expression studies, mouse isolated islets were snap-frozen in liquid nitrogen before RNA extraction. Human pancreatic tissue was harvested from brain-dead, non-diabetic adult human donors (see Supplementary Table S1 for donor information). Isolation and islets culture were performed as described elsewhere (Kerr-Conte *et al*, 2010). Human islets were treated for 48 hours with DMSO 0.1% or HLM006474 at 10 µM. Data are expressed as a ratio of total insulin content. For mRNA and protein quantification, human islets were isolated as described above and snap-frozen for further processing.

### Cell Culture and Pharmacological Treatments

Min6 cells (AddexBio) were cultured in DMEM (Gibco) with 15% fetal bovine serum, 100 mg/ml penicillin-streptomycin, and 55 mM beta-mercaptoethanol. Cells were treated with Trichostatin A (TSA) at 0.5 µM and subjected to GSIS or RNA extraction 16 hr later. Min6 cells were treated with HLM006474 (10 µM) or DMSO 0.1% (Ma *et al*., 2008; Rosales-Hurtado *et al*., 2019) for 48 hr before GSIS assay or RNA extraction.

### RNA extraction, qPCR and RNA-Sequencing

Total RNA was extracted from cells and tissues using trizol reagent (Life Technologies) as described previously. mRNA expression was measured after reverse transcription by quantitative real-time PCR (qRT-PCR) with FastStart SYBR Green master mix (Roche) using a LightCycler Nano or LC480 instruments (Roche). qRT-PCR results were normalized to endogenous cyclophilin reference mRNA levels. The results are expressed as the relative mRNA level of a specific gene expression using the formula 2^-Δ^ . The complete list of primers is presented in Supplementary Table S2. For RNA sequencing, total RNA was extracted from Min6 cells or pancreatic islets using the RNeasy Plus Microkit (Qiagen) following manufacturer’s instructions. RNA quality was verified using RNA 6000 nanochips on the Agilent 2100 bioanalyzer. Purified RNA (50ng) with RNA integrity number ≥6.5 was subsequently used for library preparation (TruSeq Stranded mRNA Library Preparation Kit, Illumina) and sequenced on a HiSeq2500 system (Illumina). 3 biological replicates per condition were sequenced using paired-end mode. A mean of 54 million paired-end reads of 75 bp were generated for each sample. After initial checks and validation of sequence quality, RNA-seq reads were aligned to the mouse reference genome (mm10) using TopHat2. Subsequently, both quantification and annotation of the reads were performed using Bioconductor package Rsubread. Finally, the differential gene expression analyses were performed using Bioconductor package DESeq2. Using a *P<0.05* adjusted for multiple comparisons as threshold, we then performed pathway analysis using Ingenuity Pathway Analysis (Ingenuity Systems, Qiagen), Metascape (Zhou *et al*, 2019) and Gene Set Enrichment Analysis (GSEA; http://software.broadinstitute.org/gsea/).

### Chromatin immunoprecipitation and ChIP sequencing

ChIP experiments were performed on formaldehyde-fixed Min6 cells. Briefly, 20.10^6^ Min6 cells were treated with formaldehyde at a final concentration of 1% to crosslink DNA and protein complexes during 10 min. The reaction was stopped by the addition of glycine (0.125 M) during 5 min. Cells were lysed and DNA-protein complexes were sheared using the Bioruptor Pico (Diagenode, ref B01060010) for 8 minutes. The sheared chromatin was immunoprecipitated with either the non-specific antibody IgG (Santa Cruz, sc2025), H3K4me3 (Active motif, #61379), H3K27me3 (Active motif, #61017) or H3K27ac (Active motif, #39685). 1 ng of eluted and purified DNA was used to prepare DNA sequencing library with the Nextflex rapid DNA seq kit 2.0 (Perkin Elmer, NOVA-5188-01) on the NextSeq 500 system (Illumina) using single read 100 base pairs mode. The demultiplexing of sequence data (from BCL files generated by Illumina sequencing systems to standard FASTQ file formats) was performed using bcl2fastq Conversion Software (Illumina; version 2.20). Trimming of residuals adapters and low quality reads was performed using TrimGalore (version 0.4.5). Subsequently, sequence reads from FASTQ files were mapped to the mouse genome (mm10) using Bowtie2 Aligner (version 2.3.5.1). Finally, peak-calling was performed with MACS2 software (version 2.2.7.1). Further bioinformatic analyses of the public data sets and those of this study were performed using the open web-based platform Galaxy Europe (https://usegalaxy.eu). A list of ChIP-seq and RNA-seq data sets used in this study can be found in Supplementary Table S3.

### Rapid immunoprecipation mass spectometry of endogenous protein (RIME)

#### Chromatin Immunoprecipitation

Min6 cells were transfected with an empty vector (pCMV-FLAG, negative control) or a vector encoding human *E2F1* gene coupled to a FLAG tag for immunoprecipitation (pCMV-hE2F1-FLAG). 48h after transfections, cells were fixed in 1% formaldehyde for 15 min and quenched with 0.125 M glycine. Chromatin was isolated by the addition of lysis buffer, followed by disruption with a Dounce homogenizer. Lysates were sonicated and the DNA sheared to an average length of 300-500 bp. Genomic DNA (Input) was prepared by treating aliquots of chromatin with RNase, proteinase K and heat for de-crosslinking, followed by ethanol precipitation. Pellets were resuspended and the resulting DNA was quantified on a NanoDrop spectrophotometer. Extrapolation to the original chromatin volume allowed quantitation of the total chromatin yield. An aliquot of chromatin (150 ug) was precleared with protein G agarose beads (Invitrogen). Proteins of interest were immunoprecipitated using 15 ug of antibody against FLAG (#F9291) and protein G magnetic beads. Protein complexes were washed then trypsin was used to remove the immunoprecipitate from beads and digested the protein sample. Protein digests were separated from the beads and purified using a C18 spin column (Harvard Apparatus). The peptides were vacuum dried using a speedvac.

#### Mass Spectrometry

Digested peptides were analyzed by LC-MS/MS on a Thermo Scientific Q Exactive Orbitrap Mass spectrometer in conjunction with a Proxeon Easy-nLC II HPLC (Thermo Scientific) and Proxeon nanospray source. The digested peptides were loaded on a 100 micron x 25 mm Magic C18 100Å 5U reverse phase trap where they were desalted online before being separated using a 75 micron x 150 mm Magic C18 200Å 3U reverse phase column. Peptides were eluted using a 90 minutes gradient with a flow rate of 300nl/min. An MS survey scan was obtained for the m/z range 300-1600, MS/MS spectra were acquired using a top 15 method, where the top 15 ions in the MS spectra were subjected to HCD (High Energy Collisional Dissociation). An isolation mass window of 1.6 m/z was for the precursor ion selection, and normalized collision energy of 27% was used for fragmentation. A five second duration was used for the dynamic exclusion.

#### Database Searching

Tandem mass spectra were extracted by Unspecified version Unspecified. Charge state deconvolution and deisotoping were not performed. All MS/MS samples were analyzed using X! Tandem (The GPM, thegpm.org; version CYCLONE (2013.02.01.1)). X! Tandem was set up to search the mouse-F-_20150428_KK0vMm database (unknown version, 90758 entries) the cRAP database of common laboratory contaminants (www.thegpm.org/crap; 114 entries) plus an equal number of reverse protein sequences assuming the digestion enzyme trypsin. X! Tandem was searched with a fragment ion mass tolerance of 20 PPM and a parent ion tolerance of 20 PPM. Carbamidomethyl of cysteine was specified in X! Tandem as a fixed modification. Glu->pyro-Glu of the n-terminus, ammonia-loss of the n-terminus, gln->pyro-Glu of the n-terminus, deamidated of asparagine and glutamine, oxidation of methionine and tryptophan, dioxidation of methionine and tryptophan and acetyl of the n-terminus were specified in X! Tandem as variable modifications.

#### Criteria for Protein Identification

Scaffold (version Scaffold_4.5.3, Proteome Software Inc., Portland, OR) was used to validate MS/MS based peptide and protein identifications. Peptide identifications were accepted if they exceeded specific database search engine thresholds. X! Tandem identifications required at least -Log(Expect Scores) scores of greater than 1.5. Protein identifications were accepted if they contained at least 1 identified peptides. Proteins that contained similar peptides and could not be differentiated based on MS/MS analysis alone were grouped to satisfy the principles of parsimony. Proteins sharing significant peptide evidence were grouped into clusters.

#### List Filtering

Final list generation was done by taking all proteins with a spectral count of five and above from each replicate reaction and comparing them in a venn-diagram against IgG control replicates. Proteins unique to both experimental replicates were then applied to the PANTHER database for protein ontology results.

### Statistical Analysis

Data are presented as mean ± s.e.m. Data are derived from multiple experiments unless stated otherwise. Statistical analysis was performed using a two-tailed unpaired *t*-test or one-way or two-way ANOVA with Tukey‘s post hoc test comparing all groups to each other, using GraphPad Prism 7.0 software. Differences were considered statistically significant at p 0.05 (*p < 0.05, ** p < 0.01, and *** p < 0.001).

## Results

### Increased alpha to beta cell ratio in the pancreas of germline *E2f1*-deficient mice

To investigate the contribution of E2f1 to islet morphology, we performed immunofluorescence staining of insulin and glucagon in the pancreas of wild-type and global *E2f1* knock-out mice. The detailed analysis of *E2f1*-deficient pancreas revealed a decreased proportion of ins+ β-cell and a concomitant expansion of the glu+ α-cell percentage per islet in chow-fed 16 week-old animals, as demonstrated through the quantification of insulin and glucagon positive cells in *E2f1*+/+ and *E2f1*-/- pancreas (Figures 1A-B, Supplementary figures S1G). In the *db/db* background, *E2f1* deficient animals displayed a further increase of the α-cell number per islet, while exhibiting an even lower β-cell count compared to *db/db::E2f1^+/+^* controls (Figures 1 C-D).

**Figure 1.**
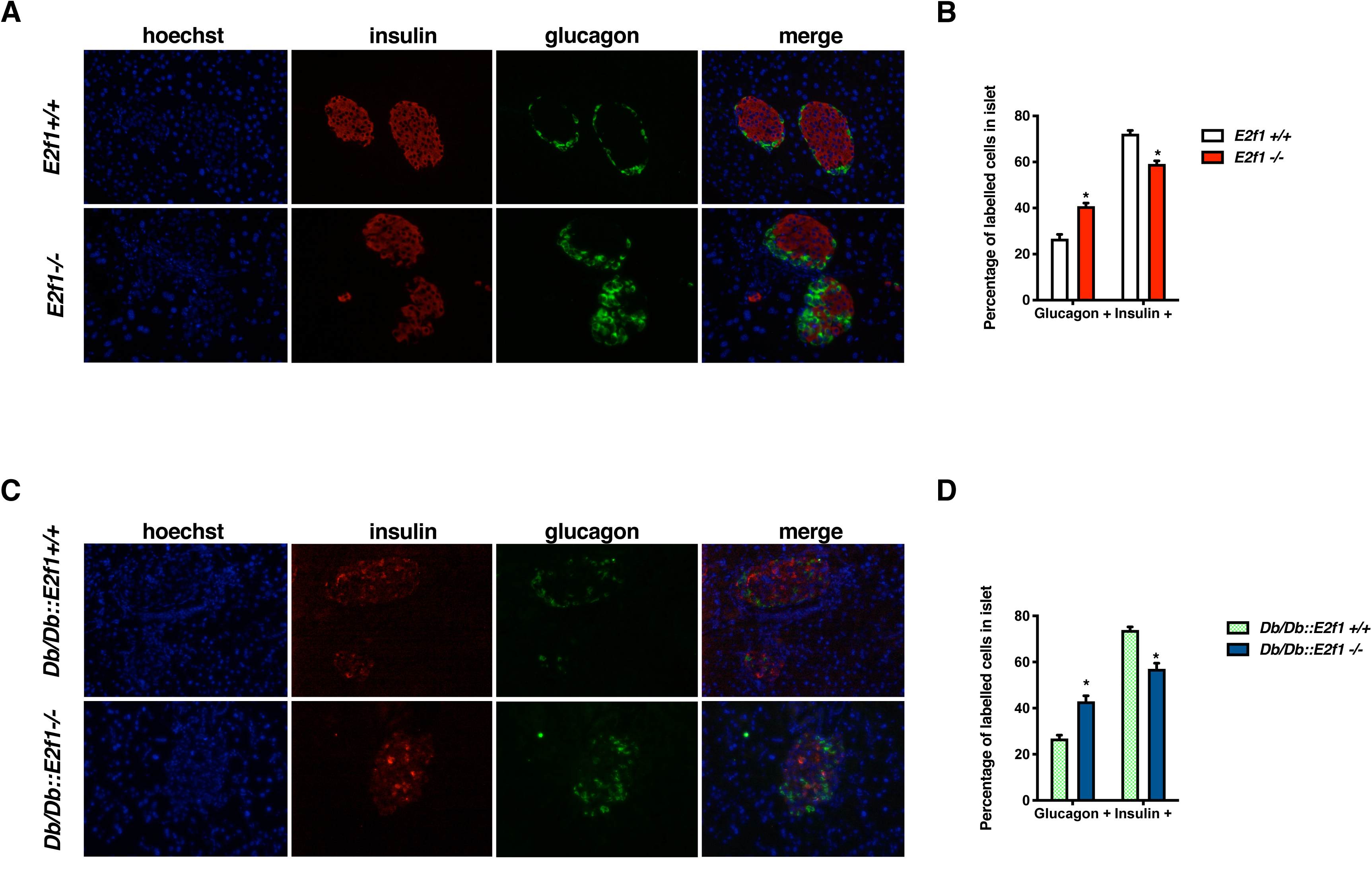
Altered α-to-β cell ratio in germline *E2f1*-deficient mice. **(A)** Representative immunofluorescent staining of insulin and glucagon in pancreatic sections from 16 week old global *E2f1* knockout male mice (*E2f1*^-/-^) compared to littermate controls (*E2f1*^+/+^). **(B)** Quantification of glucagon (Glucagon +) and insulin (Insulin +) labelled cells from (A, n=3 per gentotype). **(C)** Representative immunofluorescent staining of insulin and glucagon in pancreatic sections from 14 to 15 week old global *E2f1* knockout male mice in a *Db/Db* background (*Db/Db::E2f1*^-/-^) compared to littermate control (*Db/Db::E2f1*^+/+^). **(D)** Quantification of glucagon (Glucagon +) and insulin (Insulin +) labelled cells in *Db/Db::E2f1*^-/-^ and *Db/Db::E2f1*^+/+^ mice calculated from C (n=4 per genotype). Values in B and D are expressed as mean ± s.e.m. and were analysed by two-tailed unpaired *t*-test. *p < 0.05; **p<0.01.

To test whether *bona fide* regulators of E2F1 activity could rescue the altered α o- β-cell ratio observed in *E2f1 -/-* mice, we used 2 different genetically-engineered mouse models (Supplementary figures S1A). First, the R24C mouse model of CDK4 hyperactivation (CMV-CDK4^R24C^) constitutively expresses a mutant CDK4 protein that restores β-cell mass and function during diabetes development (Martin *et al*, 2003; Miyawaki *et al*, 2008; Rane *et al*, 1999). As previously observed (Rane *et al*., 1999), α-to-β-cell ratio was decreased in *CMV-Cdk4^R24C^* pancreas (Supplementary figures S1B, S1C and S1G). We then generated compound mutant mouse models with both *E2f1* deficiency and overactive CDK4 (*E2f1-/-::CMV-Cdk4^R24C^*). Interestingly, the increased β-cell number observed in *CMV-Cdk4^R24C^* pancreas was blunted in *E2f1-/-::CMV-Cdk4^R24C^* mice, with a concomitant increase of glucagon immunofluorescent staining (Supplementary figures S1D and S1G). We replicated these data in the second model using *Cdkn2a-*deficient mice (*Cdkn2a*-/-;(Serrano *et al*., 1996)), an upstream regulator of the E2F1-CDK4-pRb signaling pathway involved in β-cell function (Helman *et al*, 2016; Ndiaye *et al*, 2017; Pal *et al*, 2016). Although Glu+ and Ins+ immunofluorescent positive cell staining was conserved between *Cdkn2a*+/+ and *Cdkn2a*-/- pancreas (Supplementary figures S1E and S1G), *E2f1*-/-::*Cdkn2a*-/- compound mutant mice displayed an altered α-to-β-cell ratio with a decrease of Ins+ and an increase Glu+ cell numbers, respectively (Supplementary figures S1F and S1G). Altogether, these results suggest a specific role of E2f1 in maintaining pancreatic β cell numbers under normal conditions but also in a diabetic environment associated to glucose intolerance, insulin resistance and obesity.

### β-cell specific *E2f1* deficiency impairs glucose tolerance, insulin secretion and alters α-to-β cell ratio

To determine whether *E2f1* regulates insulin-producing β-cell fate and function in a cell-autonomous manner, we generated β-cell specific *E2f1* deficient mice by crossing *E2f1* floxed (*E2f1^fl/fl^)* with RIP-Cre (Rip-Cre/+) mice (Herrera, 2000). Quantitative RT-PCR showed a 91 % tissue-specific reduction in *E2f1* expression in pancreatic islets isolated from *E2f1*^β-/-^ mice (Supplementary figure S2A). β-cell specific deletion of *E2f1* was also confirmed at the protein level by immunofluorescence analysis (Supplementary figure S2B). *E2f1*^β^ mice had normal body weight (Supplementary figure S2C) and fasting glycemia when fed a chow diet (Supplementary figure S2D). When challenged with a bolus of glucose, *E2f1*^β^ mice exhibit glucose intolerance (Figures 2A-B), primarily due to decreased insulin secretion in response to glucose (Figure 2C) rather than defective insulin sensitivity (Figure 2D). GSIS experiments on mouse pancreatic islets isolated from *E2f1^fl/fl^* and *E2f1*^β-/-^ further confirmed insulin secretion defects in response to glucose (Figure 2E). Interestingly, an immunofluorescence analysis of sections from control and *E2f1*^β-/-^ pancreas demonstrated increased glucagon-positive cells and an increased α-to-β cell ratio in β-cell specific *E2f1* KO islets (Figures 2F-G). The number of cells per islet section remained unchanged between control and *E2f1*^β-/-^pancreas (Supplementary figure S2E). As observed in germline *E2f1* deficient mice (Figure 1), these data suggest that β-cell expression of *E2f1* is necessary to maintain insulin secretion and glucose homeostasis associated to normal α-to-β ell ratio.

**Figure 2.**
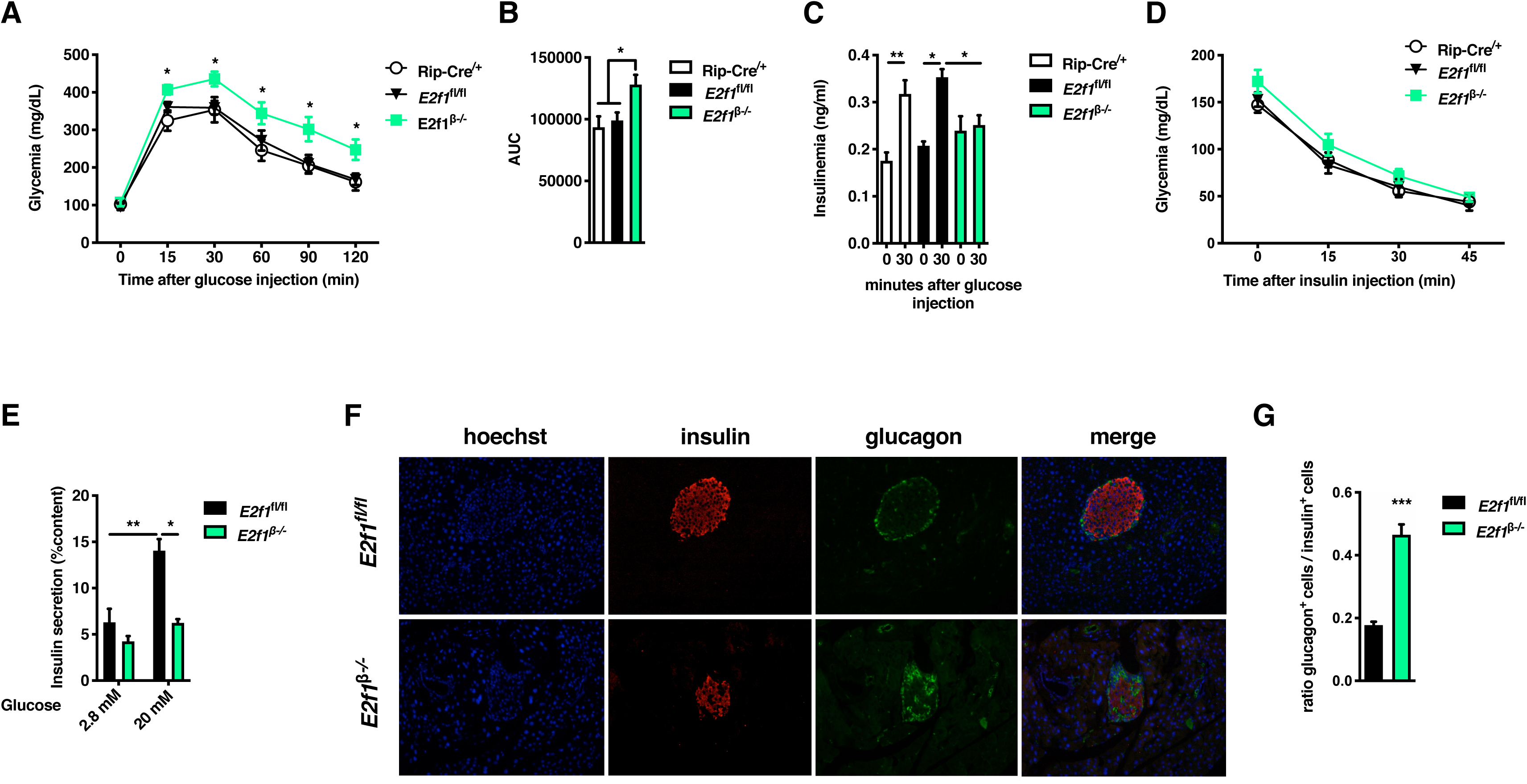
The β cell-specific deletion of *E2f1* impairs glucose tolerance and insulin secretion in mice. **(A)** Intraperitoneal glucose tolerance test (IPGTT) was performed on *E2f1*^β-/-^ (n=12), RIP-Cre^/+^ (n=7) and *E2f1*^fl/fl^ (n=8) male mice at 12 weeks of age. **(B)** Area under the curve (AUC) calculated from (B). **(C)** Plasma insulin levels at 0 and 30 min after intraperitoneal glucose injection in *E2f1*^β-/-^ (n=18), RIP-Cre^/+^ (n=15) and *E2f1*^fl/fl^ (n=6) male mice at 12 weeks of age. **(D)** Intraperitoneal insulin tolerance test (IPITT) of *E2f1*^β-/-^ (n=8), RIP-Cre^/+^ (n=8) and *E2f1*^fl/fl^ (n=8) male mice at 12 weeks of age. **(E)** Glucose-stimulated insulin secretion (GSIS) at indicated glucose concentrations on islets isolated from *E2f1*^β-/-^ (n=3) and control *E2f1*^fl/fl^ (n=5) male mice. **(F)** Representative immunofluorescent staining of insulin and glucagon in pancreatic sections from 3 months old *E2f1*^β-/-^ and control *E2f1*^fl/fl^ male mice. **(G)** Ratio of glucagon labelled cells (Glucagon +) over insulin labelled cells (Insulin +) in *E2f1*^β-/-^ and *E2f1*^fl/fl^ male mice calculated from (G). All values are expressed as mean ± s.e.m. and were analysed by one-way analysis of variance (ANOVA) with Tukey’s test (A, C, D), two-way ANOVA with Tukey’s test (B, E, F) or two-tailed unpaired *t*-test (H). *p < 0.05; **p<0.01; ***p<0.001.

### Loss of β cell identity markers in *E2f1* deficient pancreatic islets

E2F1 is a transcription factor that controls gene expression in several cellular systems. In order to get a global view of the transcriptional mechanisms associated with the loss of *E2f1* expression in the β cell, we performed RNA sequencing (RNA-seq) in control (*E2f1^fl/fl^*) and *E2f1*^β-/-^isolated islets. As expected, the floxed region of the *E2f1* gene spanning exon 2 and 3 was not covered in *E2f1*^β-/-^ isolated islets compared to *E2f1^fl/fl^* isolated islets, indicating specific and efficient gene deletion through the *Cre* recombinase activity in *E2f1*^β-/-^ isolated islets (Supplementary figure S3A). The analysis of the transcripts revealed that 692 annotated genes were differentially expressed across the two groups (adjusted P Value (AdjP) <0.05). Interestingly, a vast majority of the genes were upregulated in *E2f1*^β-/-^ isolated islets (493 genes, Figure 3A and Supplementary Table S4), with only 199 down-regulated genes associated with the loss of *E2f1* expression in β cells (Figure 3A and Supplementary Table S4). This first observation suggests that *E2f1* expression is not only necessary to activate, but also to repress gene transcription within the β cell. Analysis of the RNA-seq data of downregulated genes with metascape software revealed an enrichment of gene networks involved in signal release, regulation of exocytosis and negative regulation of secretion (Figure 3B). Conversely, upregulated genes in *E2f1*^β-/-^ isolated islets were associated with the regulation of cell adhesion, inflammatory response and cytokine production (Figure 3C). To better understand the relationship between the observed metabolic phenotype (*i.e.* glucose intolerance and defective insulin secretion), increased glu+ cells, decreased ins+ cells and *E2f1* deficiency, we filtered gene sets to focus on β- and α-cell genes being conserved between zebrafish, mouse and human (Tarifeno-Saldivia *et al*, 2017). Notably, a total of 15 genes from 109 conserved genes (82 and 27 for α-cell and β-cell genes, respectively) were differentially expressed. Interestingly, most of the conserved β-cell genes were found to be decreased in *E2f1*^β-/-^ isolated islets (Figure 3D and Supplementary figure S3B), whereas most of the conserved α-cell genes were upregulated upon *E2f1* deficiency (Figure 3D and Supplementary figure S3C). Gene Set Enrichment Analysis (GSEA (Subramanian *et al*, 2005)) further confirmed an enriched signature of decreased expression of β-cell markers (Figure 3E) and conversely, an increased expression of α-cell markers in *E2f1*^β^ isolated islets (Figure 3F). We further confirmed these changes in the expression of selected α-β-cell markers at the transcriptional level by qPCR in an independent experiment: indeed we observed a strong decrease in transcript levels of *Pdx1, Mafa (p=0.13), Ins2*, *Pcsk9*, *Foxo1 and Glp1r* in *E2f1*^β-/-^ islets (Supplementary figure S3D). Conversely, α-cell specific Aristaless-related homeobox (*Arx)* mRNA levels were increased in *E2f1*^β-/-^ islets (Supplementary figure S3E). Altogether these results suggest that the β-cell specific deletion of *E2f1* induces a transcriptional reprogramming characterized by a loss of β-cell identity genes associated to an increased expression of non β-cells markers.

**Figure 3.**
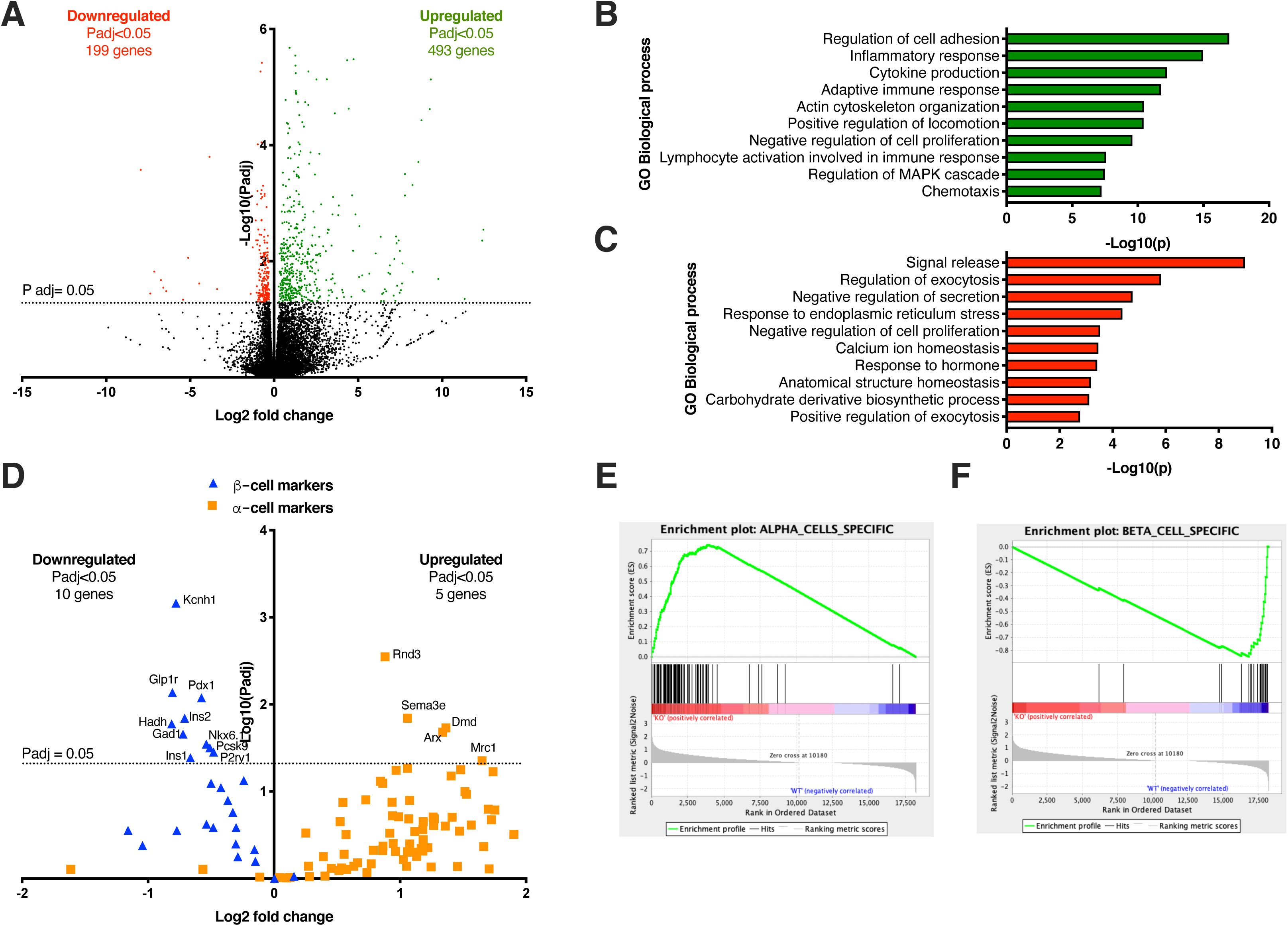
RNA-seq analysis of *E2f1* β cell-specific knockout islets reveal altered transcriptional programs. **(A)** Volcano plot providing adjusted P value (*i.e.*, FDR, false discovery rate) and fold change for all gene transcripts in islets from *E2f1*^β-/-^ mice. Genes that are differentially expressed compared to *E2f1*^fl/fl^ with an AdjP<0.05 are indicated by two-level color coding. 199 downregulated protein-coding genes are highlighted in red. 493 upregulated protein-coding genes are highlighted in green (n=3 per genotype for RNA-seq analysis). **(B-C)** Metascape enrichment analysis of downregulated (B) and upregulated (C) genes in *E2f1*^β-/-^ compared to *E2f1*^fl/fl^. Histogram of enriched terms across input gene lists are shown. **(D)** Evolutionary conserved α- and β-cell markers were recovered from Tarifeno-Saldivia E *et al*. (Tarifeno-Saldivia *et al*., 2017) and were used to filter our RNA-seq dataset. Volcano plots provides AdjP values (*i.e.*, FDR, false discovery rate) and fold change for both α- and β-cell marker transcripts in islets from *E2f1*^β-/-^ mice. The α- and β-cell markers are displayed as blue triangle and red square, respectively. Gene symbols of differentially expressed α- and β-cell markers are displayed (AdjP<0.05). **(E-F)** Enrichment plot from Gene Set Enrichment Analysis (GSEA) was conducted with 82 probe sets specifically expressed in α-cells (E) and with 26 probe sets specifically expressed in β-cells (F).

### The treatment with the E2F inhibitor *HLM006474* inhibits glucose-stimulated insulin secretion and impairs β and α-cell gene expression in Min6 cells and human islets

To assess the effect of E2F1 activity on β-cell identity markers in human islets, we made use of the E2F pan-inhibitor HLM006474 previously shown to inhibit the binding of E2Fs to their DNA target genes (Sangwan *et al*, 2012) and E2F1 transcriptional activity (Rosales-Hurtado *et al*., 2019). Consistent with our previous findings in HEK293 cells (Rosales-Hurtado *et al*., 2019), a 48h treatment of Min6 cells with this inhibitor triggered a decrease in E2f1 transcriptional activity, as measured by transient transfection experiments using an E2F reporter gene (Figure 4A). In addition, the treatment of Min6 cells with this inhibitor induced a marked decrease in glucose-stimulated insulin secretion (Figure 4B) and in the expression of several β-cell markers, including *Ins1*, *Pdx1, Pax4* and *Nkx2.2* (Figure 4C). The treatment of human islets with the E2F inhibitor for 48 hours also decreased GSIS (Figure 4D) and β-cell marker expression levels with a concomitant increase in the expression of α-cell genes (Figure 4E). Therefore, the pharmacological inhibition of E2F activity impairs β ell function and gene expression in both mouse cell line and human islets, suggesting that E2F1 activity is also required in human islets to maintain proper insulin secretion and β ell identity genes, as observed in mice.

**Figure 4.**
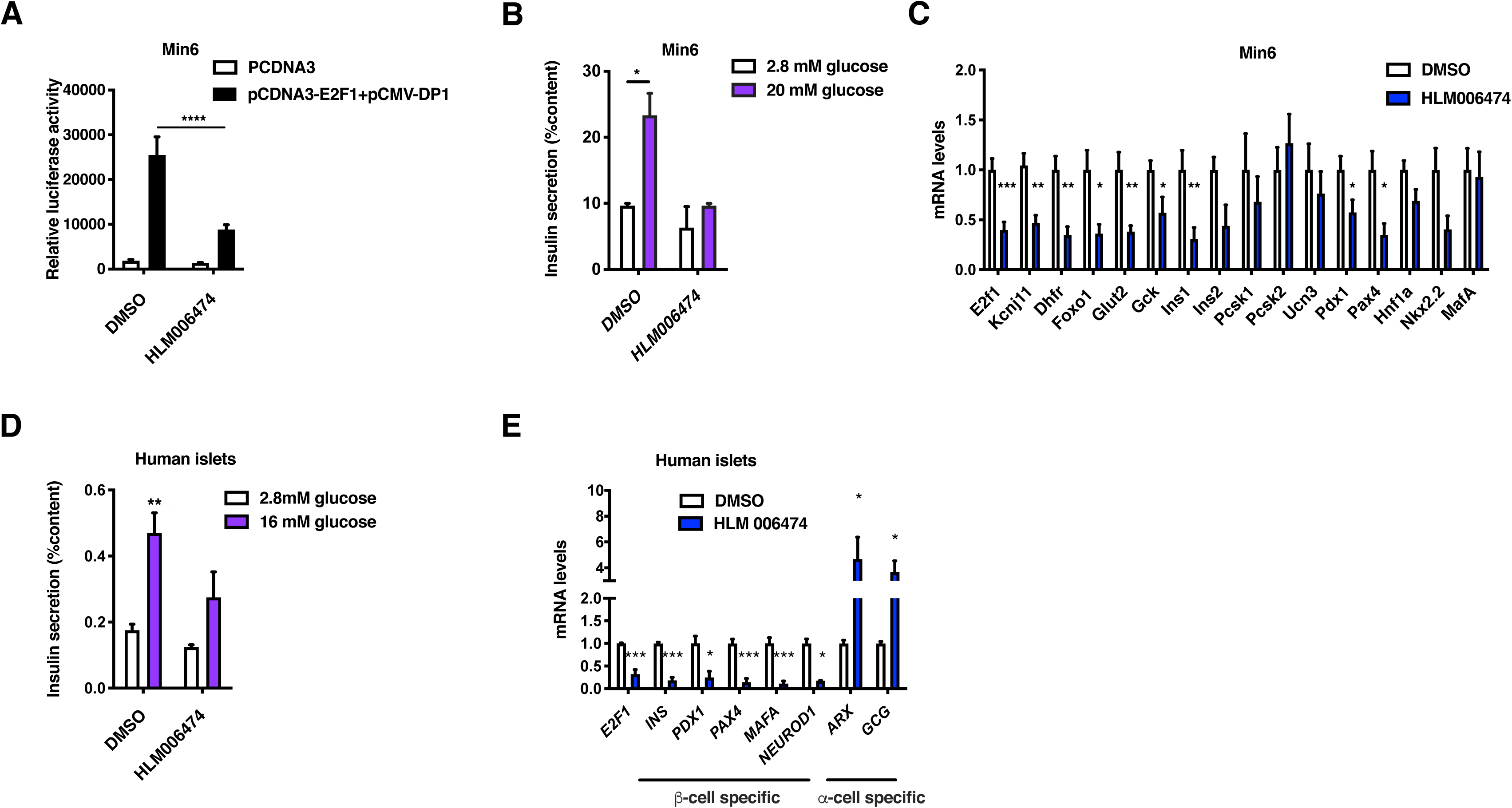
The pharmacological inhibition of E2f transcription factor activity induces a loss of β-cell function in mouse and human islets. **(A)** Min6 cells were transiently co-transfected with the E2F-RE-Tk promoter luciferase construct in the absence (PCDNA3) or presence of E2F1:DP1 heterodimer (E2F1/DP1) and were subsequently treated with DMSO (0.1 %, 48h) or HLM006474-treated (10 µM, 48h). Results were normalized to β- galactosidase activity. **(B)** Glucose stimulated insulin secretion (GSIS) at indicated glucose concentration on DMSO (0.1 %, 48h) or HLM006474-treated (10 µM, 48h) Min6 cells (n=6). **(C)** qPCR-based analysis of β-cell specific mRNA expression in DMSO (0.1 %, 48h) or HLM006474-treated (10 µM, 48h) Min6 cells (n=3). **(D)** Glucose stimulated insulin secretion (GSIS) at indicated glucose concentration on human islets treated with DMSO (0.1 %, 48h) or HLM006474 (10 µM, 48h) (n=3) (see supplementary Table S1 for donor information). **(E)** qPCR-based analysis of β-cell specific (*INS, PDX1, PAX4, MAFA* and *NEUROD1*) and α-cell specific (*ARX* and *GCG*) mRNA expression in DMSO (0.1 %, 48h) or HLM006474-treated (10 µM, 48h) human islets (n=3). Data are represented as mean ± s.e.m. and were analyzed by two-way ANOVA with Tukey’s test (A, B, D) and two-tailed unpaired *t*-test (C, E). *p < 0.05; **p<0.01; ***p<0.001.

### Maintenance of β-cell identity is dependent on histone deacetylase and E2F1 activities

Our transcriptome analysis revealed a 2.5-fold more up- than downregulation of global gene expression in *E2f1*^β-/-^ isolated islets. Comparative analysis of log10 transcription levels (Log10(TPM+1)) of up- and downregulated genes values demonstrated that the expression of upregulated genes in *E2f1*^β-/-^ islets was significantly increased upon *E2f1* deficiency compared to wild-type controls whereas the expression of downregulated genes was not significantly modulated (Figure 5A). Considering that E2f1 could play a dual role in the regulation of gene expression in pancreatic β cells, we postulated that this mechanism could be related to a distinct epigenomic profile within promoter of genes that are up- and down-regulated in *E2f1*^β-/-^ islets. Using a recently published chromatin-state segmentation model (Lu *et al*., 2018), we probed the activation/repressive level of these promoters by monitoring several epigenome marks such as active and poised promoters (tri-methylation of lysine 4 in histone H3 [H3K4me3]) and enhancers (acetylation of lysine 27 in histone H3 [H3K27ac]). Intersecting publicly available data of chromatin immunoprecipitation followed by next-generation sequencing (ChIP-seq) from healthy C57Bl6/J mouse pancreatic islets (GSE 110648 (Lu *et al*., 2018)) and our RNA-seq data, we grouped up- and down regulated genes according to their chromatin state (Figure 5B and Supplementary Table S4). 61% of the upregulated genes in *E2f1*^β-/-^ pancreatic islets were associated to a silent chromatin state in healthy C57Bl6/J mouse pancreatic islets characterized by bivalent H3K4me3/H3K27me3 and Polycomb-repressed (H3K27me3) marks. Conversely, 82% of the down-regulated genes showed an enrichment in active chromatin state characterized by RNA-Pol2 recruitment, H3K4me3 and H3K27ac histone marks.

**Figure 5.**
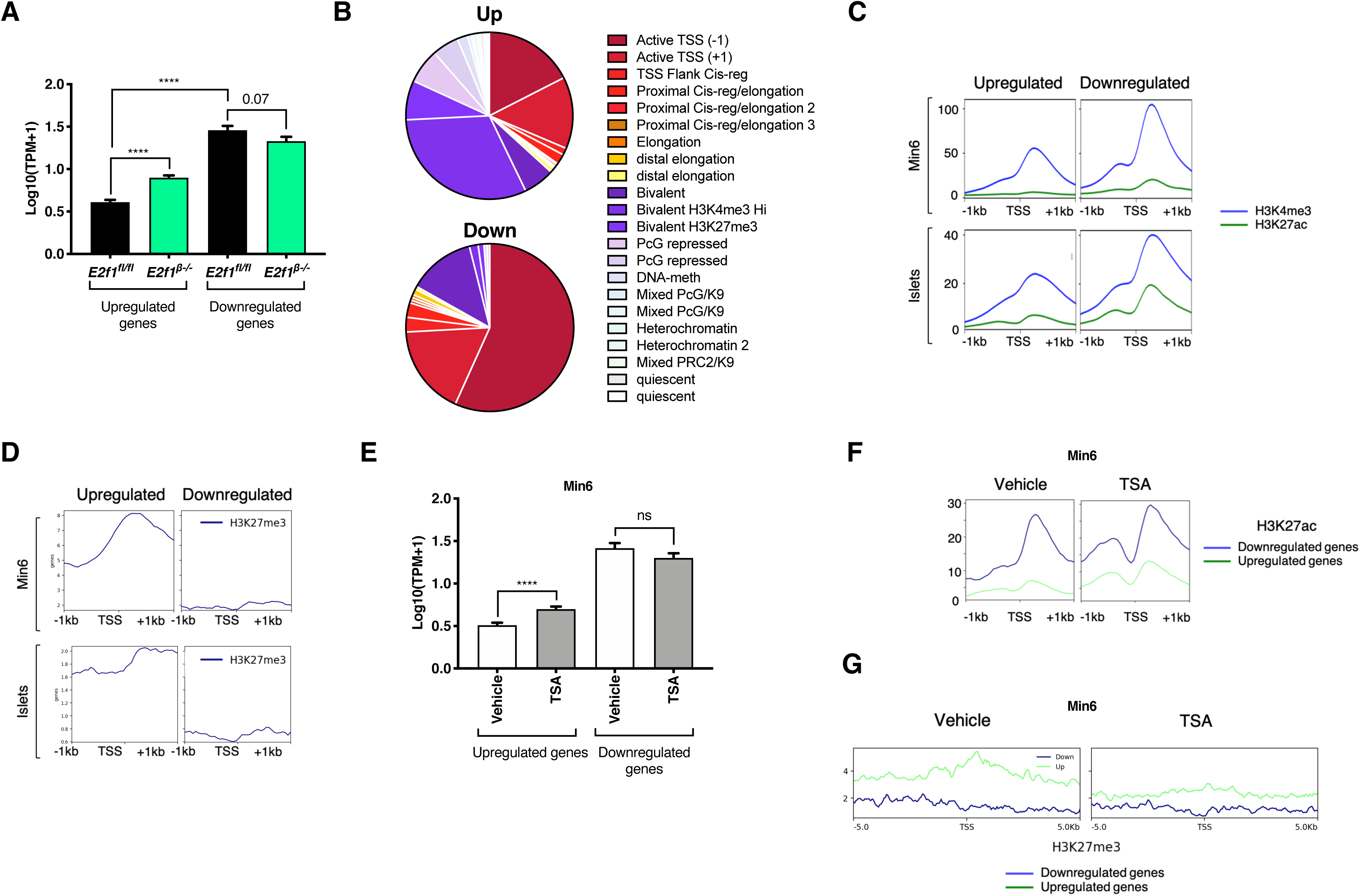
The transcriptional repression of *E2f1* β-cell target genes occurs through an HDAC-dependent mechanism. **(A)** Expression level of *E2f1*^β-/-^ up- and down- regulated genes both in pancreatic islets from control (WT) and *E2f1* knock-out (*E2f1*^β-/-^) mice. Results are displayed as Log10(TPM+1) calculated from TPM obtained from RNA-seq data. Data are represented as mean s.e.m and were analyzed by unpaired *t*-test (**** p<0.0001). **(B)** Pie-chart displaying chromatin state of *E2f1*^β-/-^ up- and down-regulated genes according to pancreatic islets genes-associated chromatin state stratification from Lu TT *et al*. (Lu *et al*., 2018) **(C)** H3K4me3 and H3K27ac ChIP-seq mean signal within promoter (centered to TSS +/- 1kb) of *E2f1*^β-/-^up- and down-regulated genes both in Min6 cells and mouse pancreatic islets (Lu *et al*., 2018). **(D)** H3K27me3 ChIP-seq mean signal within promoter (centered to TSS +/- 1kb) of *E2f1*^β-/-^up- and down-regulated genes both in Min6 cells and mouse pancreatic islets (Lu *et al*., 2018). **(E)** Expression level of *E2f1*^β-/-^ up- and down-regulated genes in TSA-treated Min6 cells. Results are displayed as Log10(TPM+1) calculated from TPM obtained from RNA-seq data. Data are represented as mean ± s.e.m and were analyzed by unpaired *t*-test (**** p<0.0001). **(F)** H3K27ac ChIP-seq mean signal within promoter (centered to TSS +/- 1kb) of *E2f1*^β-/-^up- and down-regulated genes in TSA-treated Min6 cells. **(G)** H3K27me3 ChIP-seq mean signal within promoter (centered to TSS +/- 1kb) of *E2f1*^β-/-^up- and down-regulated genes in TSA-treated Min6 cells.

While H3K4me3, H3K27ac and H3K27me3 ChIP-seq data were available for mouse islets, we then performed ChIP-seq experiments in Min6 cells as a surrogate of β cell. H3K4me3, H3K27ac and H3K27me3 ChIP-seq signals were thus interrogated within promoter (centered to transcription start site [TSS] +/- 1 kb) of up- and downregulated genes both in Min6 cells and mouse pancreatic islets (Figures 5C-D and Supplementary figures 4A-B). H3K4me3 as well as H3K27ac signals were stronger within promoter of genes that are downregulated in *E2f1*^β-/-^ isolated islets compared to upregulated genes, both in Min6 cells and mouse pancreatic islets (Figure 5C and Supplementary figures 4A-B). Conversely, H3K27me3 ChIP-seq signals were lower in the promoter region of downregulated genes compared to upregulated genes (Figure 5D and Supplementary figures 4A-B). We then analyzed our integrated RNA-seq/ChIP-seq data using the “upstream regulator analysis” function of Ingenuity Pathway Analysis (IPA) to identify potential contributors that could associate to the phenotype of *E2f1*^β-/-^ isolated islets, which would be related to epigenomics. Among the most significant upstream regulators of the upregulated genes, the HDAC inhibitor trichostatin A (TSA) signaling pathway was predicted to be significantly activated (Table 1), suggesting that HDAC could regulate an E2f1-dependent gene program. Consequently, having shown that (*i*) the levels of upregulated genes were significantly increased upon β-cell specific *E2f1* deletion (Figure 5A), (*ii*) H3K27ac signal was weaker within the promoter region of upregulated genes compared to downregulated genes (Figure 5C) and (*iii*) TSA may modulate the E2f1-dependent epigenome, we next hypothesized that E2f1 could repress the expression of upregulated genes through an HDAC (histone deacetylase)-dependent mechanism. To ascertain this hypothesis, Min6 cells were treated during 16h with the pan-HDAC inhibitor trichostatin A (TSA, 0.5 µM). Interestingly, via RNA-seq, we evidenced that the expression of upregulated genes in *E2f1*^β-/-^ islets was significantly increased in response to TSA treatment compared to vehicle-treated cells whereas the expression of downregulated genes was not significantly modulated (Figure 5E), as observed in *E2f1*^β-/-^ islets. These results showed that the pharmacological HDAC inhibition in Min6 cells partly mimicked the effects observed in *E2f1*^β-/-^ pancreatic islets on the expression of upregulated genes, suggesting that an E2f1/HDAC complex could contribute to regulate the expression of these genes in the β cell. Conversely, these results also indicated that E2f1-dependent downregulation of genes probably occurred through an HDAC-independent mechanism. To go further in the characterization of the molecular mechanisms linking E2f1 to HDAC in the β cell, we measured the acetylation levels of promoter (TSS +/- 1 kb) of up- and down-regulated genes in Min6 cells by probing H3K27ac level in response to TSA treatment through ChIP-seq experiments. Interestingly, TSA treatment increased H3K27ac levels within the promoter of upregulated genes compared to vehicle-treated cells, whereas H3K27ac levels were less affected within the promoter of down-regulated genes (Figure 5F). In addition, H3K27me3 ChIP-seq signals were lower in the promoter region of upregulated genes upon TSA treatment (Figure 5G), whereas no effects were observed upon TSA treatment on downregulated genes. These results were in accordance with gene expression analysis (Figure 5E) and support a role for HDAC activity in the transcriptional repression of E2f1 target genes in β cell whereas HDAC enzymes may probably not be involved in the E2f1-mediated transcriptional activation.

**Table 1.**
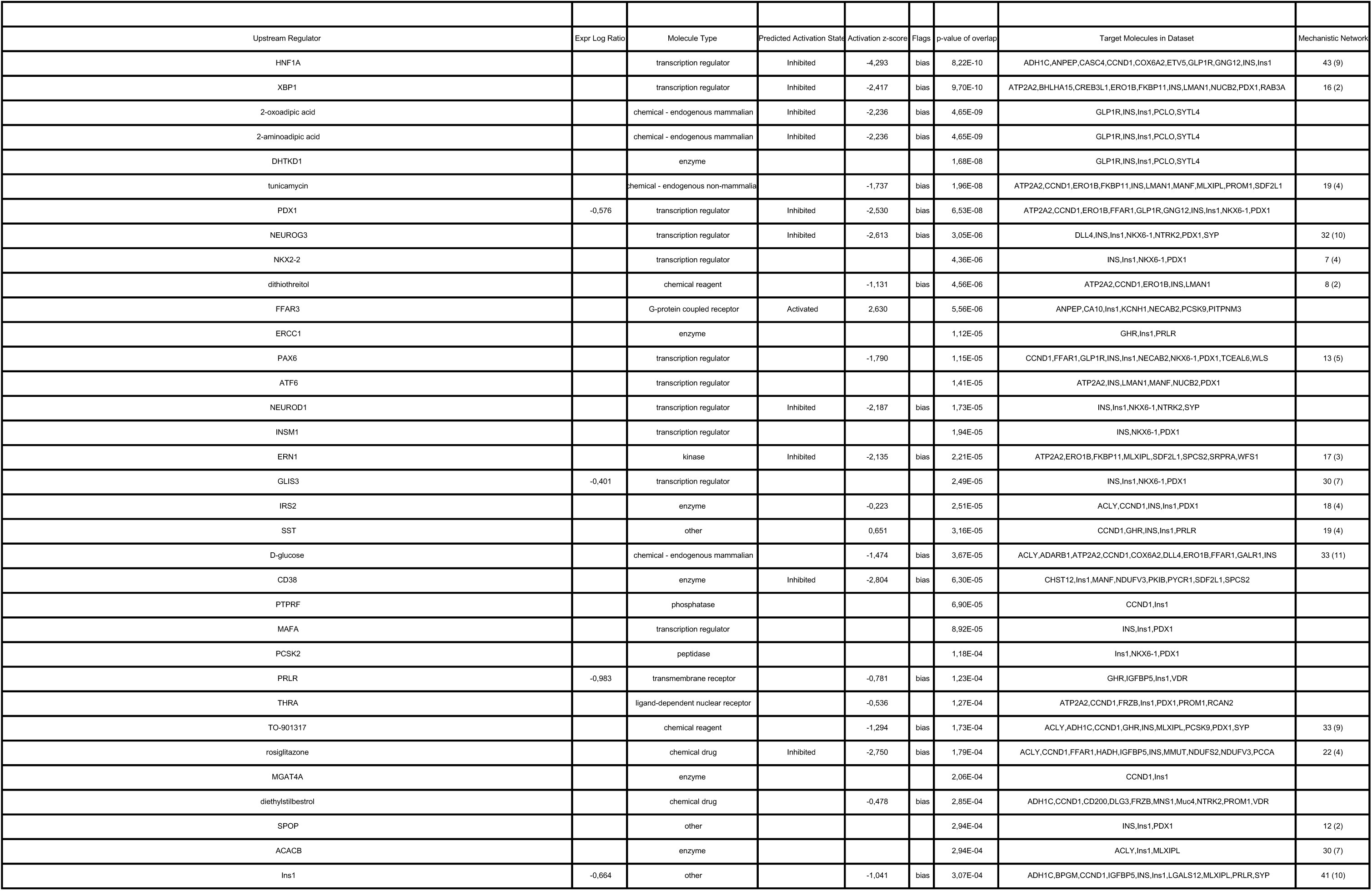
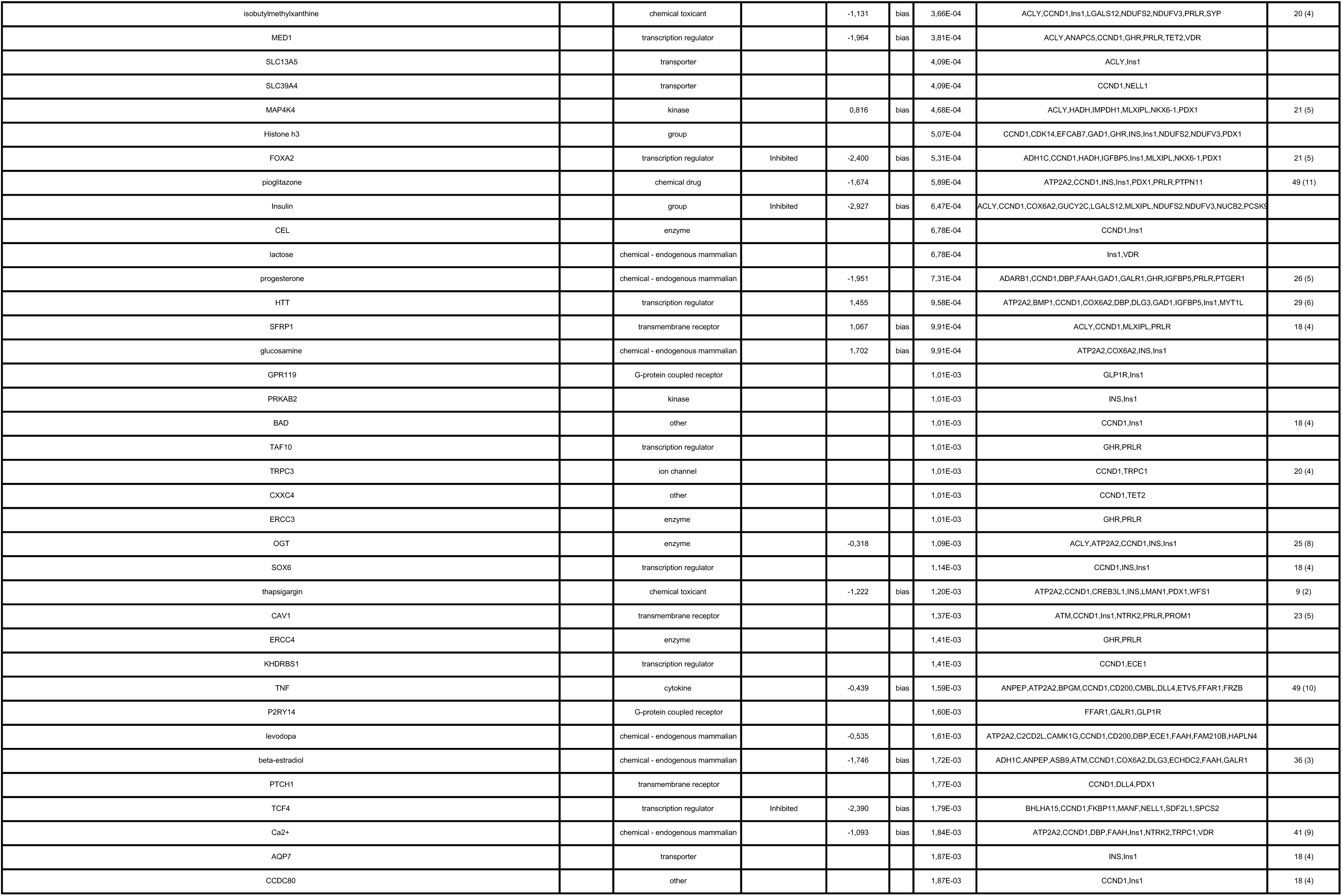
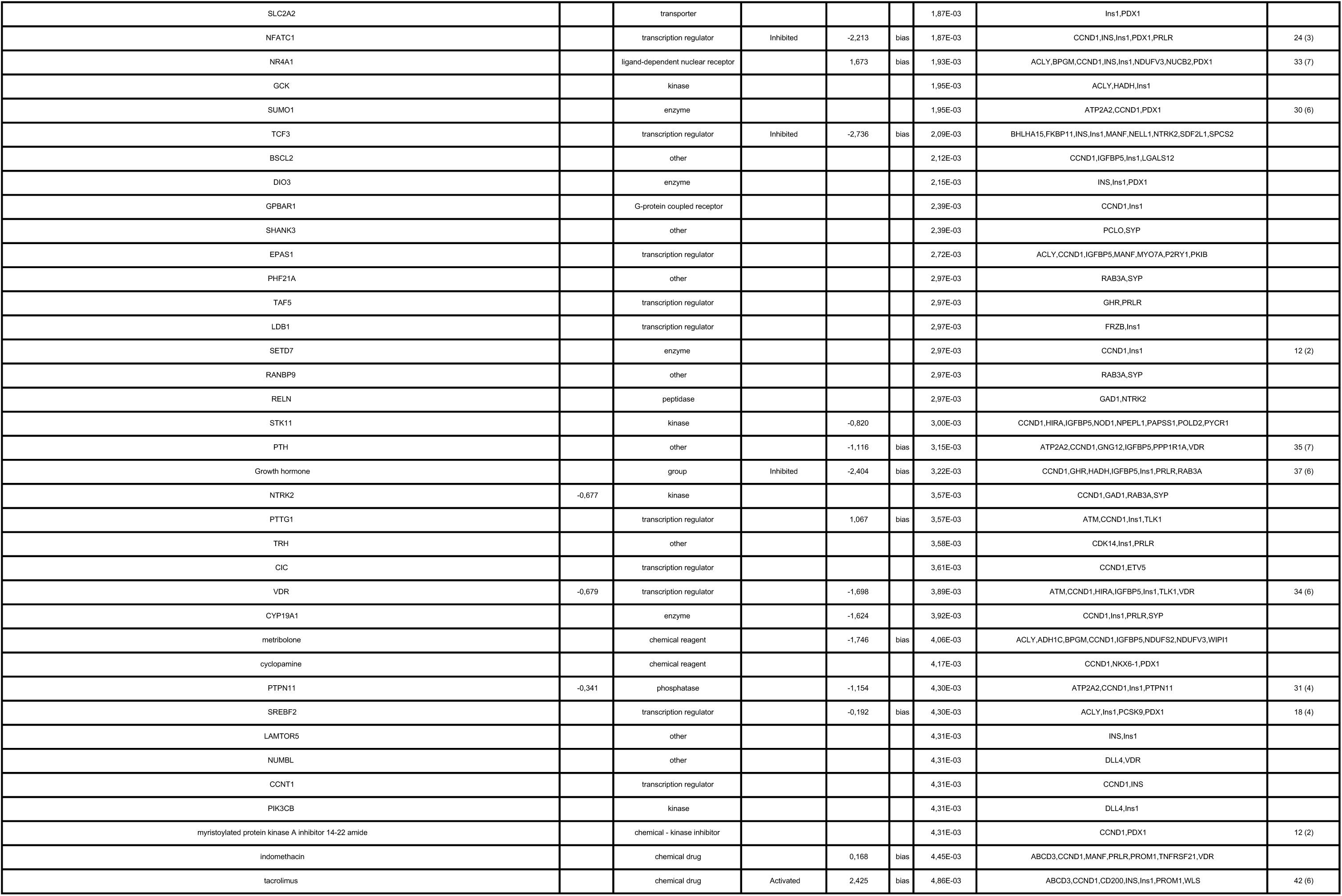
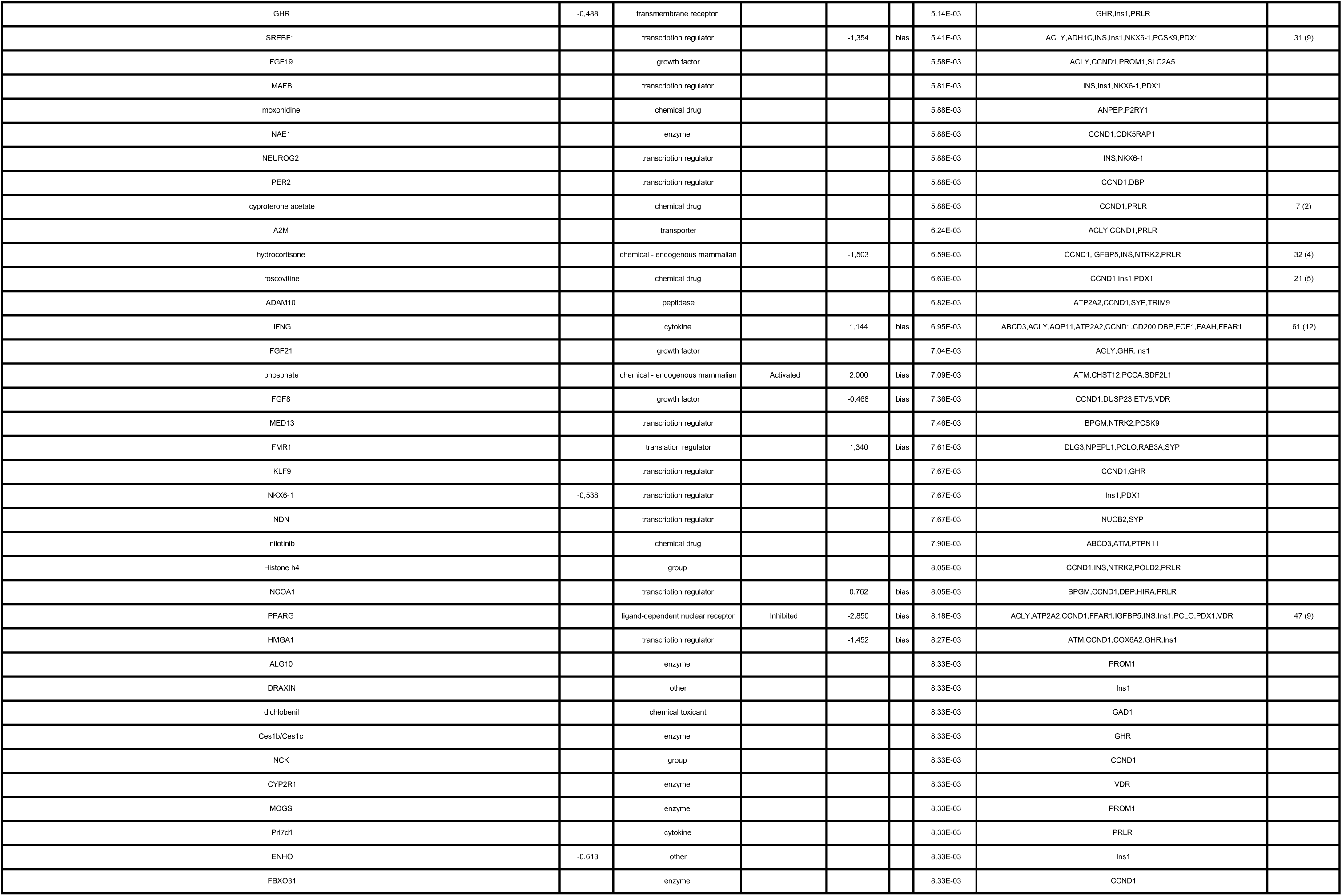
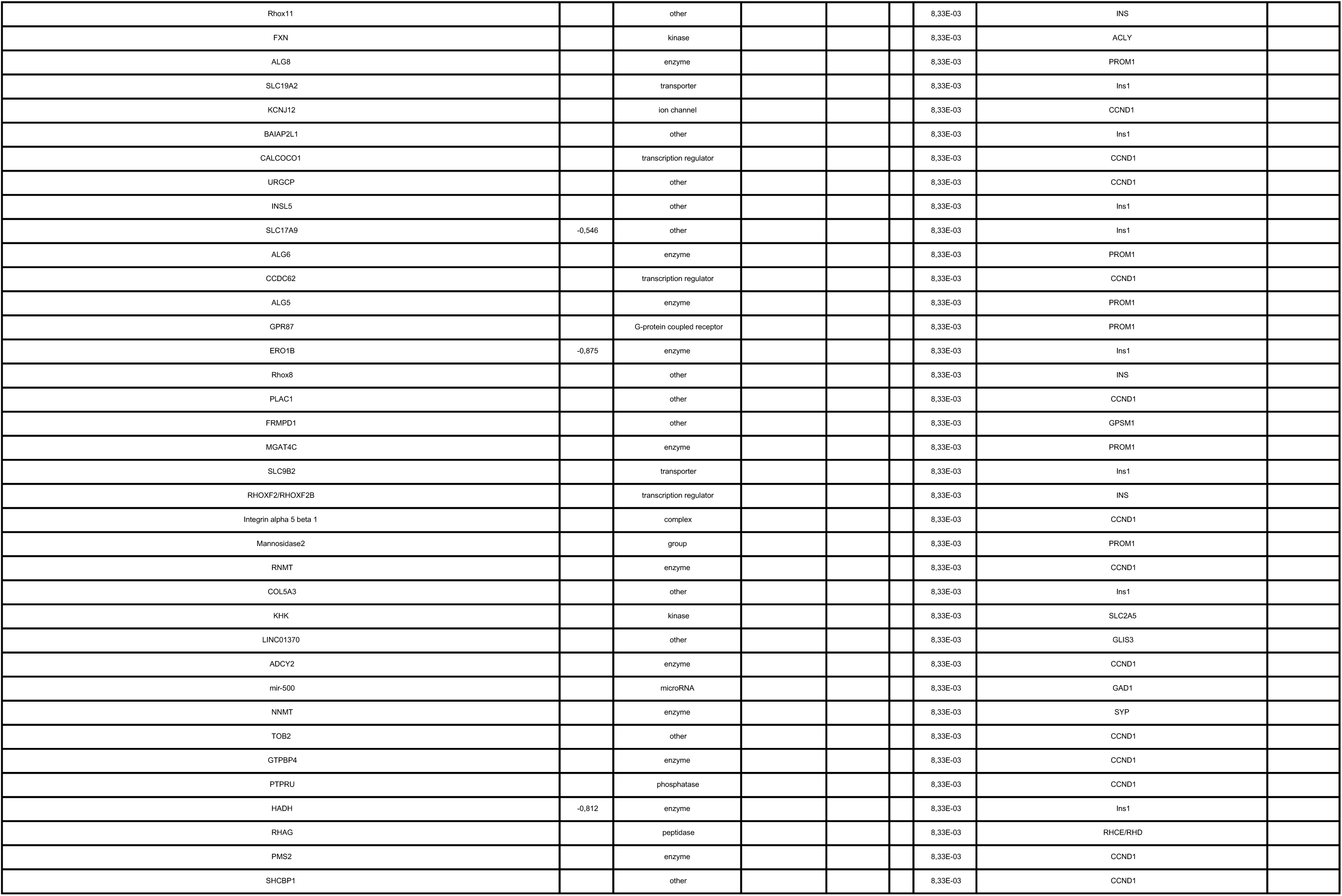
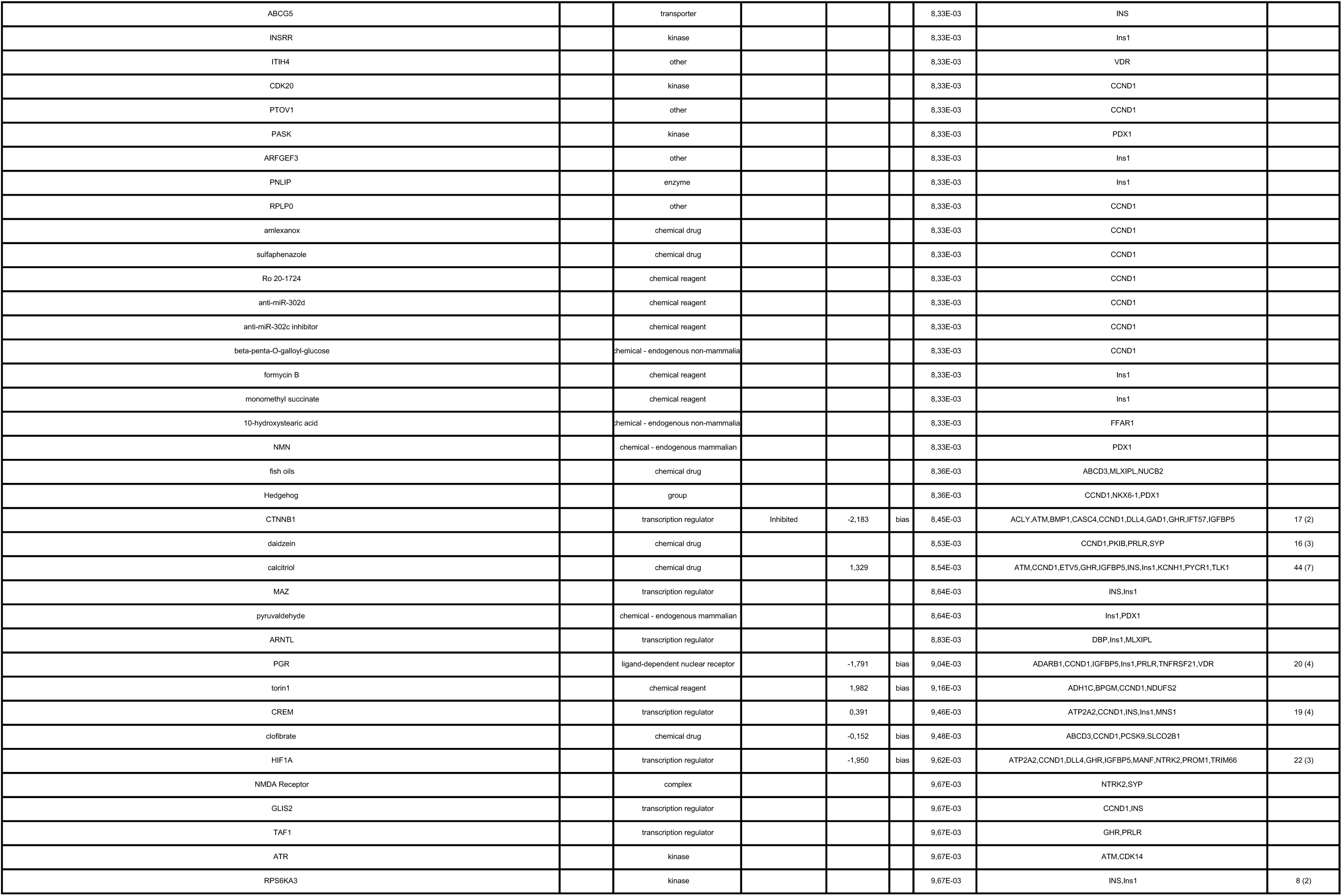
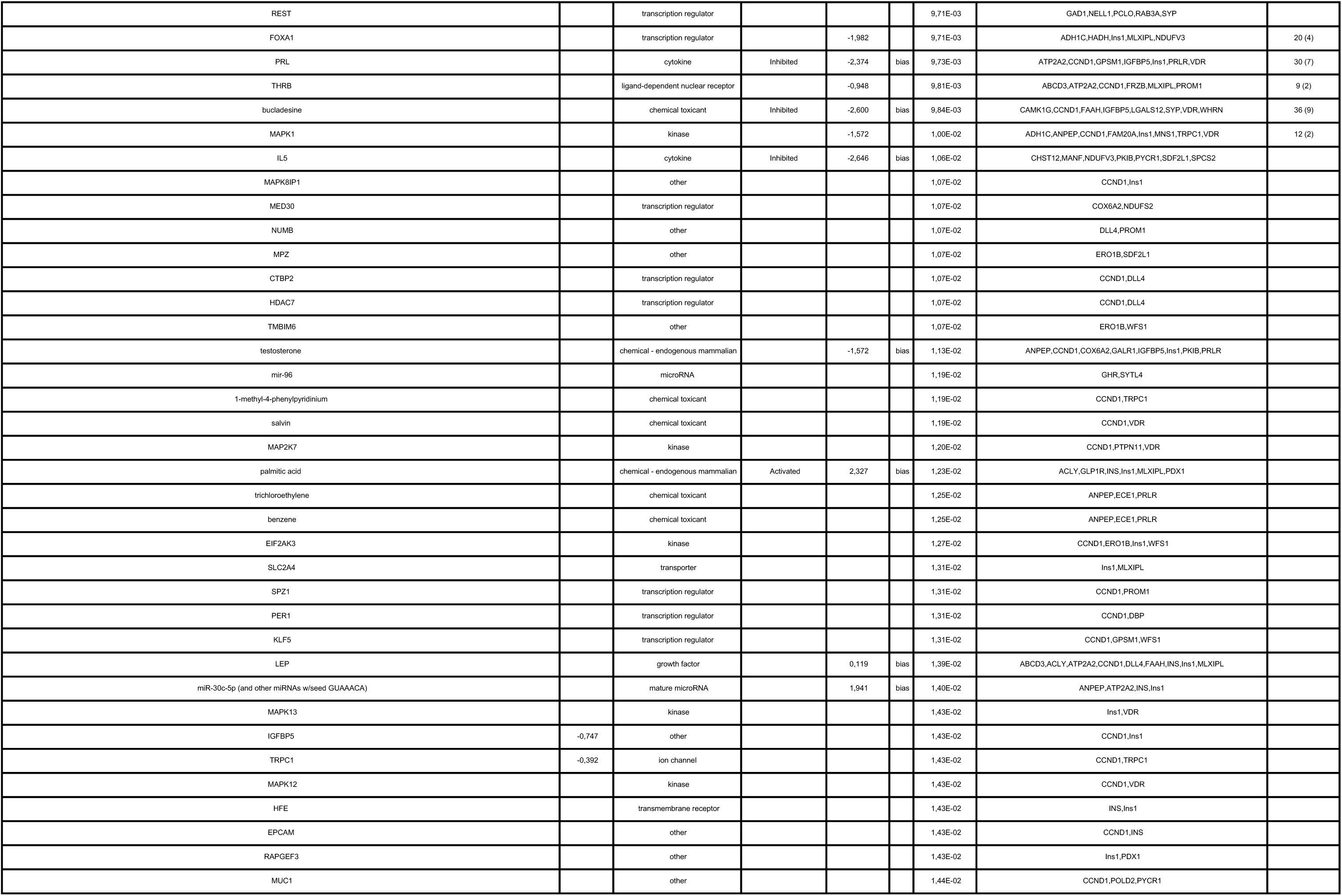
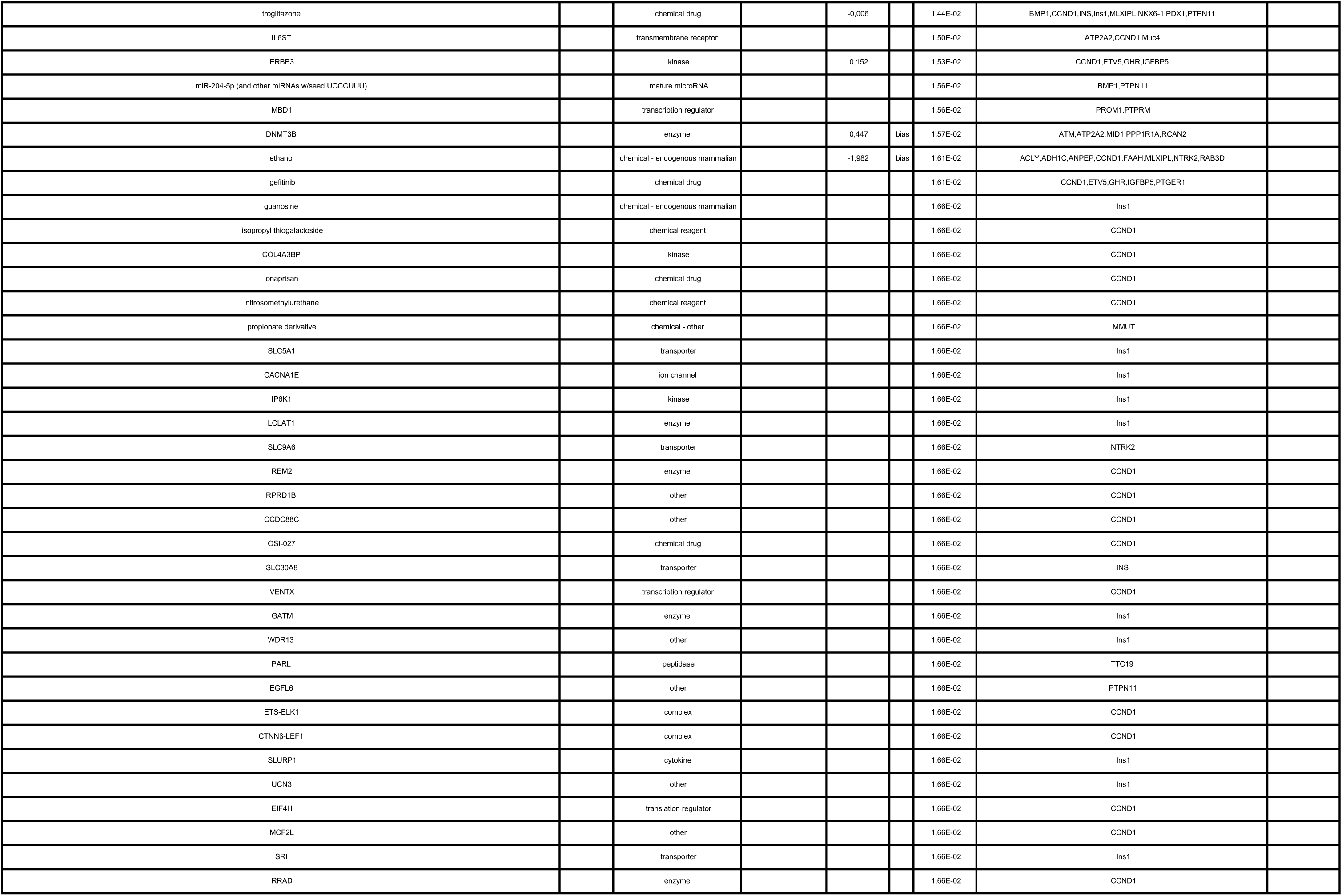
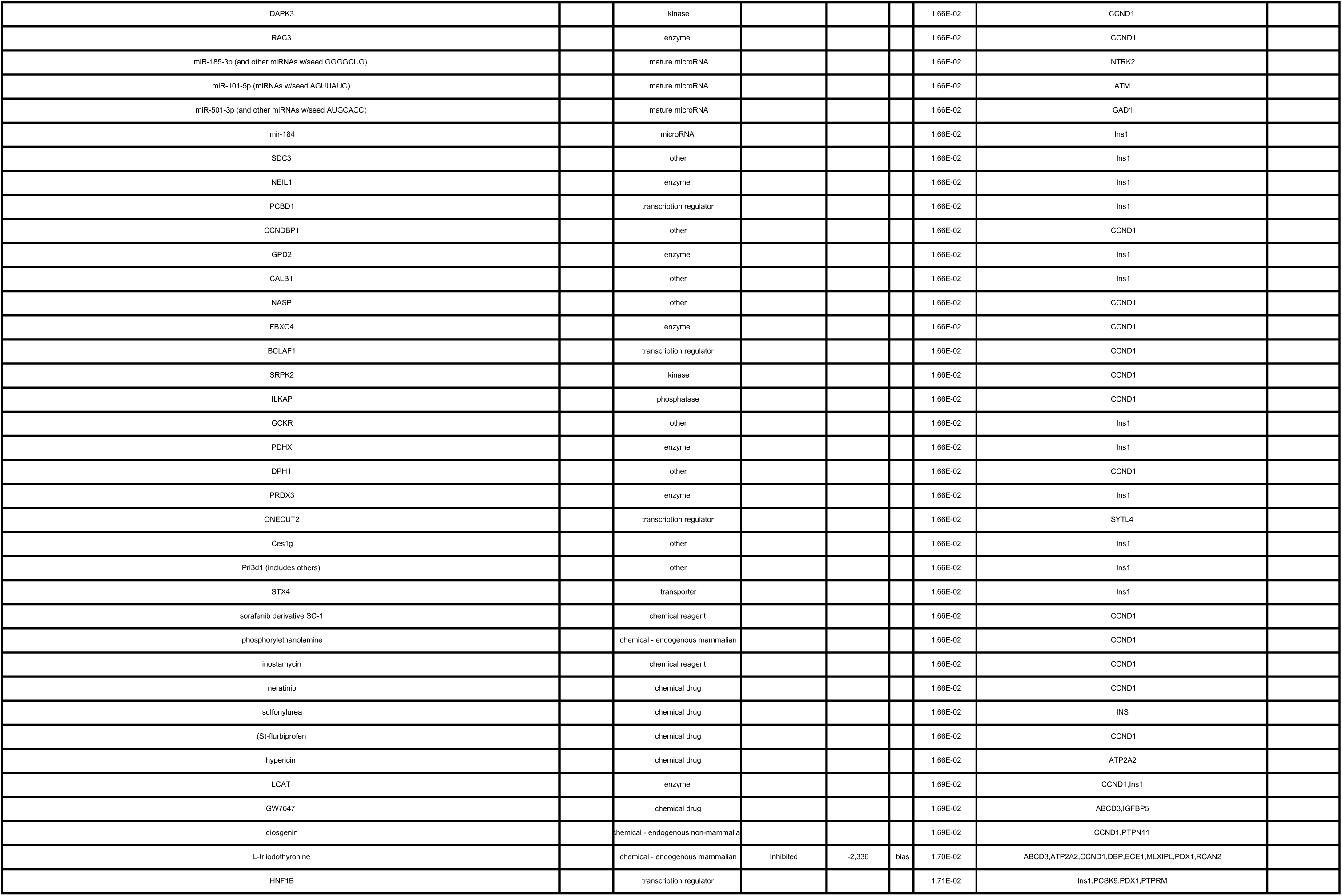
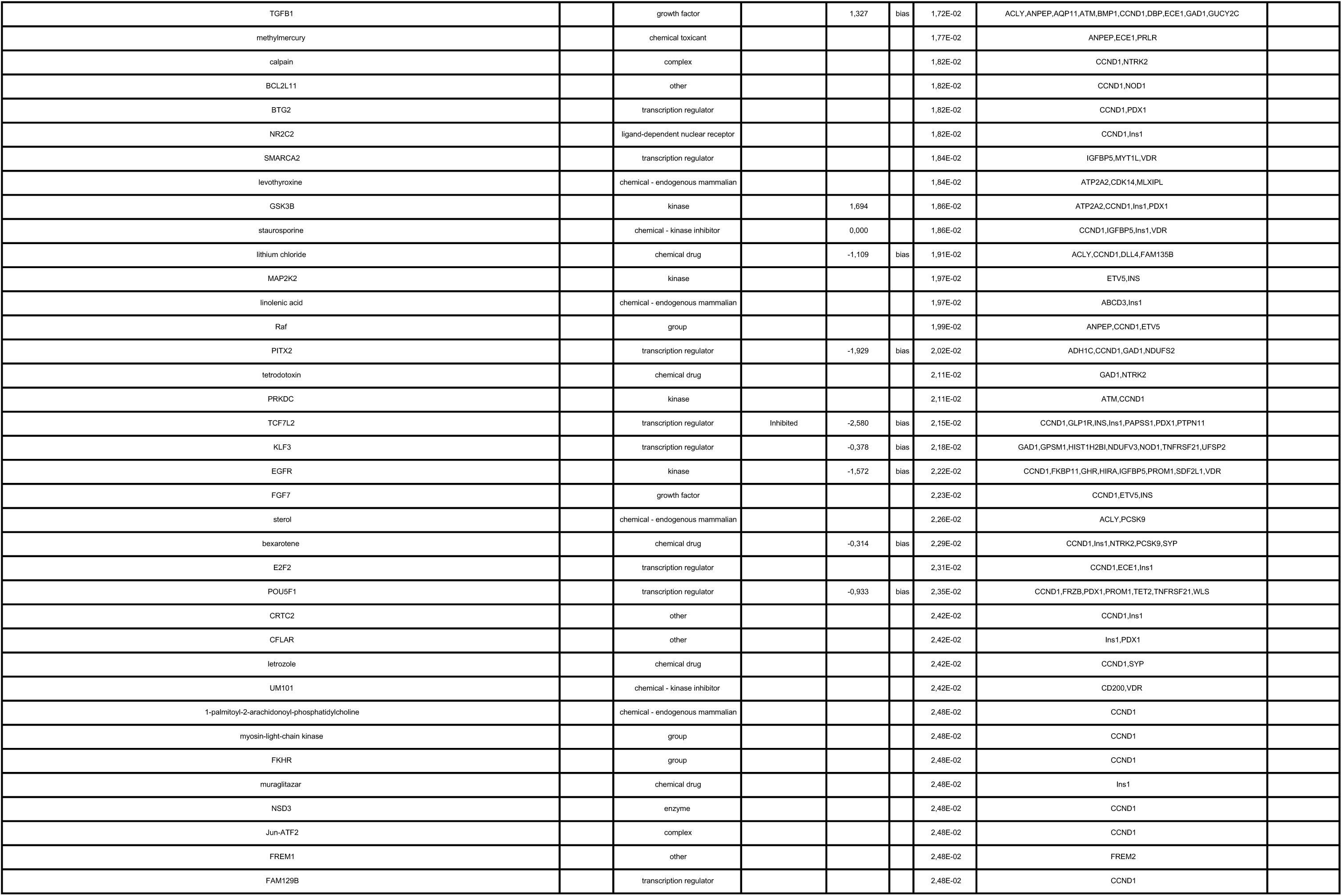
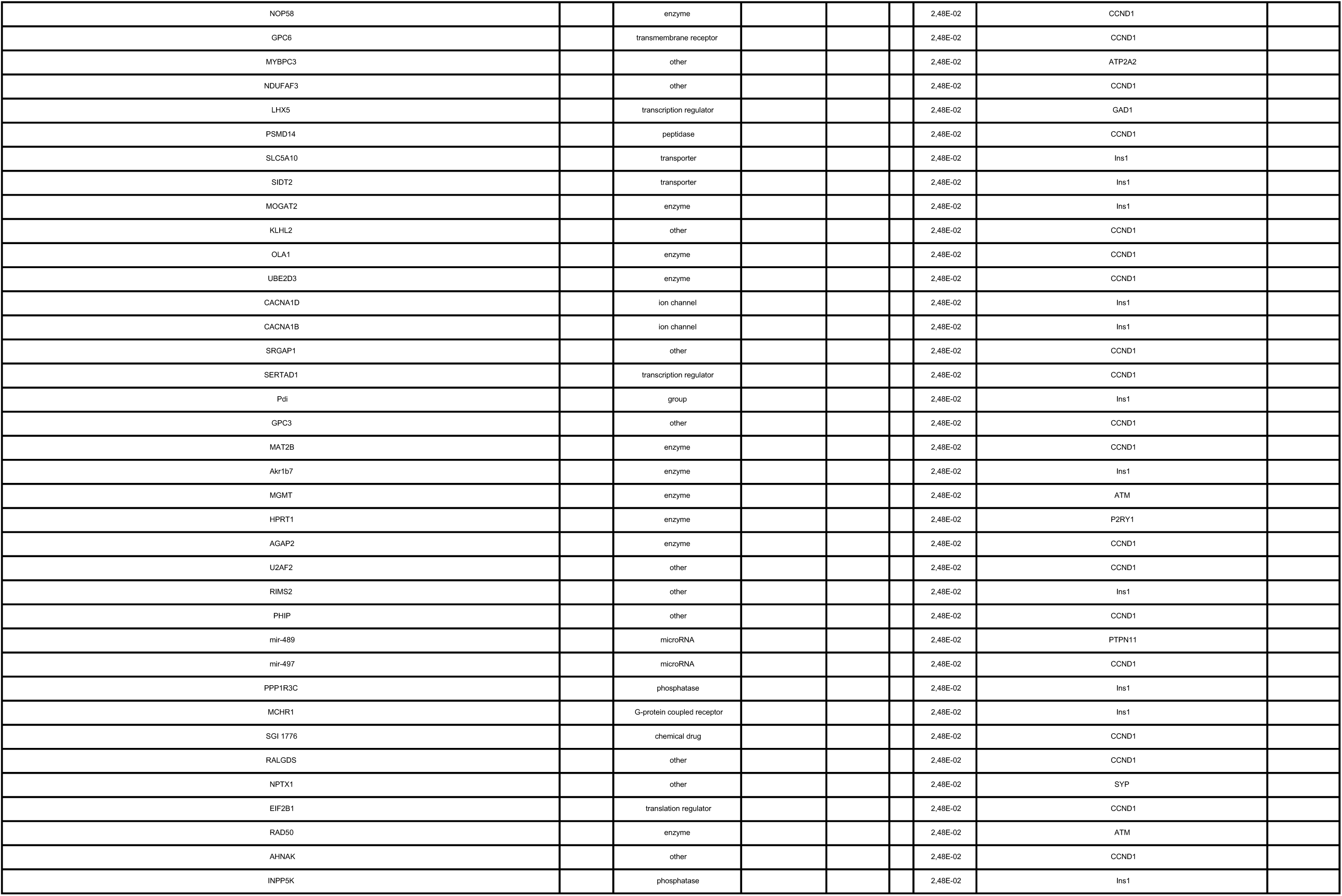
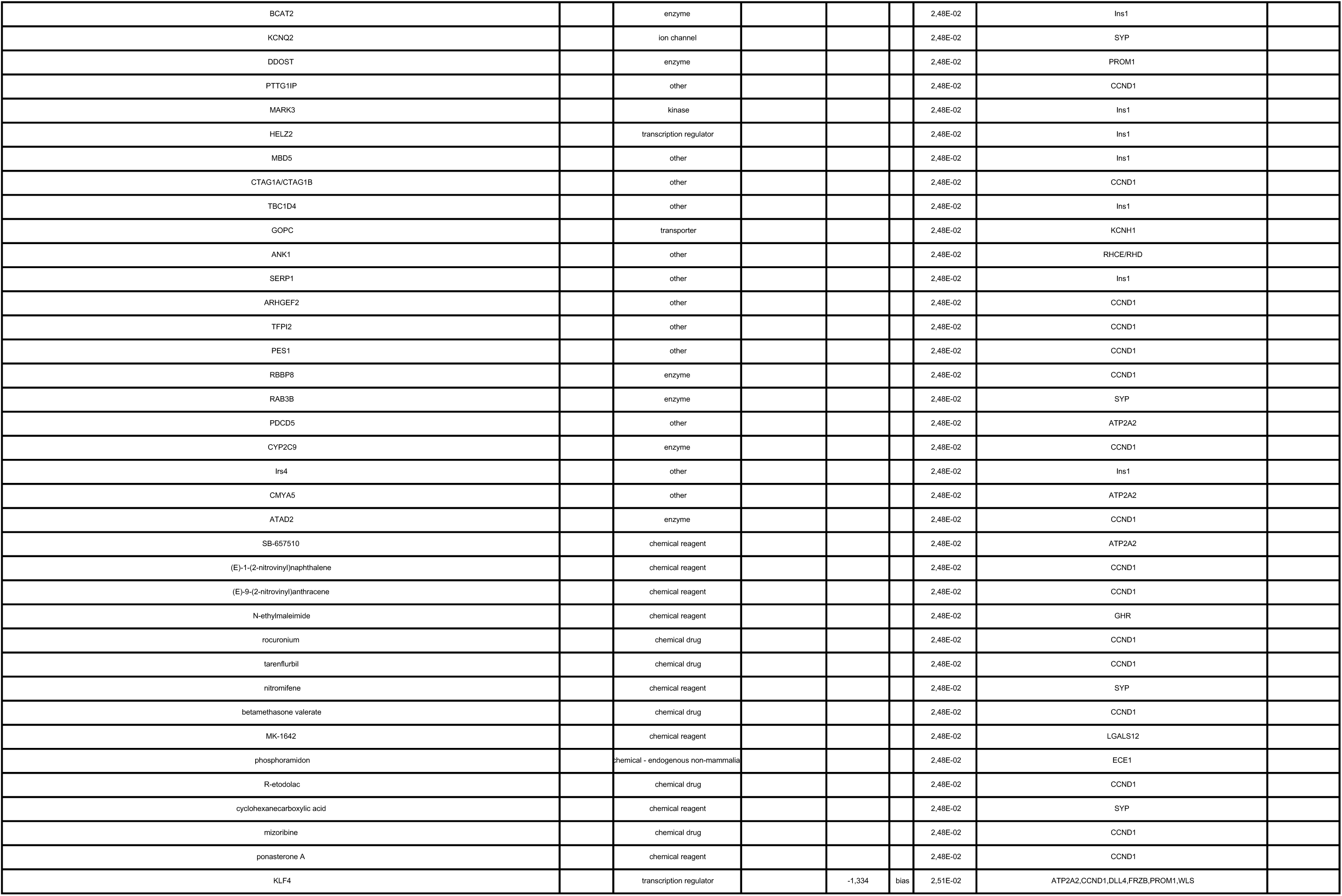
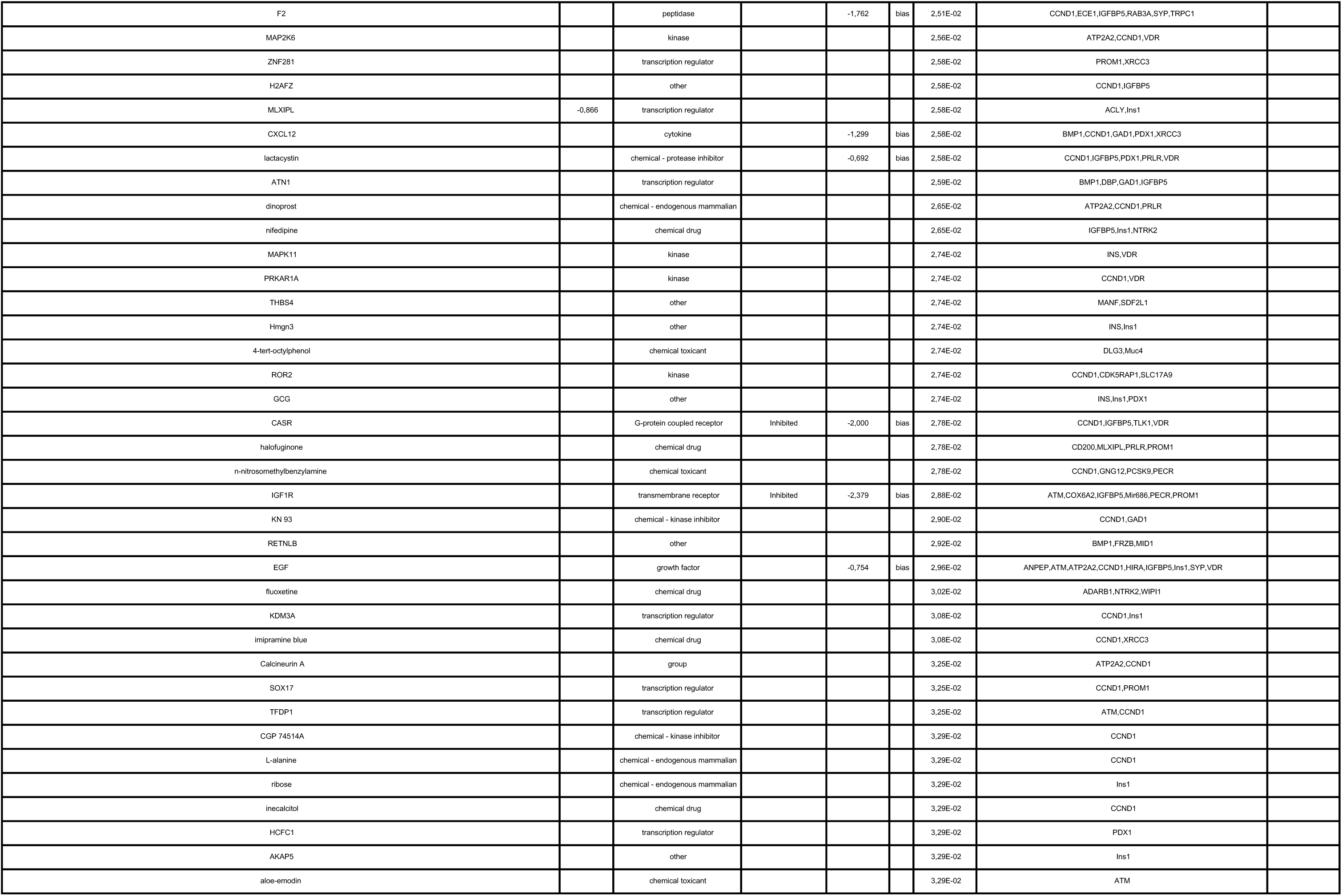
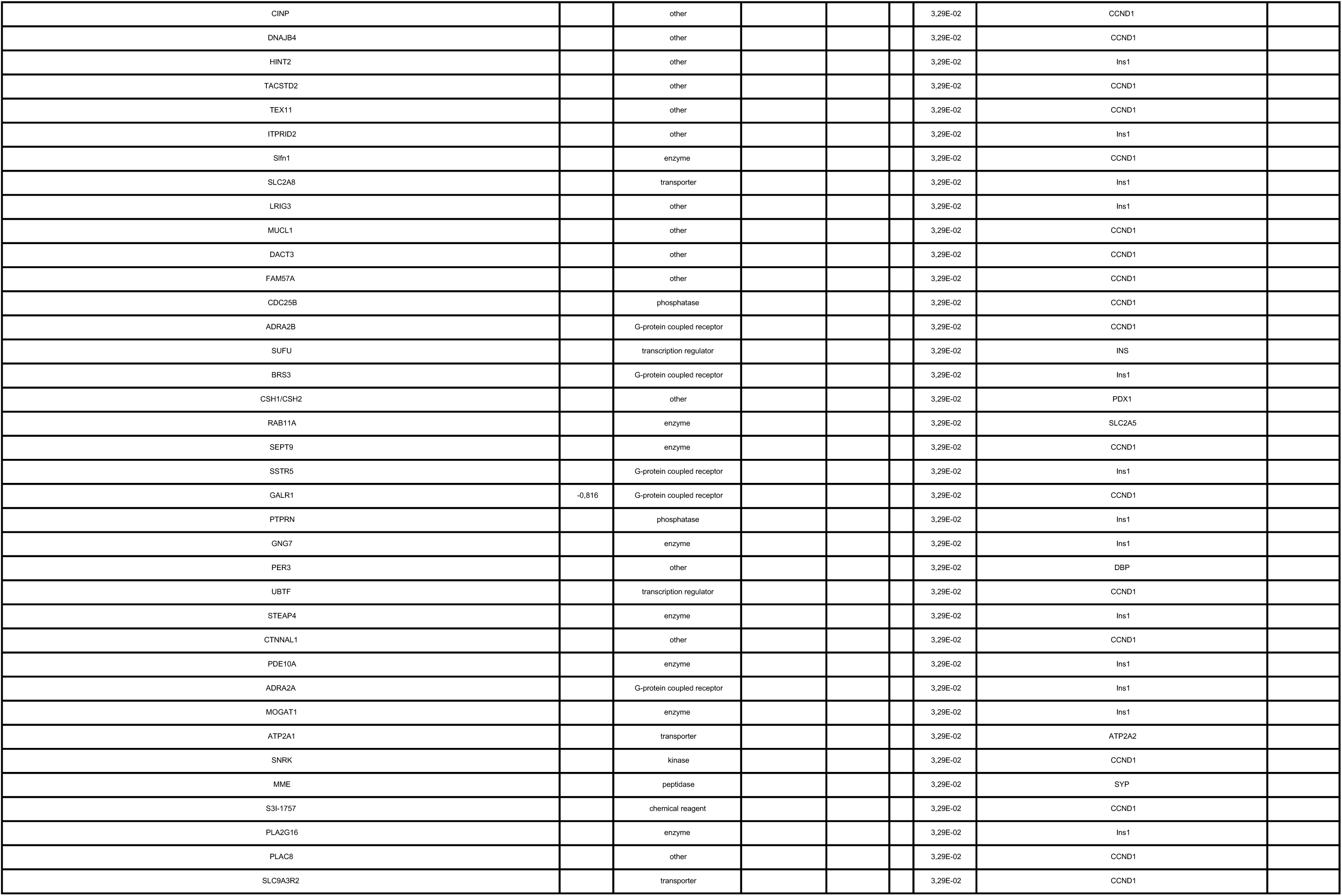
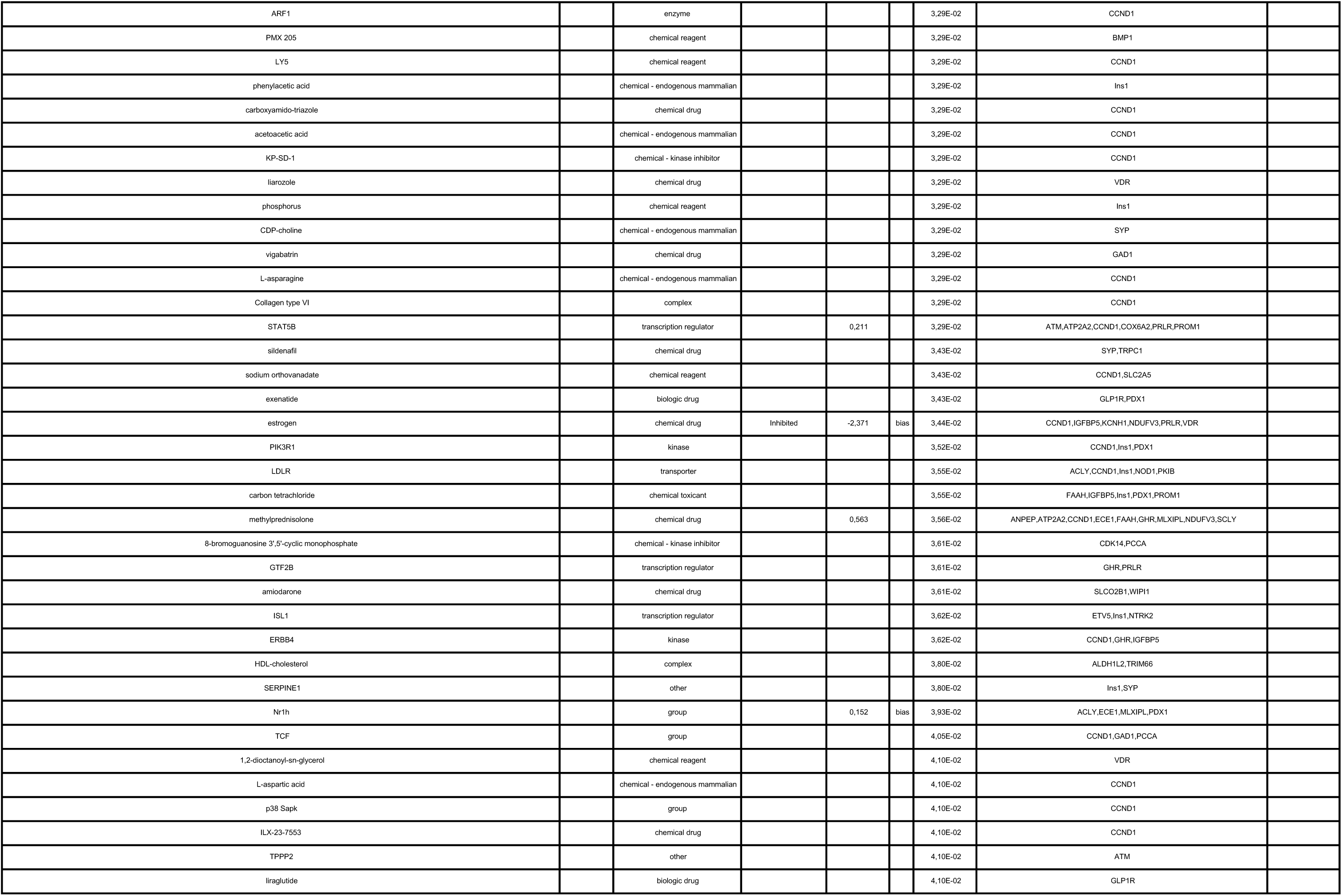
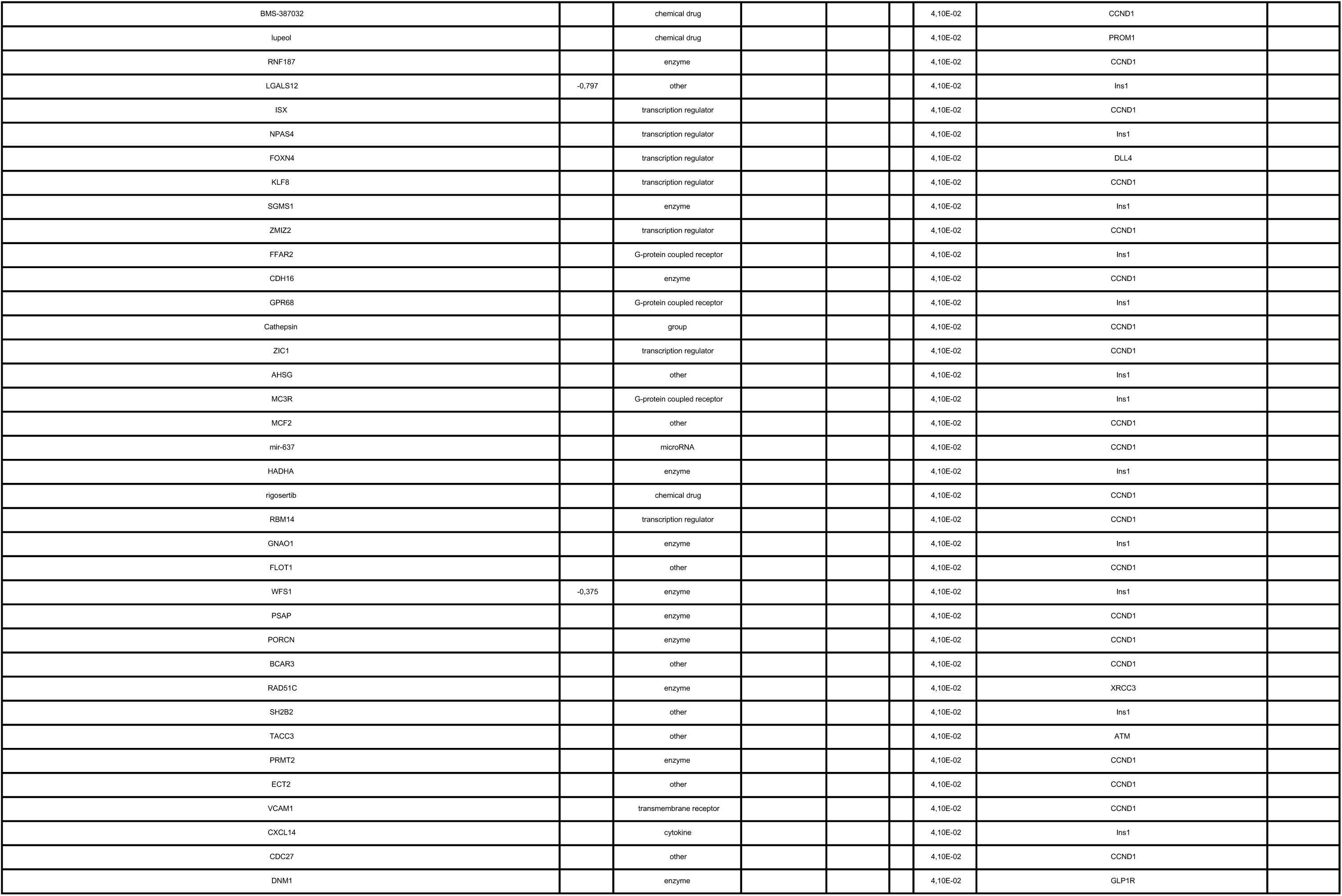
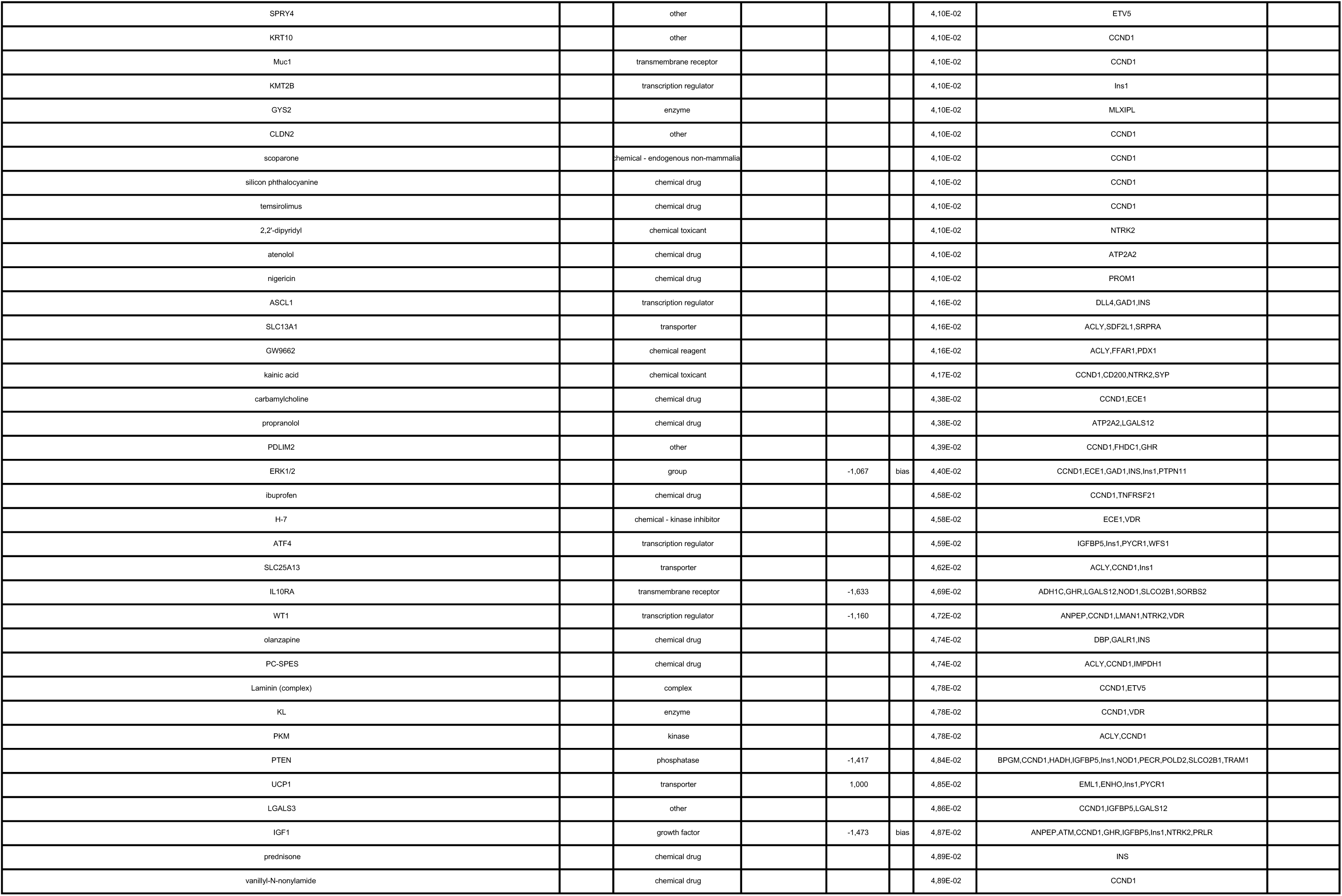
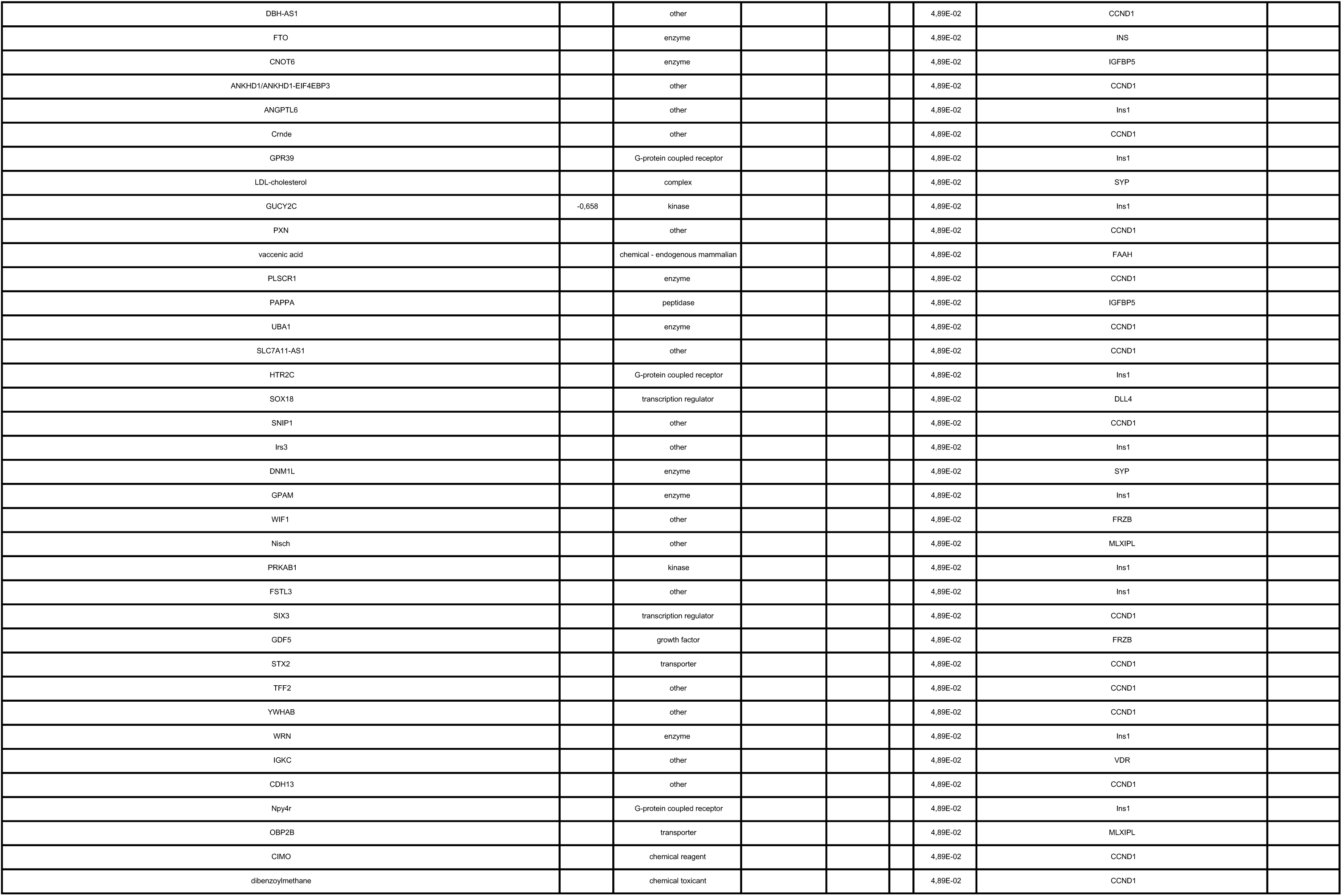
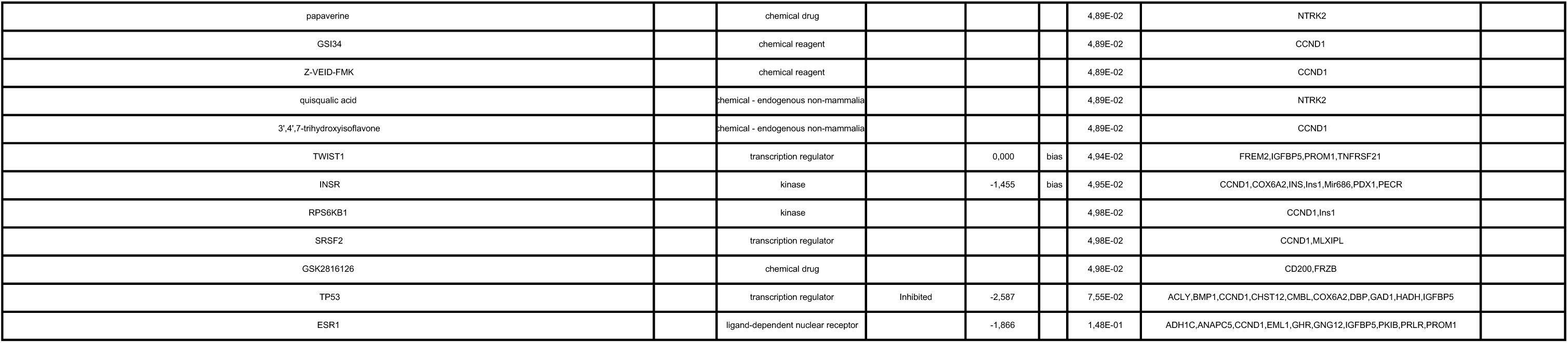
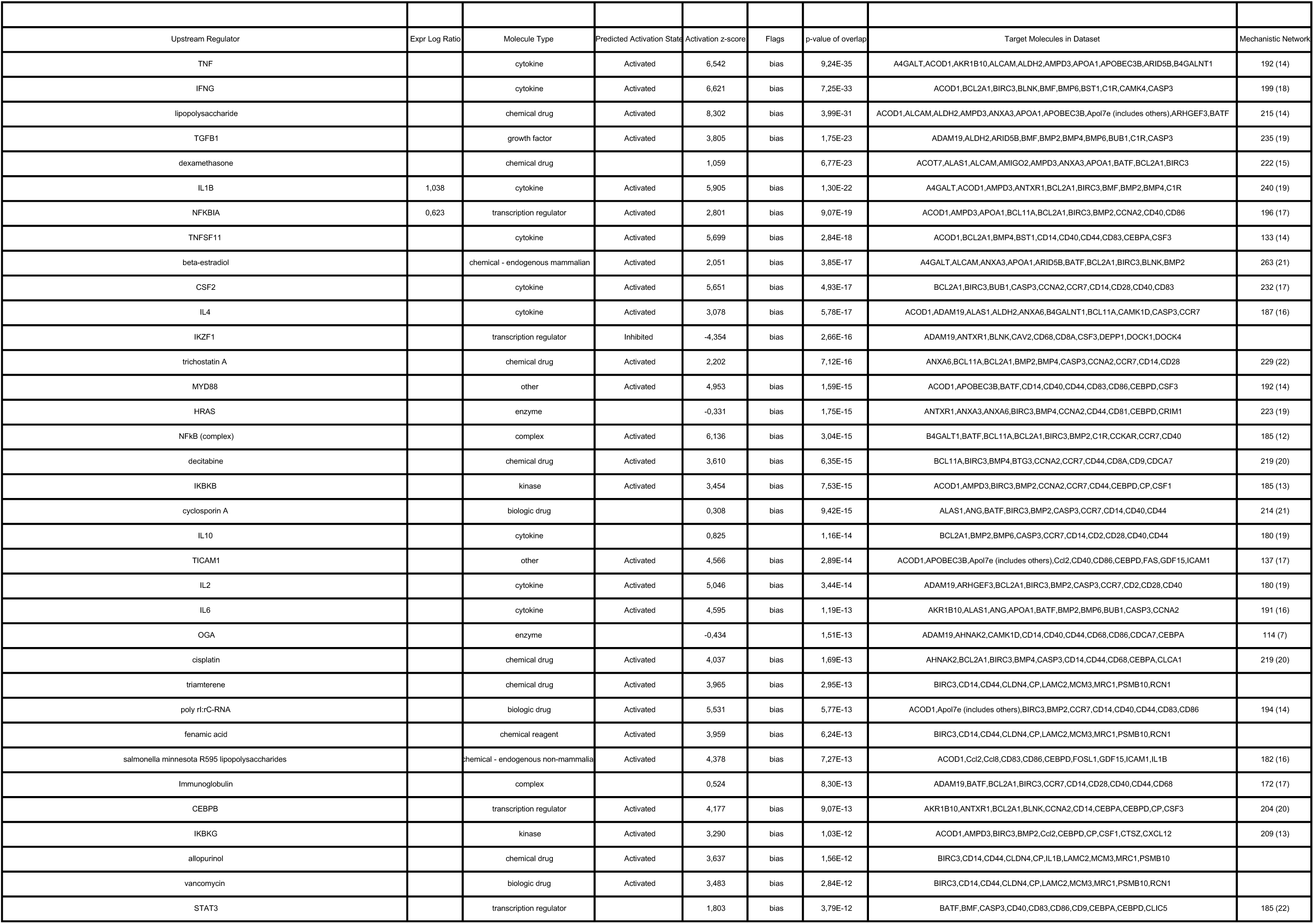
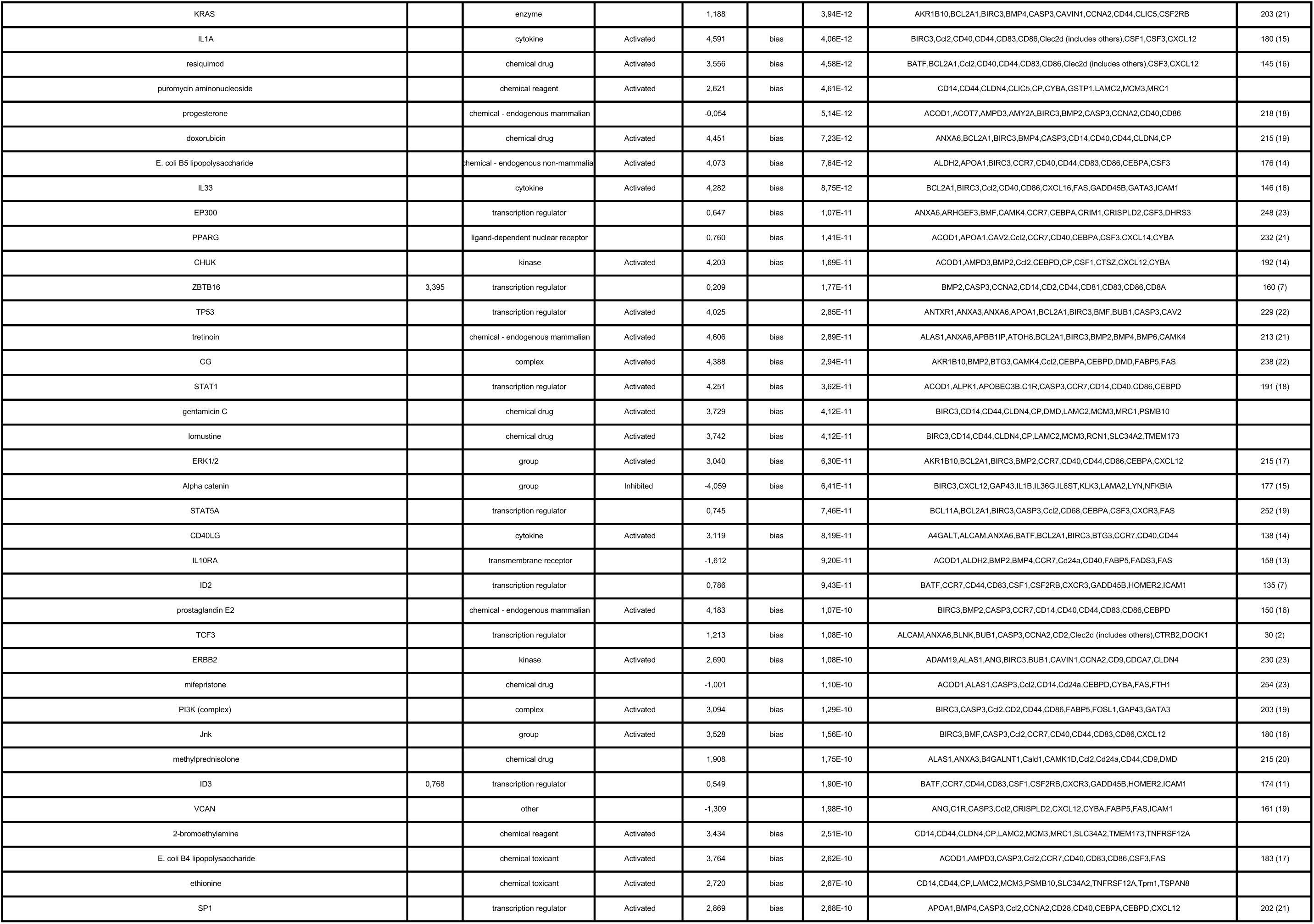
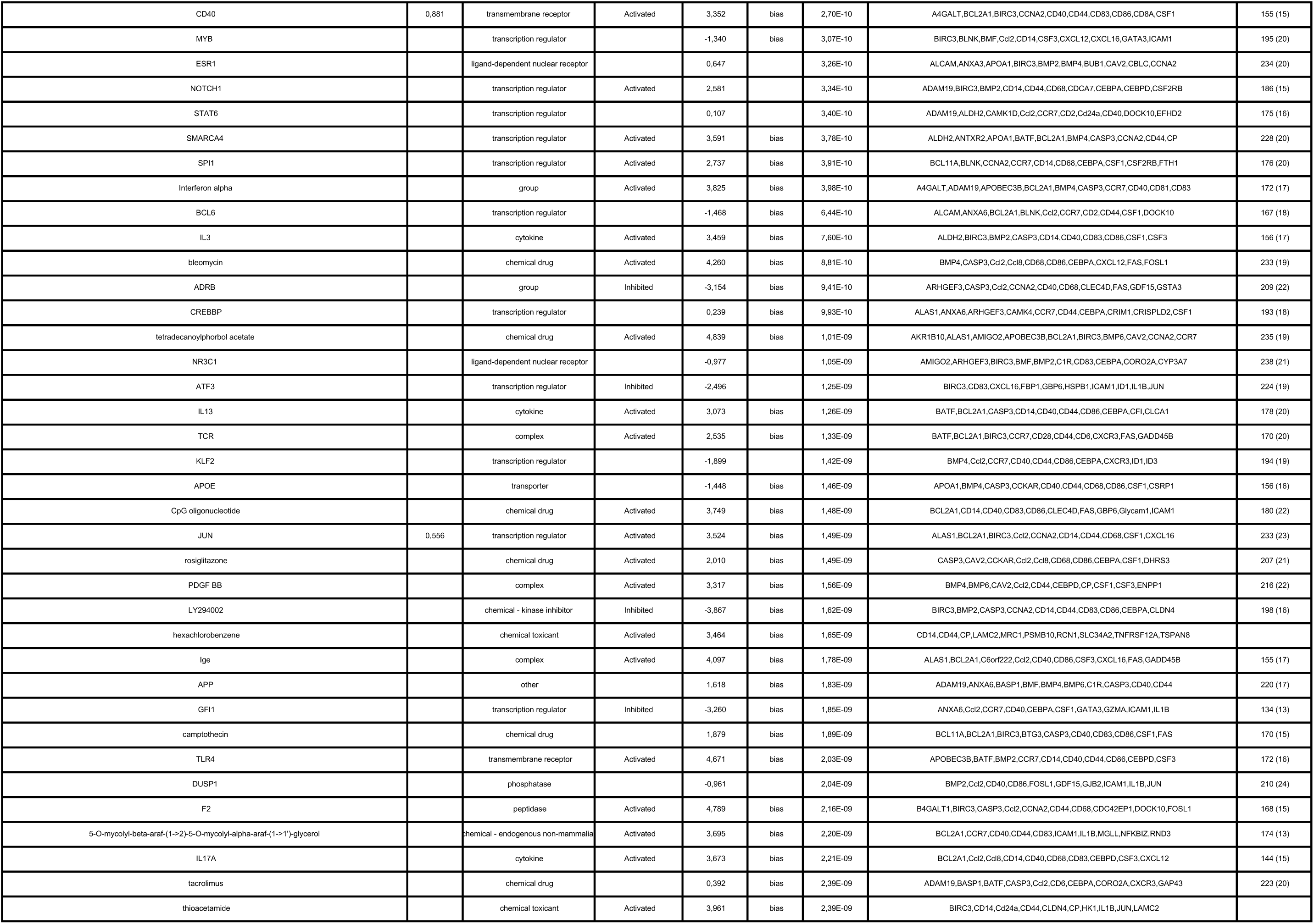
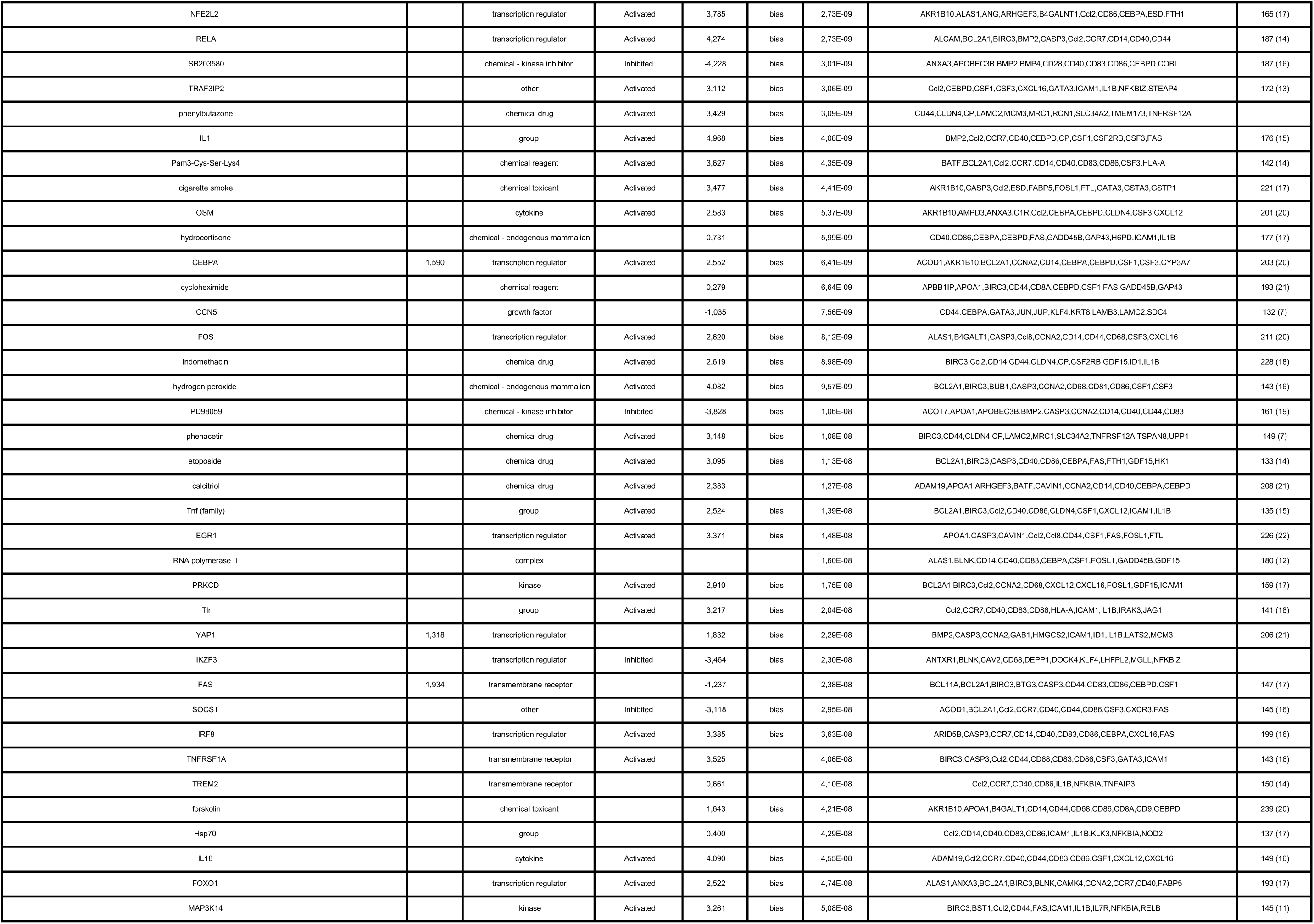
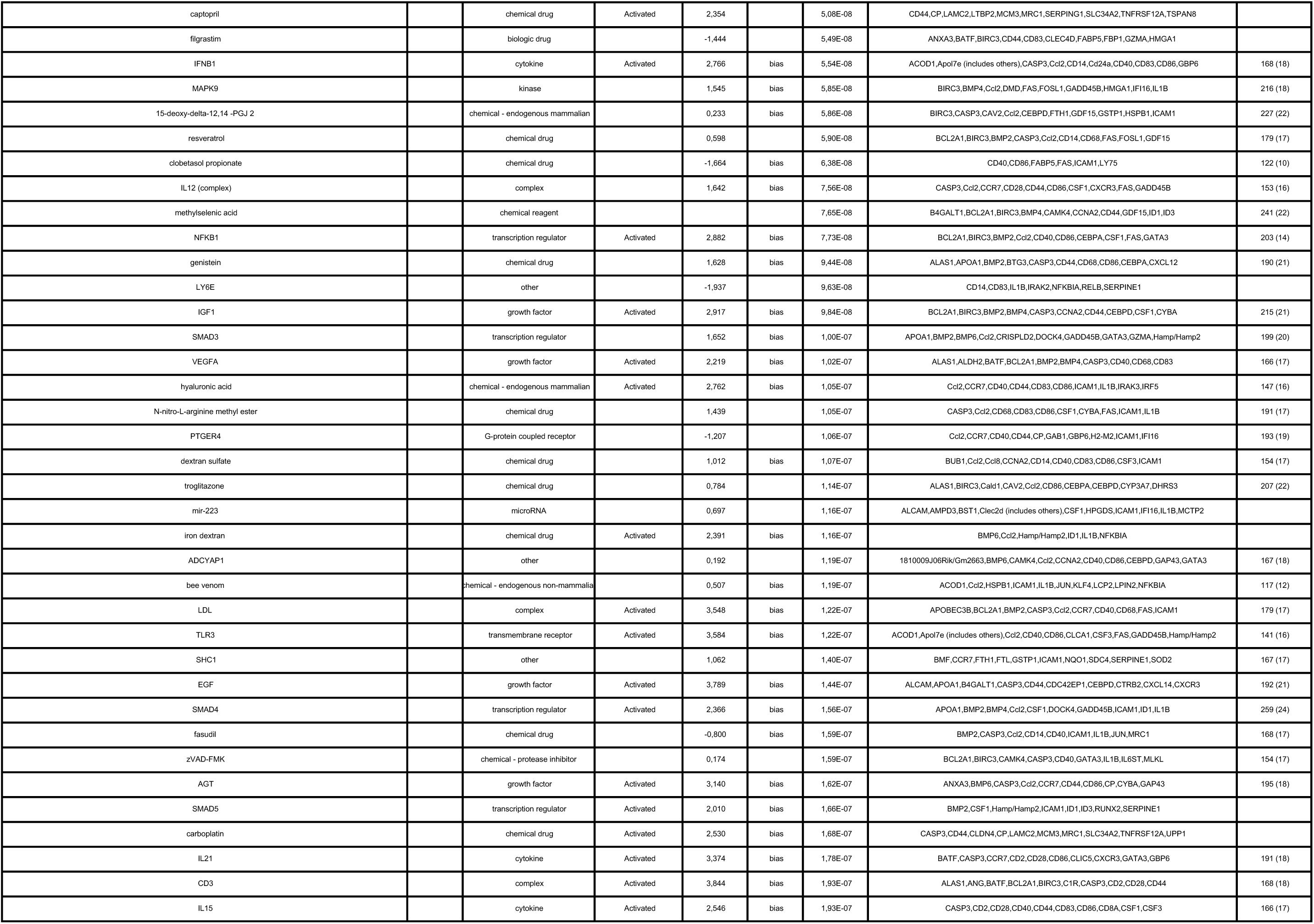
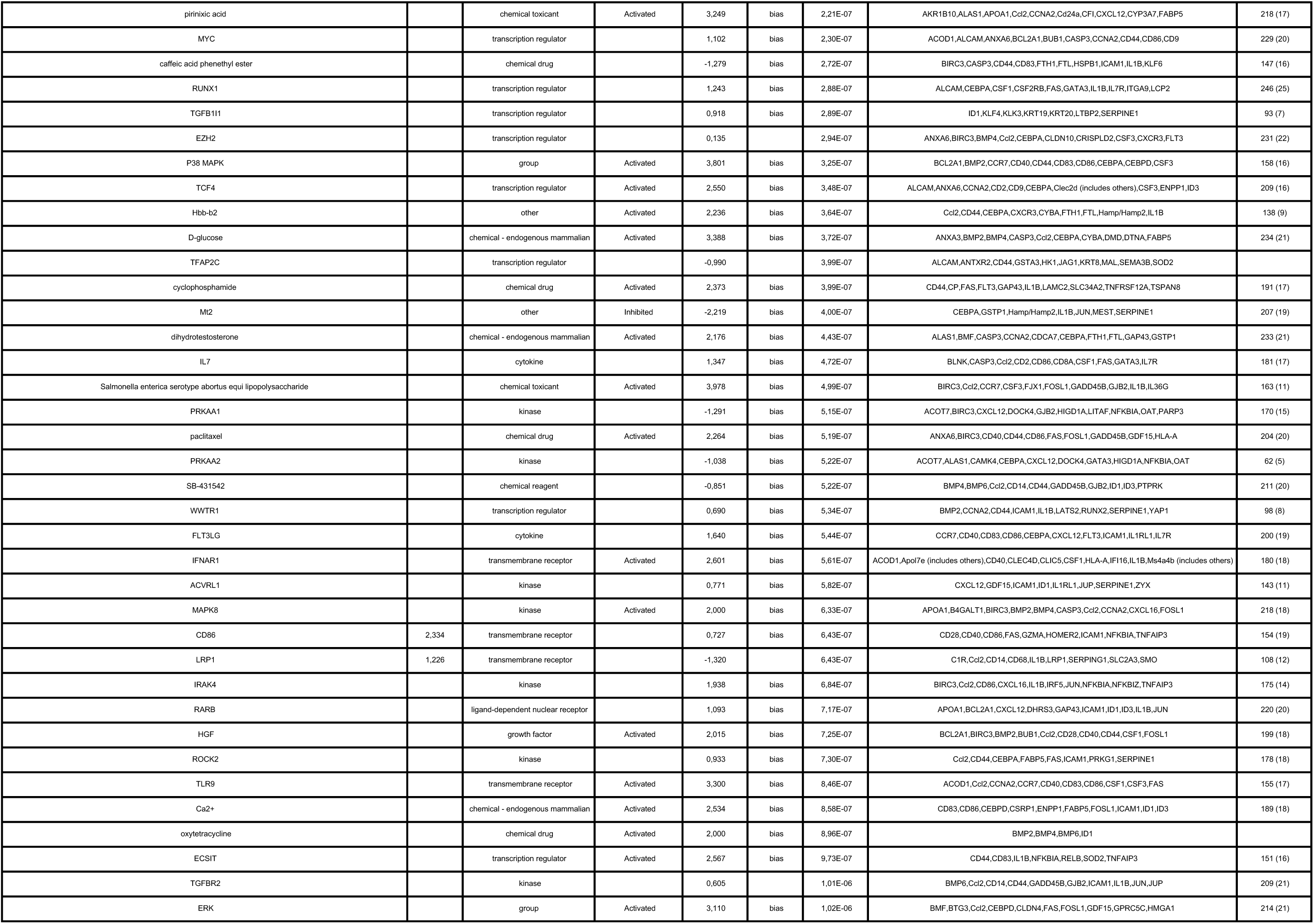
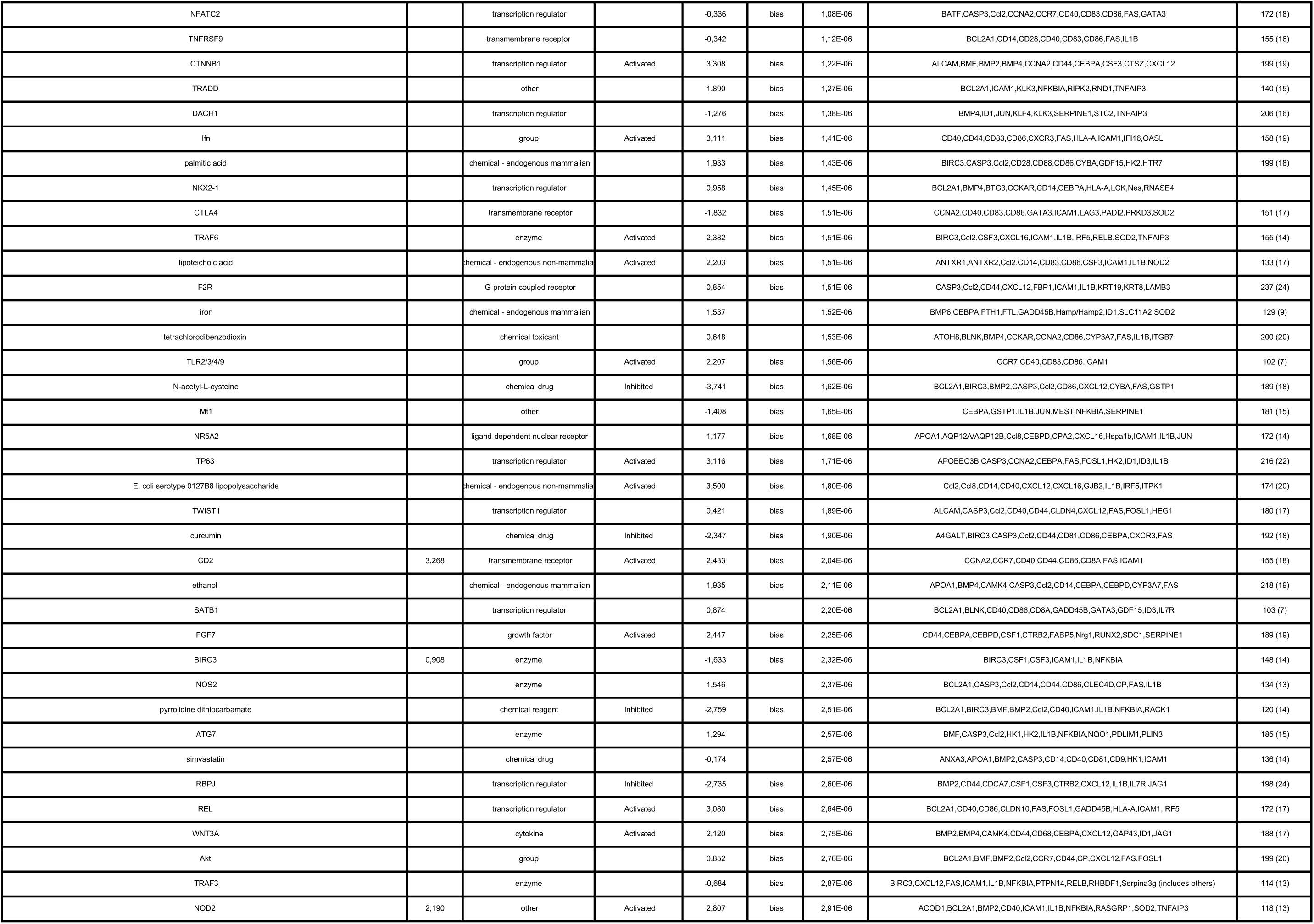
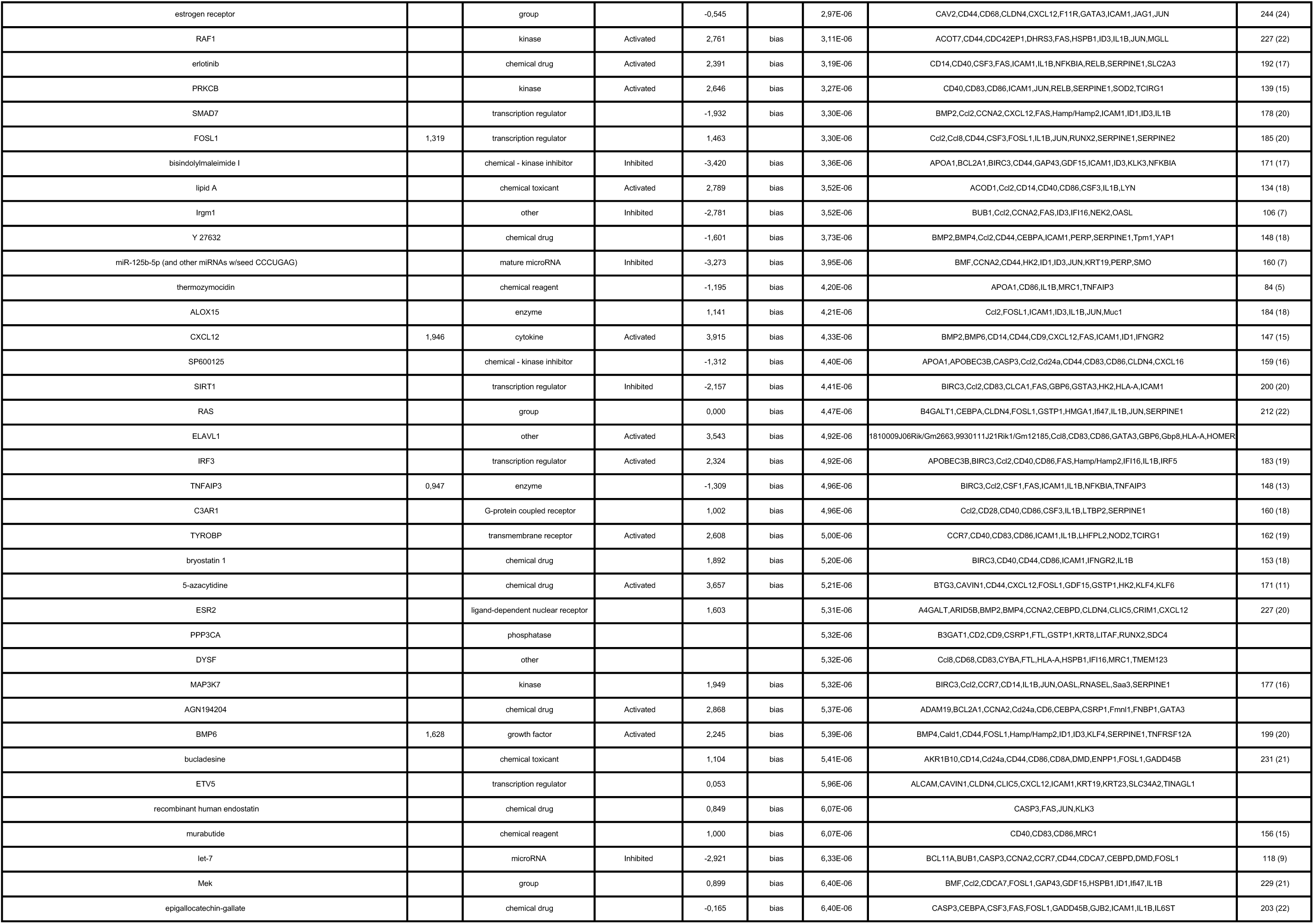
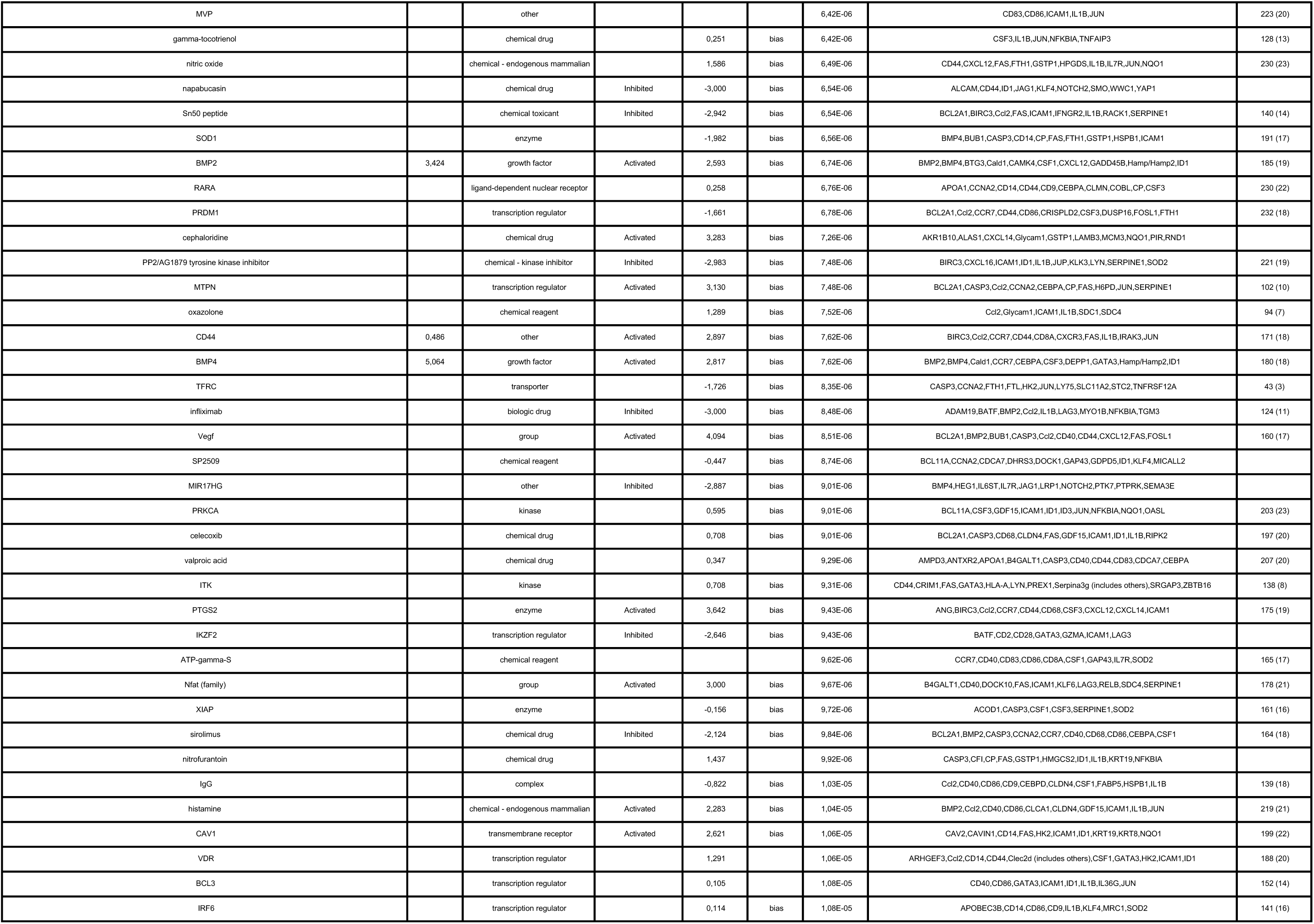
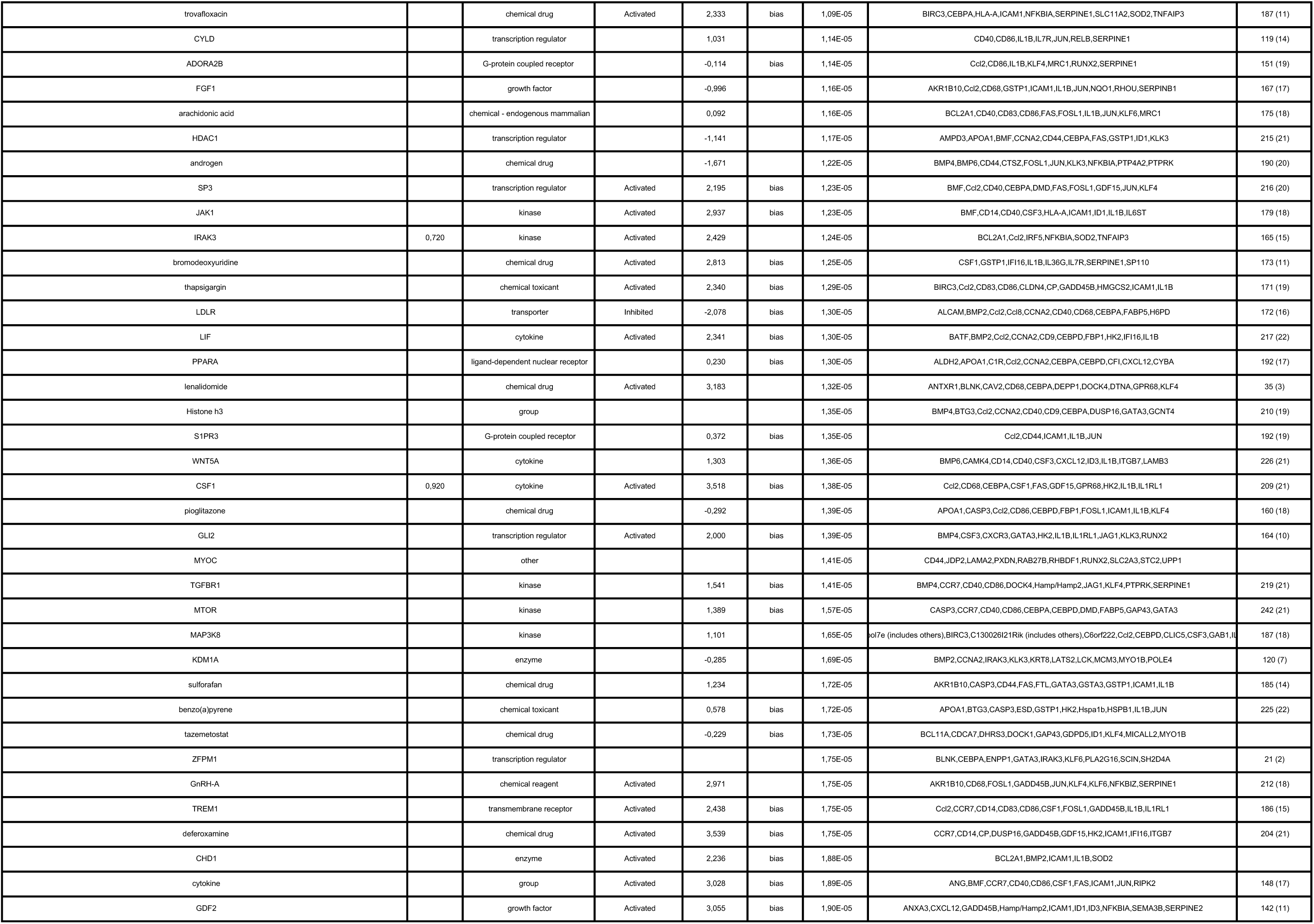
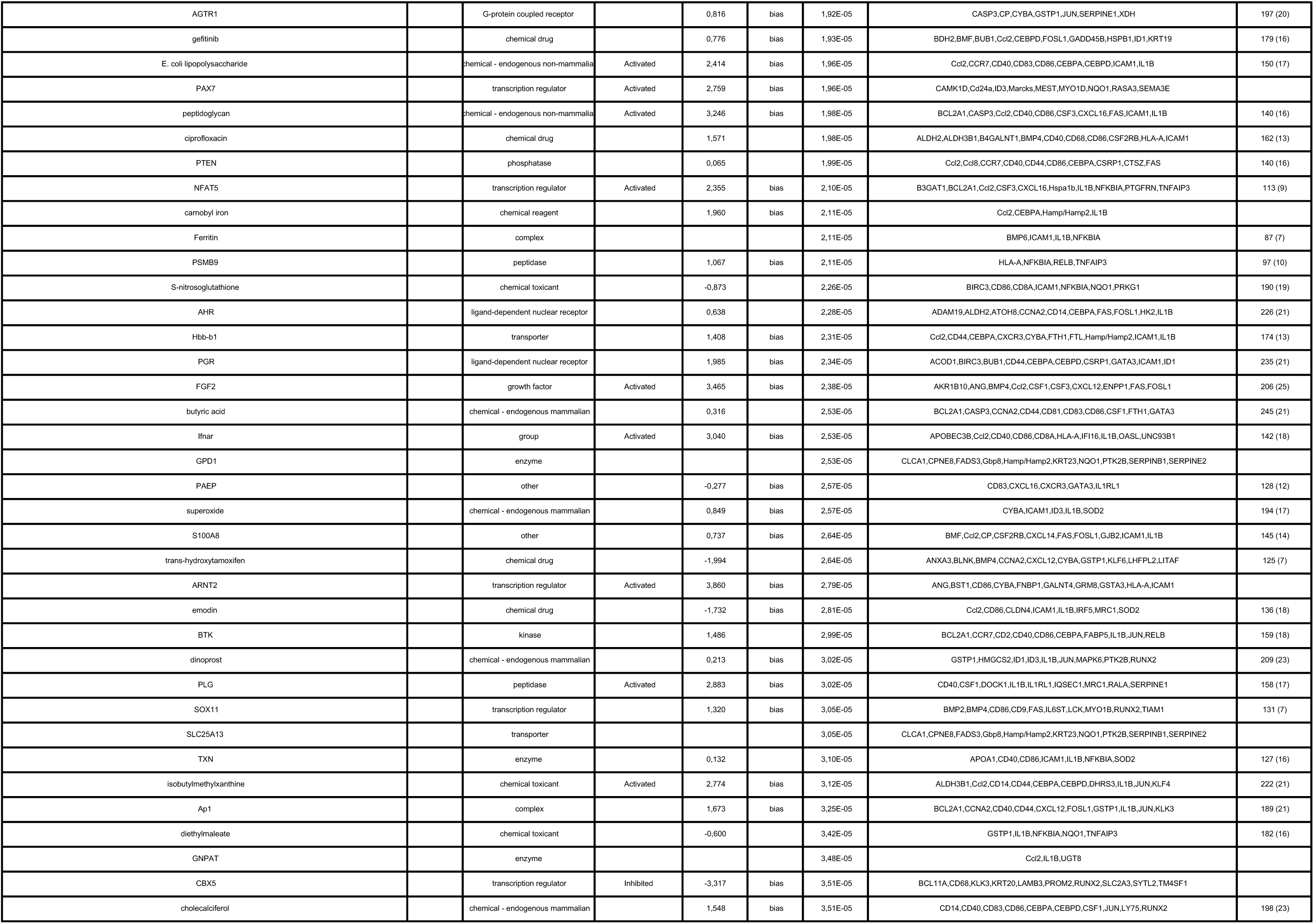
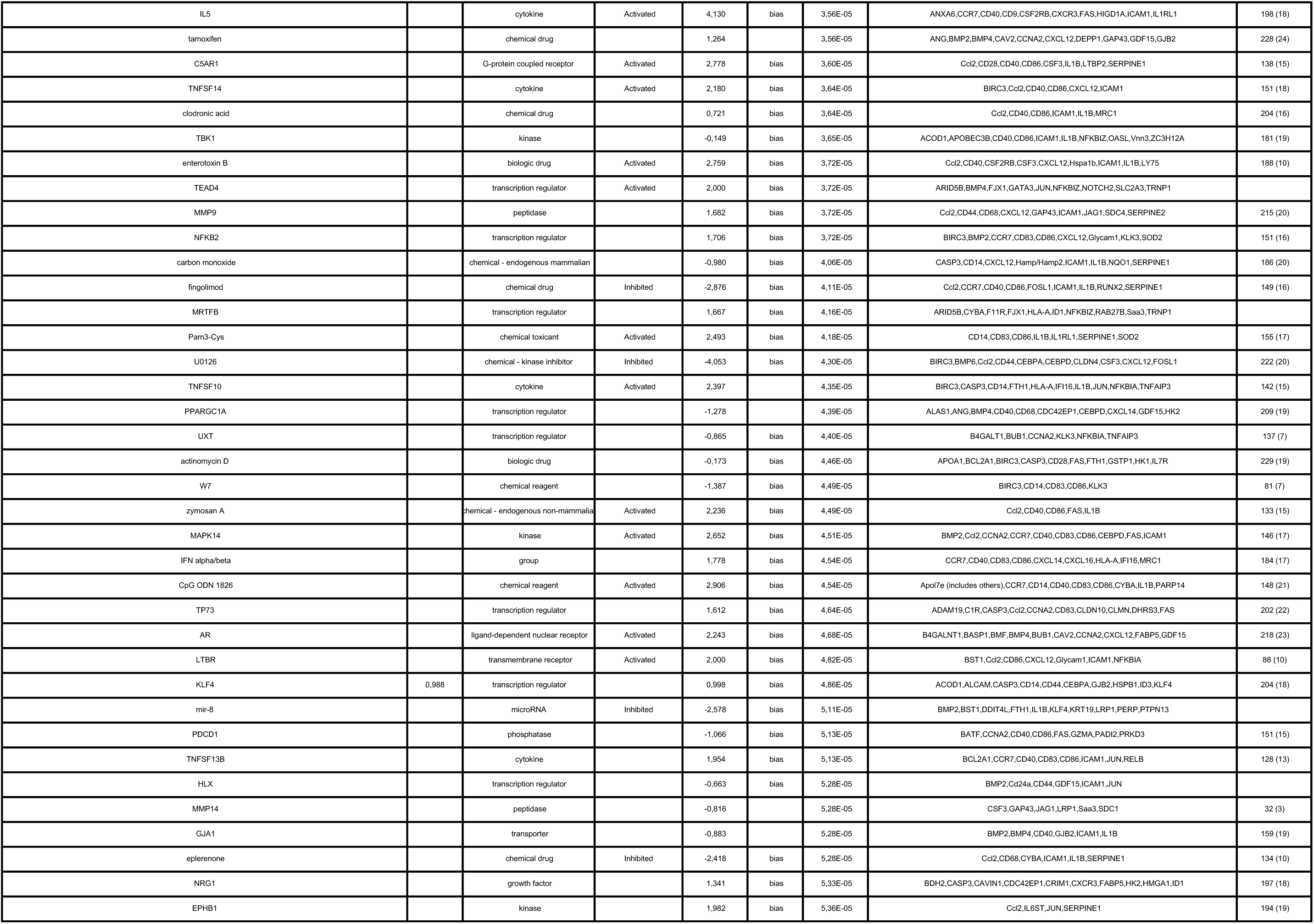
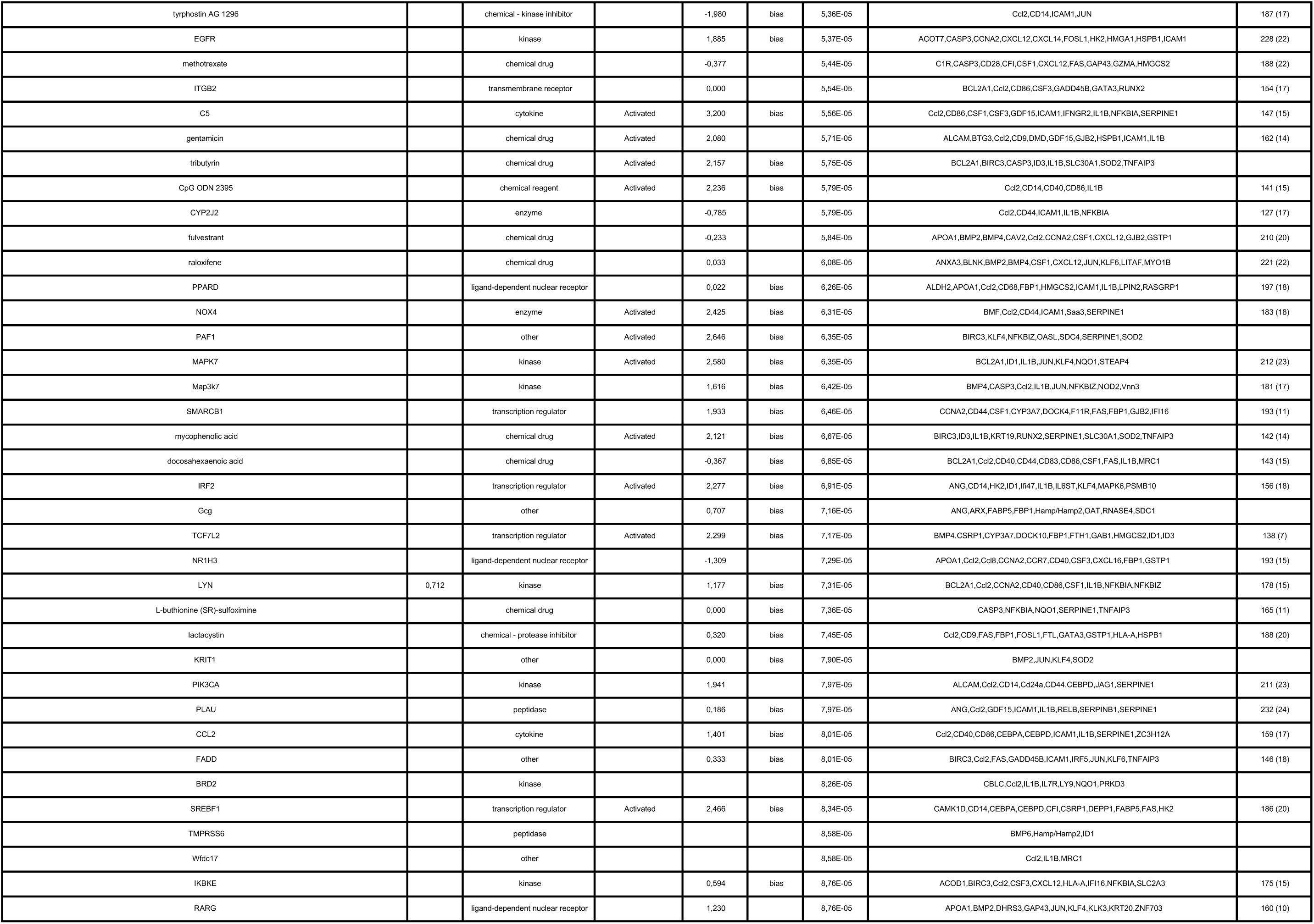
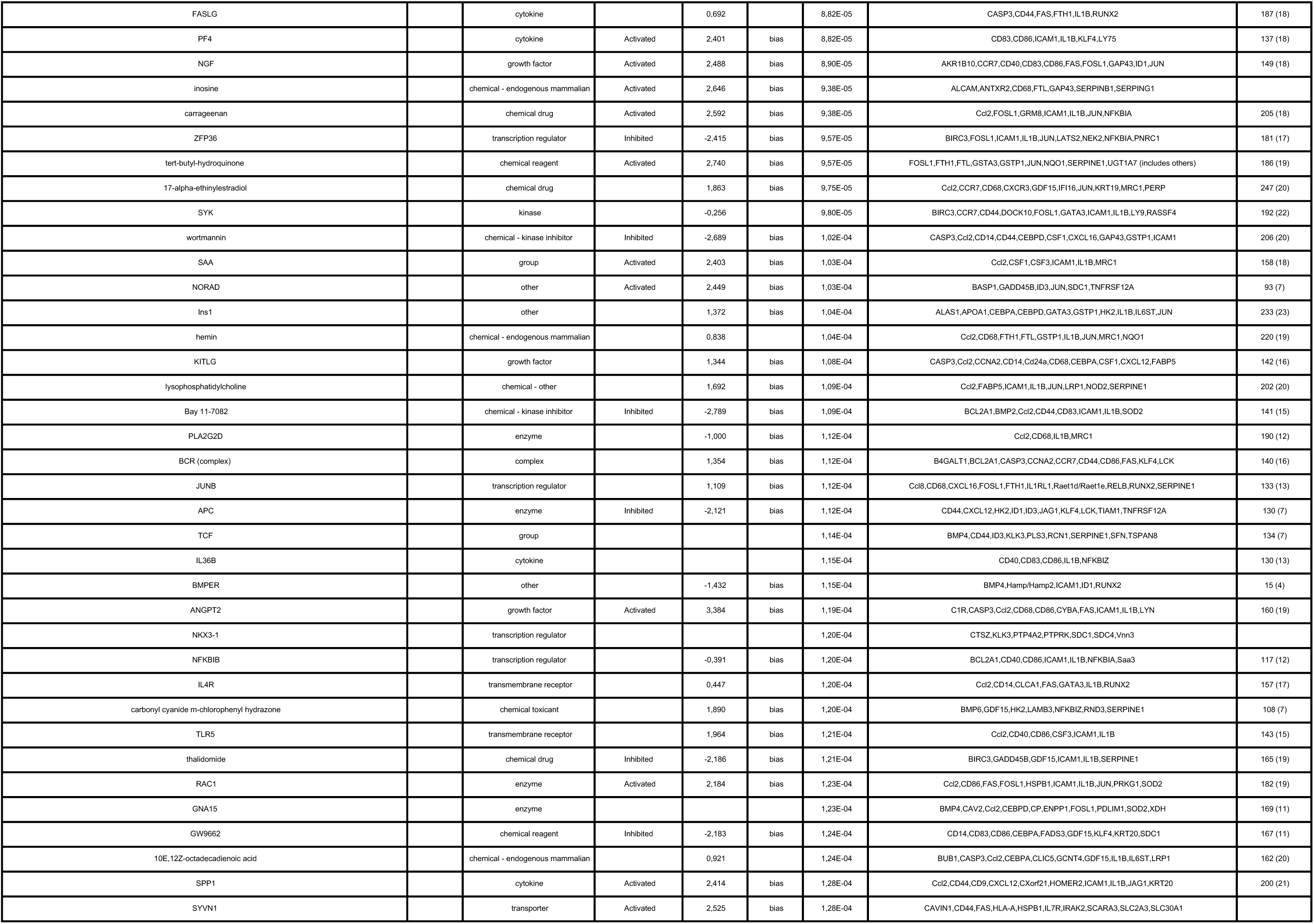
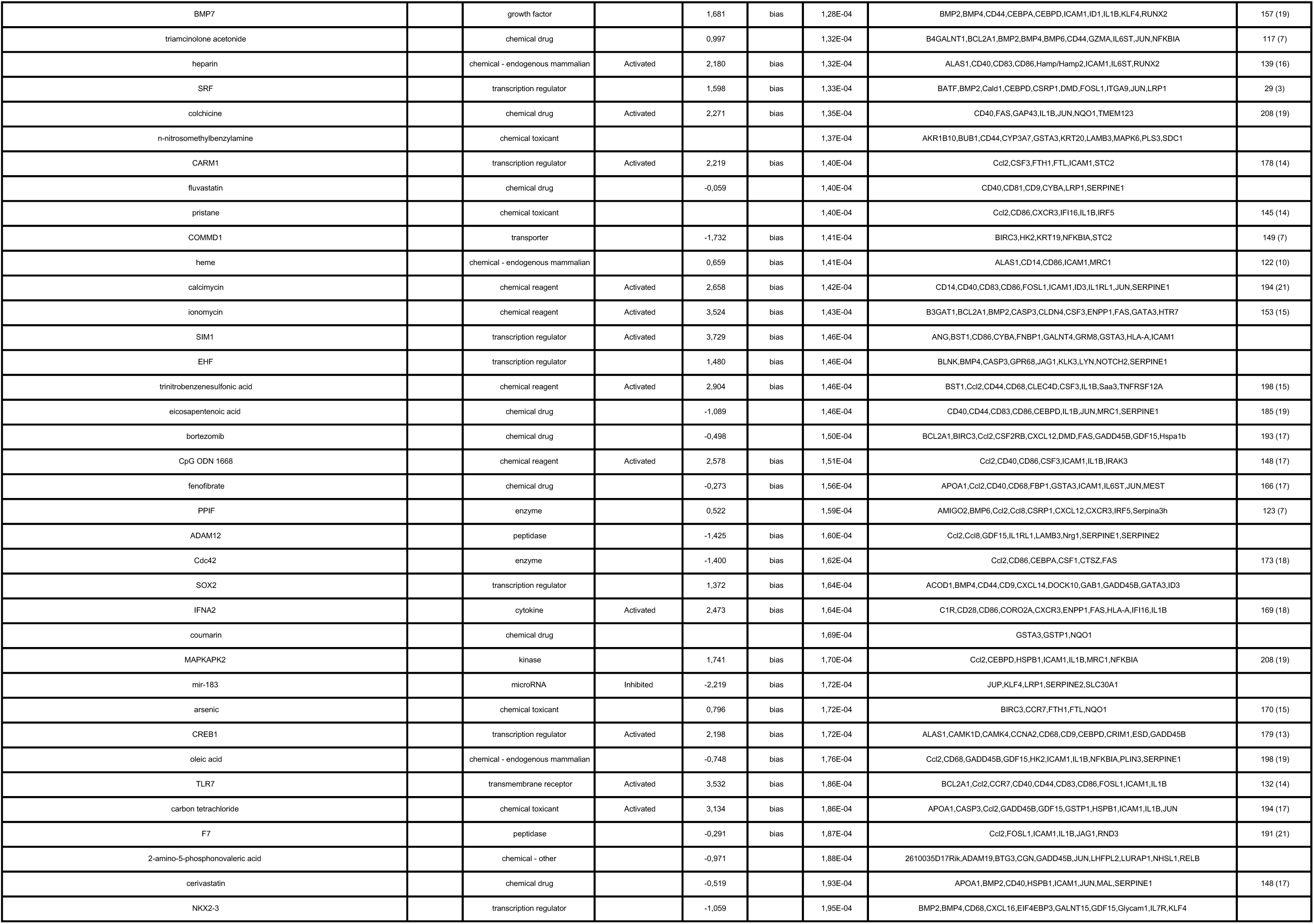
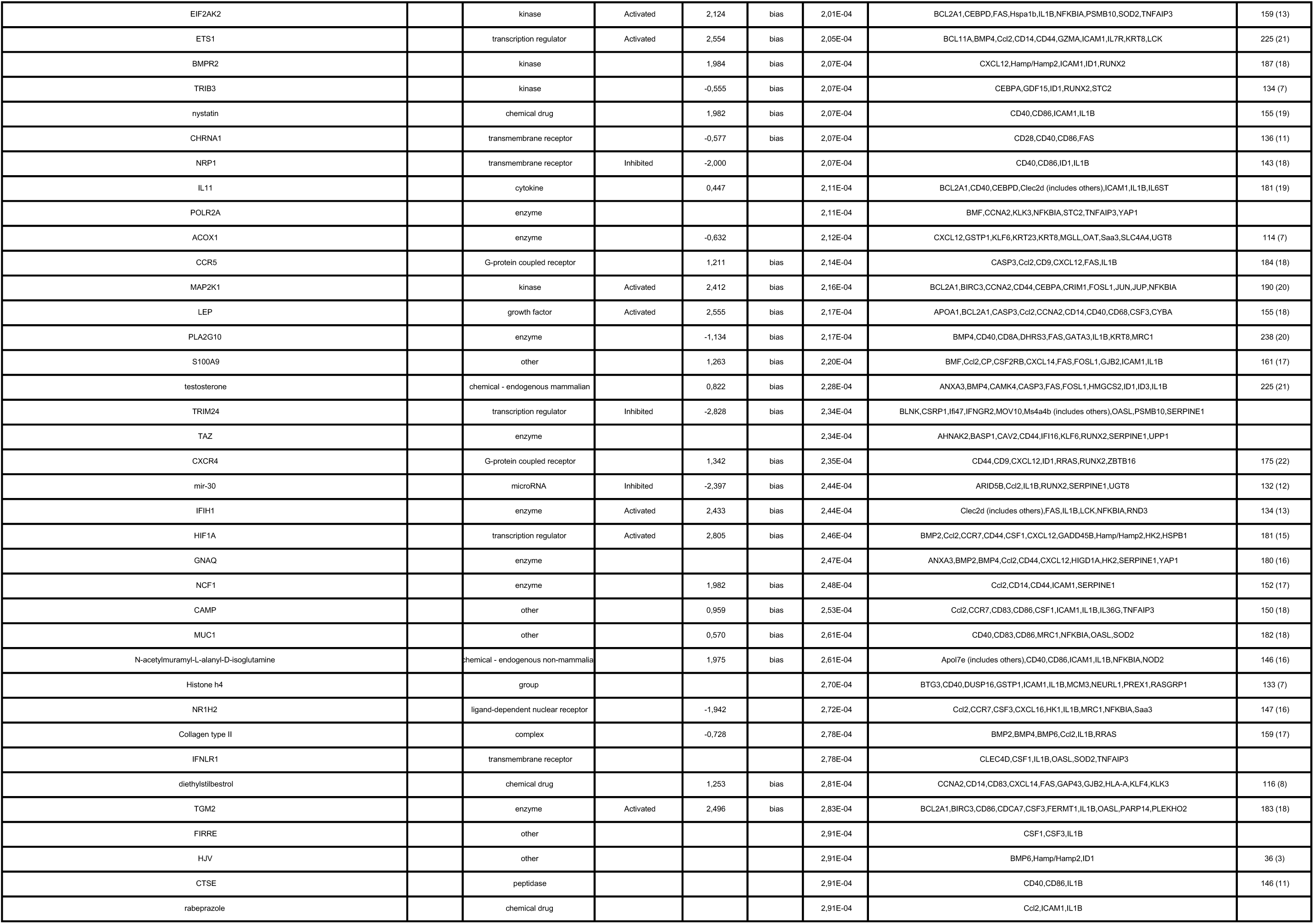
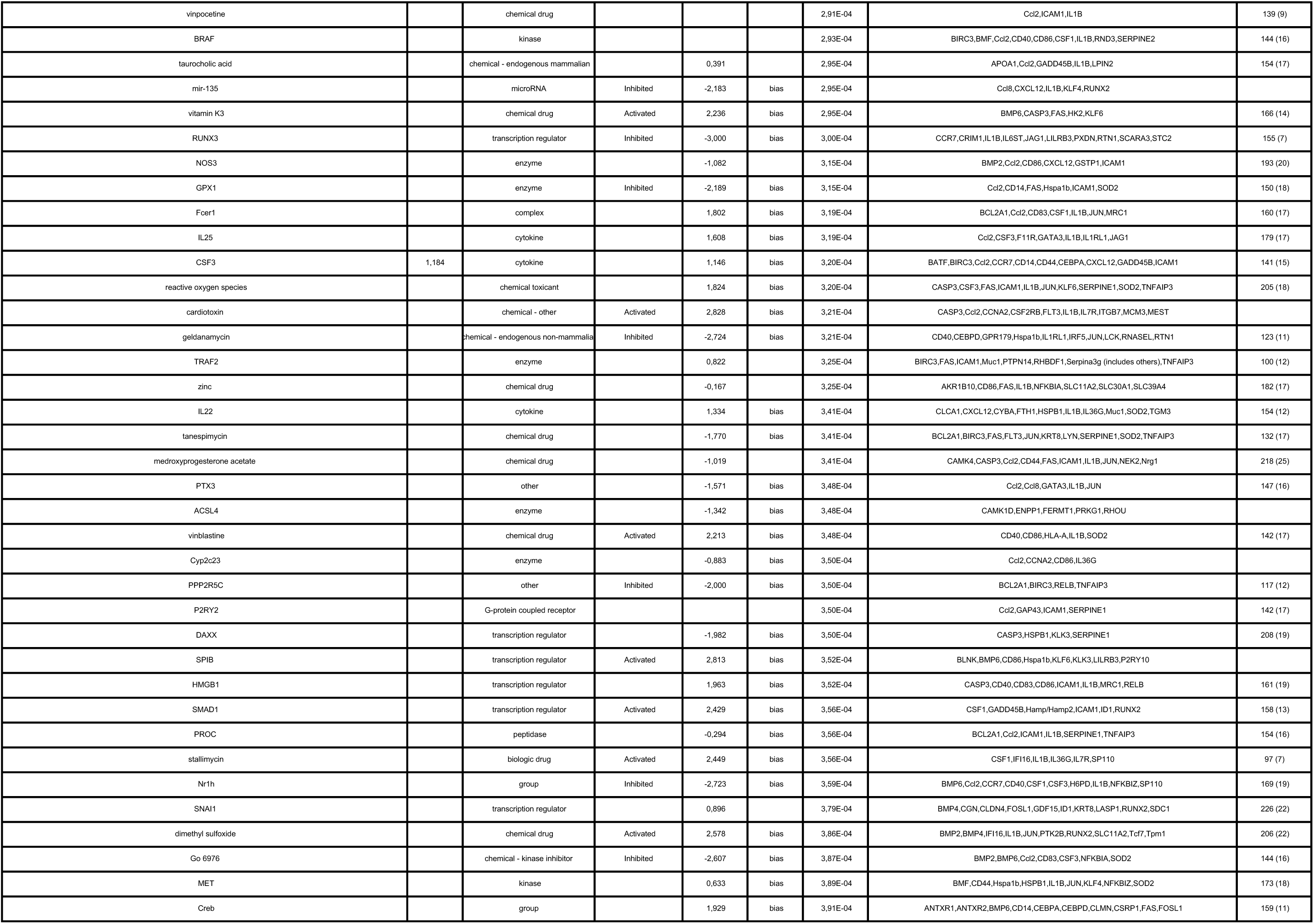
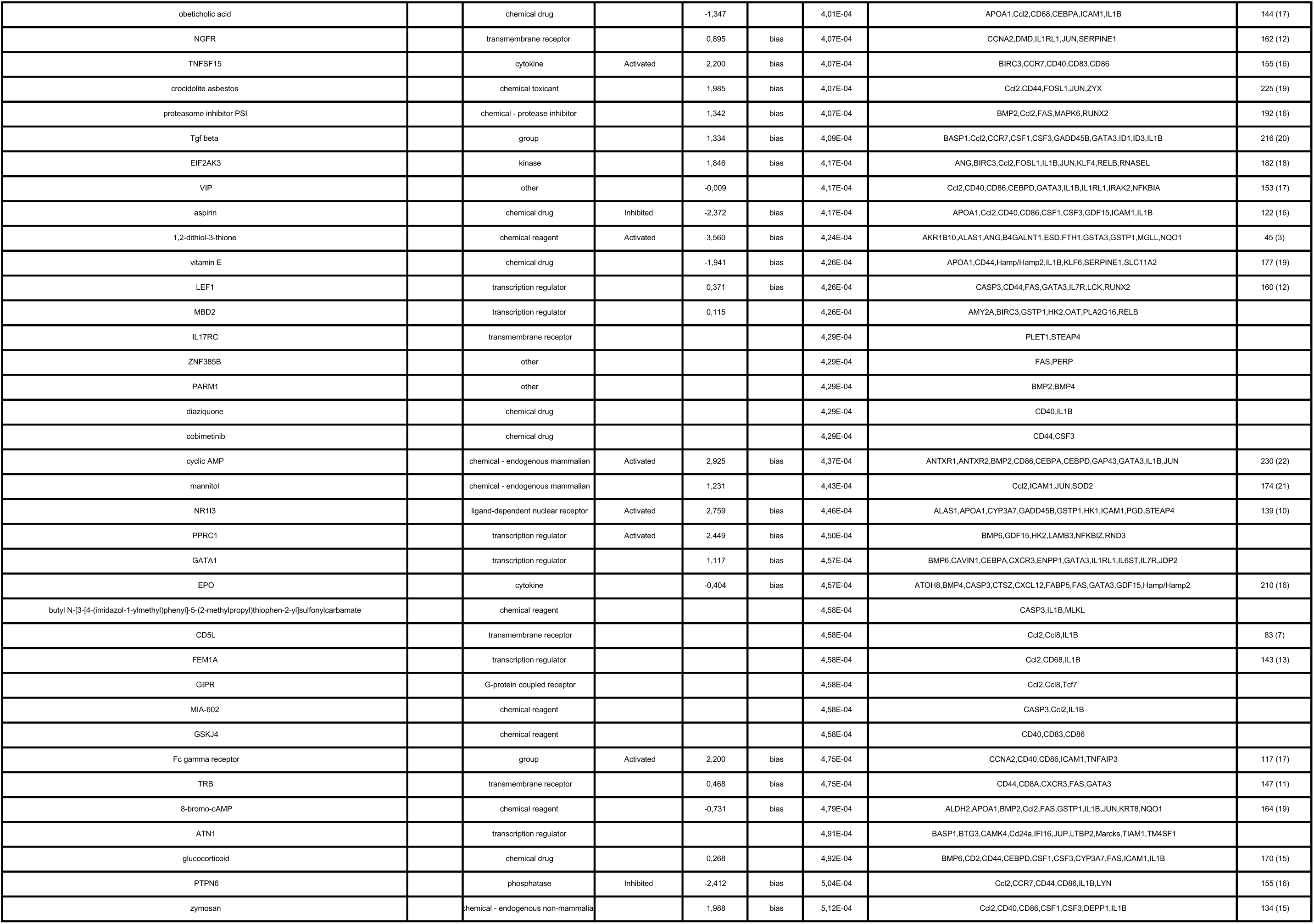
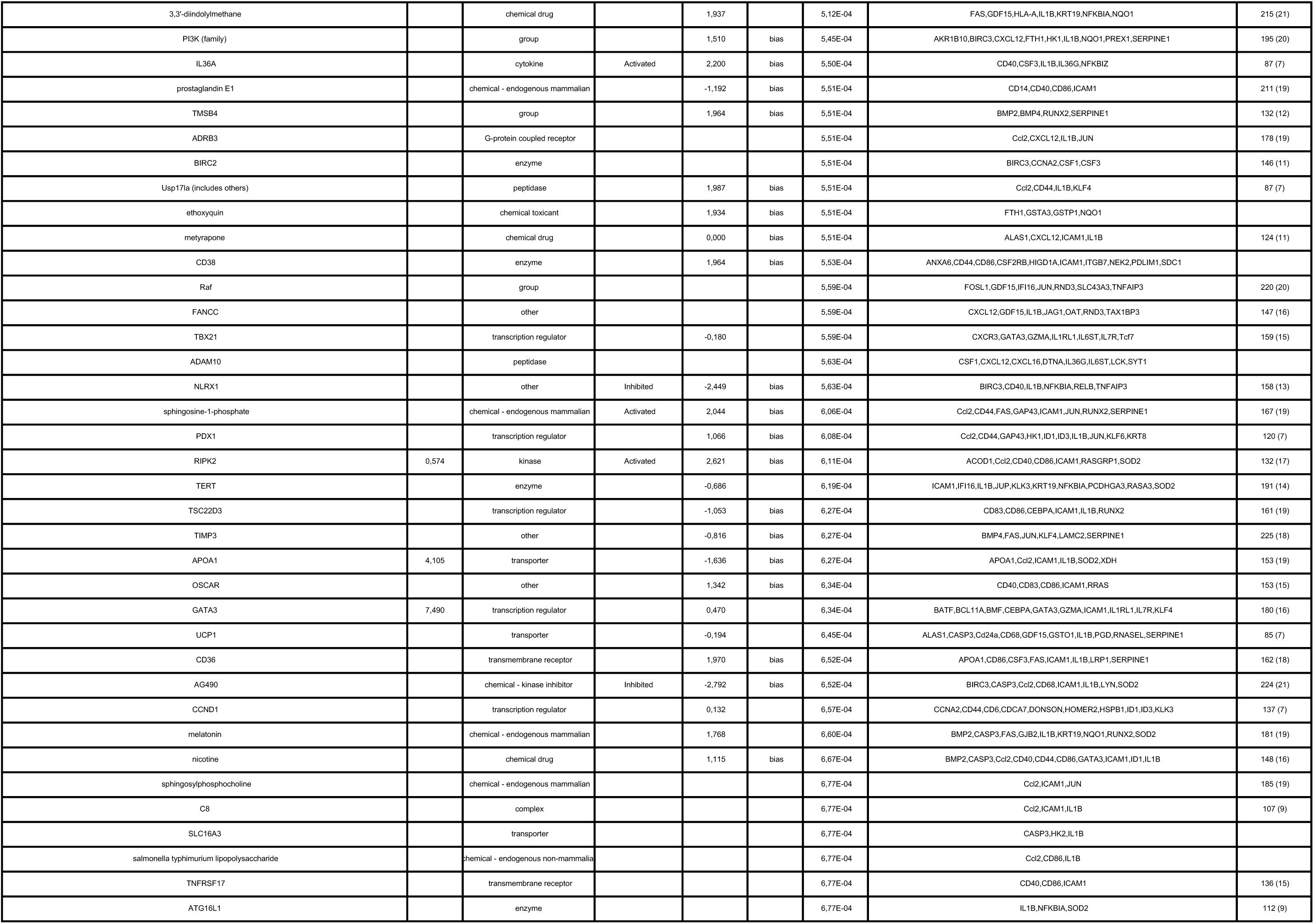
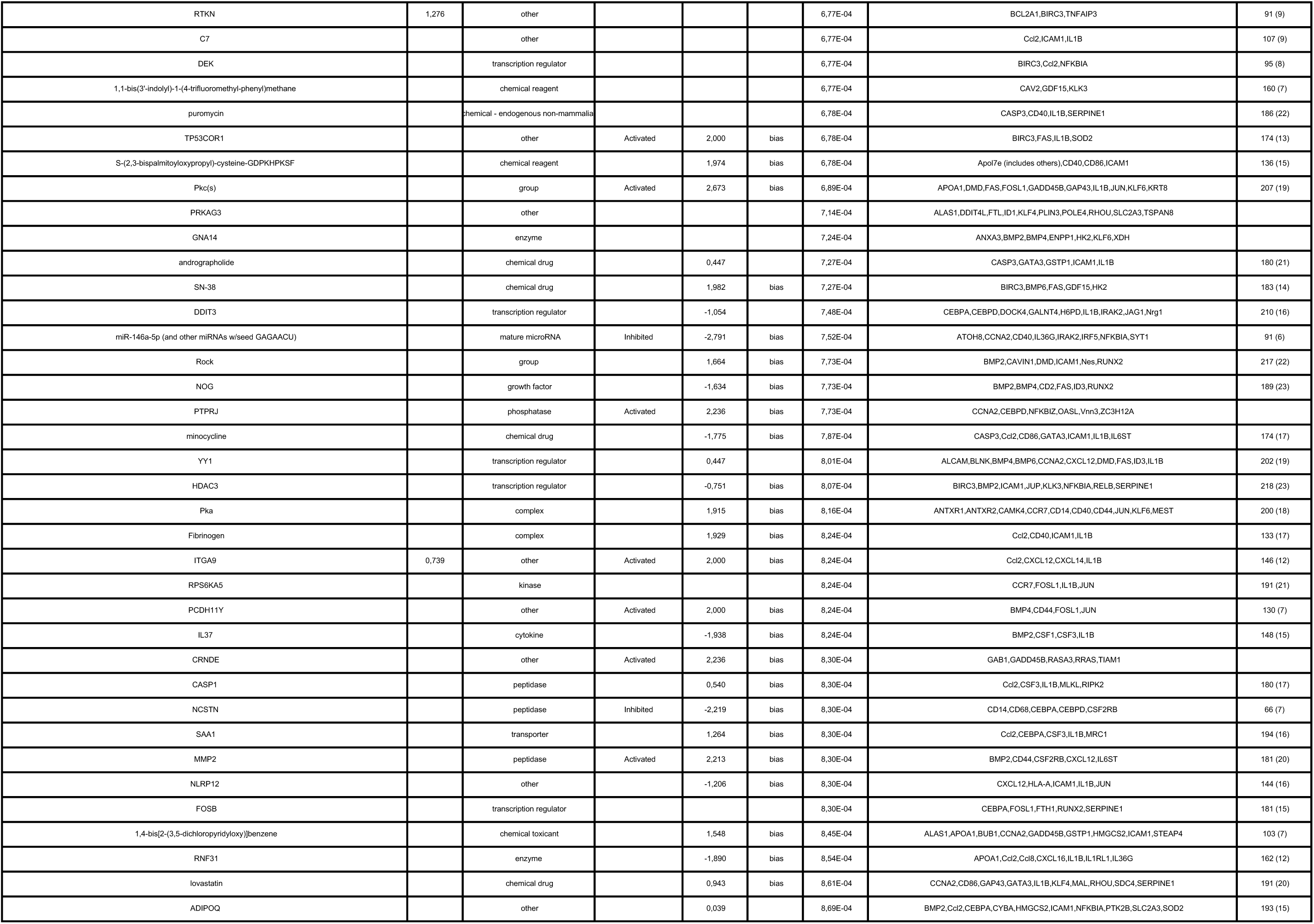
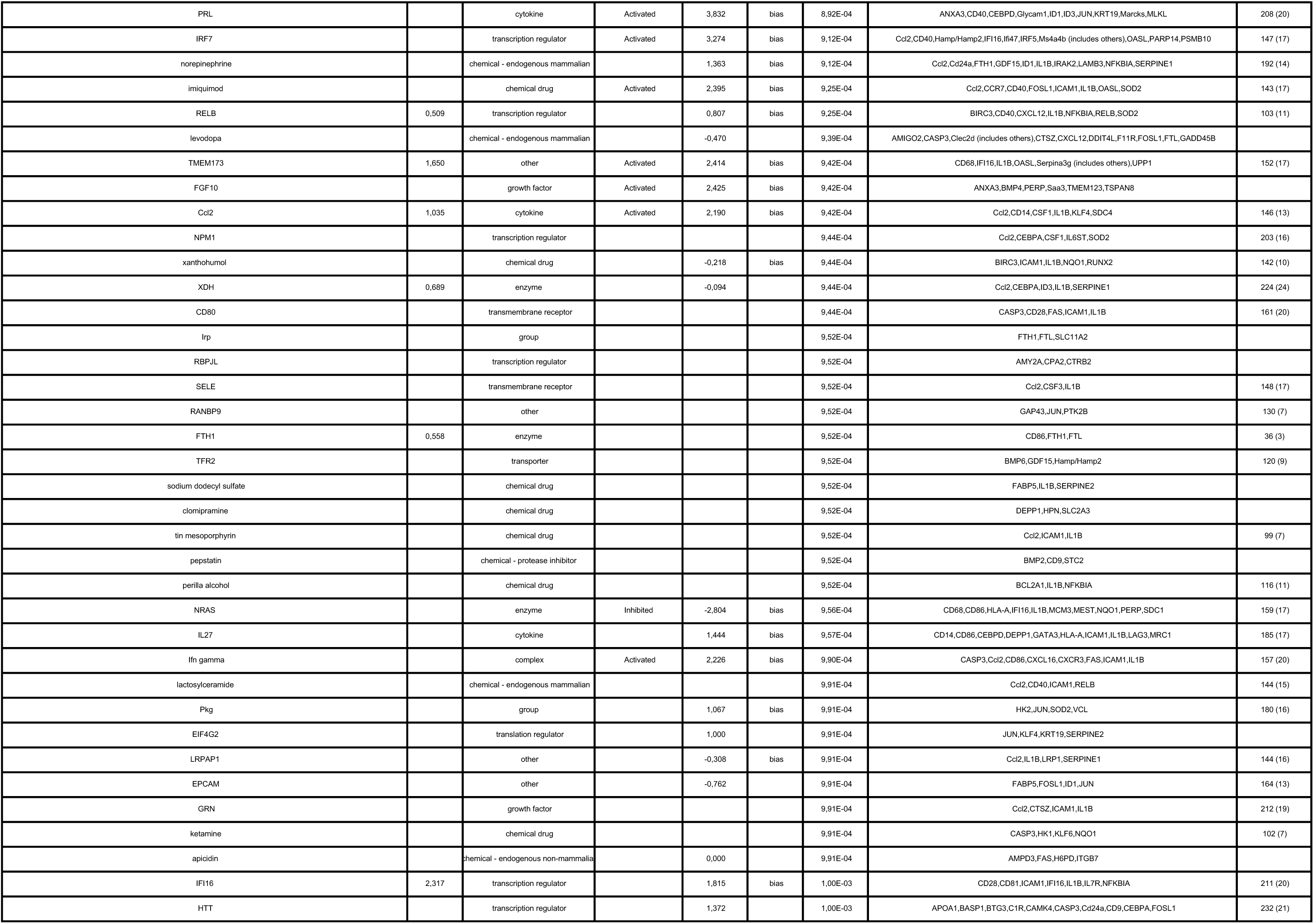
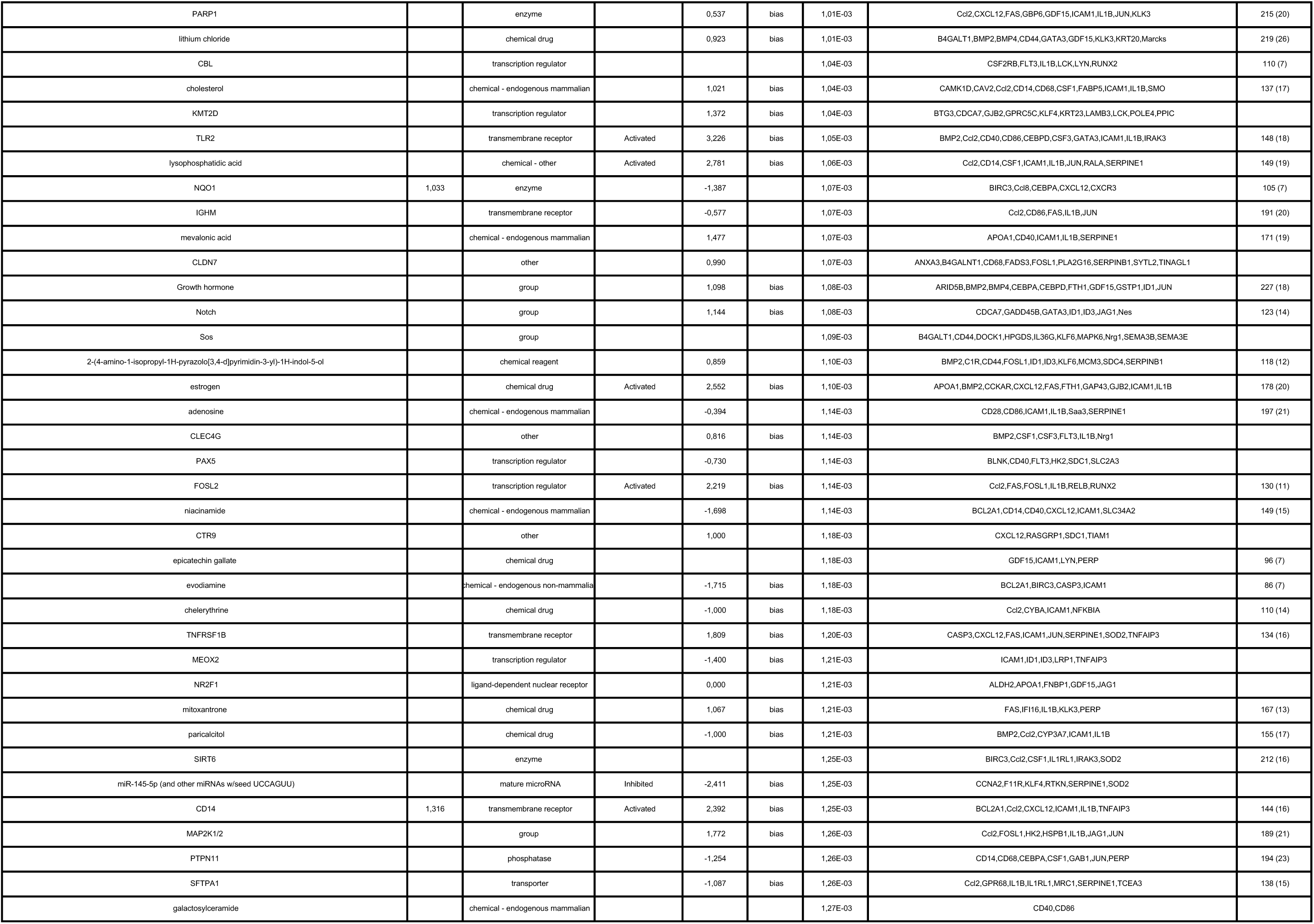
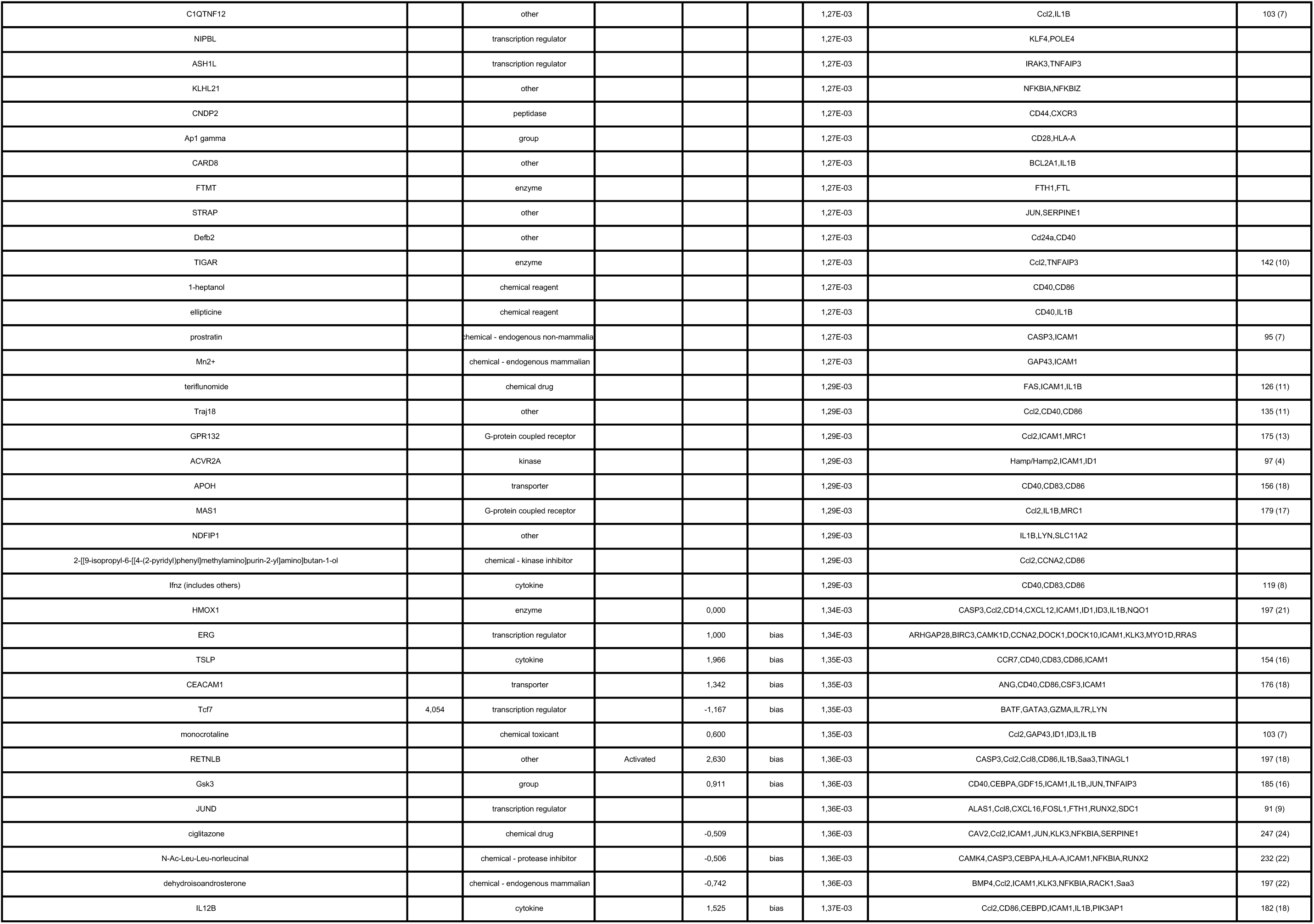
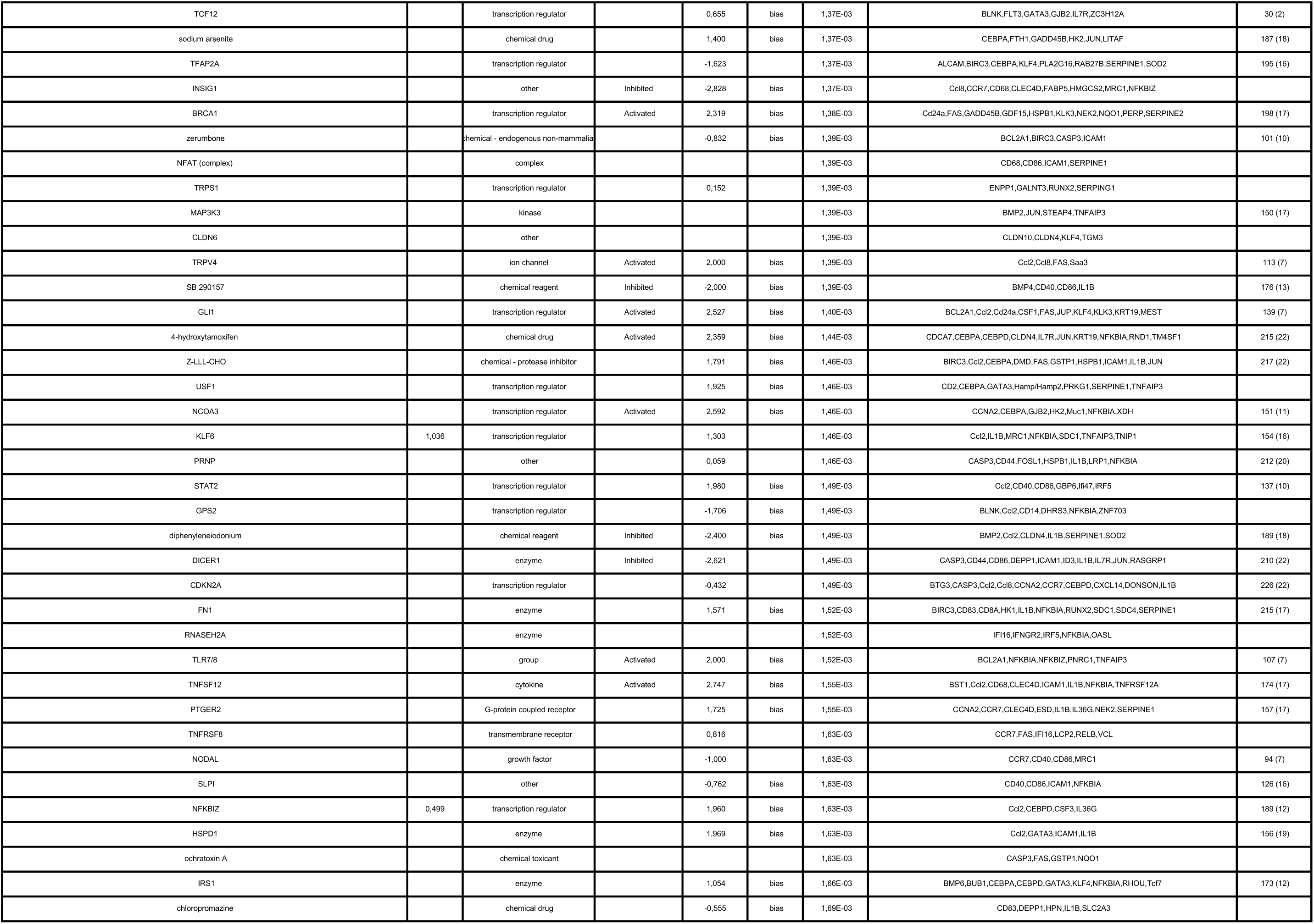
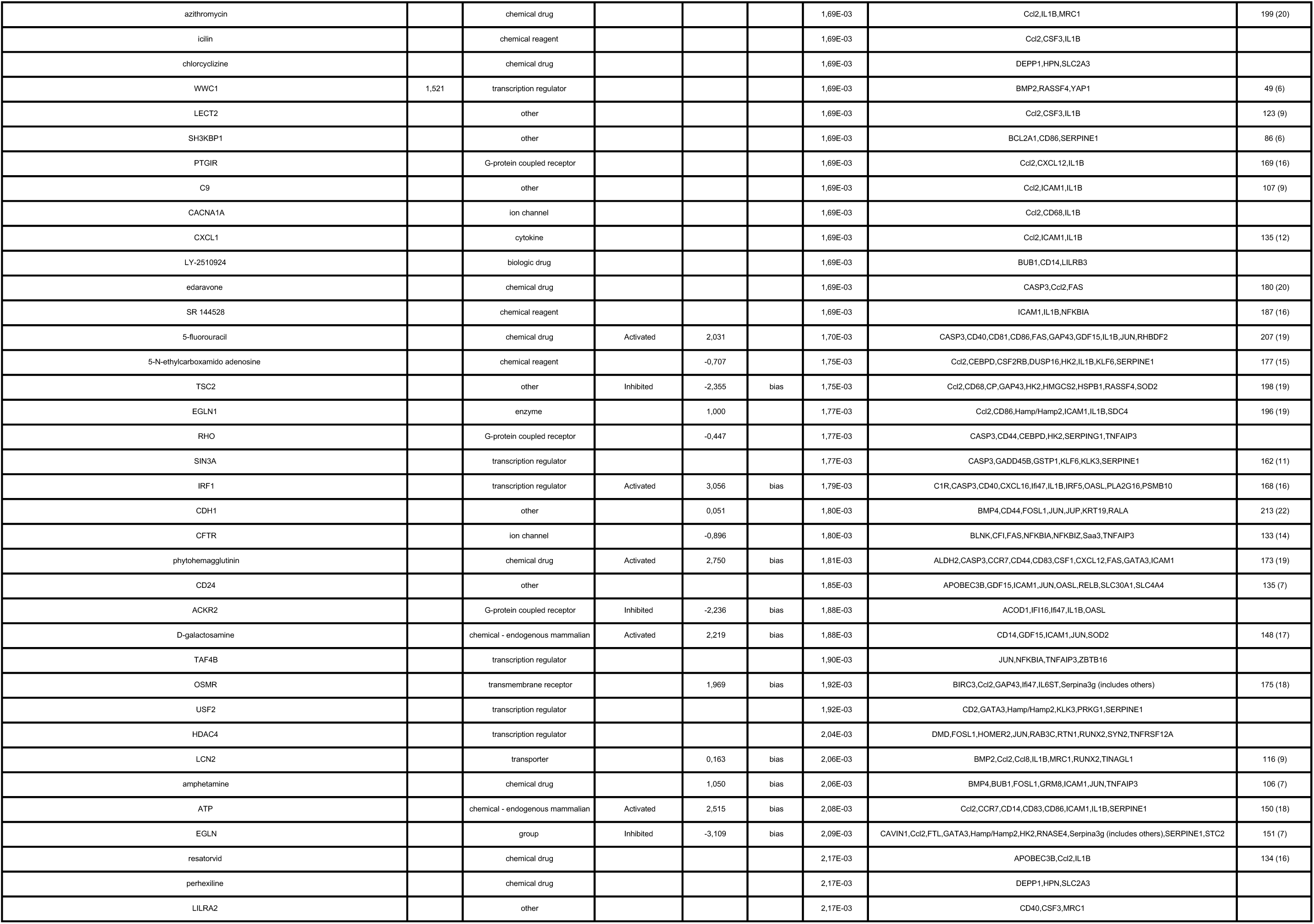
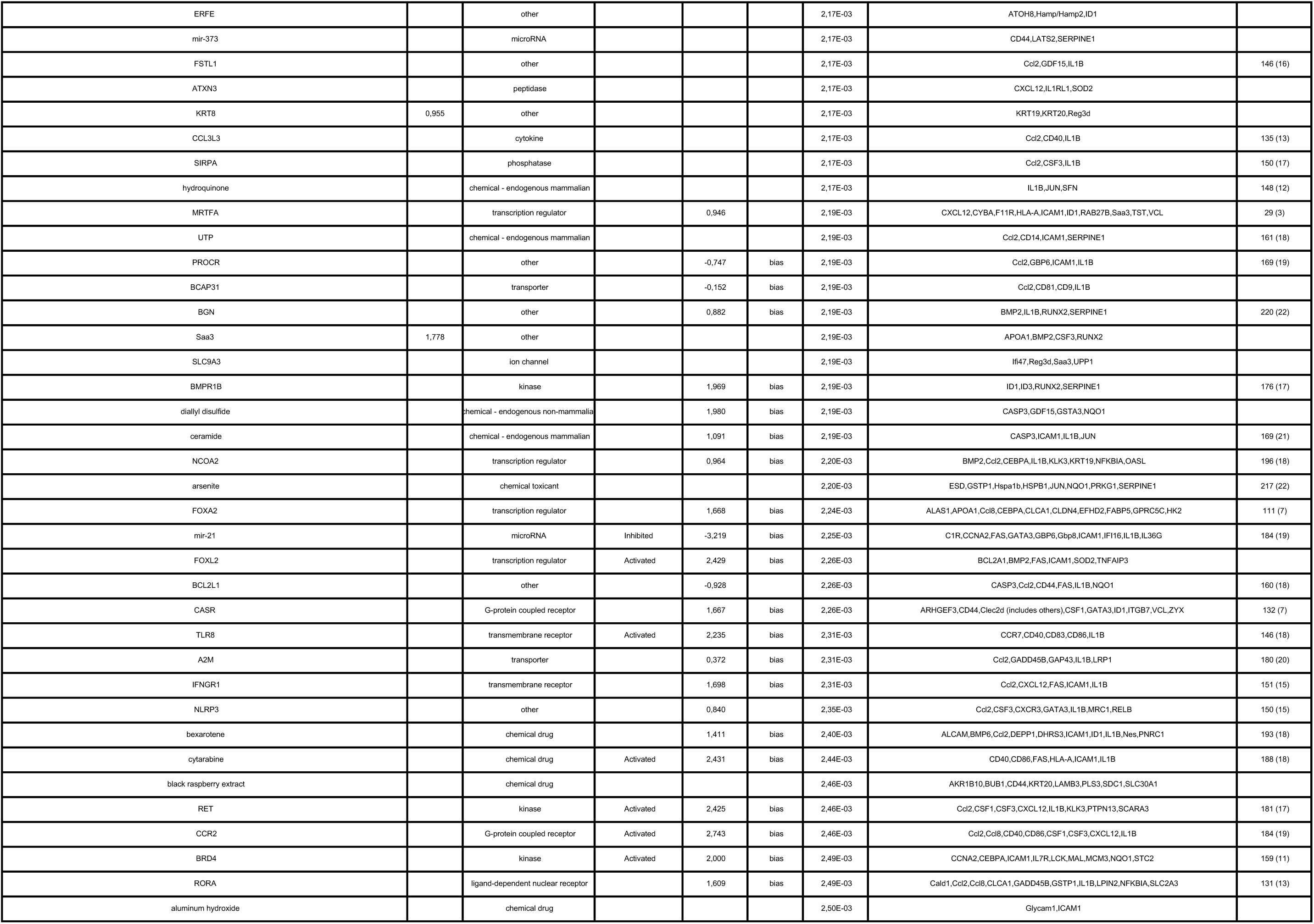
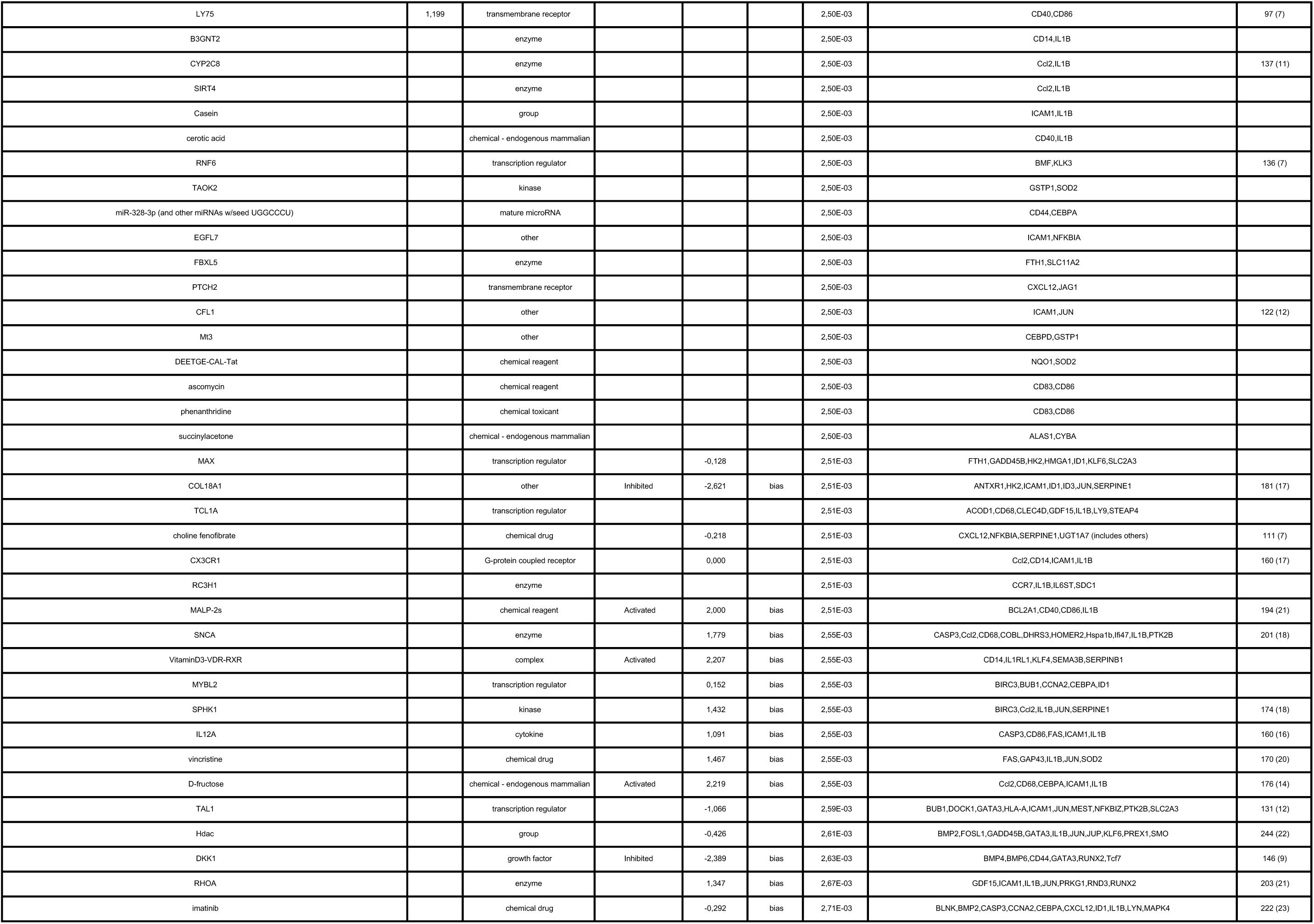
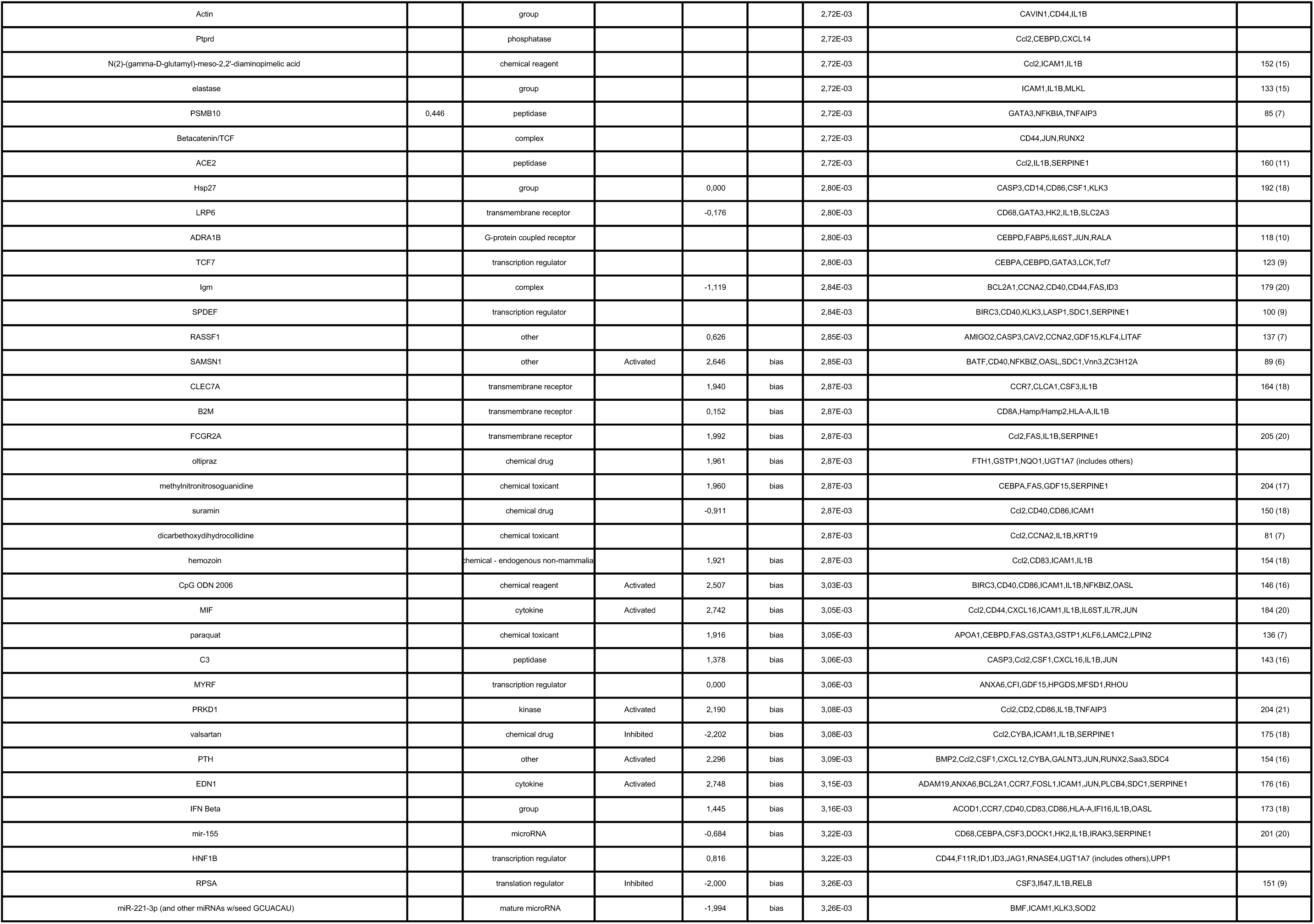
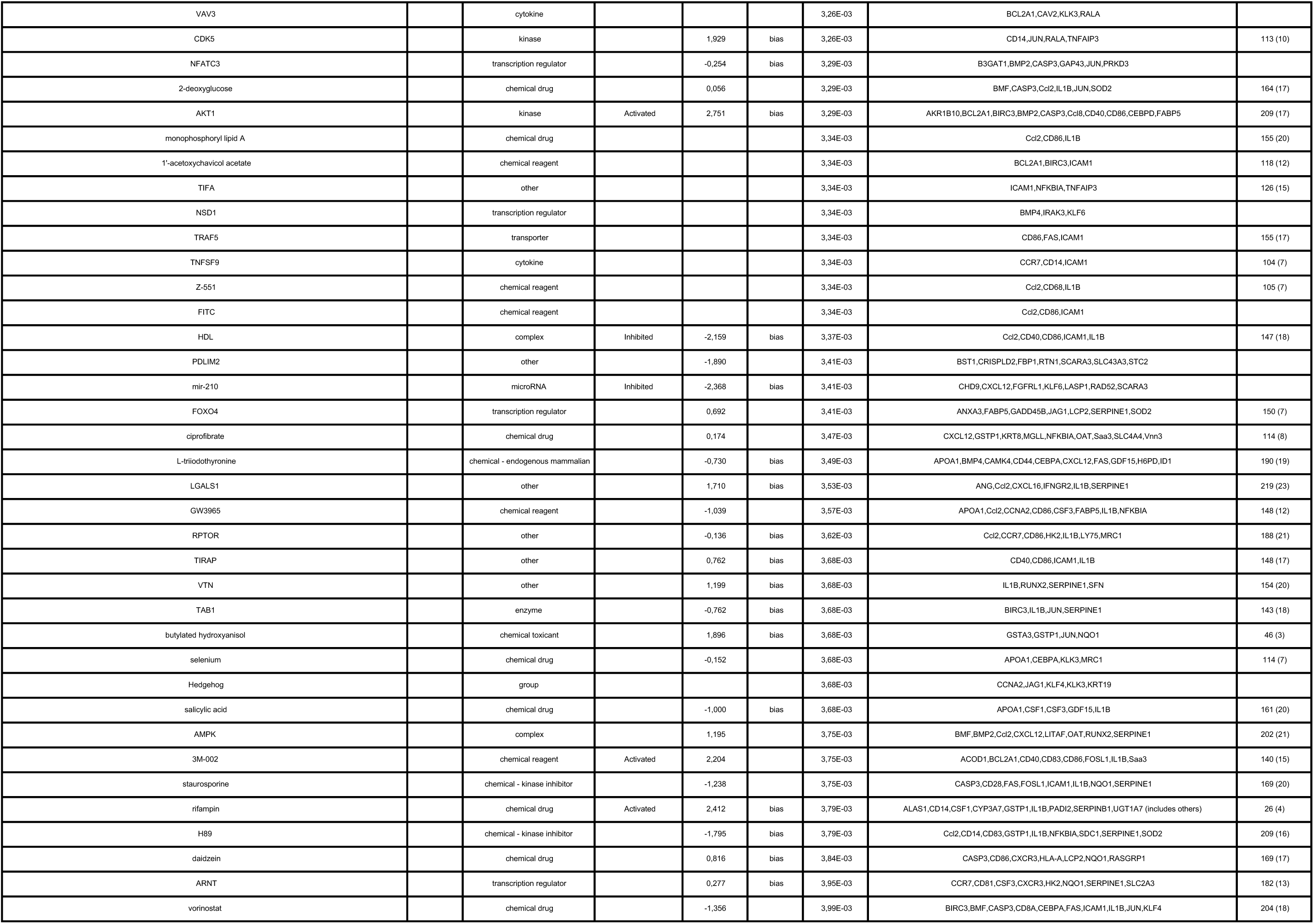
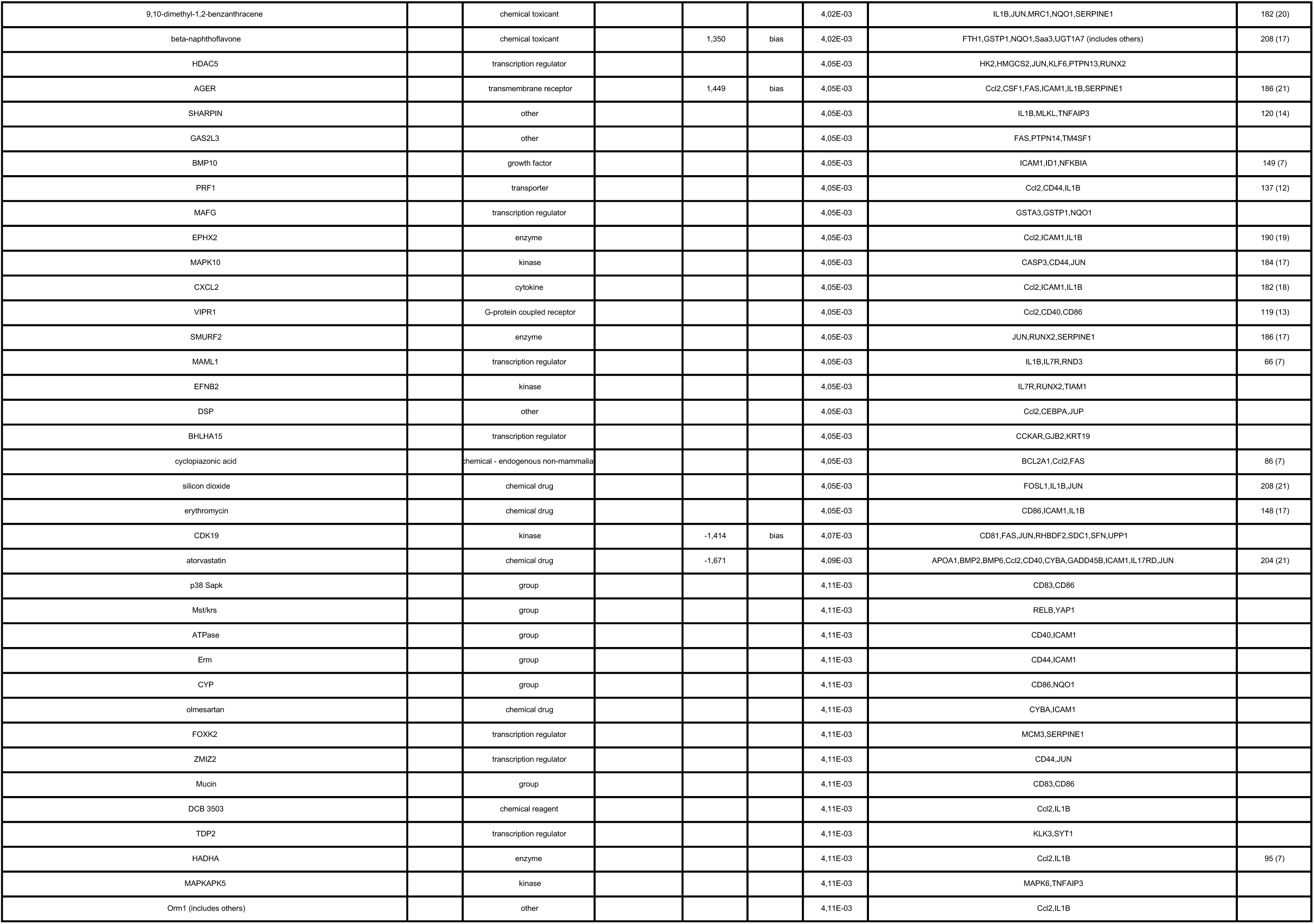
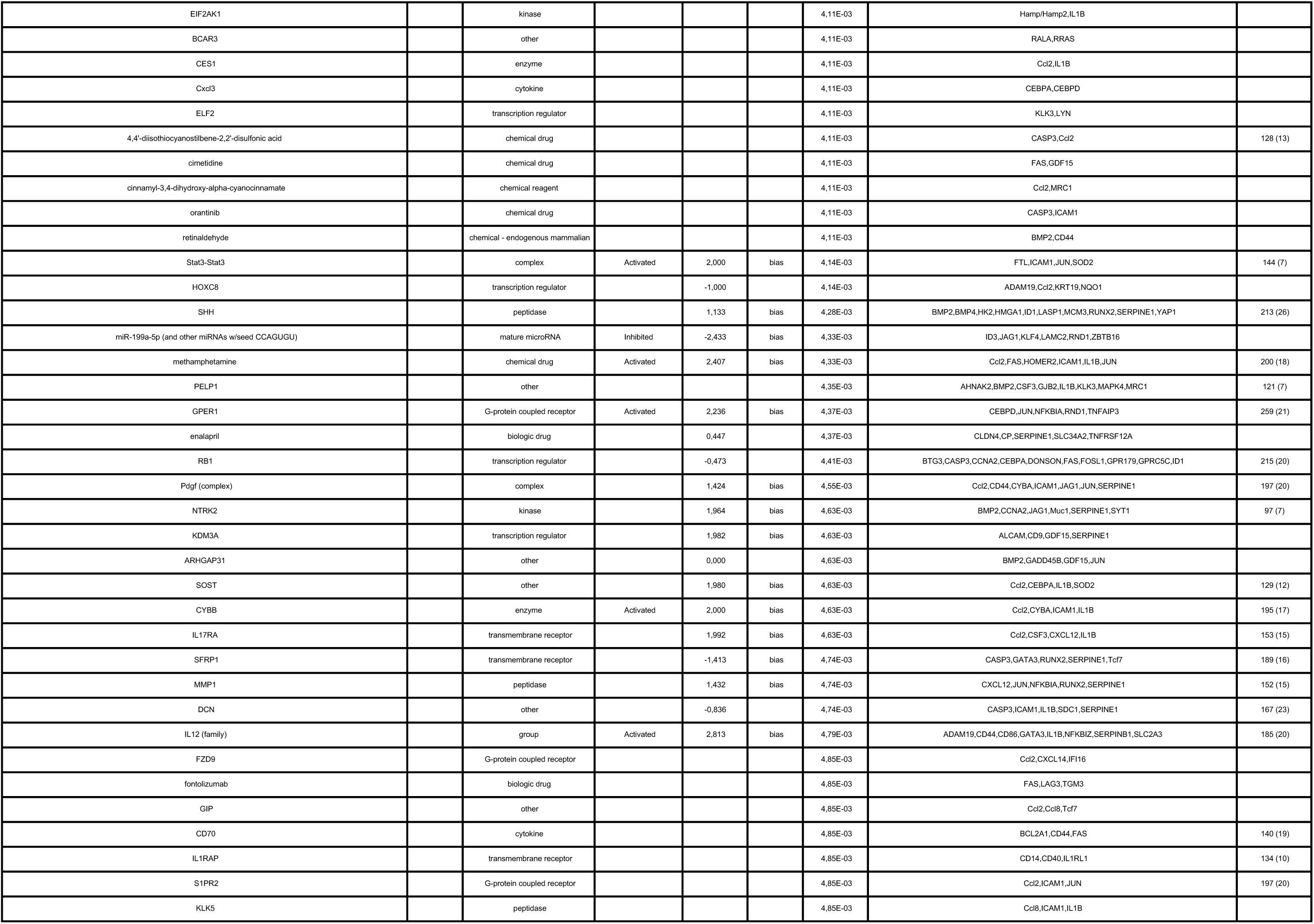
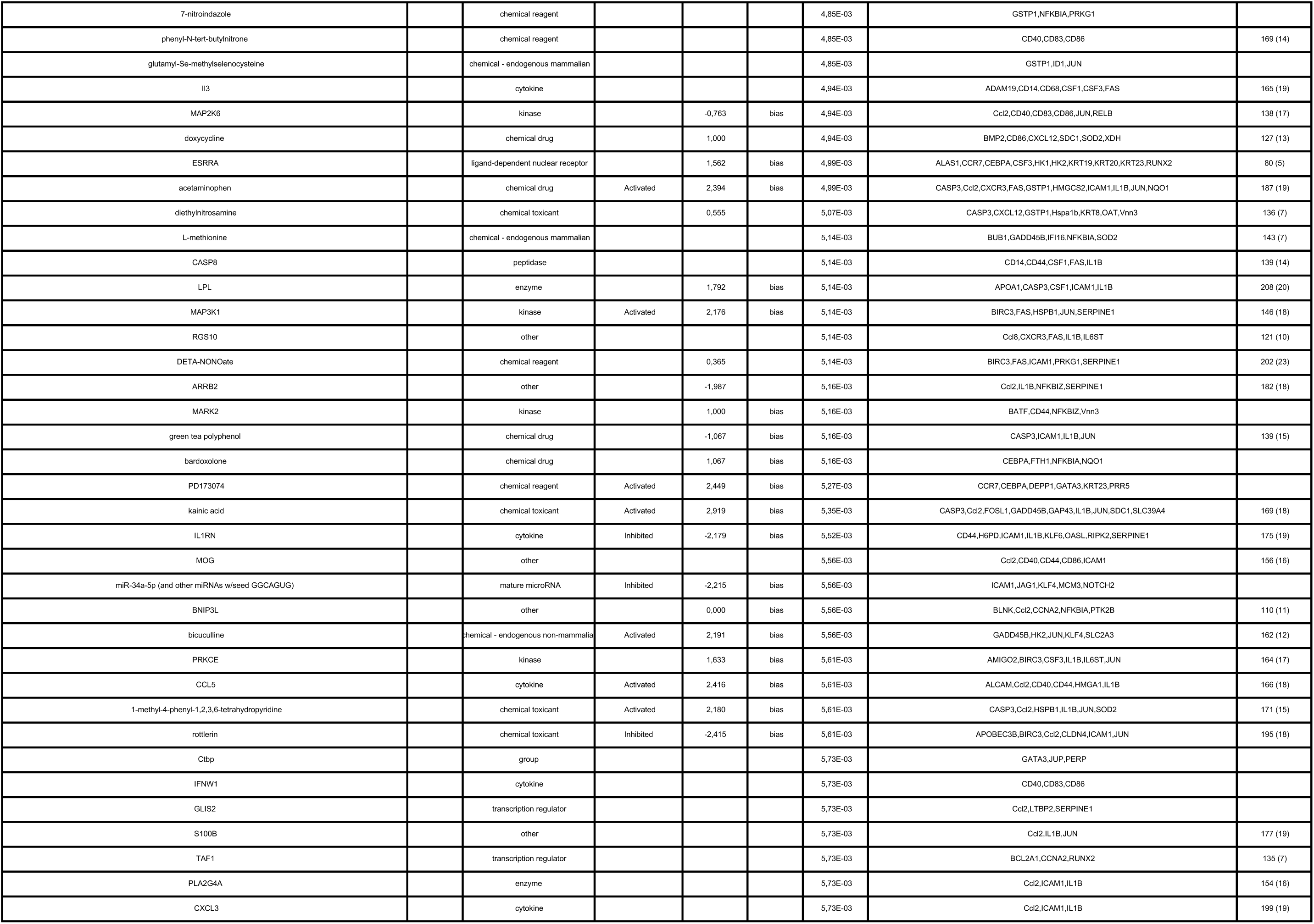
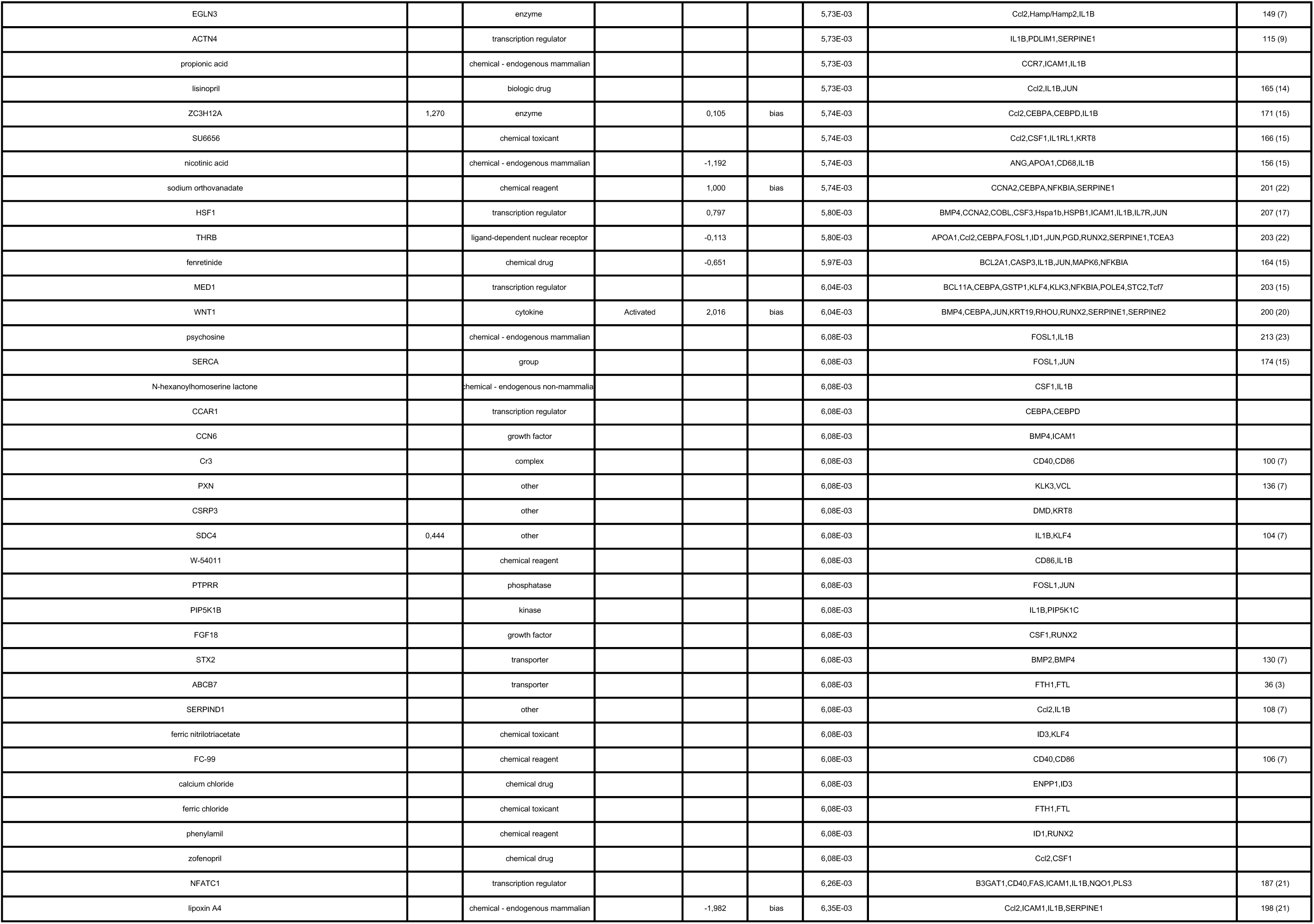
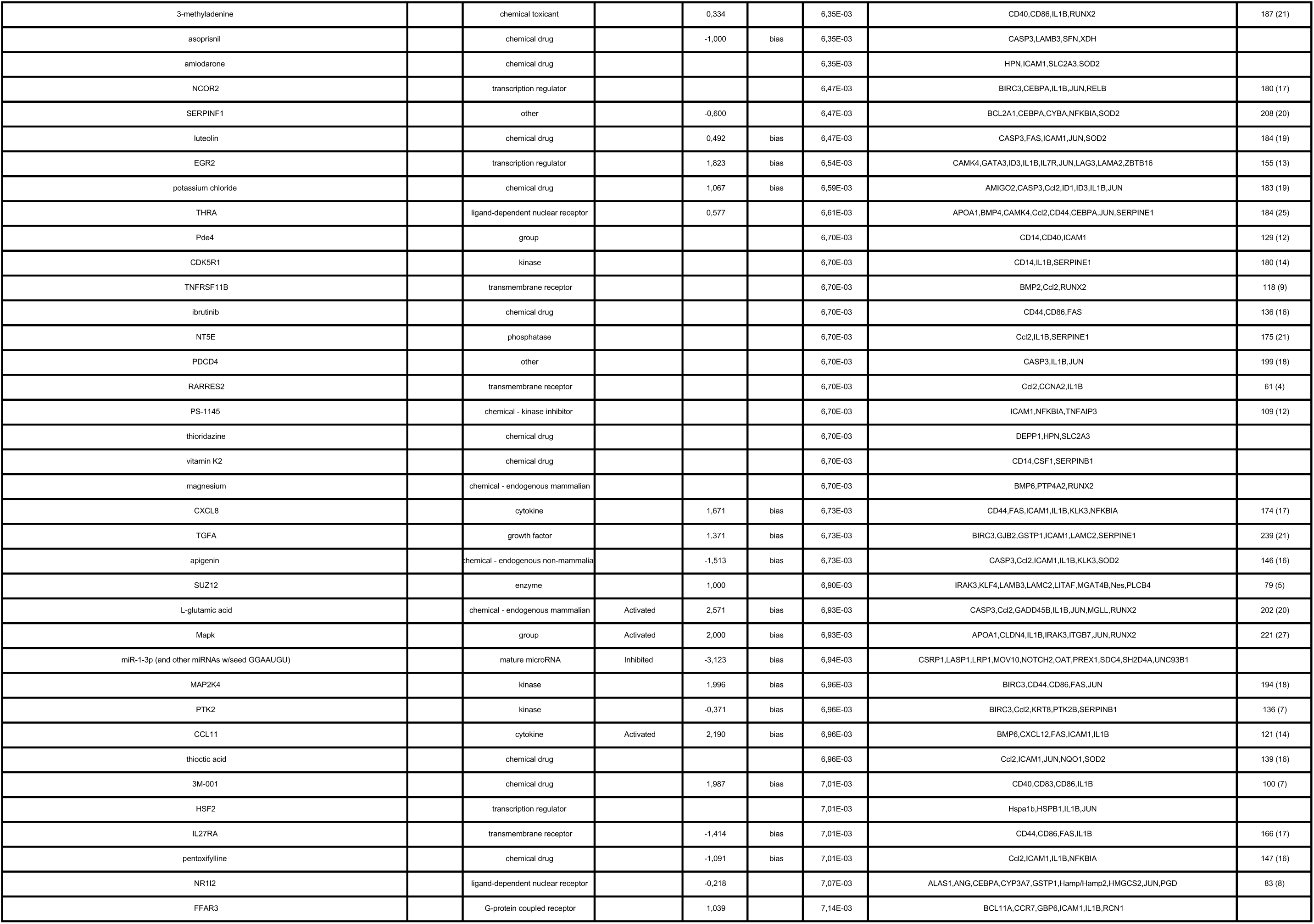
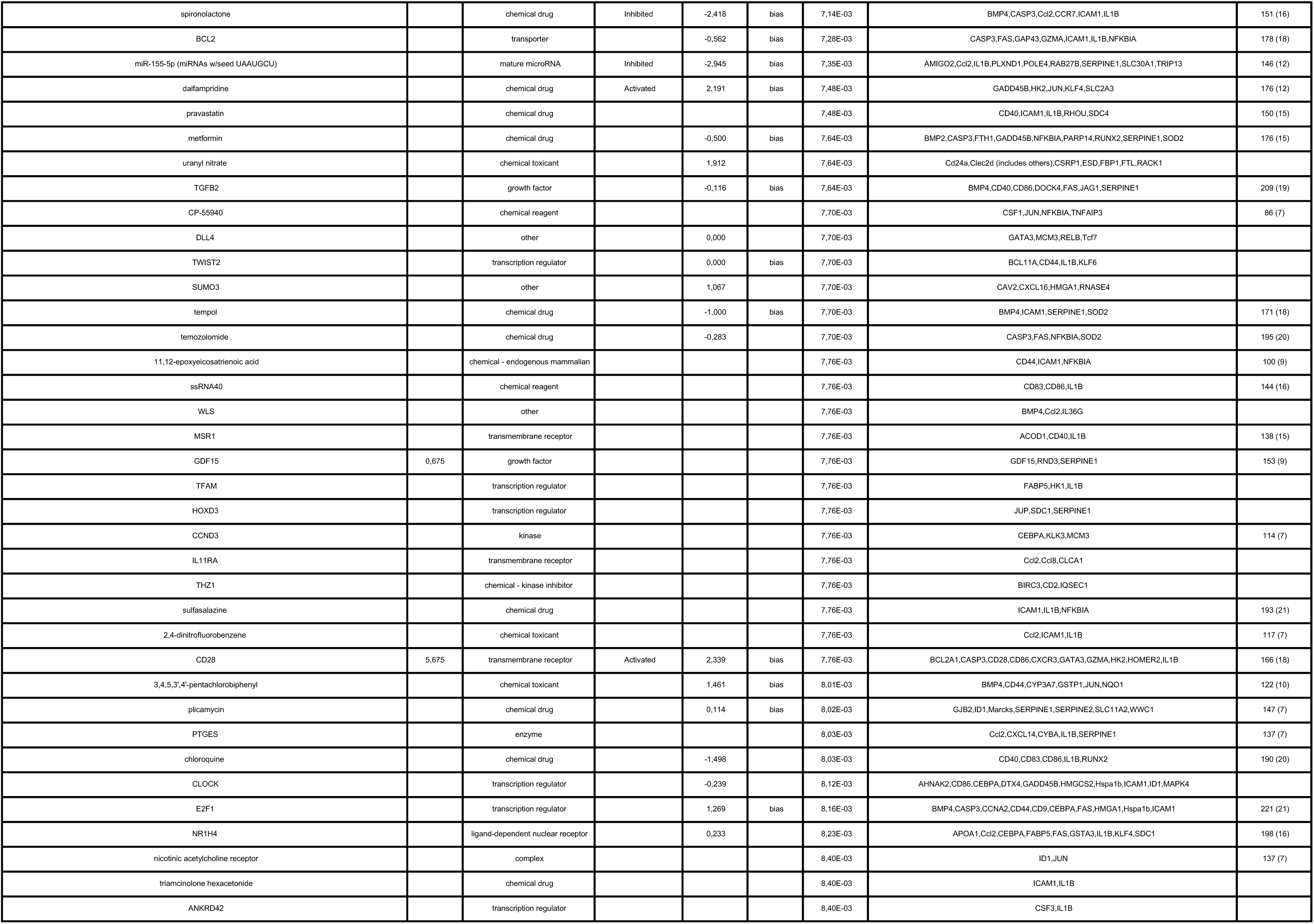
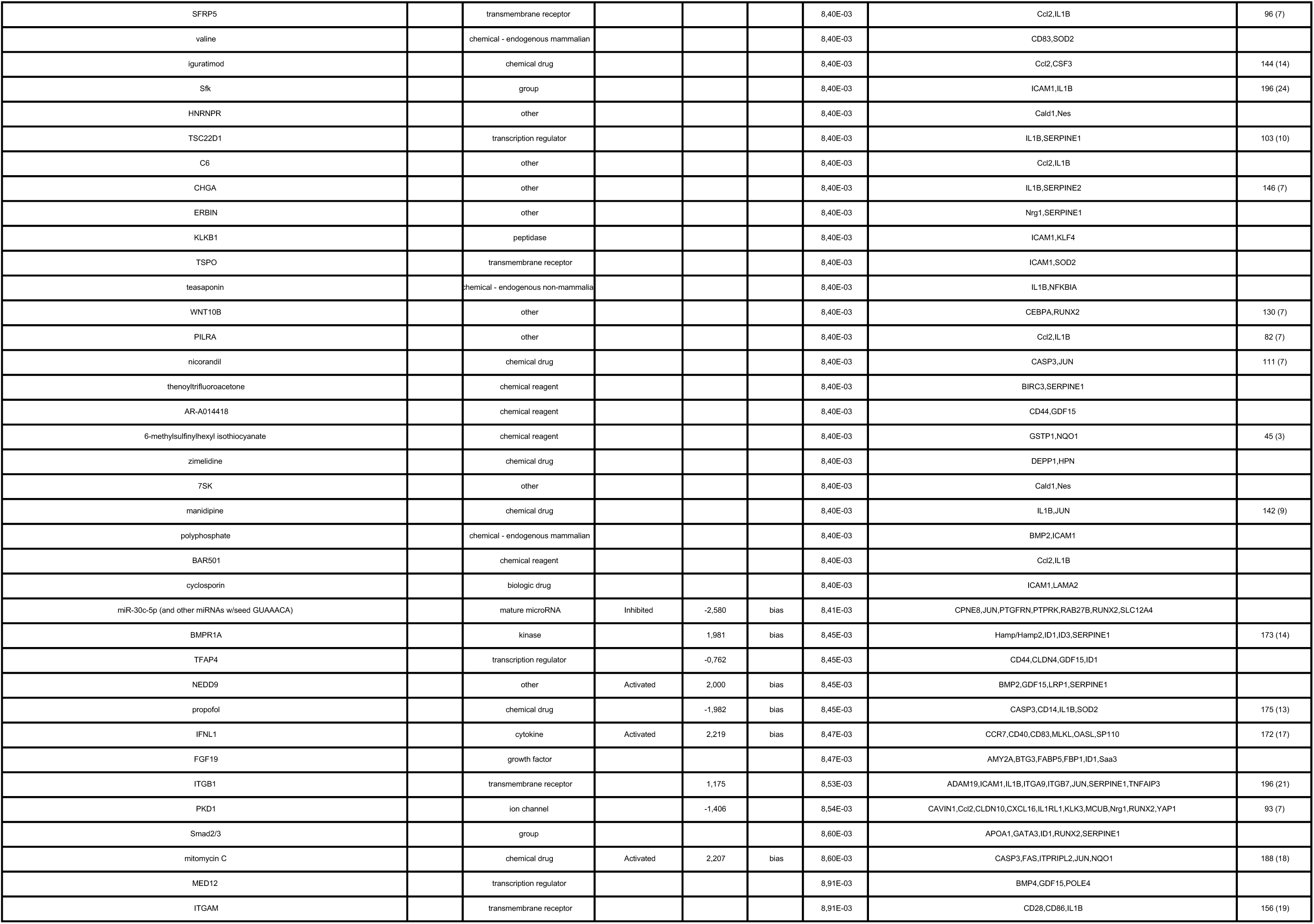
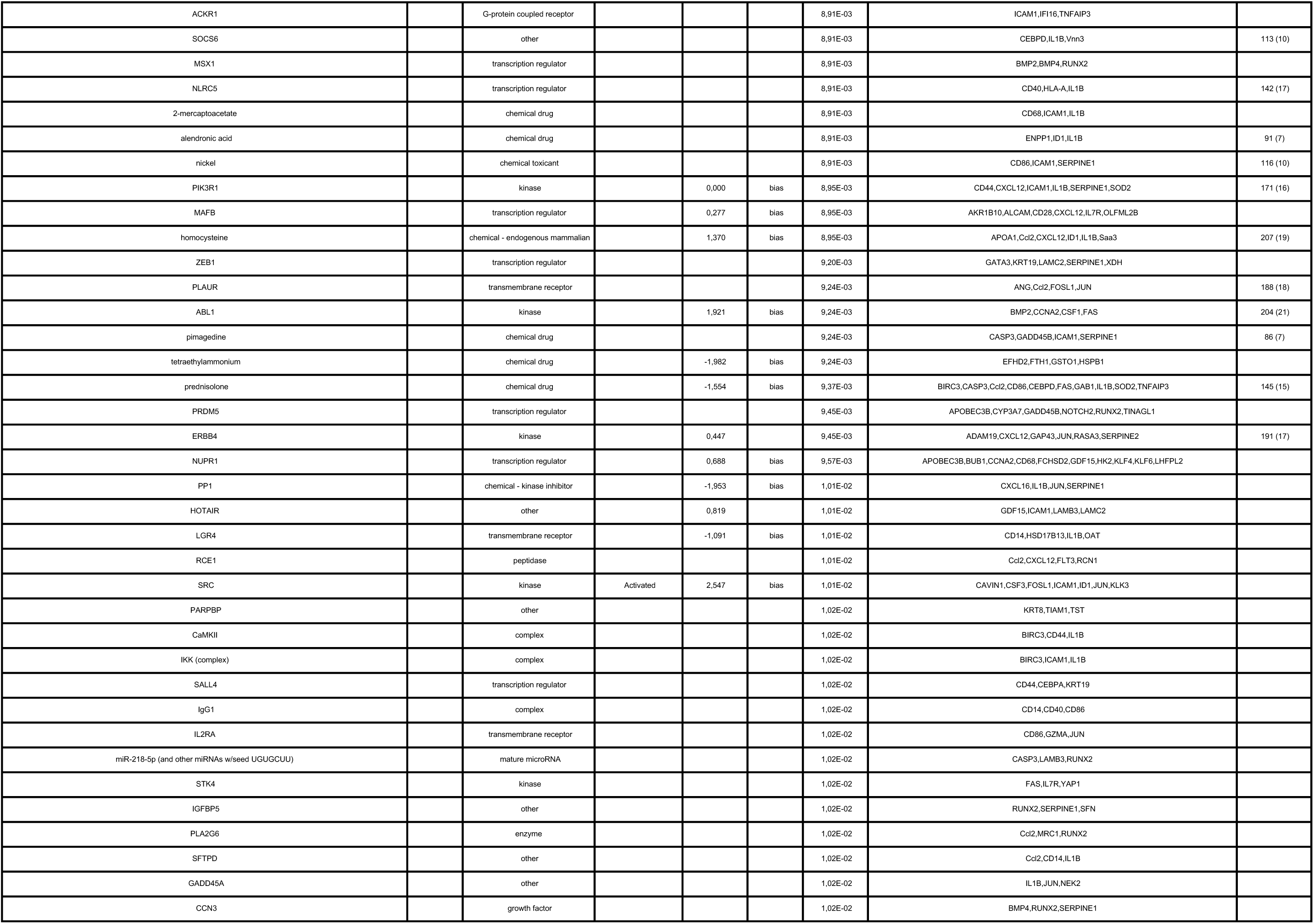
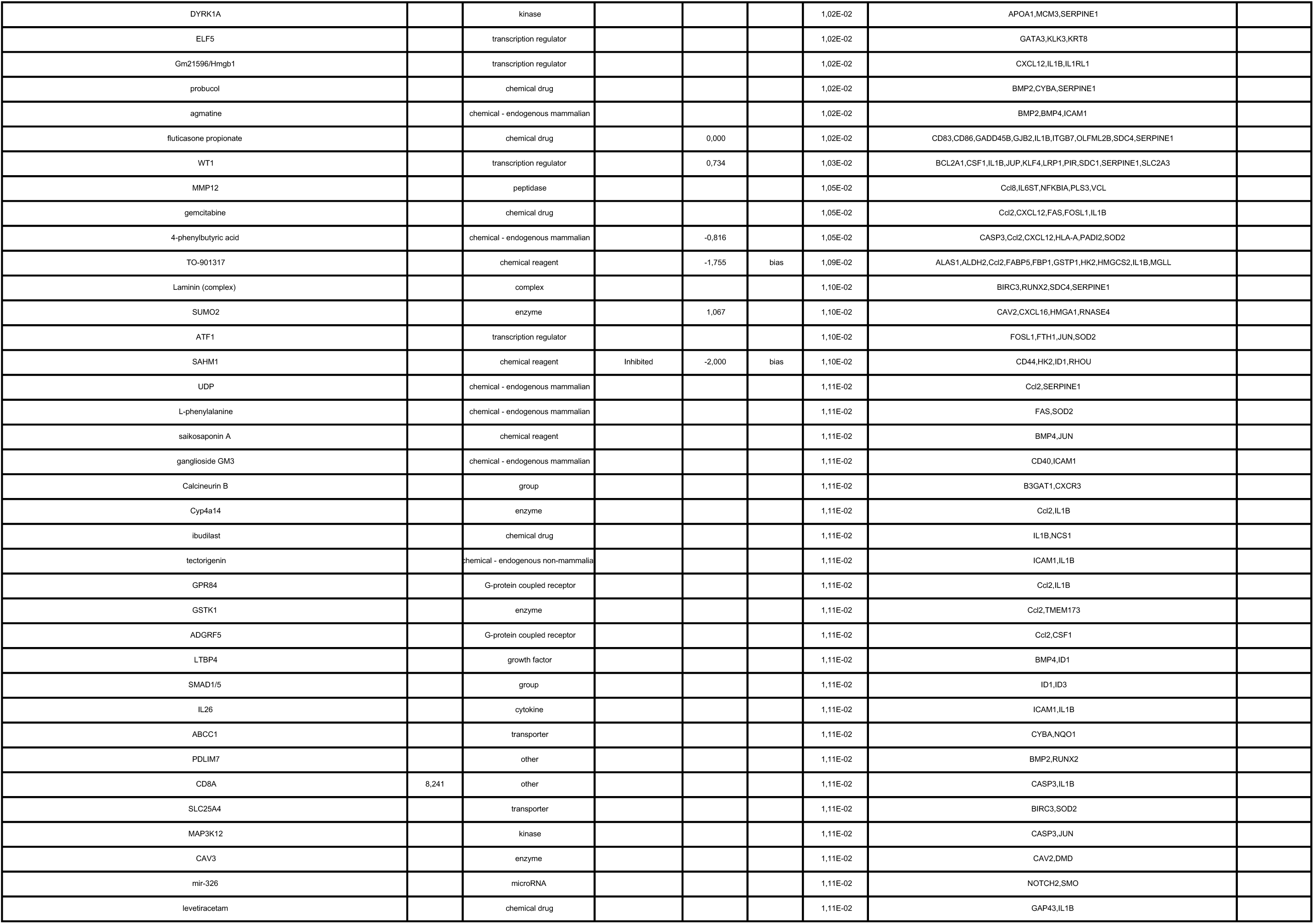
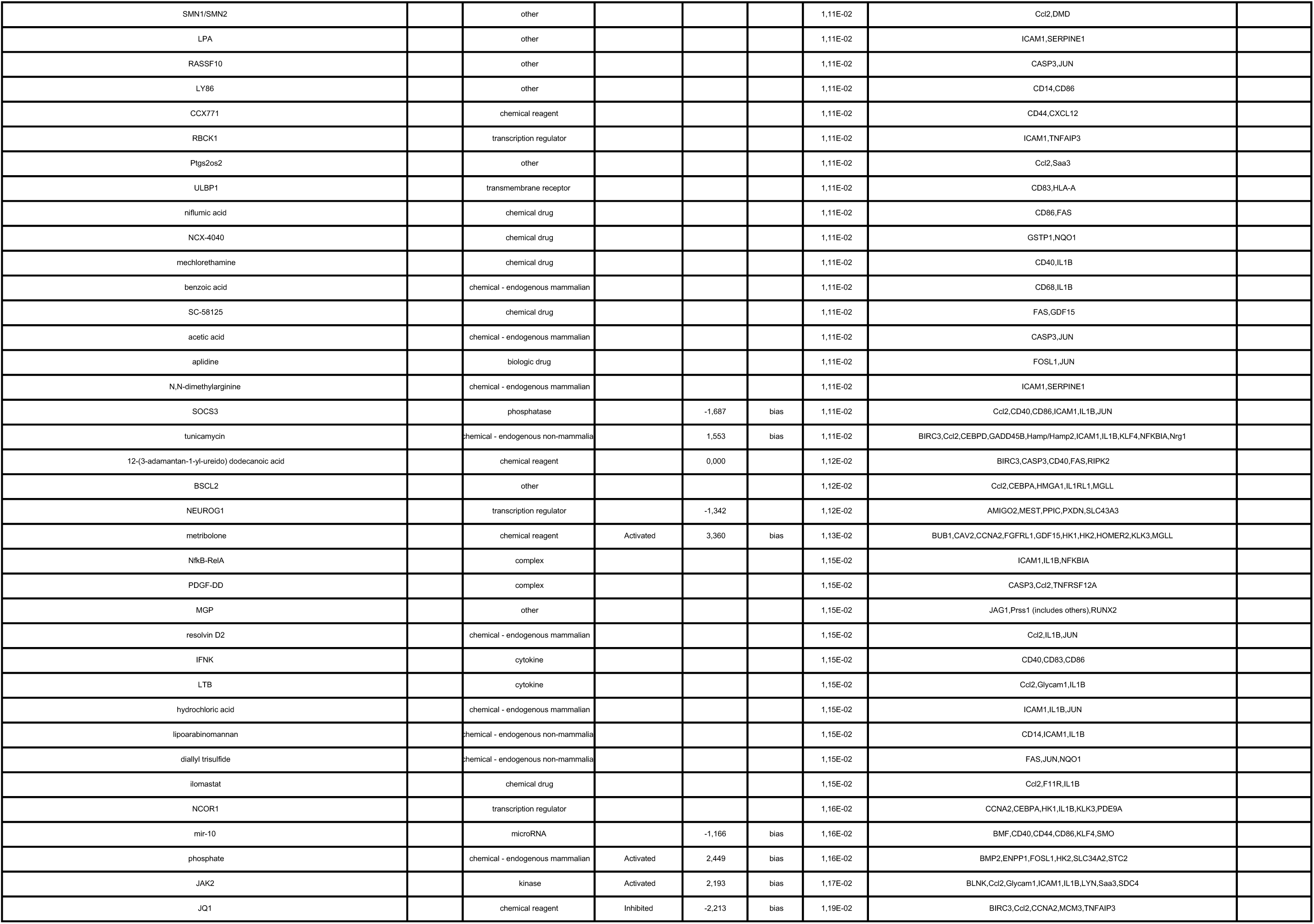
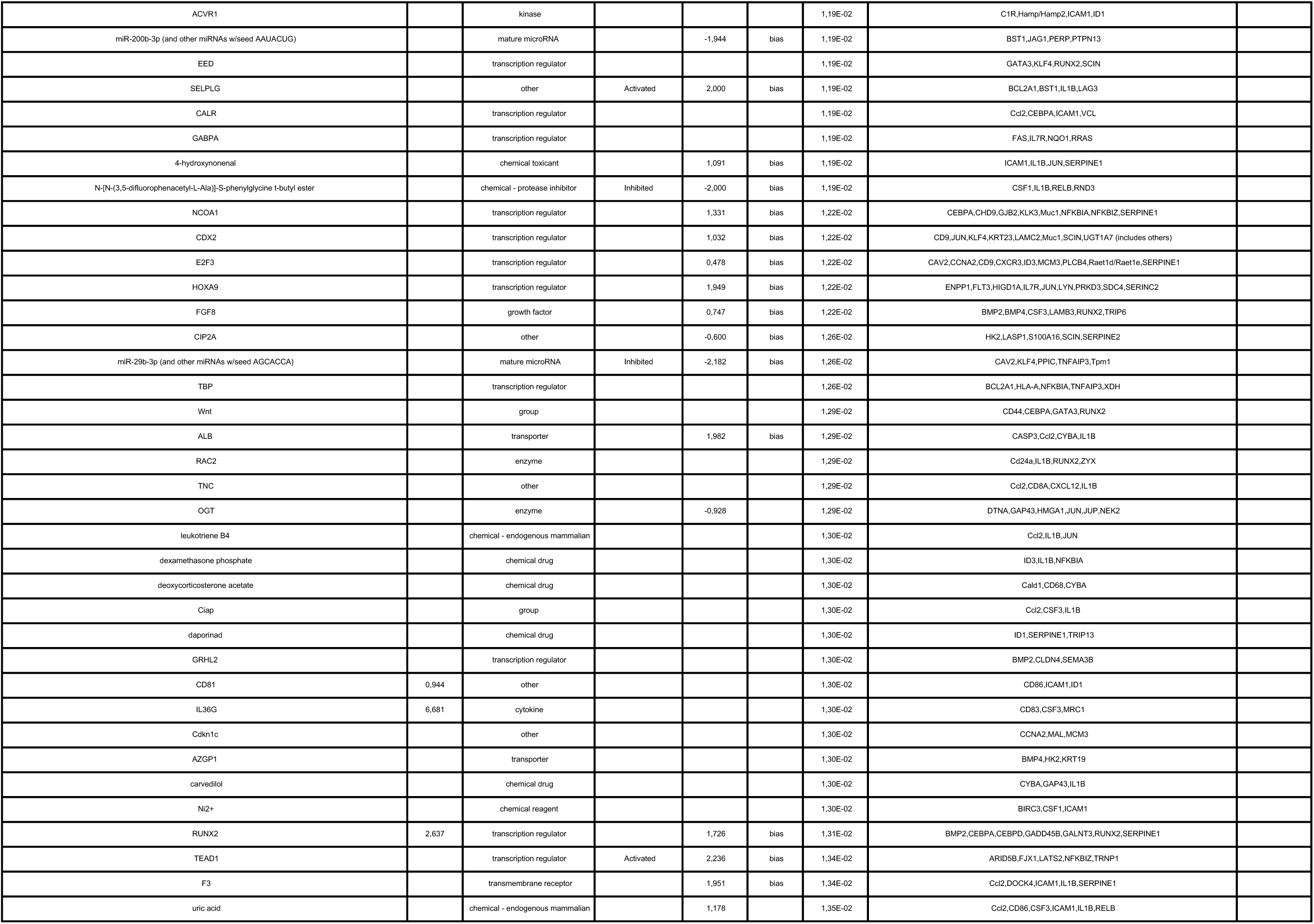
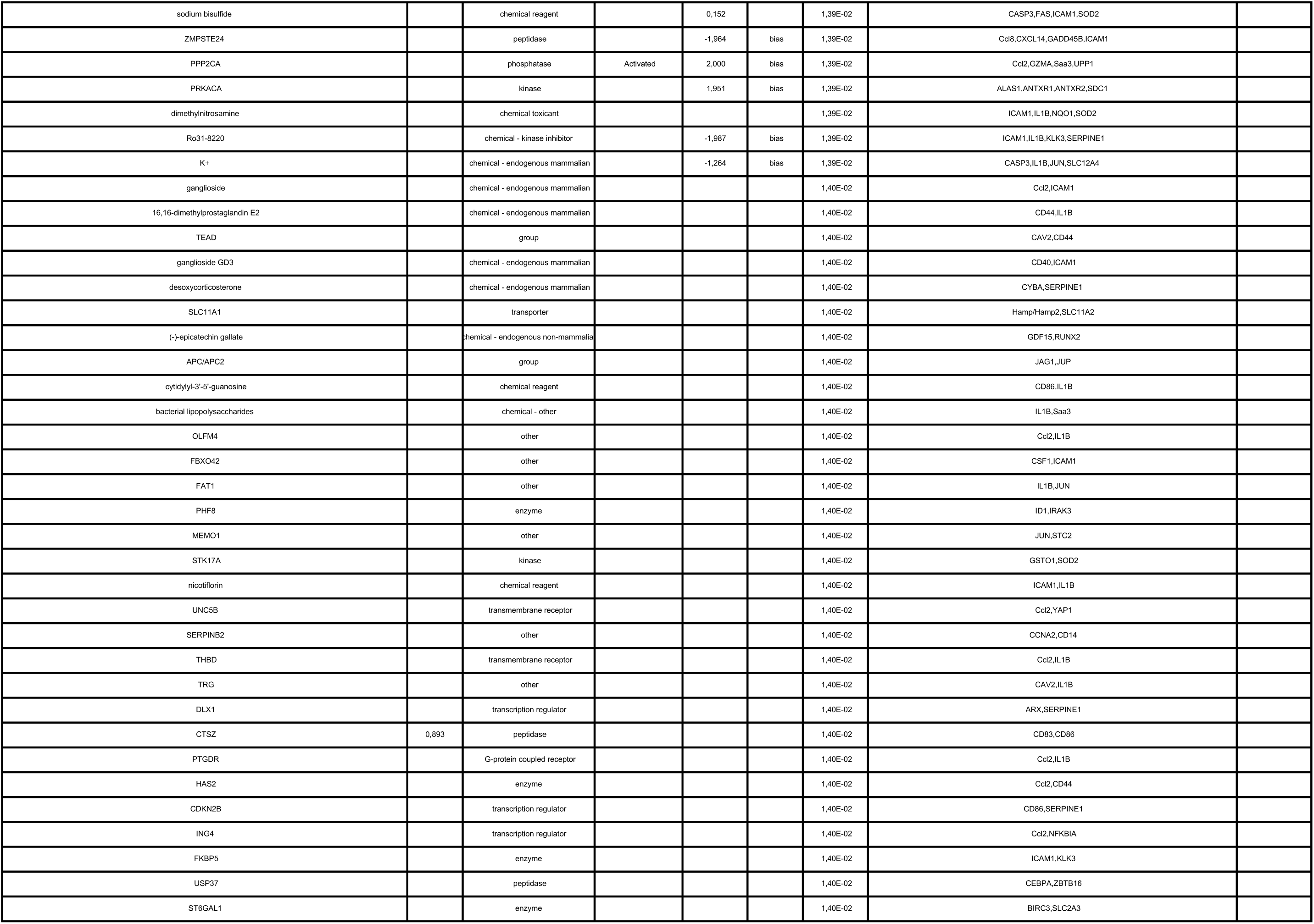
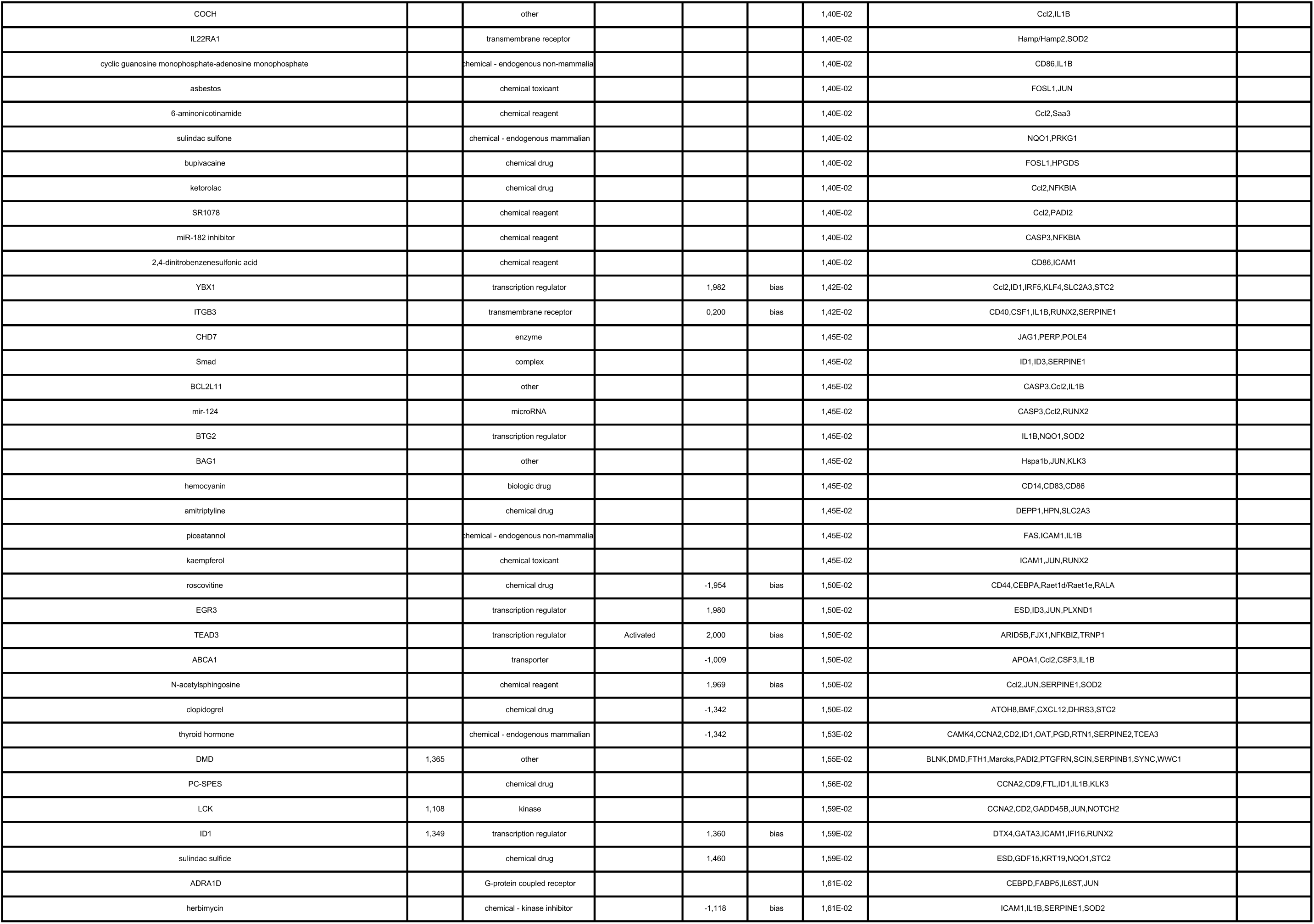
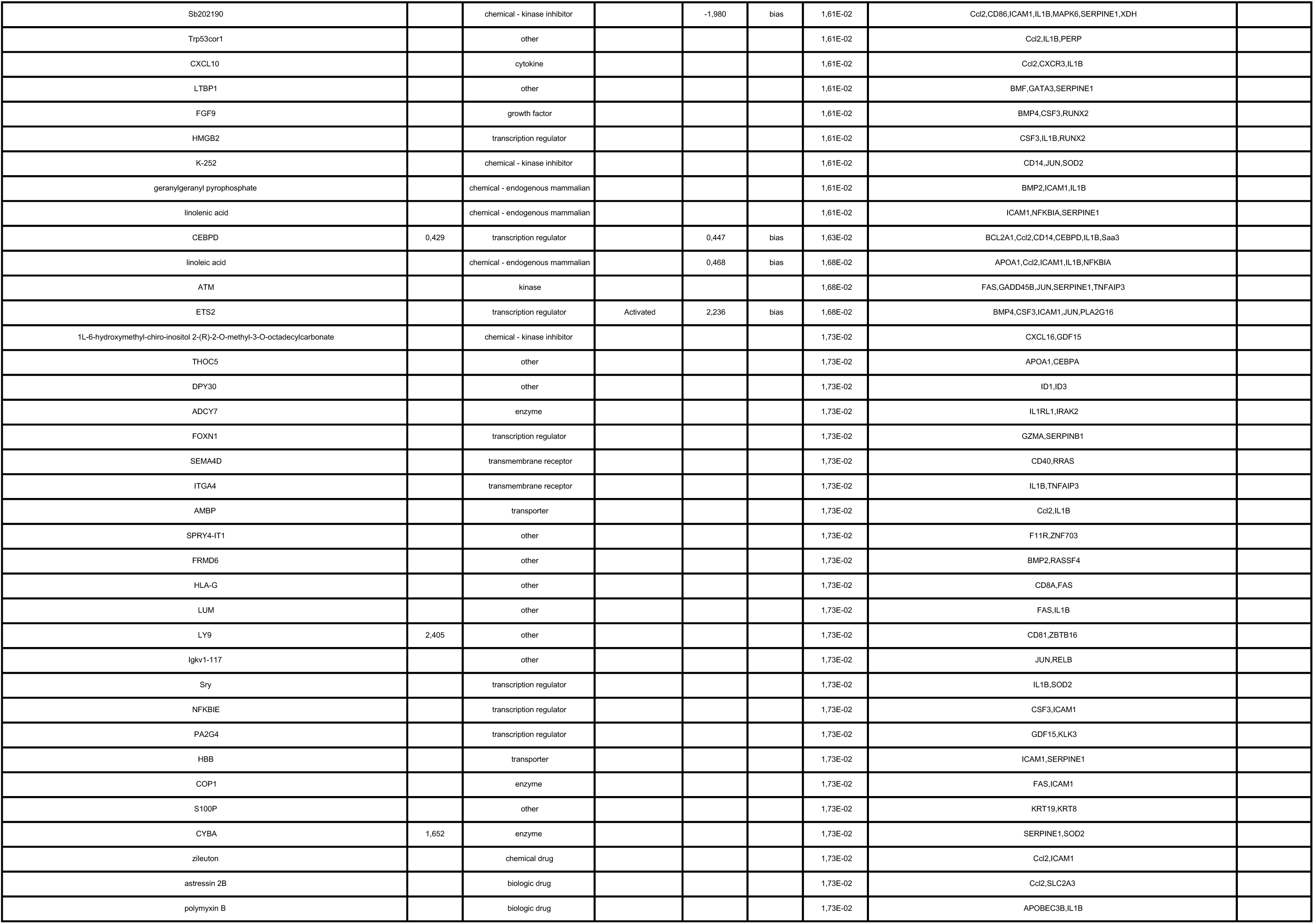
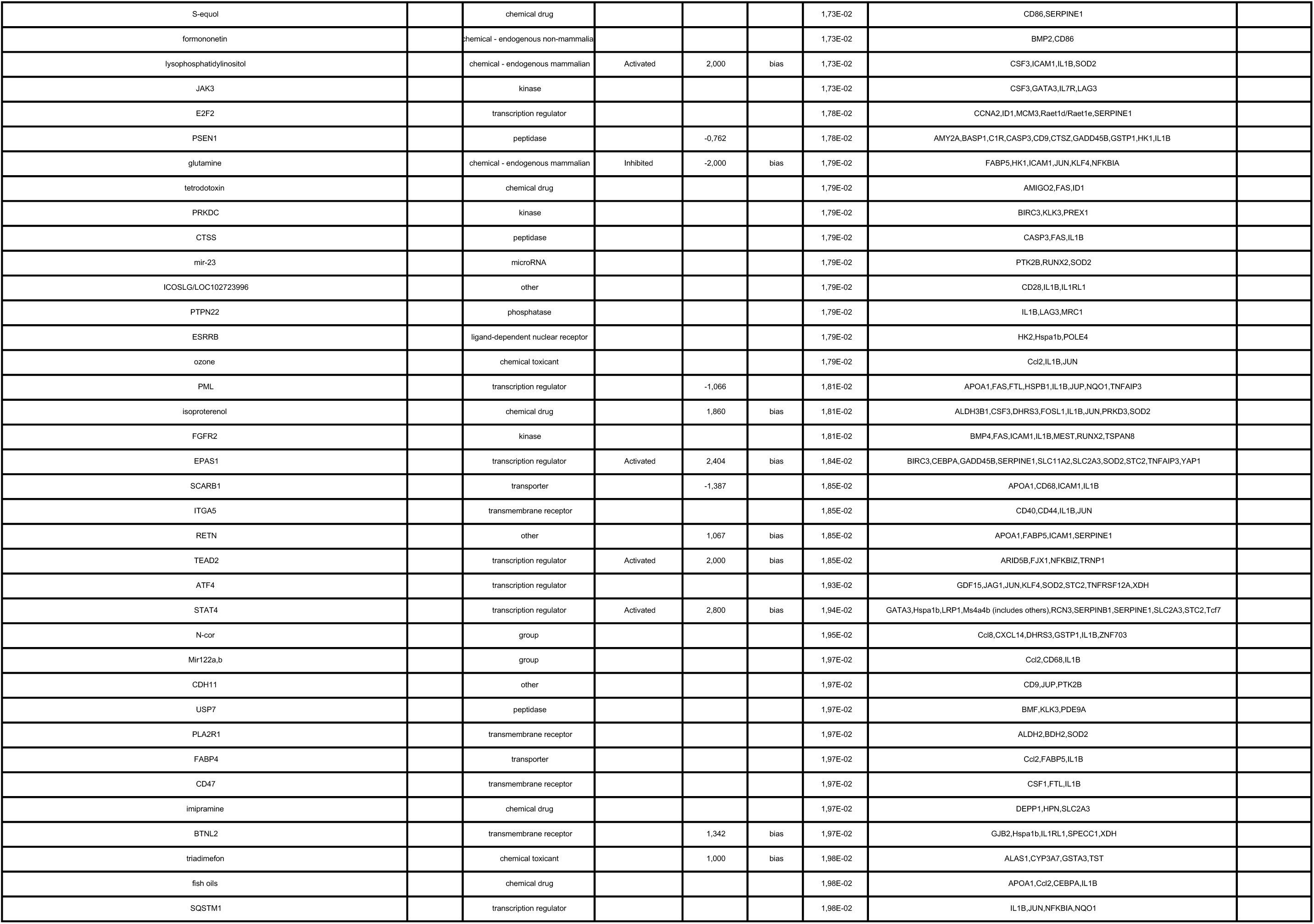
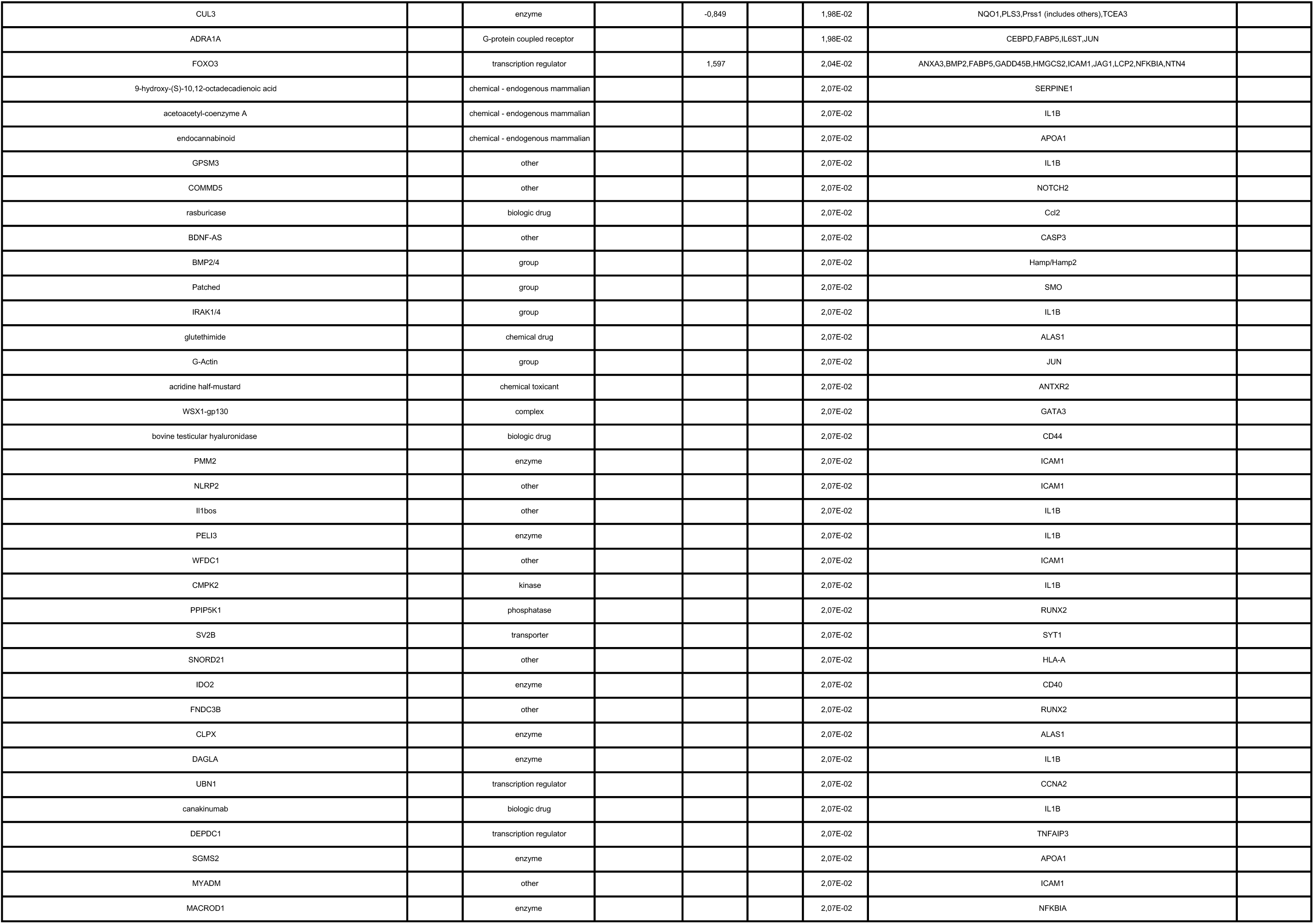
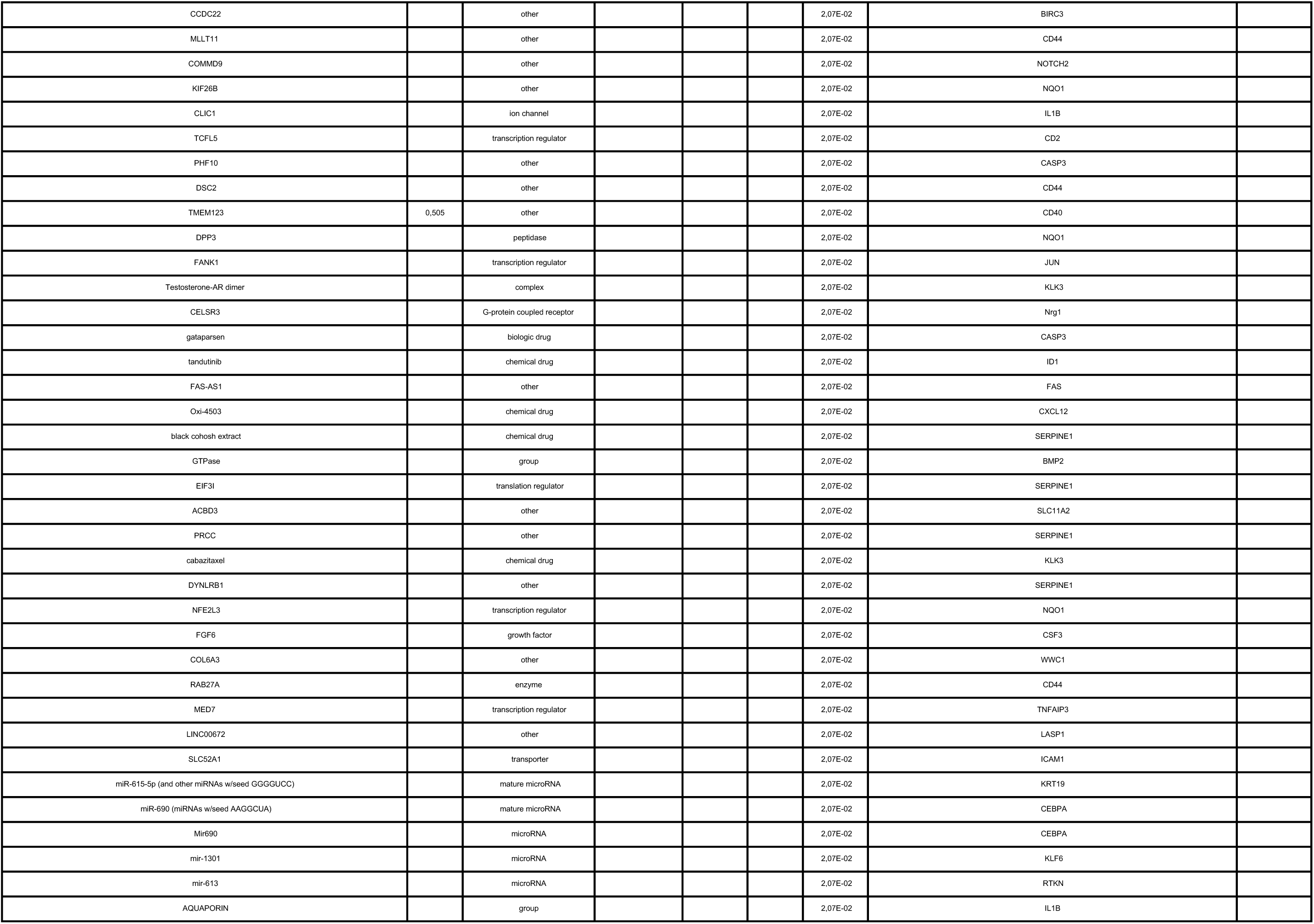
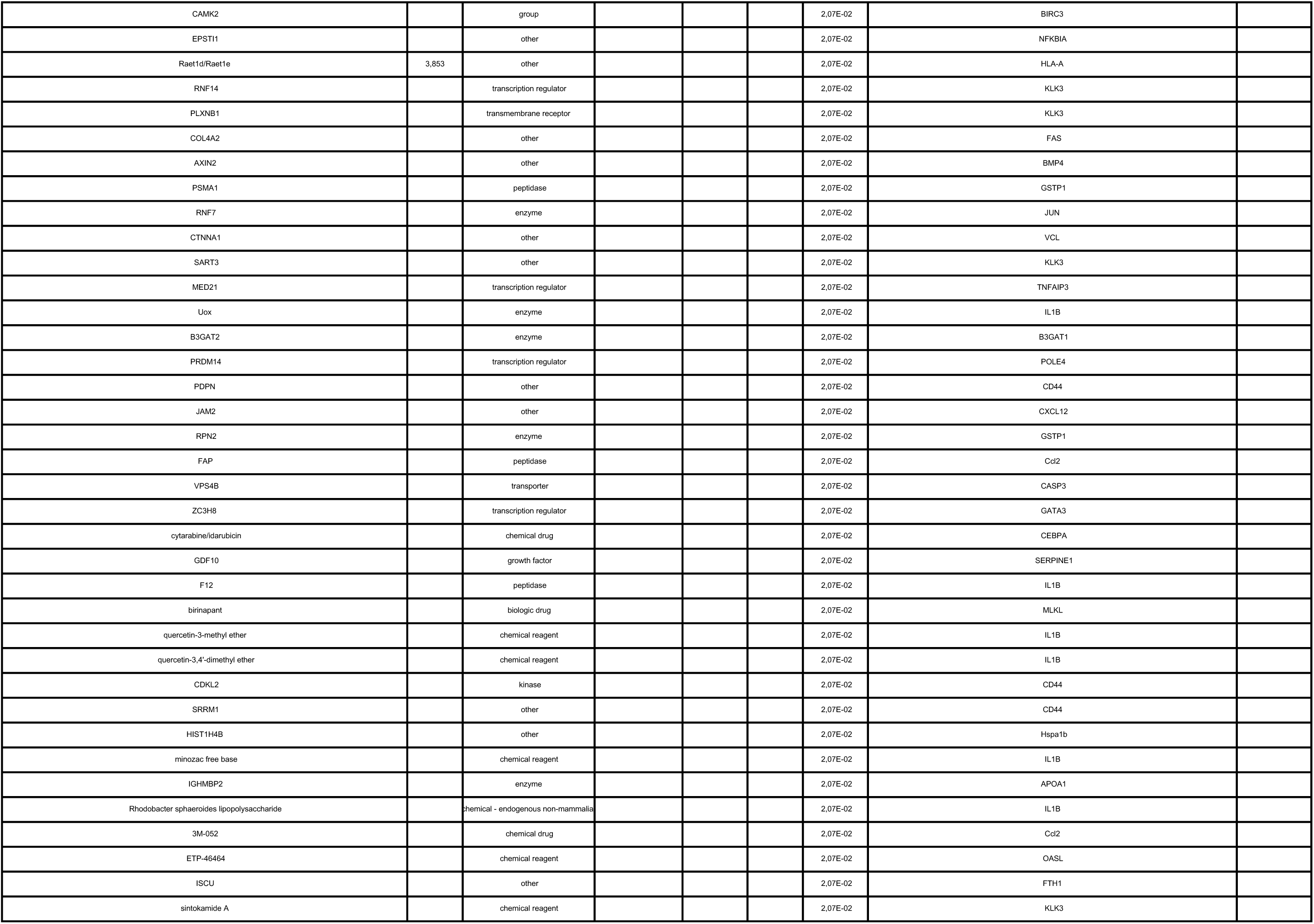
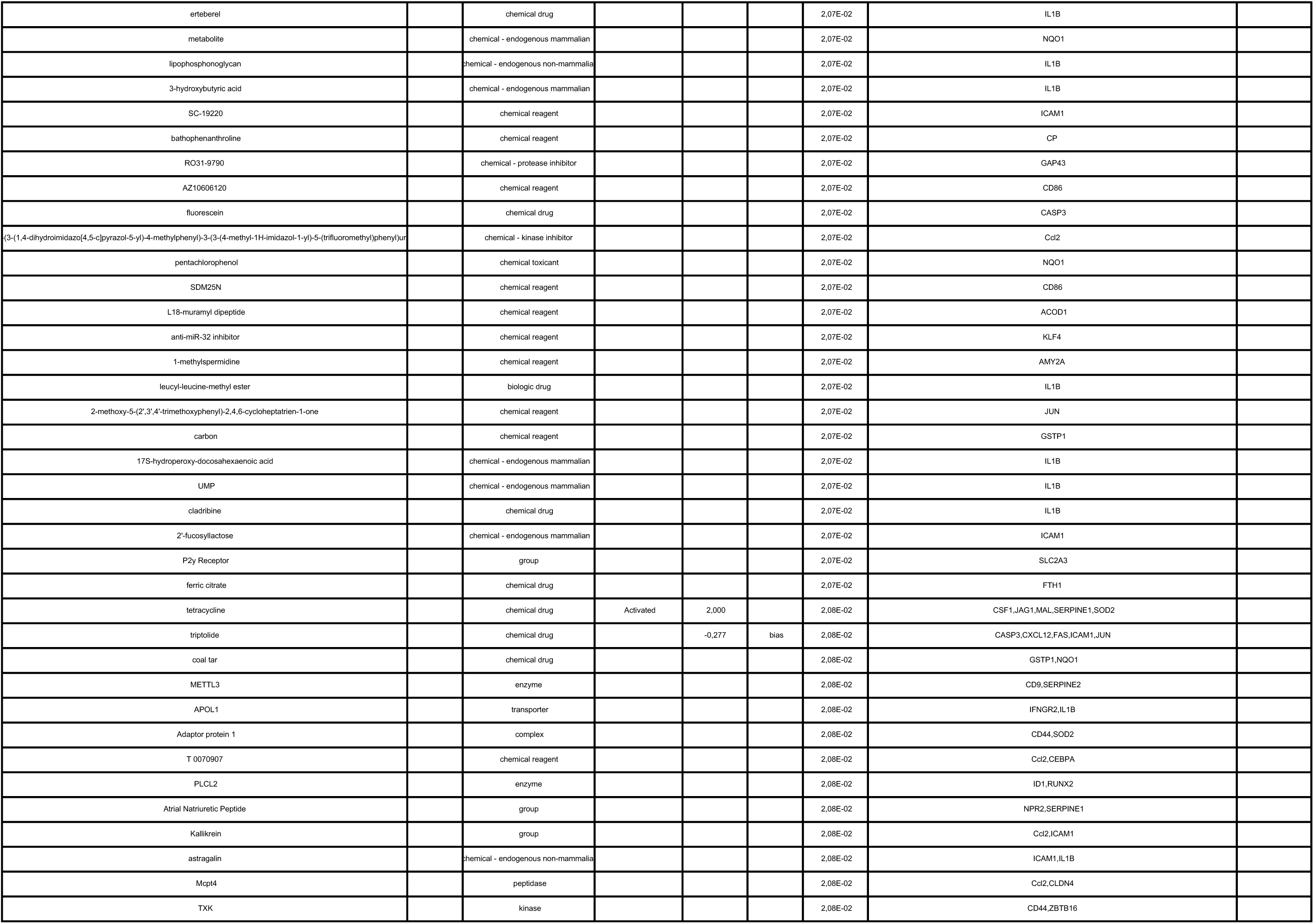
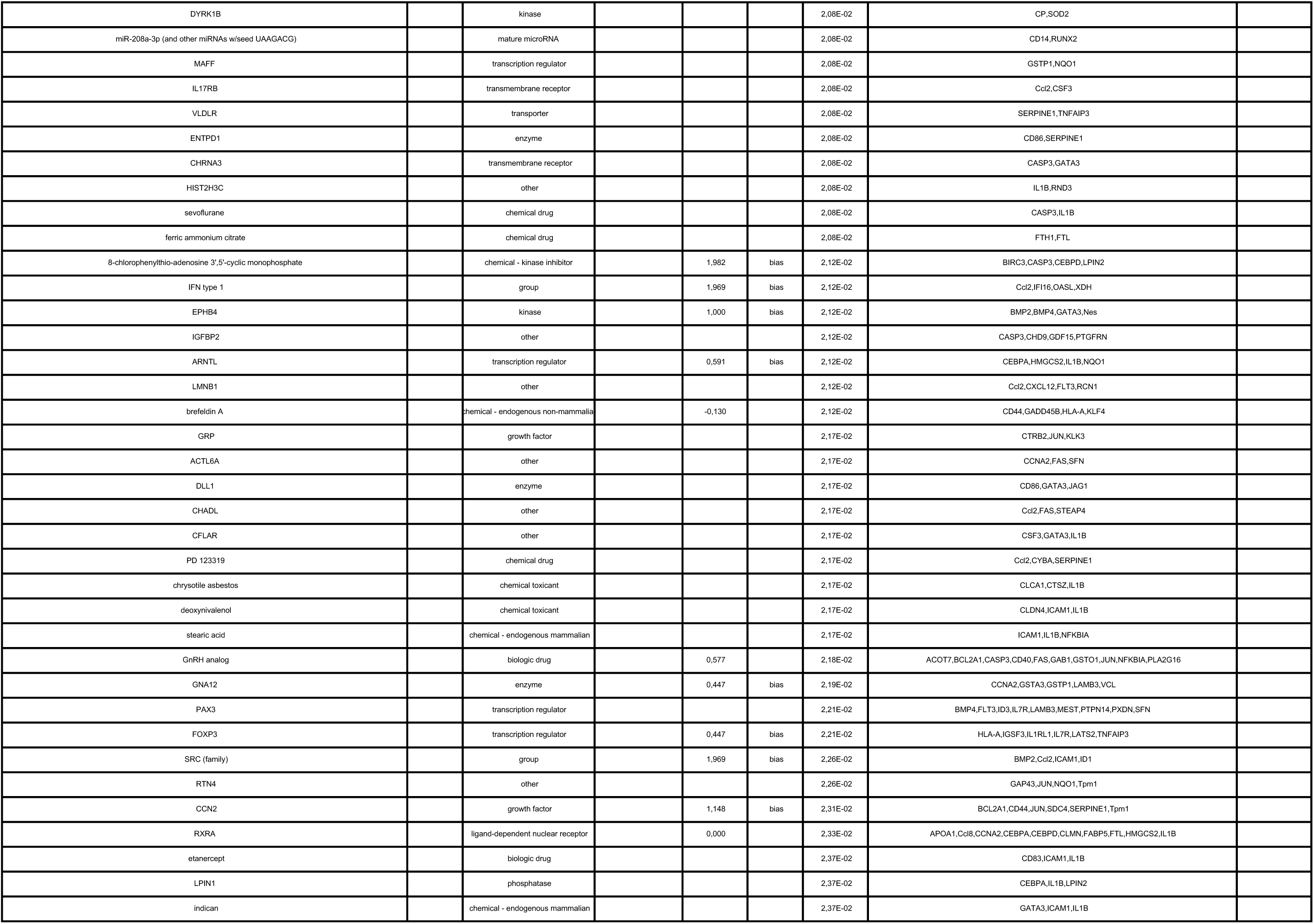
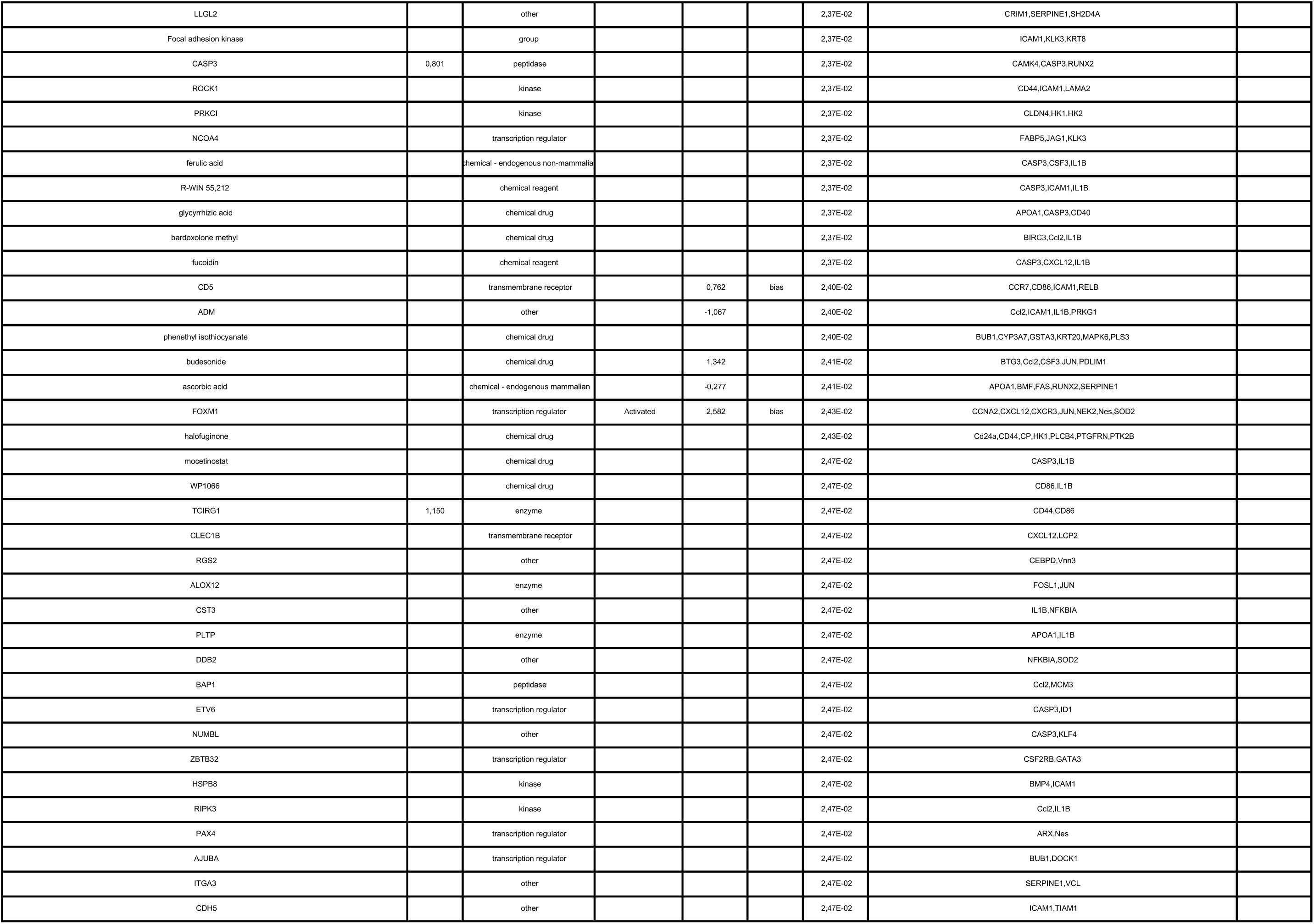
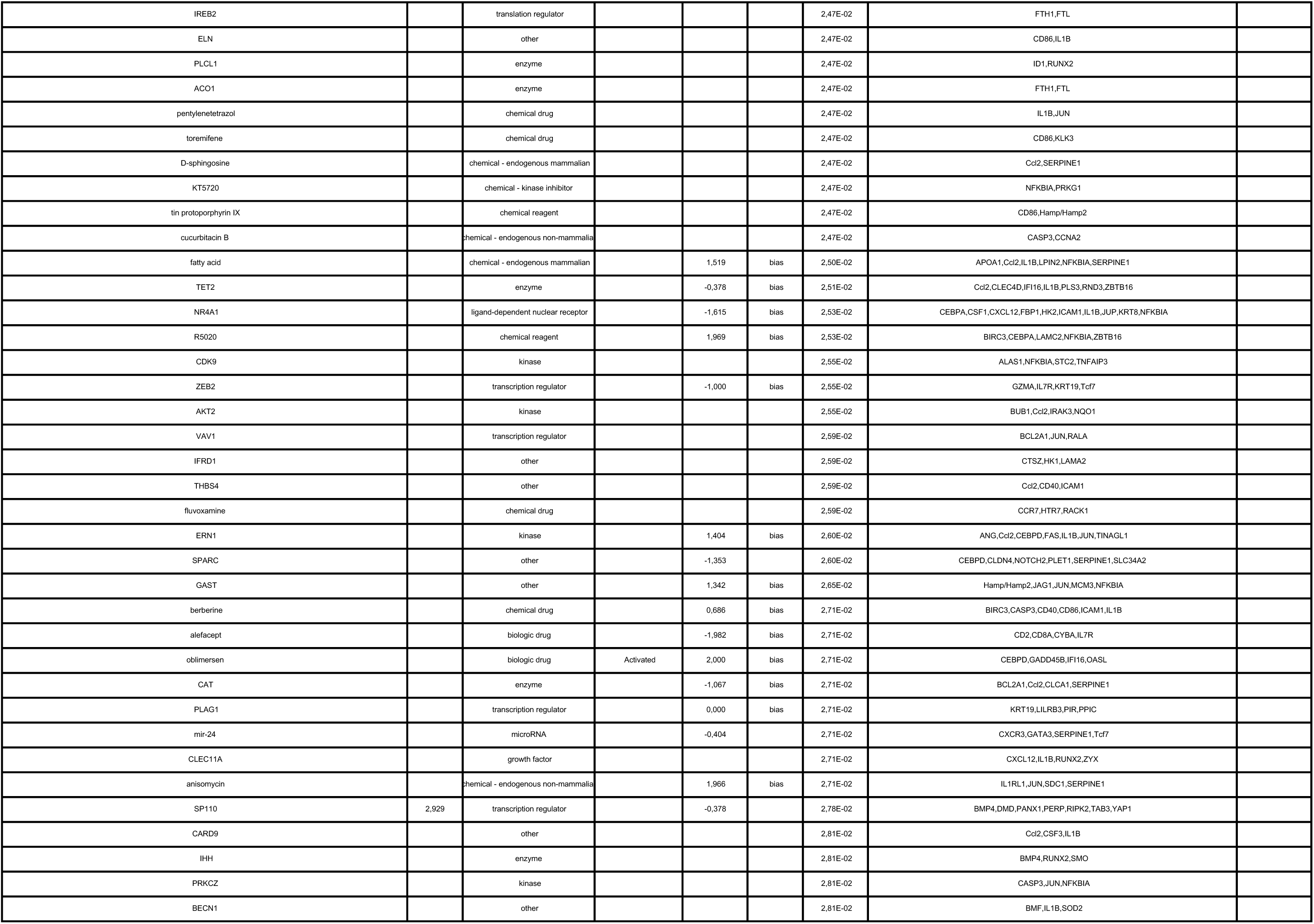
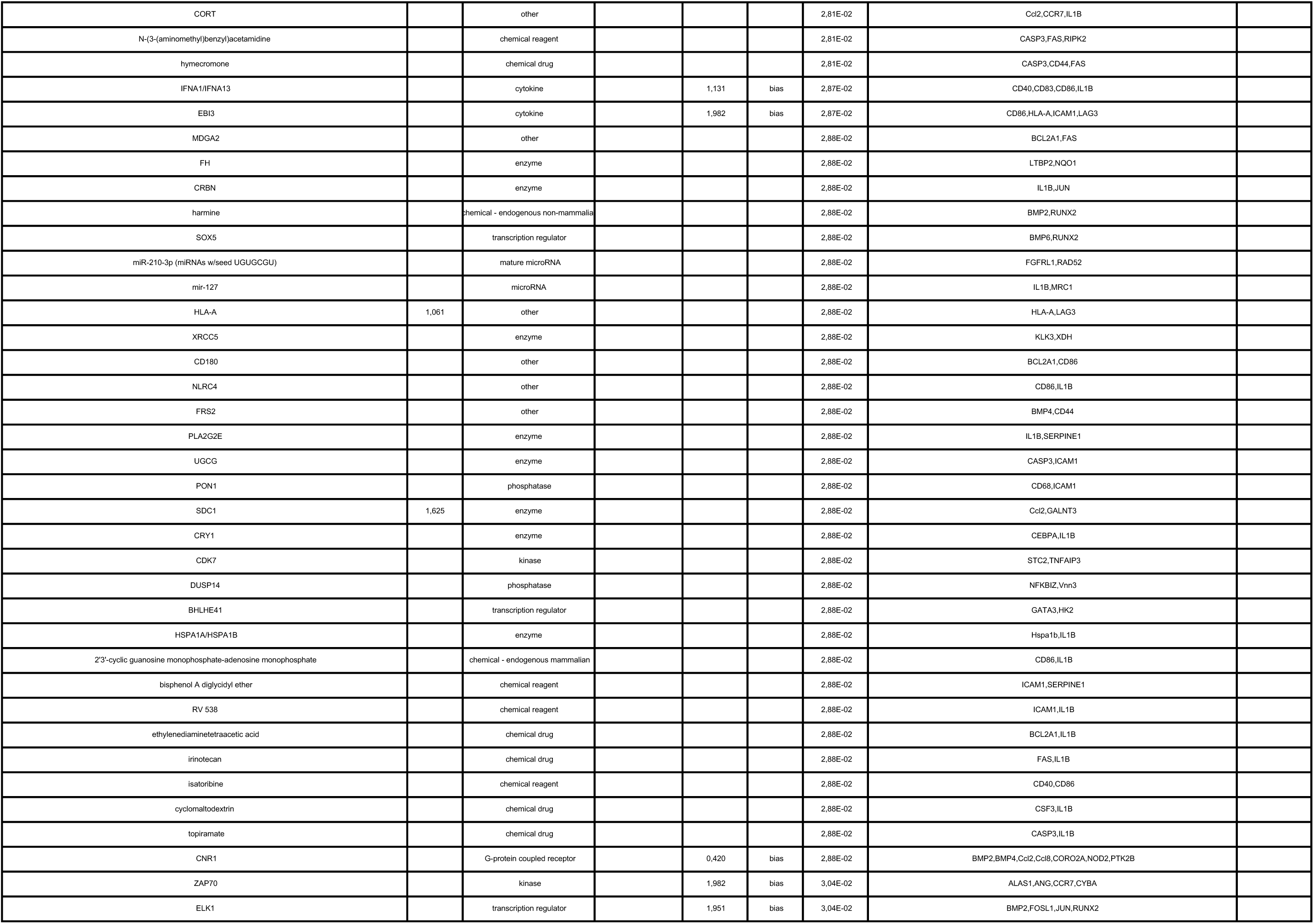
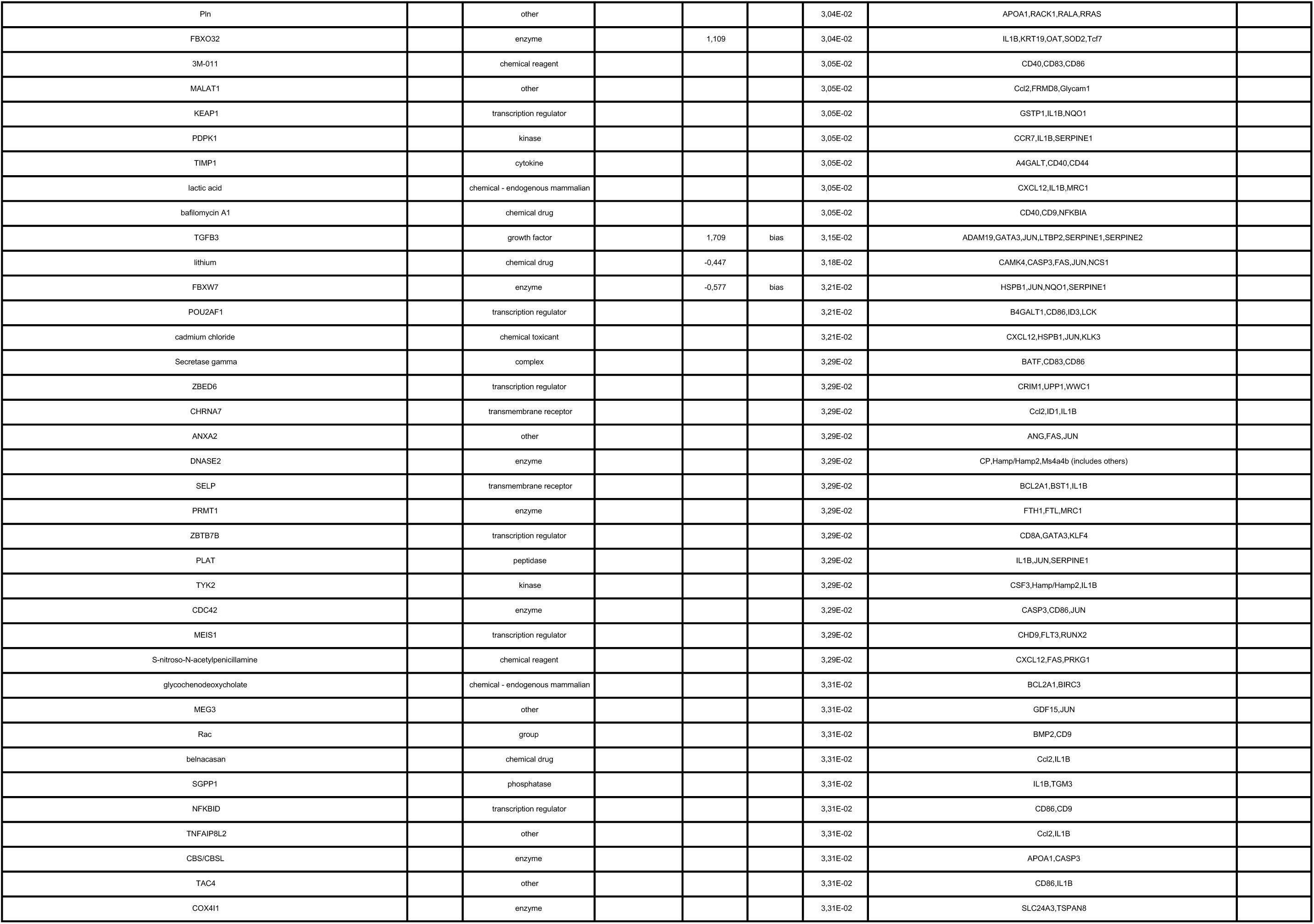
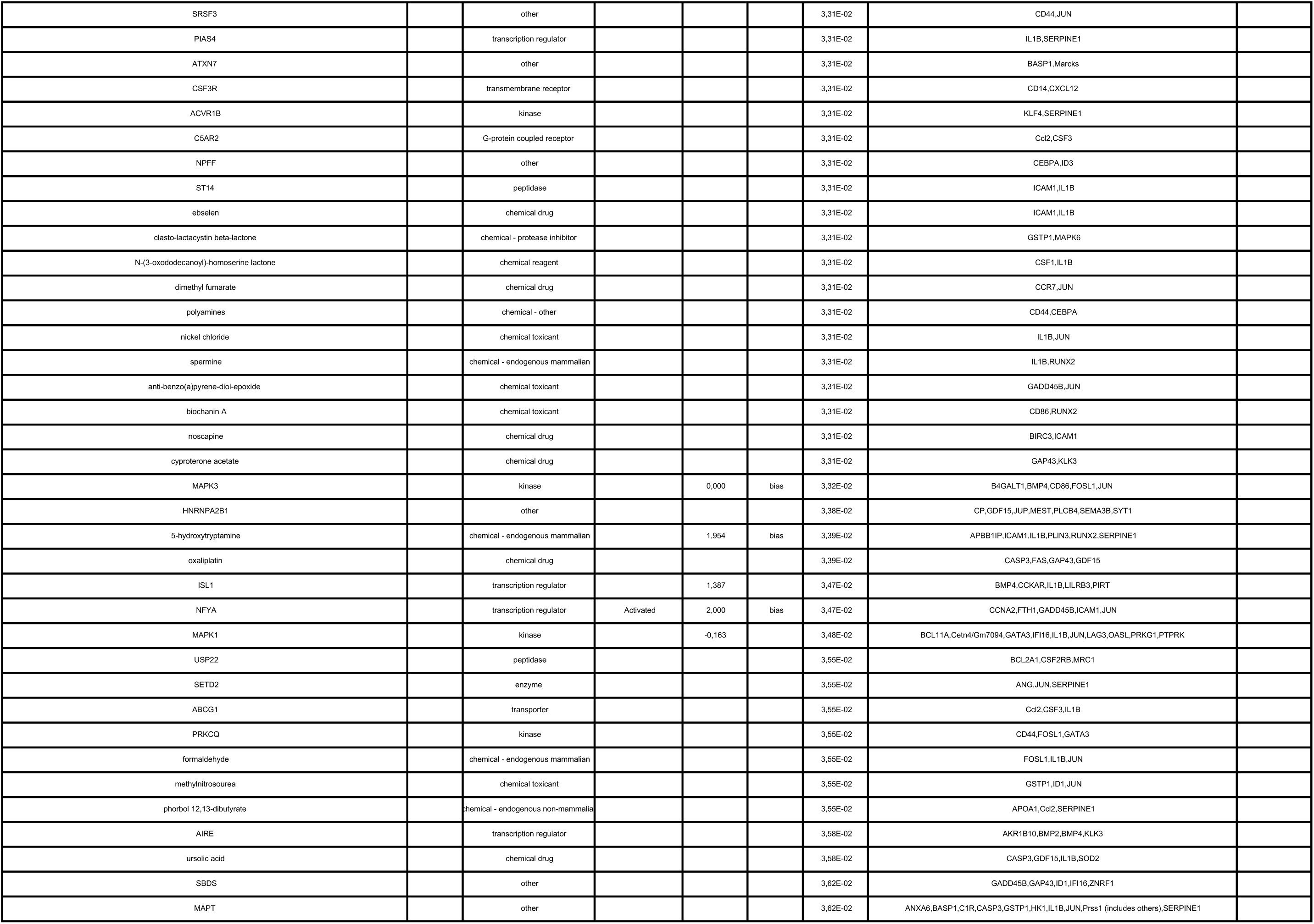
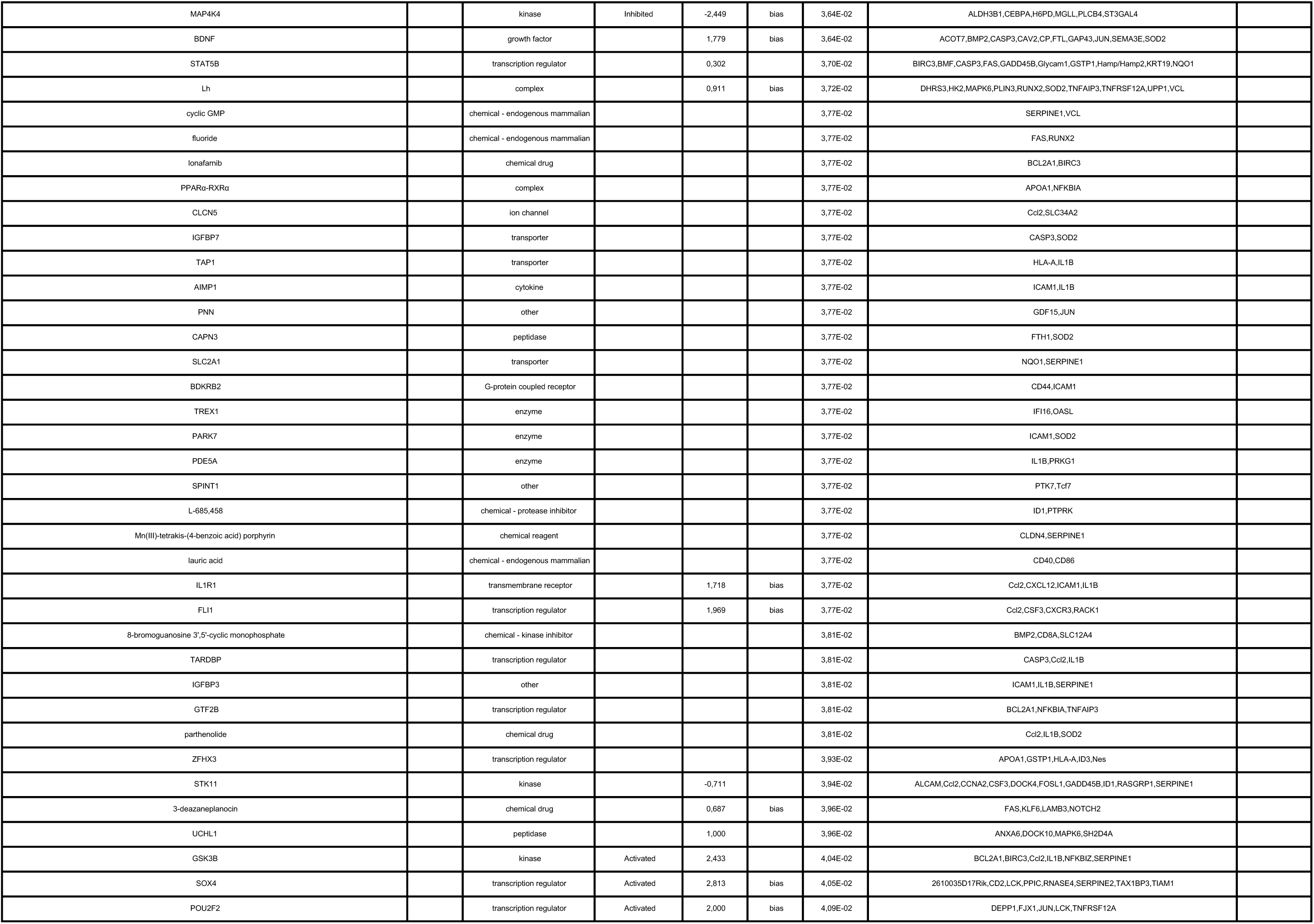
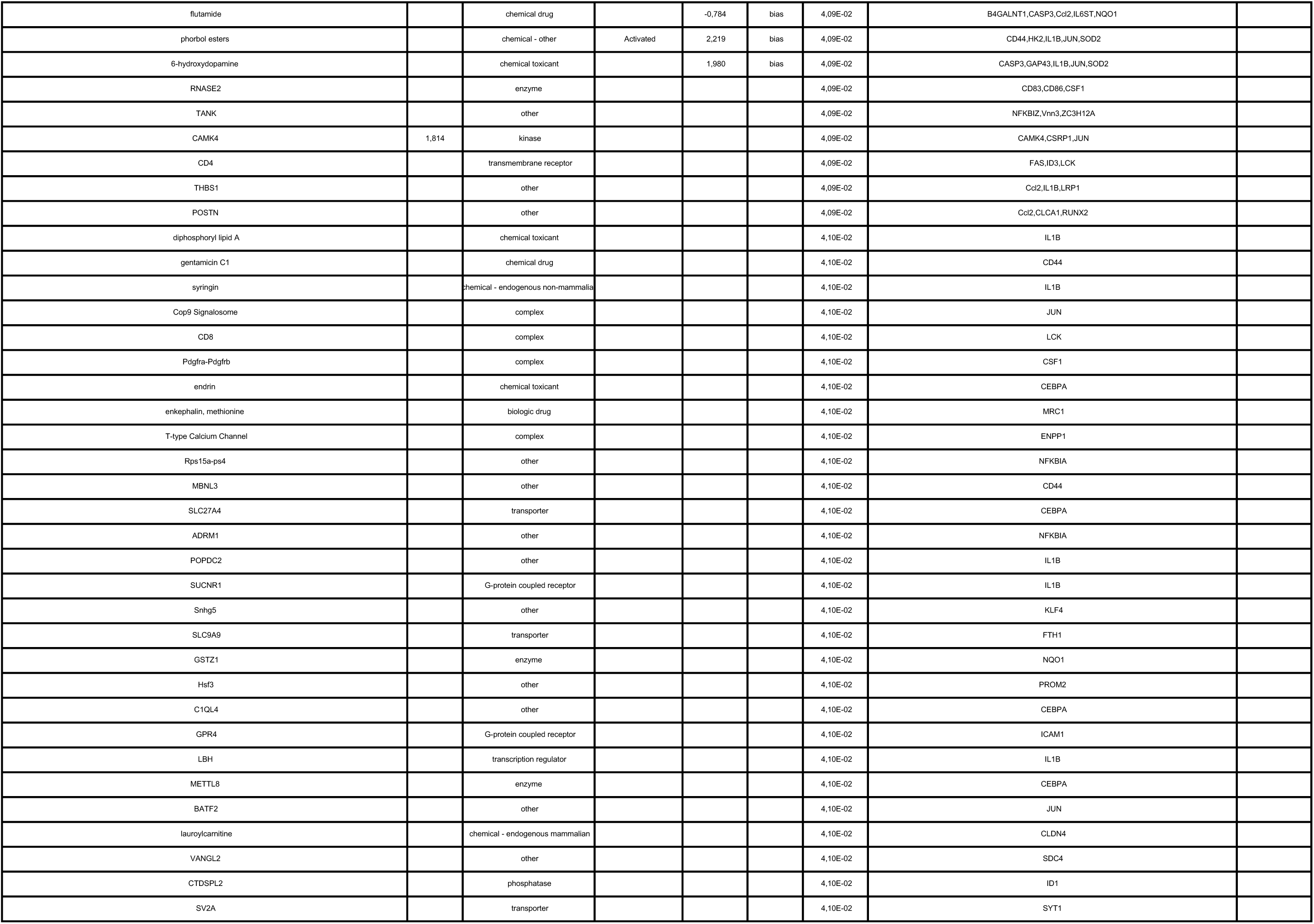
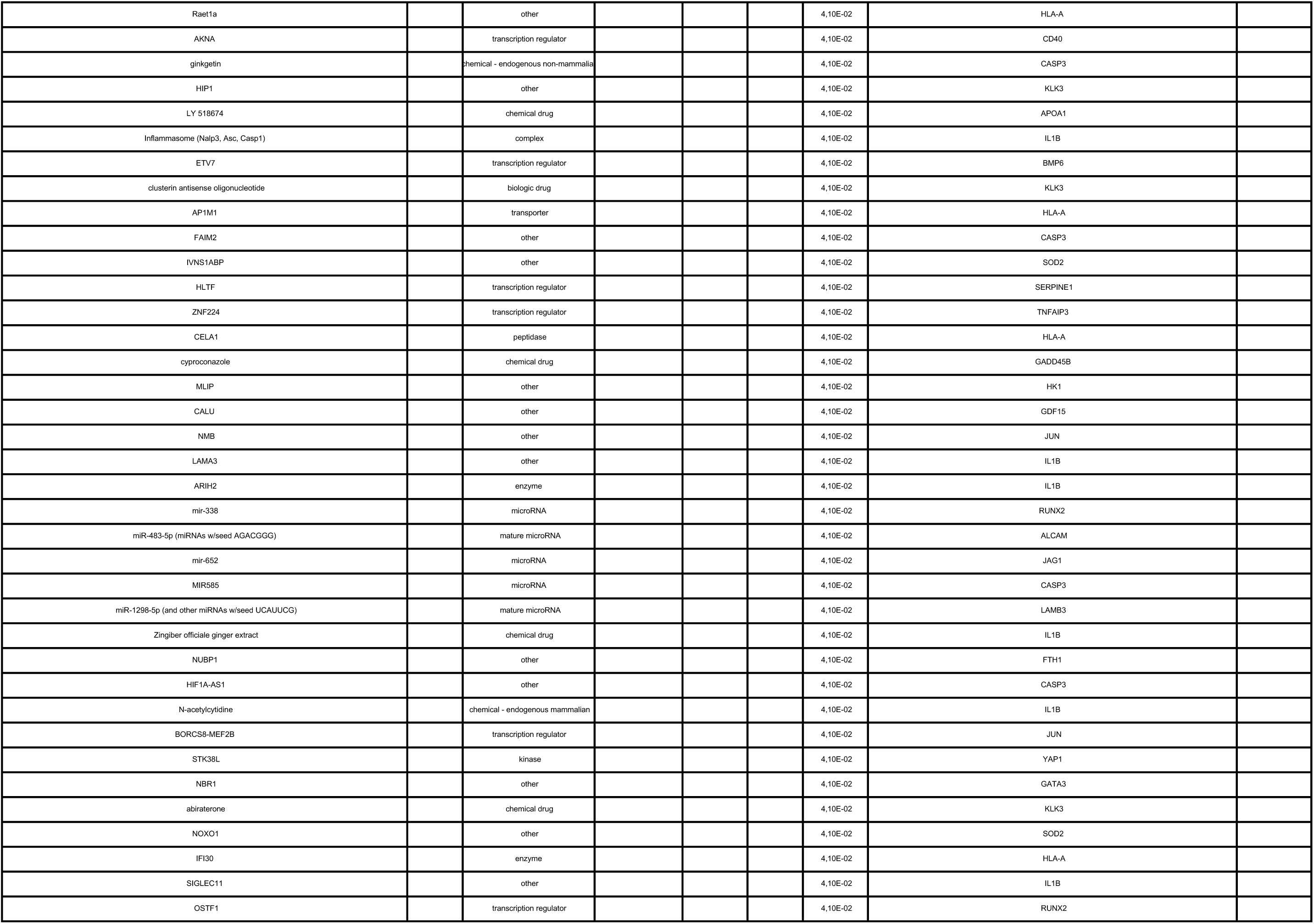
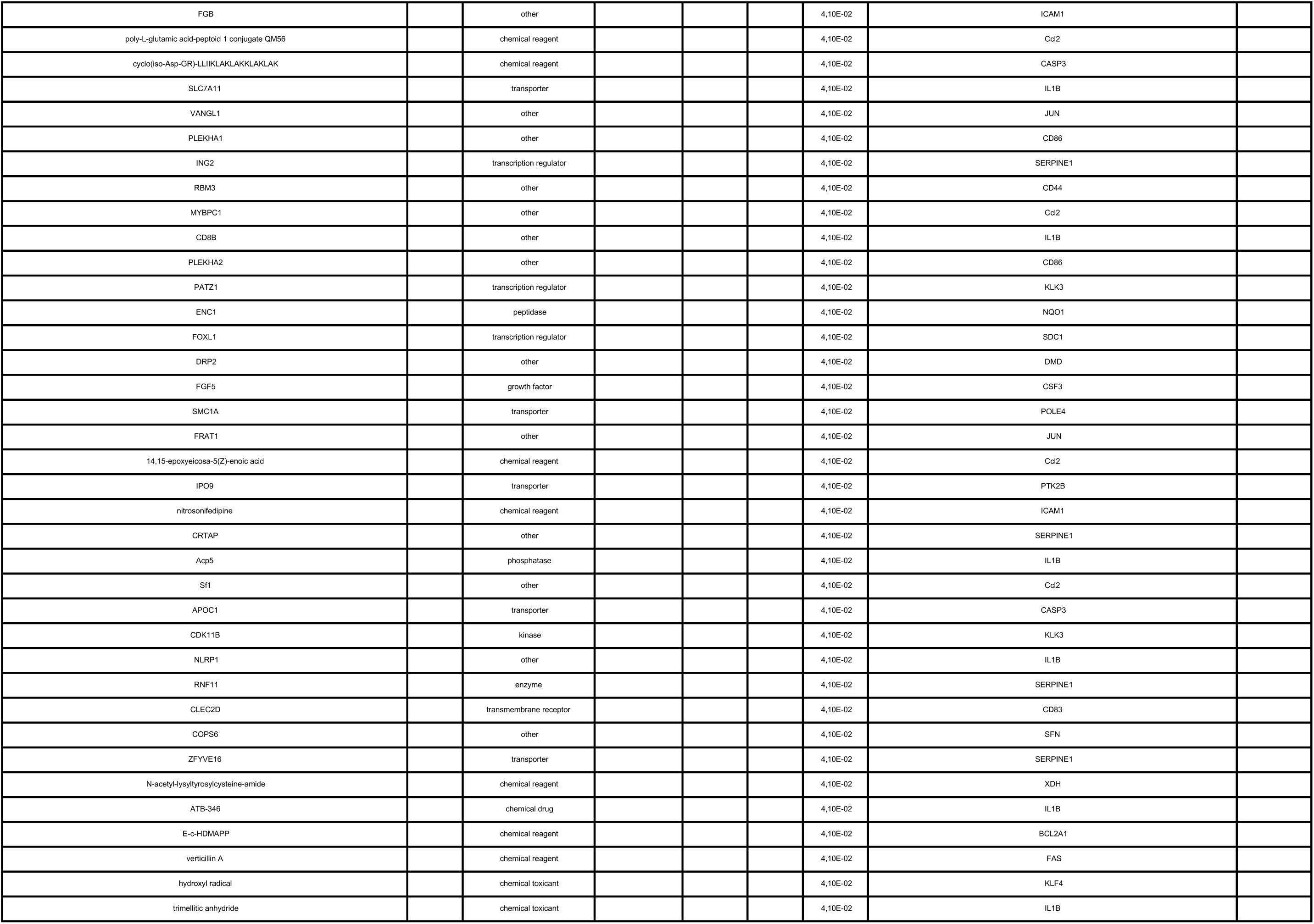
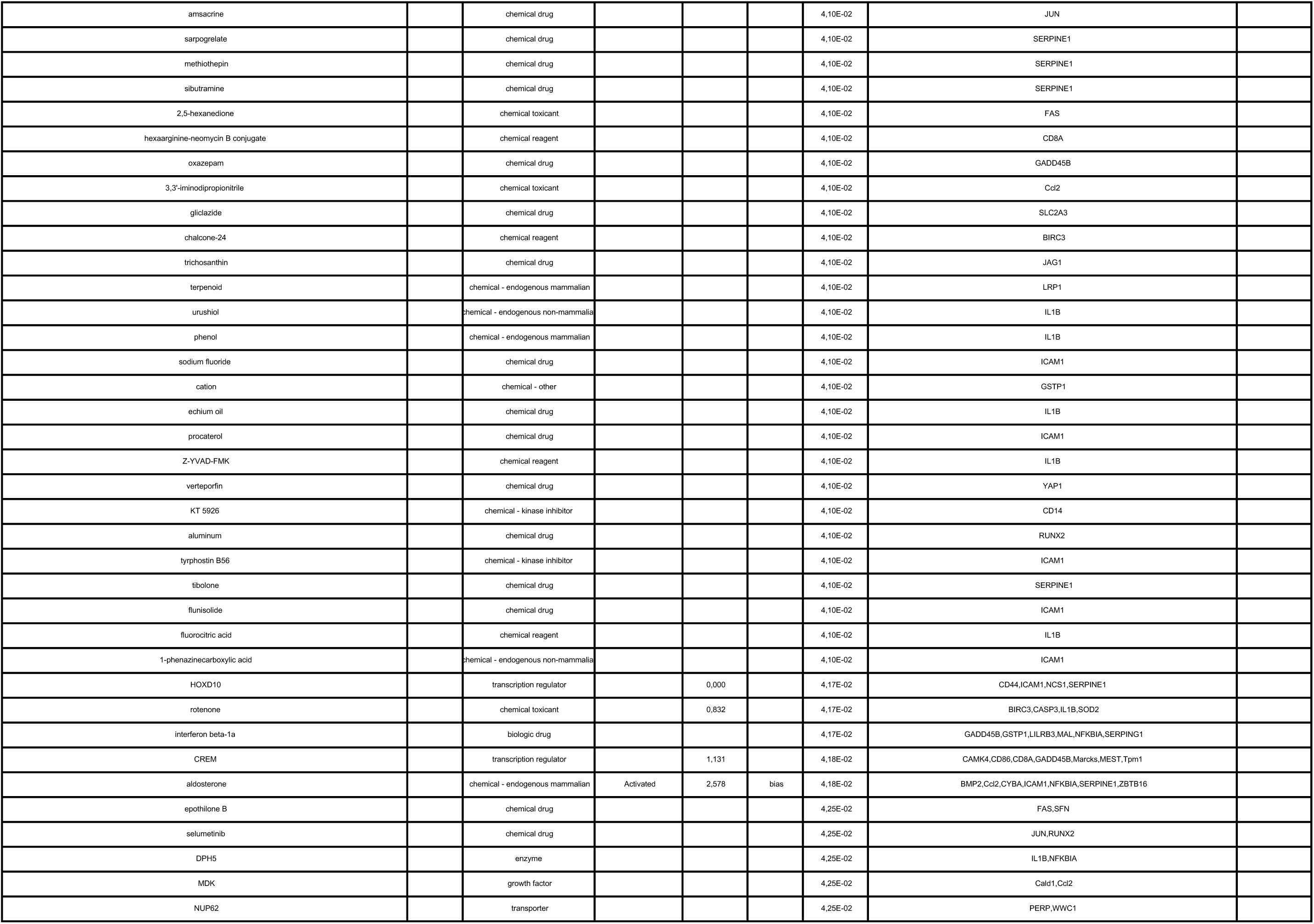
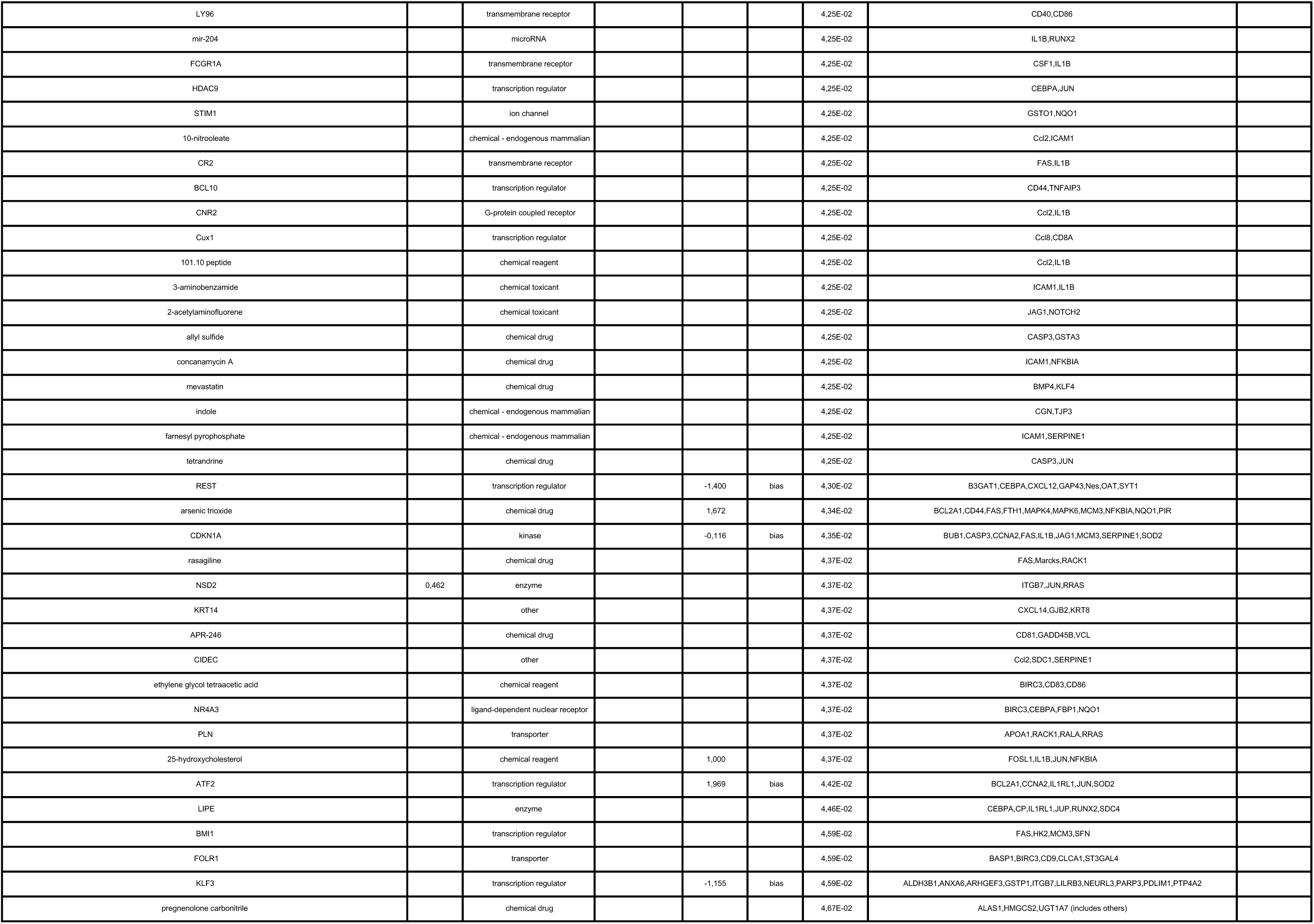
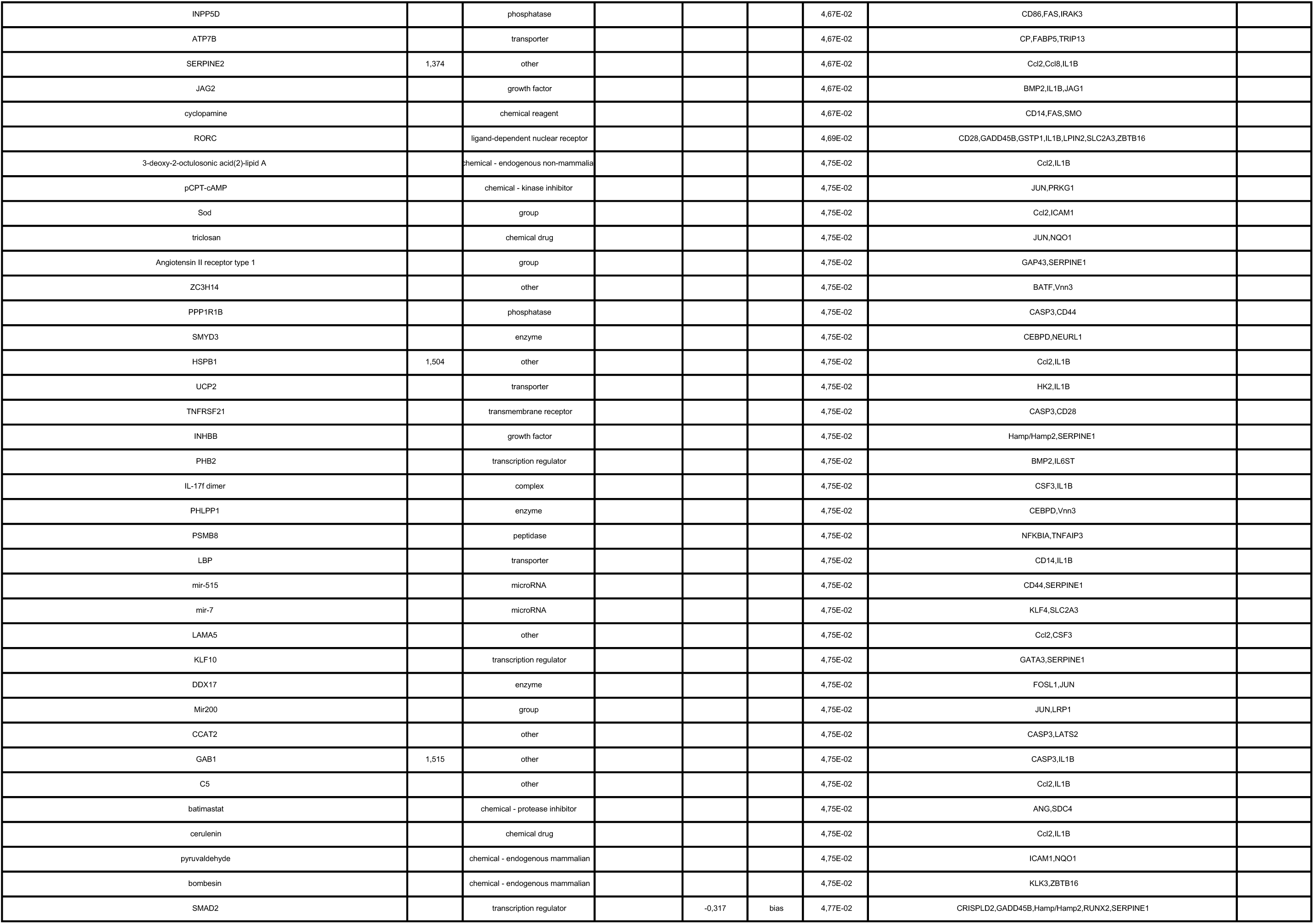
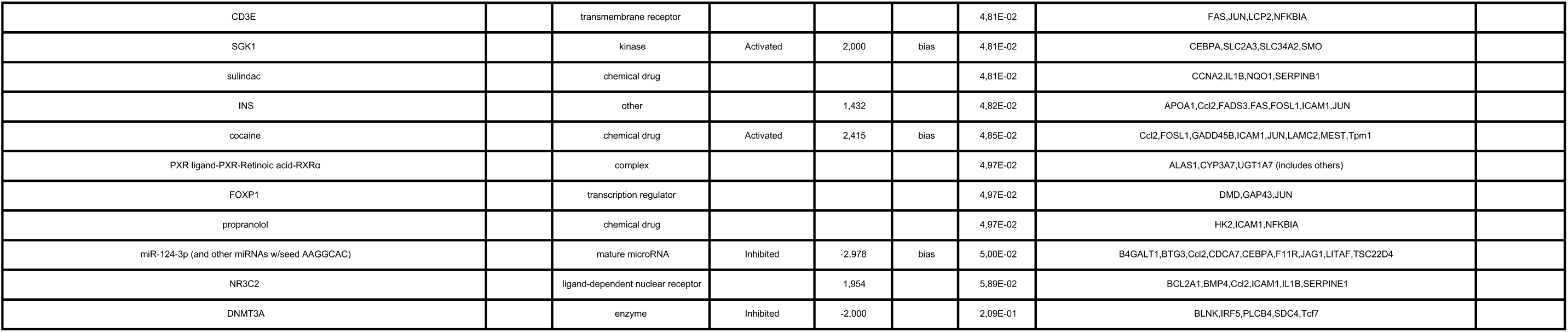
Pathway analysis of RNA-sequencing data in pancreatic islets isolated from control *E2f1*^fl/fl^ and β-cell specific *E2f1*^β-/-^ deficient mice identifies potential upstream regulators associated to *E2f1* deficiency.

### Interactomic analysis by rapid immunoprecipation mass spectometry of endogenous protein (RIME) identifies HDAC1 and HDAC6 as partners of E2f1 in β cell

We next asked whether E2f1 could directly interact with HDAC enzymes in β cell. Rather than addressing this using coimmunoprecipitation strategies targeting a series of specific HDAC, we decided to follow a global interactomic approach by using the rapid immunoprecipitation mass spectrometry of endogenous protein (RIME) technology (Mohammed *et al*, 2013) enabling the identification of E2f1-associated proteins at proteome-wide level in Min6 cells. Consequently, E2f1 was over-expressed in Min6 cells through transfection of a plasmid encoding E2f1 cDNA fused to Flag tag followed by an anti-Flag-based E2f1 immunoprecipitation in order to identify E2f1 interacting proteins through mass spectrometry analysis. RIME analysis revealed that 245 peptides were found to be associated with E2f1 in Min6 cells (Table 2 and Supplementary figure 5A). As expected, E2f1 peptides were the most significantly enriched peptides thus validating our approach. Interestingly, peptides from HDAC6 and HDAC1 were also significantly enriched providing evidence that these two HDAC enzymes were associated with E2f1 in Min6 cells at the chromatin level (Table 2). Altogether, these results suggest that E2f1-mediated transcriptional repression may occur through interactions with HDAC1 and/or HDAC6.

**Table 2.**
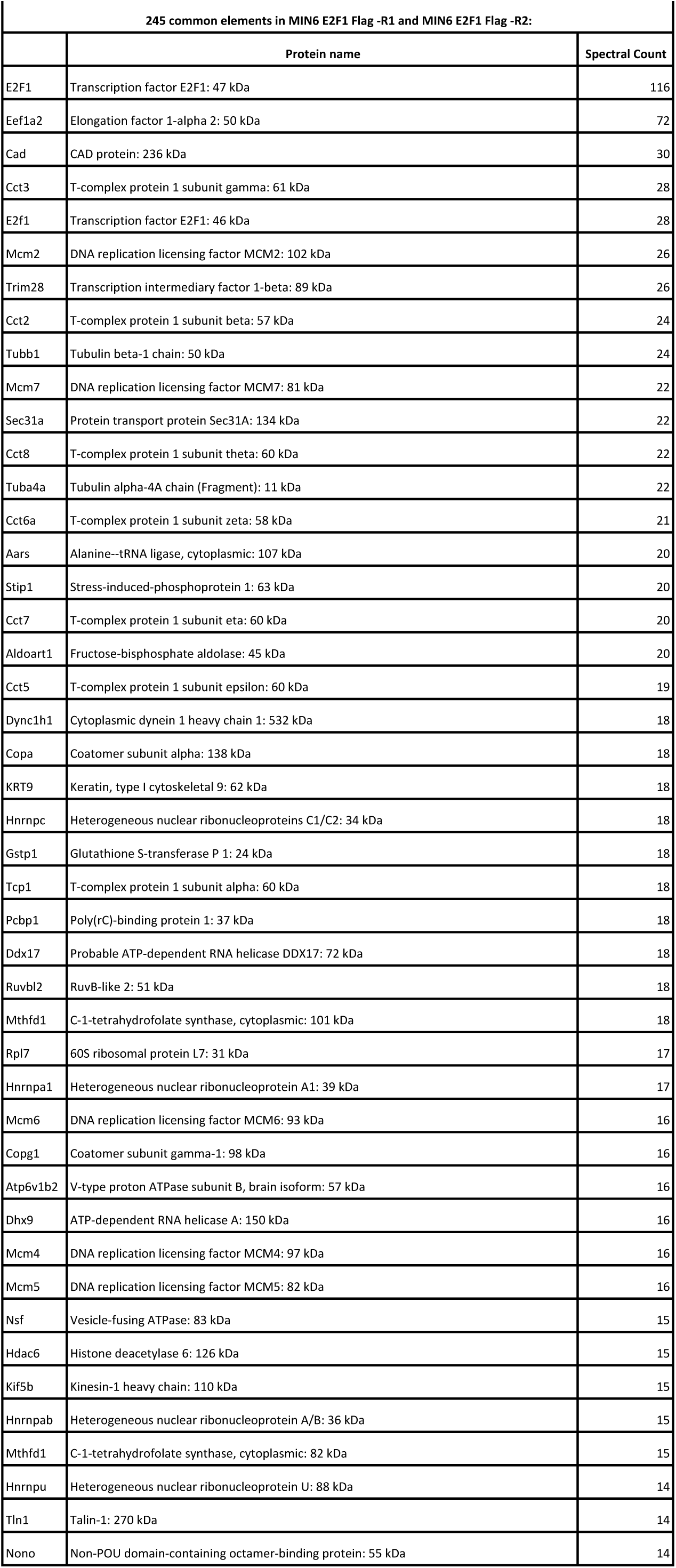
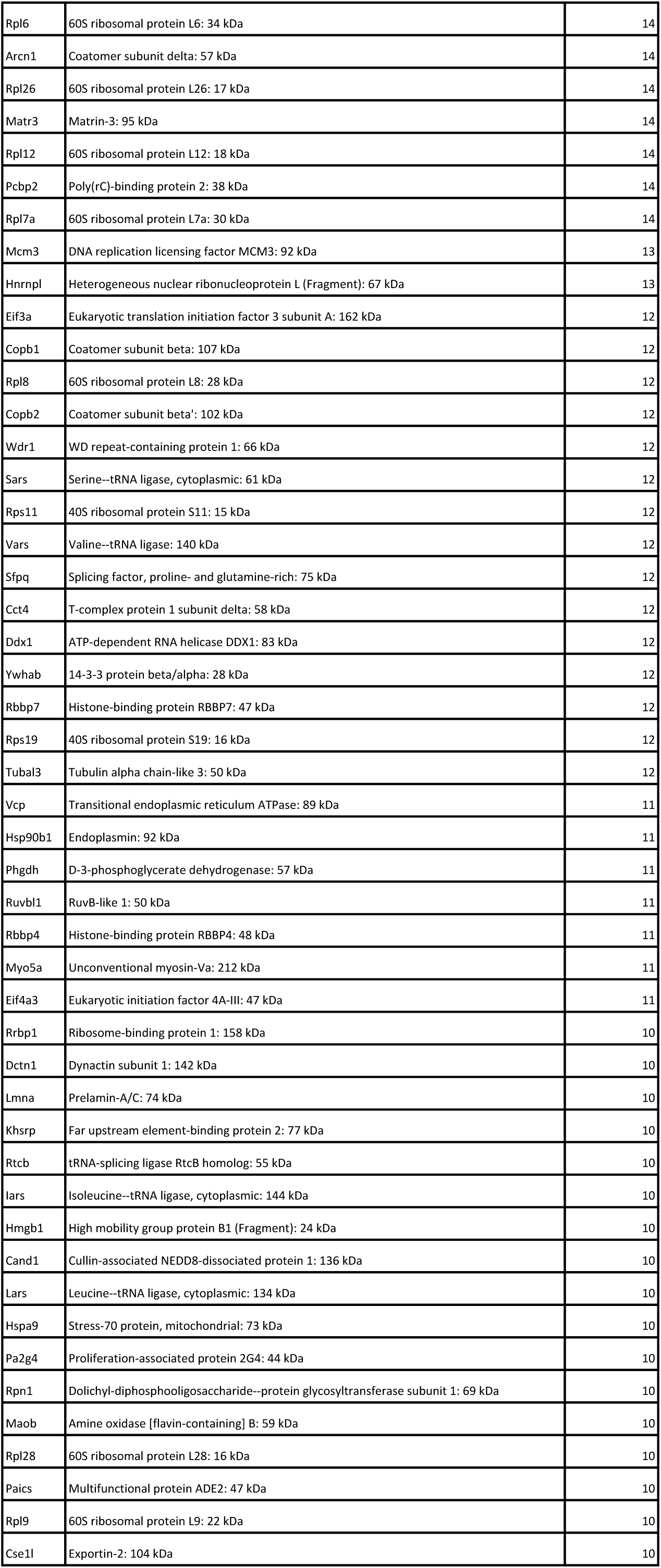
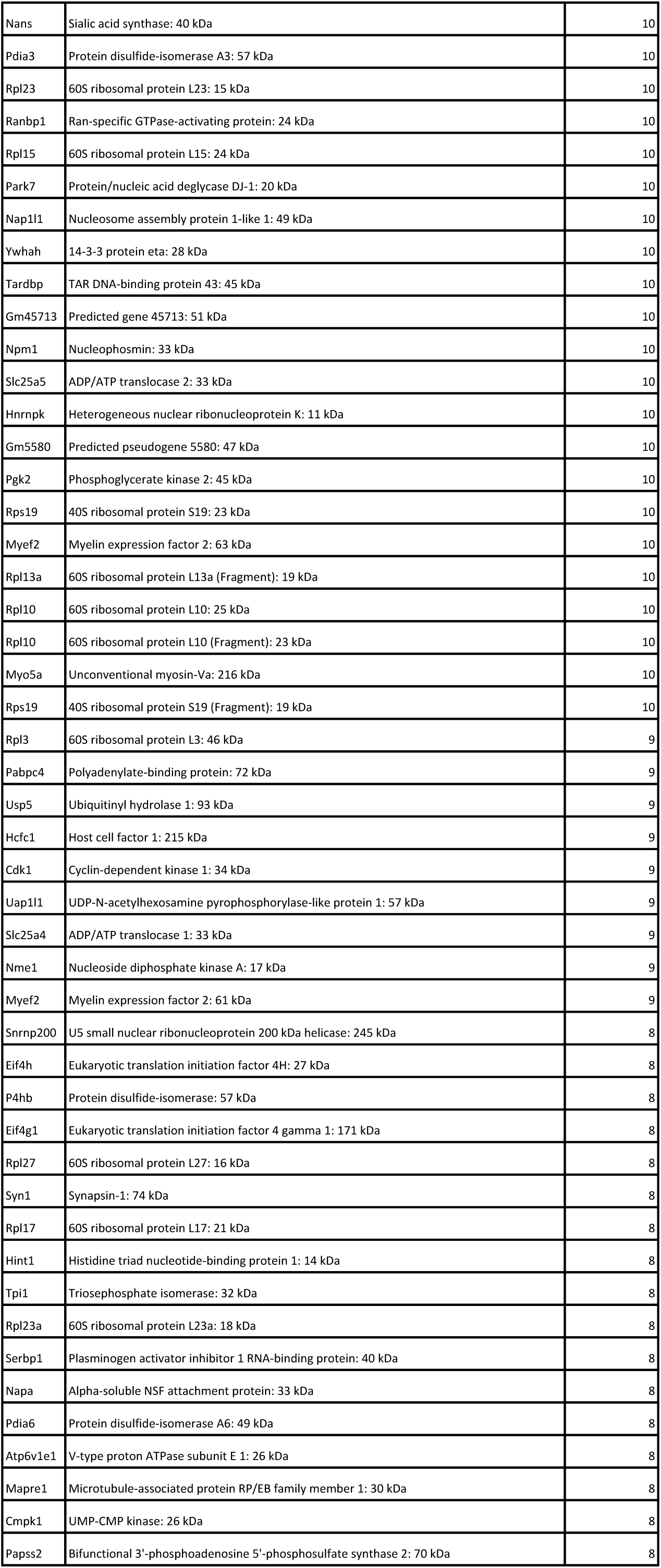
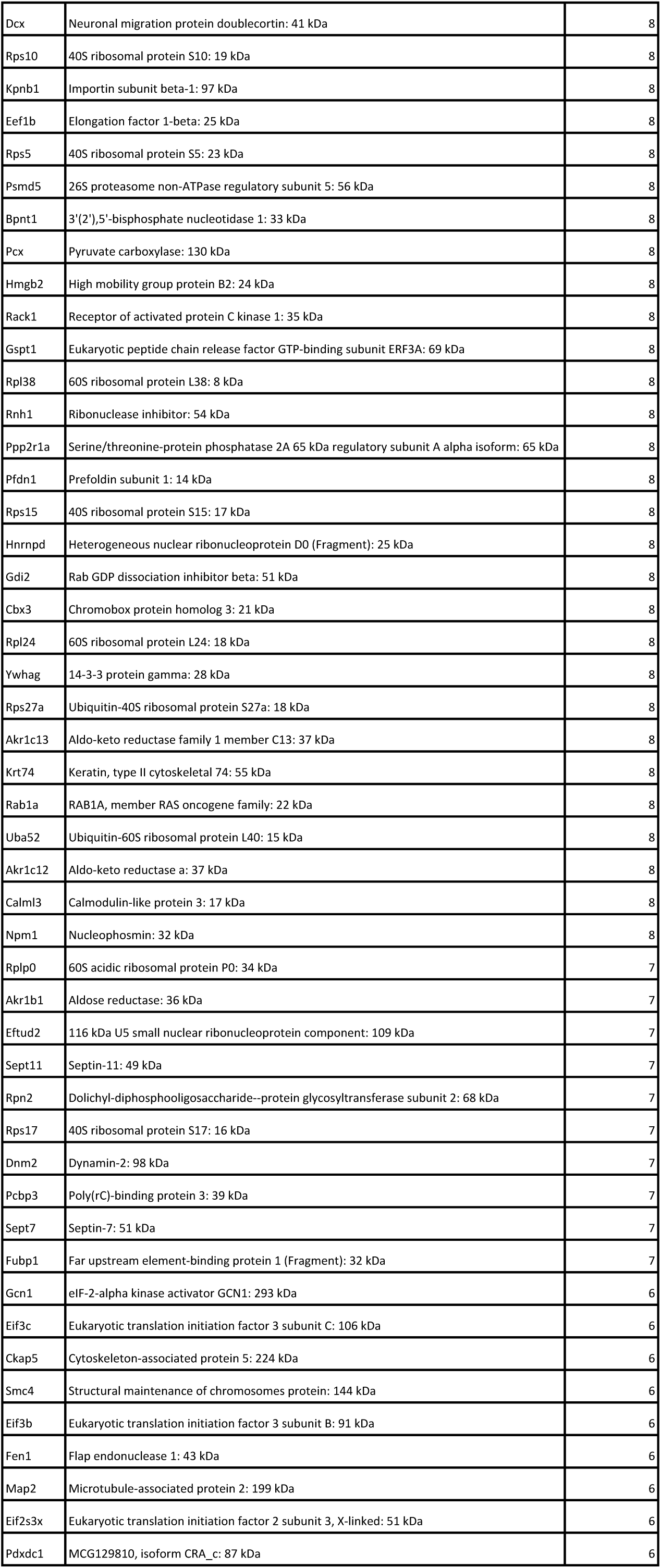
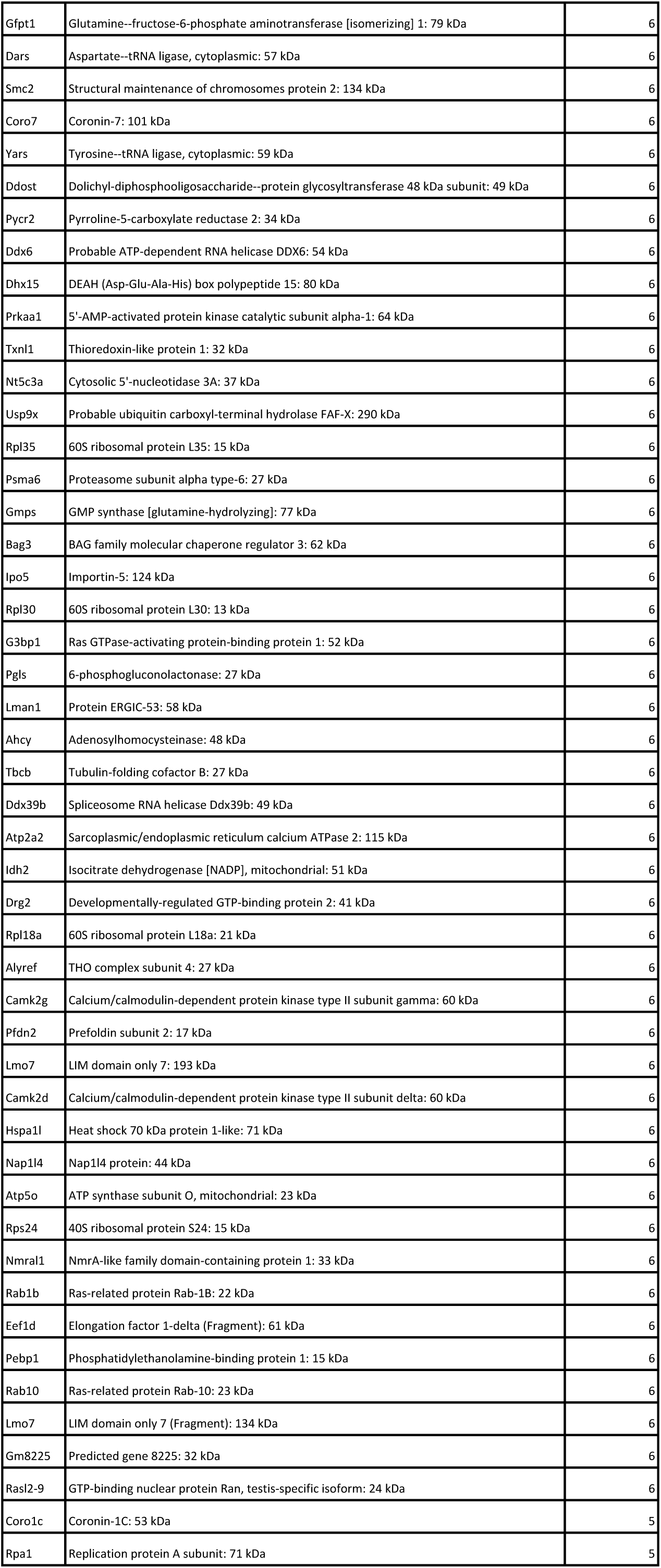
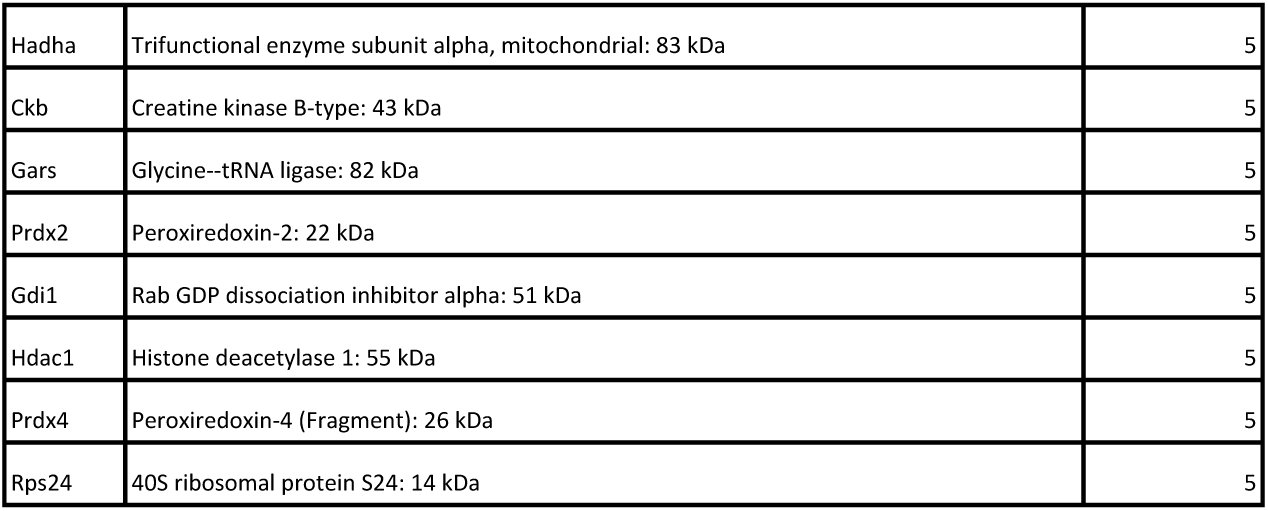
List of peptides associated to E2F1 in Min6 cells identified through RIME experiments. RIME was performed on biological duplicates and common peptides found in both replicates are shown.

## Discussion

In the present study, we show that the transcription factor E2F1 plays an essential role in maintaining β-cell function and identity through the control of transcriptomic and epigenetic programs within the pancreatic islets. Germ-line as well as β-cell specific *E2f1* deficient mice exhibit impaired glucose homeostasis and decreased β-cell functions that are associated to a gene expression reprogramming. A detailed analysis of the transcriptome and epigenome of the pancreatic islet reveals that E2f1 is necessary to maintain several key genes for cell adhesion and inflammatory processes in a repressive state through the maintenance of epigenomic marks including H3K27me3 (Figure 5). We showed that the β-cell specific loss of *E2f1* function also leads to an altered α-to-β ell ratio, while maintaining islet size, associated to insulin secretion defects in response to glucose and affects β-cell identity maintenance. In addition, through the use of the E2Fs inhibitor HLM006474, which selectively disrupts the binding of E2Fs to its target genes (Figure 4 and (Ma *et al*., 2008; Rosales-Hurtado *et al*., 2019)), we demonstrated here that a pharmacological intervention lowering the E2F1 transcriptional activity could mimic the genetic effects of *E2f1* deficiency on the maintenance of β-cell identity and function, in mouse β-cell lines and human islets. Indeed, the treatment of human islets with this compound not only blunts β-cell function by impairing GSIS, but also leads to an alternative transcriptional program that affects the expression of α- and β-cell markers. Our results, associated to those showing that E2F1 overexpression stimulates β-cell proliferation and function (Grouwels *et al*, 2010), suggest that carefully targeting this pathway might be of interest for the maintenance of β-cell functions in the context of diabetes.

Our study is the first one to demonstrate a direct, cell-autonomous contribution of E2f1 in controlling not only GSIS but also β-cell identity maintenance, without affecting the pancreatic islet cell number. The role of E2F1 in the control of cell cycle and proliferation has been extensively studied, particularly in the context of cancer (Poppy Roworth *et al*., 2015). However, its function in non-proliferating fully differentiated cells, including β cells, remains to be precisely deciphered. The observation that cell cycle regulators, including E2F1, are expressed in cells that are not proliferating, suggest that they may be involved in adaptive pathways that are independent of cellular proliferation. We and others have demonstrated that this pathway plays a key role in post-natal β-cell proliferation (Fajas *et al*, 2004; Iglesias *et al*, 2004; Li *et al*, 2003), glucose homeostasis and insulin secretion (Annicotte *et al*., 2009; Boni-Schnetzler *et al*, 2018; Grouwels *et al*., 2010). Studies also revealed that the Cdk4-E2F1-pRb pathway controls the fate of pancreatic progenitors through the transcriptional control of the expression of *Ngn3* and *Pdx1* as well as Pdx1 protein stability (Cai *et al*, 2013; Kim & Rane, 2011; Kim *et al*, 2011). Most of these studies, including those from our group, were performed using germ-line *E2f1* deficient mice, which precludes to ascertain a cell-autonomous role of E2f1 in the endocrine pancreas and β-cell functions. To better appreciate the specific role of E2F1 in β cells, we specifically knocked-down *E2f1* in the β cell using the Cre/loxP technology. Although we cannot rule out an early role for E2f1 in pancreatic progenitors due to the use of the RIP-Cre mice (Herrera, 2000), our data demonstrate that E2f1 expression within the β cell is necessary to maintain a proper gene expression to regulate insulin secretion and glucose homeostasis.

The complexity of E2F1 biology resides in the fact that it can positively or negatively regulate the expression of its target genes. Our data suggest that E2F1 is part of a repressor complex that regulates β cell identity. In line with this, we observe that most of the genes with altered expression in *E2f1*^β-/-^ isolated islets were upregulated, suggesting that the repressive effects of E2F1 are key in maintaining β-cell identity and subsequent β-cell function. Accordingly, repression of non β-cell programs is crucial for maintaining β-cell identity. Pdx1 (Gao *et al*., 2014), Pax6 (Swisa *et al*., 2017), Nkx6.1 (Schaffer *et al*., 2013) and Nkx2.2 (Gutierrez *et al*., 2017) are key transcription factor required to maintain gene repression of non-β-cell programs. Interestingly, *Nkx6.1* mRNA levels are decreased in *E2f1*^β-/-^ isolated islets, suggesting that E2F1 could modulate β cell functions through *Nkx6.1* regulation. Chromatin regulators and epigenomic features play important roles in the control of β-cell identity and plasticity (Arda *et al*, 2016; Avrahami *et al*, 2015; Campbell & Hoffman, 2016). Indeed, the modulation of islet enriched transcription factor activity, including Pdx1 or Nkx2.2, involves their interaction with coregulators such as Dnmt1, Dmnt3a or Hdac1 (Dhawan *et al*, 2011; Papizan *et al*, 2011; Spaeth *et al*, 2016) or their accessibility to chromatin, as demonstrated for the β-cell specific deletion of protein arginine methyltransferase 1 (Prmt1) that results in the loss of β-cell identity and diabetes development (Kim *et al*, 2020). In addition, recent studies also indicate that loss of Polycomb silencing in human and mouse β-cells contributes to loss of β-cell identity in diabetes (Lu *et al*., 2018). Indeed, the β-cell specific deletion of *Eed*, a component of the Polycomb repressive complex (PRC)-2, triggers β-cell dysfunctions, dedifferentiation and diabetes development associated to chromatin-state-associated transcriptional dysregulation (Lu *et al*., 2018). The findings that loss of PRC2 activity induces β-cell plasticity through epigenomic reprogramming at both active and silent genes suggest that maintaining proper and specific histone marks and chromatin state at precise loci is crucial to maintain normal β-cell functions and avoid T2D development. Interestingly, most of the upregulated genes in *E2f1*^β-/-^ pancreatic islets were characterized by bivalent H3K4me3/H3K27me3 and Polycomb-repressed (H3K27me3) marks. In addition, epigenomic interventions triggering ectopic acetylation and gene derepression contribute to β-cell dysfunctions. Indeed, blocking HDAC activity through the use of the HDAC inhibitor SAHA, impairs glucose intolerance in mice fed a HFD (Lu *et al*., 2018). Here we show that the loss of *E2f1* function triggers transcriptional dysregulation of specific genes that are in a bivalent (*i.e.* H3K4me3/H3K27me3 and Polycomb-repressed (H3K27me3) marks) and active (*i.e.* RNA-Pol2 recruitment, H3K4me3 and H3K27ac histone marks) state in healthy pancreatic islets, suggesting an ectopic acetylation in the promoter region of upregulated genes in *E2f1*^β-/-^ islets and decreased activation of the promoter region of downregulated genes. Accordingly, the use of the HDACi TSA also impairs the expression of genes regulated by *E2f1* in pancreatic islets. Although E2F1 is a ubiquitous transcription factor that is weakly expressed within β cell compared to *bona fide* β-cell genes, we speculate that this transcriptional regulator may cooperate with β-cell transcription factors and/or the chromatin machinery such as HDAC or the PRC2 complex to integrate some signals necessary for β-cell maintenance in physiological conditions. Since E2F1 regulates gene expression through its interaction with repressor complexes including pRb, SWI/SNF and HDACs (Brehm *et al*, 1998; Luo *et al*, 1998; Magnaghi-Jaulin *et al*, 1998), our finding that E2F1 mediates repression of non β-cell programs deserves deeper investigations to identify the E2F1 complexes that can trigger these transcriptomic and epigenomic effects in the β-cell and their physio(patho)logical consequences.

In summary, the present data highlight that E2F1 transcriptional activity within pancreatic islets is key for maintaining glucose homeostasis and insulin secretion through the regulation of key β-cell identity genes and the repression of non β-cell programs, both in mouse and human islets. The observation that E2F1 levels are decreased in human T2D islets (Lupi *et al*., 2008) suggests that a reduced E2F1 expression or activity may contribute to β-cell failure in diabetes.

## Author contributions

F.O., C.B., X.G., M.M., C.C., E.C., N.R., L.R. and S.A.H. contributed to the *in vivo* and cellular experiments. E.D., S.A., L.B., M.D. and A.B. performed the RNA-seq and ChIP-seq experiments and analysis. P.D.D., Z.B., P.M., L.F., J.K.C. and F.P. provided reagents and data. P.F. discussed the results from the study. J.-S.A. designed the study, supervised the project and contributed to experiments and/or their analysis and the funding of this project. F.O., C.B., N.R., P.F., A.B. and J.-S.A. wrote and/or edited the manuscript.

## Acknowledgements

We thank Dr Patrick Collombat and Dr Raphael Scharfmann and members of the INSERM U1283/CNRS UMR 8199 for helpful discussions, and Céline Gheeraert for excellent help with ChIP experiments. Human islets were provided through the JDRF award 31-2008-416 (ECIT Islet for Basic Research program). The authors thank the Experimental Resources platform from Université de Lille, especially Cyrille Degraeve, Yann Lepage, Mélanie Besegher and Julien Devassine for animal care. We thank the Department of Histology from the Lille Medicine Faculty, particularly M.H. Gevaert and R.M. Siminski, for histological preparations. This work was supported by grants from « European Genomic Institute for Diabetes » E.G.I.D, ANR-10-LABX-46 and Equipex 2010 ANR-10-EQPX-07-01; ’LIGAN-PM’ Genomics platform, a French State fund managed by the Agence Nationale de la Recherche under the frame program Investissements d’Avenir I-SITE ULNE / ANR-16-IDEX-0004 ULNE (to P.F., A.B. and J-S. A), Agence Nationale pour la Recherche (BETAPLASTICITY, ANR-17-CE14-0034 to J-S. A.), European Foundation for the Study of Diabetes (EFSD, to J-S.A.), European Commission, European Research Council (GEPIDIAB 294785 to P.F.), INSERM, CNRS, Institut Pasteur de Lille, Association pour la Recherche sur le Diabète (to J-S.A.), Université de Lille (to F.O., C.B., X.G., N.R. and J-S.A.), I-SITE ULNE (EpiRNAdiab Sustain grant to J-S.A.), Conseil Régional Hauts de France and Métropole Européenne de Lille (to X.G., N.R. and J-S.A.), F.E.D.E.R. (Fonds Européen de Développement Régional, to N.R., P.F. and J-S.A.) and Société Francophone du Diabète (to S.A.H. and J-S.A).

