## Supplemental Data 1 for "Pancreatic β-cell specific loss of *E2f1* impairs insulin secretion and β-cell identity through the epigenetic repression of non β-cell programs"

**Supplemental information**

**Supplemental figures and legends**

**Supplementary Figure S1, related to Figure 1. The E2f1-CDK4-Cdkn2a pathway is required to maintain normal α-to-β cell ratio. (A)** Scheme representing the different genetically-engineered mouse models used in this study. **(B to F)**. Representative immunofluorescent staining of insulin and glucagon in pancreatic sections from control male *E2f1*$^{+/+}$ (B), CMV-CDK4$^{R24C}$ (C), *E2f1*$^{-/-}$::CMV-CDK4$^{R24C}$ (D), *Cdkn2a*$^{-/-}$ (E), *E2f1*$^{-/-}$::*Cdkn2a*$^{-/-}$ (F) mice. (**G**) Quantification of glucagon labelled cells (Glucagon +) and insulin labelled cells (Insulin +) from (B to F). Values in G are expressed as mean ± s.e.m. and were analysed by one-way ANOVA with Tukey's test. ***p<0.001 and ****p<0.0001 compared to *E2f1*$^{+/+}$; $^{\#\#\#\#}$p<0.0001 compared to *E2f1*$^{-/-}$.

**Supplementary Figure S2, related to Figure 2. Validation of the *E2f1*$^{ß-/-}$ mouse model. (A)** qPCR-based analysis of *E2f1* mRNA expression in different tissues (pancreatic islets, hypothalamus (Hypo), heart, epididymal white adipose tissue (eWAT), brown adipose tissue (BAT) and muscle) isolated from 3 month old male *E2f1*$^{ß-/-}$, RIP-Cre$^{/+}$ and *E2f1*$^{fl/fl}$ control mice (n=3-6). **(B)** Representative immunofluorescent staining of E2f1, insulin and glucagon in pancreatic sections from 3 month old *E2f1*$^{ß-/-}$ and *E2f1*$^{fl/fl}$ control male mice. **(C and D)** Body weight (C, n=5-8) and fasting glycaemia (D, n=7-12) of 3 month old male *E2f1*$^{ß-/-}$, RIP-Cre$^{/+}$ and control *E2f1*$^{fl/fl}$ mice. **(E)** Cell number per islet section in 3 month old male *E2f1*$^{ß-/-}$ and *E2f1*$^{fl/fl}$ mice. Statistical analysis for A, C, D, E were performed using one-way ANOVA with Tukey's test. ***p<0.005. Results are represented as mean ± s.e.m.

**Supplementary Figure S3, related to Figure 3. Gene expression analysis of pancreatic islets isolated from β cell-specific *E2f1* knockout mouse (E2f1$^{β-/-}$) compared to littermate controls (*E2f1*$^{fl/fl}$). (A)** Aligned reads and sequencing coverage of *E2f1* gene exons in *E2f1*$^{fl/fl}$ and *E2f1*$^{β-/-}$ isolated islets in RNA-sequencing experiments. **(B and C)** Heatmap from gene set enrichment analysis (GSEA) displaying differentially expressed β-cell specific genes **(B)** and α-cell specific genes **(C)** in islets isolated from *E2f1*$^{β-/-}$ and *E2f1*$^{fl/fl}$ mice. **(D)** qPCR-based analysis of *E2f1,*

*Pcsk9, Foxo1* and some β-cell specific genes (*Pdx1*, *Mafa*, *Ins2, Glp1r*) in islets isolated from 6 month old $E2f1^{ß-/-}$ and $E2f1^{fl/fl}$ control male mice (n=5-11). **(E)** qPCR-based analysis of *Arx* (α-cell specific gene) in islets isolated from 6 month old $E2f1^{ß-/-}$ and $E2f1^{fl/fl}$ control male mice (n=3-5). Results are represented as mean ± s.e.m. Statistical analysis for D and E were performed using one-way ANOVA with Tukey's test. *p < 0.05; **p<0.01; ***p<0.001.

**Supplementary Figure S4, related to Figure 5. Chromatin state of specific examples of *E2f1*$^{β-/-}$ up- and down- regulated genes in pancreatic islets and Min6 cells.** IGB (Integrated Genome Browser, (Nicol *et al*, 2009)) was used to visualize H3K4m3, H3K27ac and H3K27me3 ChIP-seq signal intensities from Min6 cells and mouse pancreatic islets (Lu *et al.*, 2018) within a series of **(A)** *E2f1*$^{β-/-}$ down-regulated genes and **(B)** *E2f1*$^{β-/-}$ up-regulated genes.

**Supplementary Figure S5, related to Table 2. Overlap of proteins identified in each RIME replicates.** Venn diagram displaying the overlap of proteins identified for each replicate of Flag-E2F1 and IgG RIME in the Min6 E2F1 flag sample or the negative controls (Min6 pCNA Flag sample).

**Supplementary Table S1, related to Figure 4.** Donor informations.

**Supplementary Table S2, related to Figures 4, S2 and S3.** List of oligonucleotides used in this study.

**Supplementary Table S3, related to Figures 5 and S4.** List of ChIP-seq and RNA-seq data sets used in this study.

**Supplementary Table S4, related to Figures 3 and S3.** List of transcripts differentially regulated in $E2f1^{β-/-}$ and *E2f1*$^{fl/fl}$ pancreatic islets identified by RNA-sequencing and their chromatin state.

Oger *et al.*, supplementary figure S1

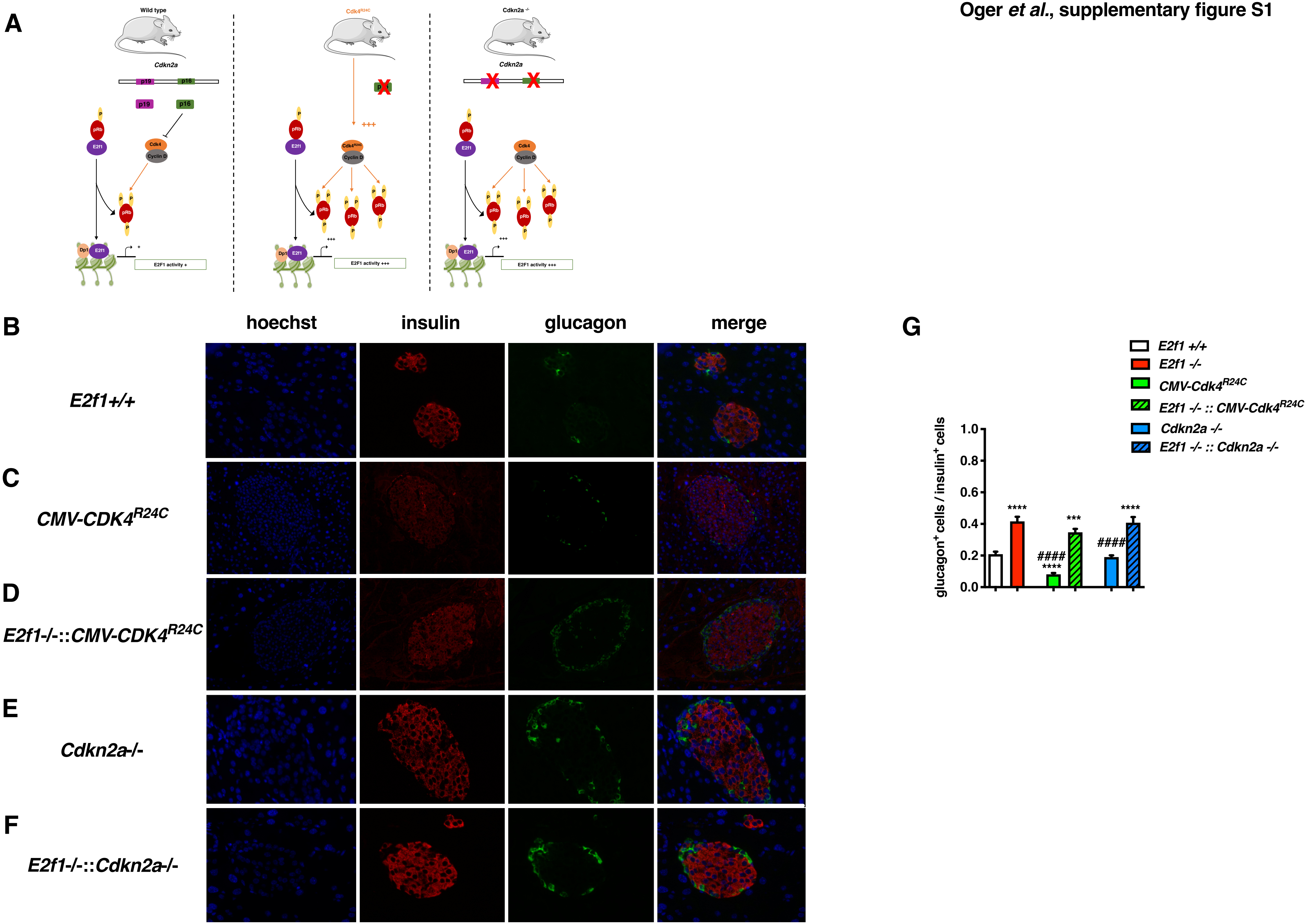

Oger *et al.*, supplementary figure S2

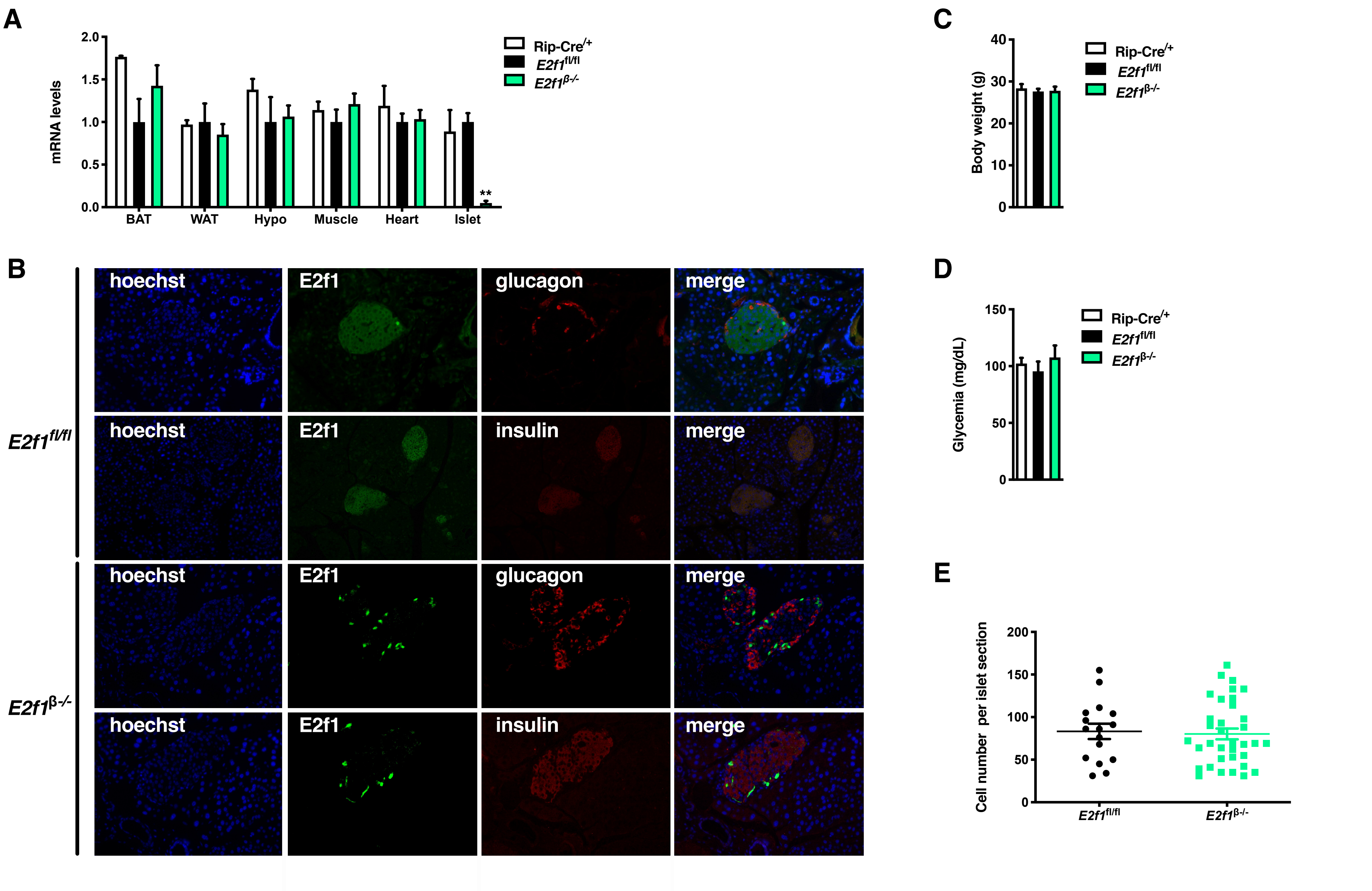

Oger *et al.*, supplementary figure S3

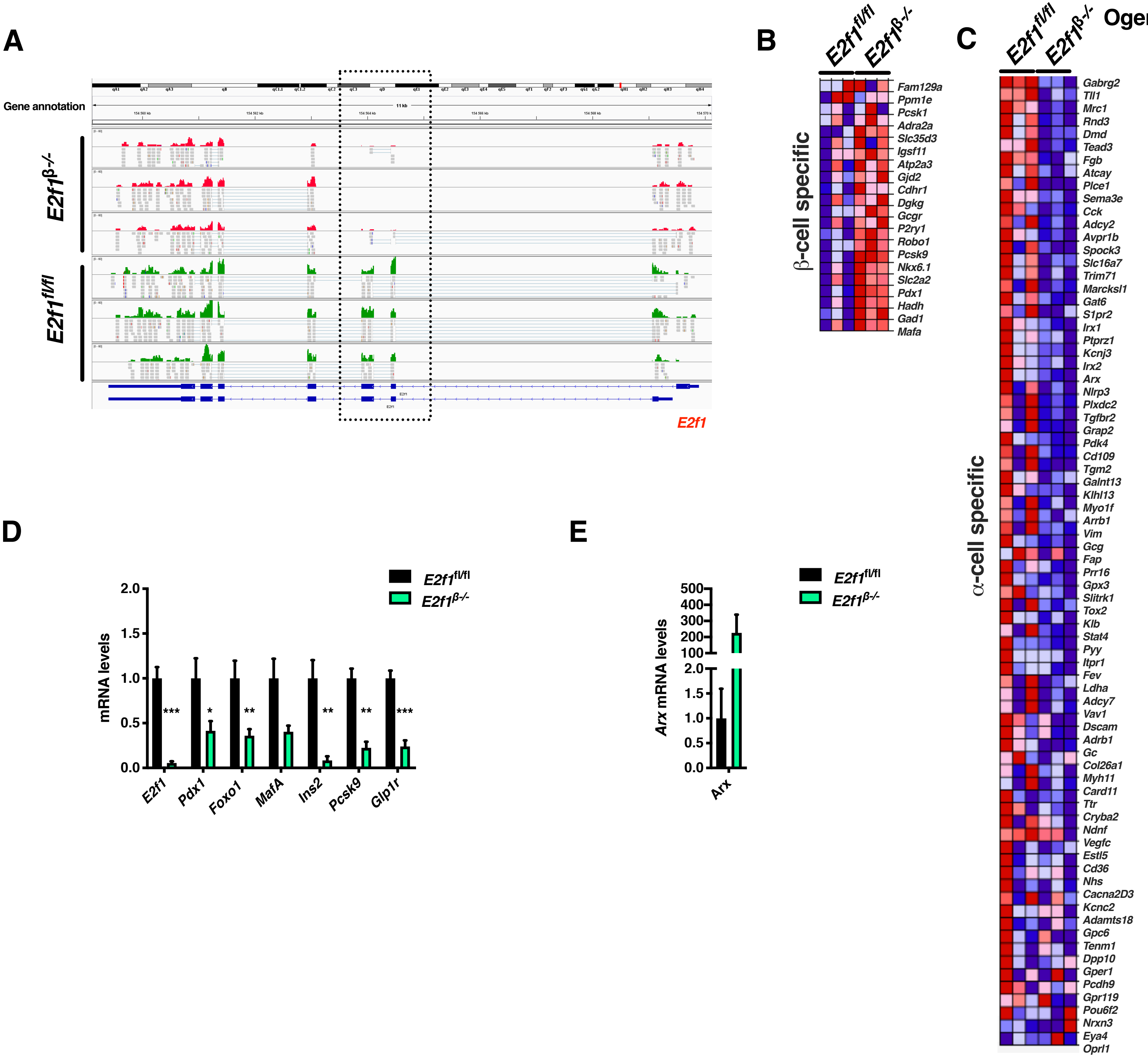

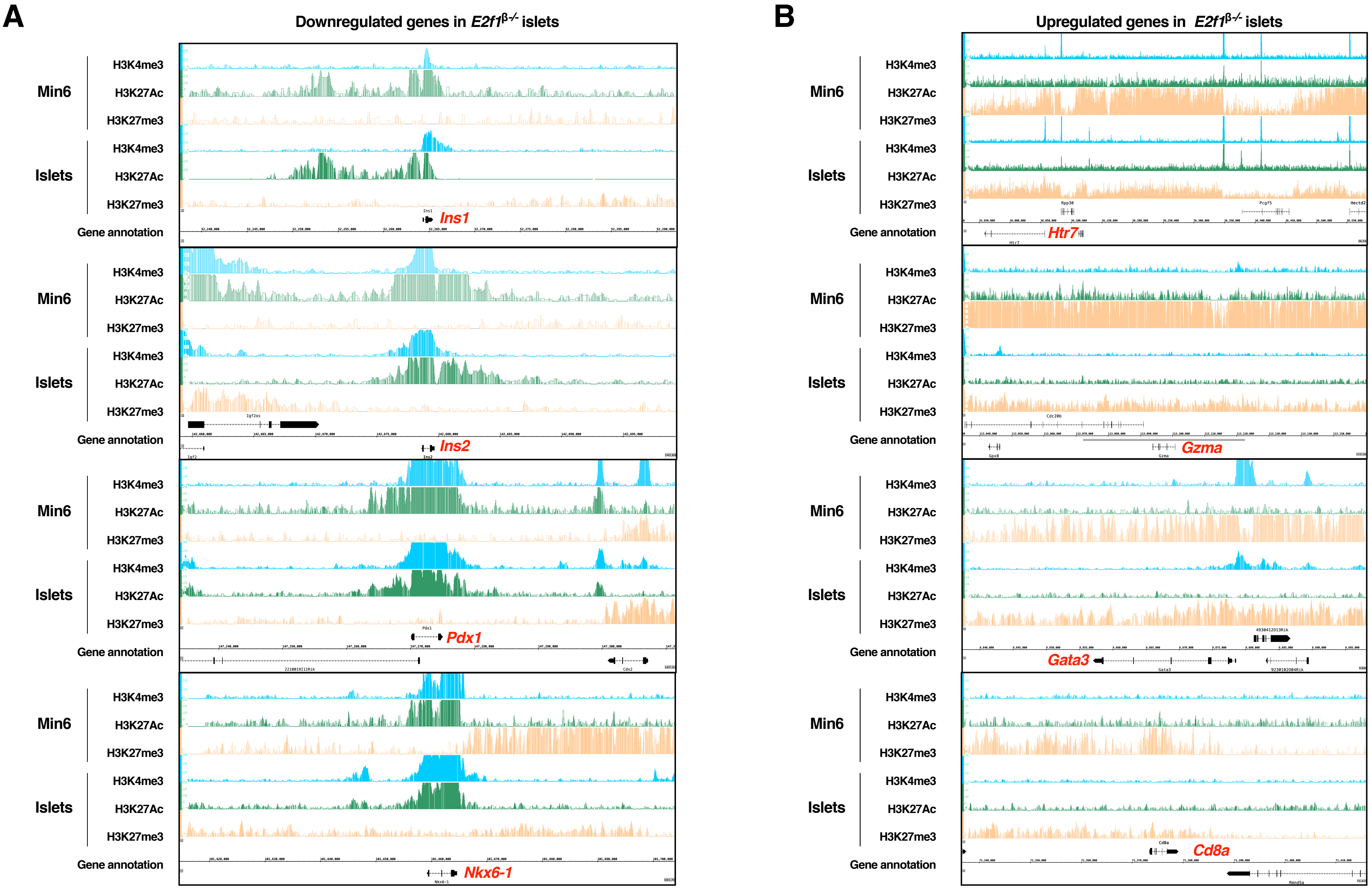
A
Downregulated genes in E2f1β-/- islets
B
Upregulated genes in E2f1β-/- islets
Min6
Islets
H3K4me3
H3K27Ac
H3K27me3
Gene annotation
Ins1
Ins2
Pdx1
Nkx6-1
Htr7
Gzma
Gata3
Cd8a

A

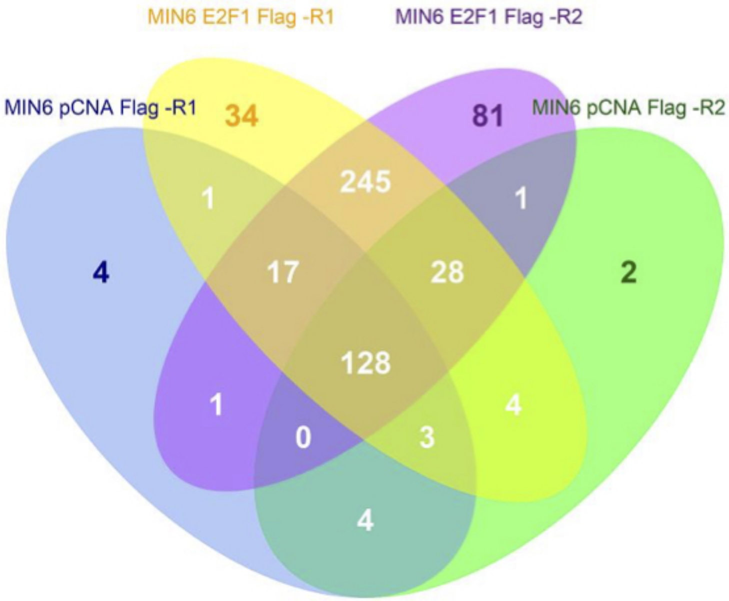
MIN6 E2F1 Flag -R1
MIN6 E2F1 Flag -R2
MIN6 pCNA Flag -R1
MIN6 pCNA Flag -R2
34
81
245
1
1
4
17
28
2
128
1
4
0
3
4

Supplementary Table S1, related to Figure 4. Donor informations.

| Islet number | Condition | Age | BMI | Hba1c | Gender |
|---|---|---|---|---|---|
| H937 | Control islets | 54 | 24.1 | N.D* | M |
| H940 | Control islets | 53 | 32,8 | 6.2% | M |
| H949 | Control islets | 60 | 24,8 | 5.8% | F |

* Not determined

| Cause of death |
| --- |
| Subarachnoid hemorrhage |
| Subarachnoid hemorrhage |
| Stroke |

Supplementary Table S2, related to Figure 4, S2 and S3. List of oligonuc

| Gene Name | Gene symbol | Species | Primer |
|---|---|---|---|
| *Cyclophilin* | *Cyclo* | Mouse | ATGGCACTGGCGGCAGGTCC |
| | | | TTGCCATTCCTGGACCCAAA |
| *E2f1* | *E2f1* | Mouse | GATCGAAGCTTTAATGGAGCG |
| | | | CCCTTGCTTCAGAGAACAGG |
| *E2f1 Exon3* | *E2f1 Exon3* | Mouse | ACAGCTGCAACTGCTTTCGGAG |
| | | | AGCTTGTAGTTGGGTCTCAGGAGG |
| *Paired Box 4* | *Pax4* | Mouse | GTGTACCCTCAGCTGCCTTG |
| | | | ATAGGCCTGGGATGAGGTGT |
| *Pancreatic and duodenal homebox 1* | *Pdx1* | Mouse | ATTGTGCGGTGACCTCGGGC |
| | | | GATGCTGGAGGGCTGTGGCG |
| *Aristaless-related homebox* | *Arx* | Mouse | GCGGGCTTTCCAGAAGACGC |
| | | | CTGGAACCACACCTGGACTCGG |
| *Homebox protein Nkx2.2* | *Nkx2.2* | Mouse | GTGCAGGGAGTATTGGAGGC |
| | | | GAAGGGCCAGAGGAGGAGA |
| *Forkhead box protein O1* | *Foxo1* | Mouse | TGCCCAACCAAAGCTTCCCACA |
| | | | TGGACTGCTCCTCAGTTCCTGCT |
| *Insulin1* | *Ins1* | Mouse | GCCAAACAGCAAAGTCCAGG |
| | | | GTTGAAACAATGACCTGCTTGC |
| *Insulin2* | *Ins2* | Mouse | CAGCAAGCAGGAAGCCTATCT |
| | | | CAGGTGGGAACCACAAAGGT |
| *Glucagon* | *Gcg* | Mouse | AGGCCGAGGAAGGCGAGACT |
| | | | GGAGCCATCAGCGTGCCTGC |
| *Potassium voltage-gated channel subfamily J member 11* | *Kcnj11* | Mouse | CACAAGCTGGGTTGGGGGCTC |
| | | | TGCCCCTCAGCTGGGTTCTGC |
| *Dihydrofolate reductase* | *Dhfr* | Mouse | ATGGTTCGACCATTGAACTGCATCG |
| | | | TCTTCTCAGGAATGGAGAACCAGG |
| *Glucokinase* | *Gck* | Mouse | GCTCAGTGAACCCCGGTCAGC |
| | | | TGTGCGCAGCTGCTCTGAGG |
| *v-maf musculoaponeurotic fibrosarcoma oncogene family, protein A* | *MafA* | Mouse | CCTGTAGAGGAAGCCGAGGAA |
| | | | CCTCCCCCAGTCGAGTATAGC |
| *v-maf musculoaponeurotic fibrosarcoma oncogene family, protein B* | *MafB* | Mouse | TTAGCGCAGACAGAGCTACCGAAA |
| | | | ATACTCTTTACACTCCCACCCTCG |
| *Solute carrier family 2 (facilitated glucose transporter)* | *Slc2a2* | Mouse | AACCGGGATGATTGGCATGT |
| | | | GGCGAATTTATCCAGCAGCA |
| *Proprotein convertase subtilisin/kexin type1* | *Pcsk1* | Mouse | CCGGAGGGCAAGAAGCTGCC |
| | | | TGTTCAGCCCACGTCACACGAT |
| *Proprotein convertase subtilisin/kexin type2* | *Pcsk2* | Mouse | AAAGATGGCGCTGCAACAAG |
| | | | TTGCCCAGTGTTGAACAGGT |
| *Proprotein convertase subtilisin/kexin type9* | *Pcsk9* | Mouse | TTGCAGCAGCTGGGAACTT |
| | | | CCGACTGTGATGACCTCTGGA |

| | | | |
|---|---|---|---|
| *E2F1* | *hE2F1* | Human | ATGGCACTGGCGGCAGGTCC |
| | | | TTGCCATTCCTGGACCCAAA |

| | | | |
|---|---|---|---|
| *Paired Box 4* | *hPAX4* | Human | AGCAGAGGCACTGGAGAAAGAGTT |
| | | | CAGCTGCATTTCCCACTTGAGCTT |
| *Pancreatic and duodenal homebox 1* | *hPDX1* | Human | CGTCCAGCTGCCTTTCCCAT |
| | | | CCGTGAGATGTACTTGTTGAATAGGA |
| *Aristaless-related homebox* | *hARX* | Human | CTGCTGAAACGCAAACAGAGGC |
| | | | CTCGGTCAAGTCCAGCCTCATG |
| *v-maf musculoaponeurotic fibrosarcoma oncogene family, protein A* | *hMAFA* | Human | TTGTACAGGTCCCGCTCTTT |
| | | | AGCAGAGGCACTGGAGAAAGAGTT |
| *Neurogenic Differentiation* | *hNEUROD1* | Human | GCCCCAGGGTTATGAGACTAT |
| | | | GAGAACTGAGACACTCGTCTGT |
| *Insulin* | *hINS* | Human | AGCCTTTGTGAACCAACACC |
| | | | GCTGGTAGAGGGAGCAGATG |
| *Glucagon* | *hGCG* | Human | CATTCACAGGGCACATTCAC |
| | | | CGGCCAAGTTCTTCAACAAT |

Supplementary Table S3, related to Figures 5 and S4 . List of ChIP-seq and RNA-seq data sets used in this study.

| References | Experiment | Mouse strain | Genotype | Biological samples | Antibody | Accession number |
|---|---|---|---|---|---|---|
| Lu et al., Cell Metab, 2018 | ChIP-seq | C57B6/J (8-10 weeks) | WT | Pancreatic islets | H3K4me3 | GSE110648 |
| | | | | | H3K27ac | |
| | | | | | H3K27me3 | |
| This study | ChIP-seq | N/A | N/A | Min6 cells (DMSO) | H3K4me3 | In progress |
| | | | | | H3K27ac | In progress |
| | | | | | H3K27me3 | In progress |
| | | | | Min6 cells (TSA) | H3K27ac | In progress |
| This study | RNA-seq | C57B6/J (12 weeks) | $E2f1^{fl/fl}$ (ctrl) | Pancreatic islets (n=3) | N/A | In progress |
| | | C57B6/J (12 weeks) | $E2f1^{\beta-/-}$(KO) | Pancreatic islets (n=3) | | In progress |
| | | N/A | N/A | Min6 cells (n=3) | | In progress |

| Database | Related figures |
|---|---|
| Gene expression omnibus (https://www.ncbi.nlm.nih.gov/geo/) | Figure 5 and Figure S5 |
| | Figure 5 and Figure S5 |
| | Figure 5 and Figure S5 |
| | Figure 5 and Figure S5 |
| | Figure 5 and Figure S5 |
| | Figure 5 and Figure S5 |
| | Figure 5 and Figure S5 |
| | Figure 3, Figure 5 and Figure S3 |
| | Figure 3, Figure 5 and Figure S3 |
| | Figure 5 |

y regulated in E2f1🚲-/- and E2f1fl/fl pancreatic islets identified by RNA-sequencing and their chromatin state.

| | baseMean | log2FoldChange | lfcSE | stat | pvalue | padj | ensembl_gene_id | external_gene_name | tss_chromatine_ state | A | Active TSS (-1) |
|---|---|---|---|---|---|---|---|---|---|---|---|
| ENSMUSG00000096515 | 493,7382422 | 25,67664338 | 3,906855553 | 6,572201872 | 4,95766E-11 | 8,29732E-08 | ENSMUSG00000096515 | Igkv14-100 | | B | Active TSS (+1) |
| ENSMUSG00000028255 | 15,93590967 | 20,94433456 | 2,81784241 | 7,432755814 | 1,06358E-13 | 3,26342E-10 | ENSMUSG00000028255 | Clca1 | N | C | TSS Flank Cis-reg |
| ENSMUSG00000076576 | 484,1335785 | 12,46127603 | 2,934936975 | 4,245841098 | 2,17775E-05 | 0,002855725 | ENSMUSG00000076576 | Igkv6-32 | | D | Proximal Cis-reg/elongation |
| ENSMUSG00000095335 | 460,3636887 | 12,38863751 | 3,013469649 | 4,111087533 | 3,938E-05 | 0,004393852 | ENSMUSG00000095335 | Igkv3-5 | | E | Proximal Cis-reg/elongation 2 |
| ENSMUSG00000095630 | 697,2342764 | 11,34603634 | 3,589659146 | 3,160755904 | 0,001573603 | 0,044491982 | ENSMUSG00000095630 | Igkv6-23 | | F | Proximal Cis-reg/elongation 3 |
| ENSMUSG00000076556 | 75,3782068 | 9,778196893 | 2,775040987 | 3,523622512 | 0,00042569 | 0,02014642 | ENSMUSG00000076556 | Igkv4-57 | | G | Elongation |
| ENSMUSG00000095442 | 55,29094767 | 9,331579038 | 1,633979485 | 5,710952387 | 1,12346E-08 | 7,38673E-06 | ENSMUSG00000095442 | Ighv1-4 | | H | distal elongation |
| ENSMUSG00000056290 | 46,12129767 | 9,263777657 | 1,696963974 | 5,459030244 | 4,78742E-08 | 2,38207E-05 | ENSMUSG00000056290 | Ms4a4b | W | I | distal elongation 2 |
| ENSMUSG00000021223 | 38,37098841 | 8,772852677 | 1,635152411 | 5,365159002 | 8,08779E-08 | 3,72241E-05 | ENSMUSG00000021223 | Papln | L | J | Active TES/distal enhancer |
| ENSMUSG00000023132 | 33,11067298 | 8,59154443 | 1,726165639 | 4,97724218 | 6,44966E-07 | 0,000194653 | ENSMUSG00000023132 | Gzma | N | K | Bivalent |
| ENSMUSG00000093861 | 917,6157786 | 8,246225838 | 2,267411573 | 3,636845615 | 0,000275997 | 0,015205994 | ENSMUSG00000093861 | Igkv1-110 | | L | Bivalent H3K4me3 High |
| ENSMUSG00000053977 | 26,91486277 | 8,240684634 | 1,731746262 | 4,758598192 | 1,94942E-06 | 0,000484984 | ENSMUSG00000053977 | Cd8a | N | M | Bivalent H3K27me3 High |
| ENSMUSG00000106797 | 19,09370507 | 7,797063646 | 1,602764619 | 4,864759026 | 1,14596E-06 | 0,000314883 | ENSMUSG00000106797 | Gm42936 | | N | PcG repressed |
| ENSMUSG00000024798 | 17,52679147 | 7,789610914 | 1,676962951 | 4,645070369 | 3,39961E-06 | 0,000736315 | ENSMUSG00000024798 | Htr7 | L | O | PcG repressed 2 |
| ENSMUSG00000041449 | 17,16188954 | 7,64681725 | 2,033170045 | 3,761031828 | 0,000169214 | 0,010906871 | ENSMUSG00000041449 | Serpina3h | | P | DNA-meth |
| ENSMUSG00000071716 | 16,84112609 | 7,616630477 | 2,205079796 | 3,454129185 | 0,000552073 | 0,024106381 | ENSMUSG00000071716 | Apol7e | R | Q | Mixed PcG/K9 |
| ENSMUSG00000076652 | 16,09520681 | 7,551529017 | 2,268560172 | 3,328776159 | 0,000872285 | 0,031526499 | ENSMUSG00000076652 | Ighv7-3 | | R | Mixed PcG/K9 |
| ENSMUSG00000015619 | 19,53736019 | 7,489807109 | 1,891461094 | 3,959799719 | 7,50126E-05 | 0,006639341 | ENSMUSG00000015619 | Gata3 | M | S | Heterochromatin |
| ENSMUSG00000076522 | 15,27985042 | 7,475498061 | 2,278786007 | 3,280473917 | 0,001036328 | 0,034878989 | ENSMUSG00000076522 | Igkv16-104 | | T | Heterochromatin 2 |
| ENSMUSG00000009185 | 14,62072498 | 7,413407455 | 1,801918732 | 4,114174143 | 3,88568E-05 | 0,004388673 | ENSMUSG00000009185 | Ccl8 | N | U | Mixed PRC2/K9 |
| ENSMUSG00000095130 | 13,77127619 | 7,327103007 | 2,147052321 | 3,412633654 | 0,000643384 | 0,026321542 | ENSMUSG00000095130 | Ighv1-39 | | V | ZFP genes + repeats |
| ENSMUSG00000083161 | 11,91656623 | 7,289934592 | 1,787173534 | 4,079030075 | 4,5224E-05 | 0,004897492 | ENSMUSG00000083161 | Gm11427 | | W | quiescent |
| ENSMUSG00000094808 | 13,19076197 | 7,264189589 | 1,742007926 | 4,170009492 | 3,04587E-05 | 0,003738297 | ENSMUSG00000094808 | 1810009J06Rik | Q | X | quiescent 2 |
| ENSMUSG00000022491 | 12,83740959 | 7,224913162 | 2,310281854 | 3,127286461 | 0,001764279 | 0,047976931 | ENSMUSG00000022491 | Glycam1 | I | | |
| ENSMUSG00000028167 | 9,510678808 | 7,165071738 | 1,872650456 | 3,826166124 | 0,000130154 | 0,009145587 | ENSMUSG00000028167 | Bdh2 | K | | |
| ENSMUSG00000024211 | 9,201281748 | 7,117088093 | 2,211062626 | 3,218854143 | 0,001287039 | 0,03909966 | ENSMUSG00000024211 | Grm8 | L | | |
| ENSMUSG00000027379 | 9,406954178 | 6,910462926 | 1,920883936 | 3,597543191 | 0,000321237 | 0,01656576 | ENSMUSG00000027379 | Bub1 | L | | |
| ENSMUSG00000090264 | 10,15501244 | 6,886247967 | 1,845322867 | 3,731730685 | 0,000190169 | 0,011788077 | ENSMUSG00000090264 | Eif4ebp3 | G | | |

| | | | | | | | | | |
|---|---|---|---|---|---|---|---|---|---|
| ENSMUSG00000113841 | 9,634962891 | 6,810322486 | 1,878183807 | 3,626014909 | 0,000287829 | 0,01572382 | ENSMUSG00000113841 | Gm48701 | |
| ENSMUSG00000044103 | 8,797830101 | 6,680655999 | 2,021943684 | 3,304076198 | 0,000952899 | 0,033037431 | ENSMUSG00000044103 | Il1f9 | N |
| ENSMUSG00000096375 | 8,30968383 | 6,596255546 | 1,970892533 | 3,346836742 | 0,000817393 | 0,030408377 | ENSMUSG00000096375 | Gm7094 | |
| ENSMUSG00000033882 | 7,814807895 | 6,5281565 | 2,090065342 | 3,123422205 | 0,001787611 | 0,048396943 | ENSMUSG00000033882 | Rbm46 | N |
| ENSMUSG00000081665 | 7,805856912 | 6,506881988 | 1,982376385 | 3,282364559 | 0,001029404 | 0,034837008 | ENSMUSG00000081665 | Gm15922 | |
| ENSMUSG00000022132 | 8,13504613 | 6,433753301 | 1,977689038 | 3,253167296 | 0,001141263 | 0,036603918 | ENSMUSG00000022132 | Cldn10 | N |
| ENSMUSG00000026069 | 272,0811594 | 6,375359854 | 2,042725793 | 3,121006195 | 0,001802342 | 0,048515218 | ENSMUSG00000026069 | Il1rl1 | N |
| ENSMUSG00000100001 | 23,87743635 | 6,346466386 | 1,524909628 | 4,161863937 | 3,1566E-05 | 0,003784327 | ENSMUSG00000100001 | 1810007D17Rik | Q |
| ENSMUSG00000035775 | 22,69017844 | 6,05170726 | 1,544550459 | 3,918102659 | 8,92487E-05 | 0,007335127 | ENSMUSG00000035775 | Krt20 | N |
| ENSMUSG00000058119 | 121,7412868 | 5,970186761 | 0,70309687 | 8,491271992 | 2,04388E-17 | 1,25426E-13 | ENSMUSG00000058119 | Gm5771 | P |
| ENSMUSG00000026012 | 27,42073306 | 5,675164935 | 1,57411845 | 3,605297261 | 0,000311796 | 0,016304531 | ENSMUSG00000026012 | Cd28 | W |
| ENSMUSG00000025058 | 13,19711246 | 5,370625992 | 1,626206094 | 3,30254942 | 0,000958102 | 0,033155373 | ENSMUSG00000025058 | 5430427O19Rik | |
| ENSMUSG00000079455 | 18,31529583 | 5,280215983 | 1,583387894 | 3,334758338 | 0,000853736 | 0,031172947 | ENSMUSG00000079455 | Gm16026 | |
| ENSMUSG00000026622 | 30,65657216 | 5,252353208 | 1,468110524 | 3,577627924 | 0,000346727 | 0,017536361 | ENSMUSG00000026622 | Nek2 | L |
| ENSMUSG00000094561 | 47,65603487 | 5,084640809 | 1,183755905 | 4,295345677 | 1,74421E-05 | 0,002489222 | ENSMUSG00000094561 | Ighv1-22 | |
| ENSMUSG00000021835 | 39,27876404 | 5,064484846 | 1,42572229 | 3,552223936 | 0,00038199 | 0,018653657 | ENSMUSG00000021835 | Bmp4 | M |
| ENSMUSG00000024670 | 37,25426678 | 4,845004518 | 1,093882042 | 4,429183708 | 9,45904E-06 | 0,00155483 | ENSMUSG00000024670 | Cd6 | O |
| ENSMUSG00000071519 | 1206,790689 | 4,729010537 | 0,800137599 | 5,910246613 | 3,41596E-09 | 3,30988E-06 | ENSMUSG00000071519 | Prss3 | T |
| ENSMUSG00000041479 | 24,78766966 | 4,709553933 | 1,18819762 | 3,963611653 | 7,38243E-05 | 0,0065976 | ENSMUSG00000041479 | Syt15 | M |
| ENSMUSG00000071517 | 71,07675696 | 4,640419309 | 0,998127843 | 4,649123196 | 3,33349E-06 | 0,00073059 | ENSMUSG00000071517 | Gm10334 | P |
| ENSMUSG00000029188 | 17,14644104 | 4,601826481 | 1,438779851 | 3,19842294 | 0,001381815 | 0,040997983 | ENSMUSG00000029188 | Slc34a2 | M |
| ENSMUSG00000066513 | 82,081214 | 4,444719588 | 0,813105641 | 5,466349465 | 4,59398E-08 | 2,34931E-05 | ENSMUSG00000066513 | Klk1b4 | N |
| ENSMUSG00000042195 | 89,05745752 | 4,35131527 | 0,738143248 | 5,894946925 | 3,74802E-09 | 3,45005E-06 | ENSMUSG00000042195 | Slc35f2 | L |
| ENSMUSG00000076490 | 30,62424688 | 4,165058627 | 1,305840282 | 3,189562065 | 0,001424885 | 0,041506551 | ENSMUSG00000076490 | Trbc1 | |
| ENSMUSG00000032083 | 50,82340123 | 4,104863884 | 1,248405075 | 3,288086509 | 0,001008708 | 0,034516855 | ENSMUSG00000032083 | Apoa1 | O |
| ENSMUSG00000000782 | 208,8707391 | 4,054063538 | 1,213807761 | 3,339955196 | 0,000837919 | 0,031038411 | ENSMUSG00000000782 | Tcf7 | M |
| ENSMUSG00000078249 | 283,4976504 | 3,874701905 | 0,371997798 | 10,415927 | 2,09744E-25 | 3,86139E-21 | ENSMUSG00000078249 | Hmga1b | |
| ENSMUSG00000053219 | 57,84121675 | 3,853258099 | 0,851034485 | 4,527734383 | 5,96195E-06 | 0,001097594 | ENSMUSG00000053219 | Raet1e | L |
| ENSMUSG00000024857 | 142,5143771 | 3,822309397 | 0,865227417 | 4,417693342 | 9,97598E-06 | 0,001597025 | ENSMUSG00000024857 | Cabp2 | O |
| ENSMUSG00000050232 | 42,7377981 | 3,739944627 | 1,179407369 | 3,171037188 | 0,001518957 | 0,043363298 | ENSMUSG00000050232 | Cxcr3 | |

| | | | | | | | | | |
|---|---|---|---|---|---|---|---|---|---|
| ENSMUSG00000016283 | 271,7240211 | 3,622323657 | 0,667890933 | 5,423525728 | 5,84348E-08 | 2,83101E-05 | ENSMUSG00000016283 | H2-M2 | T |
| ENSMUSG00000029082 | 59,80482846 | 3,613922334 | 1,04708432 | 3,451414815 | 0,000557656 | 0,024106381 | ENSMUSG00000029082 | Bst1 | N |
| ENSMUSG00000066512 | 197,1244242 | 3,514914591 | 0,760120276 | 4,624155811 | 3,76127E-06 | 0,000778034 | ENSMUSG00000066512 | Klk1b5 | N |
| ENSMUSG00000042817 | 45,80278391 | 3,488823231 | 0,986407243 | 3,536899445 | 0,000404854 | 0,019665852 | ENSMUSG00000042817 | Flt3 | L |
| ENSMUSG00000050921 | 72,50075765 | 3,48294648 | 0,969469108 | 3,592632762 | 0,000327354 | 0,016740508 | ENSMUSG00000050921 | P2ry10 | |
| ENSMUSG00000048852 | 80,82502379 | 3,480698754 | 0,786464018 | 4,425757151 | 9,61046E-06 | 0,001556549 | ENSMUSG00000048852 | Gm12185 | K |
| ENSMUSG00000027715 | 48,06128975 | 3,459486926 | 1,05015541 | 3,294261871 | 0,000986805 | 0,033957165 | ENSMUSG00000027715 | Ccna2 | A |
| ENSMUSG00000027358 | 157,1916444 | 3,423850858 | 0,84809304 | 4,037117034 | 5,41121E-05 | 0,005384883 | ENSMUSG00000027358 | Bmp2 | L |
| ENSMUSG00000043003 | 63,67700275 | 3,40152204 | 0,967339238 | 3,516369341 | 0,000437492 | 0,020540463 | ENSMUSG00000043003 | Rasef | M |
| ENSMUSG00000066687 | 25,18424988 | 3,395426509 | 1,017812764 | 3,336003074 | 0,000849922 | 0,031172947 | ENSMUSG00000066687 | Zbtb16 | L |
| ENSMUSG00000027863 | 107,3001113 | 3,267693733 | 1,013133436 | 3,225334015 | 0,001258258 | 0,038866653 | ENSMUSG00000027863 | Cd2 | N |
| ENSMUSG00000027375 | 97,07406245 | 3,256705878 | 0,767776943 | 4,241734411 | 2,21799E-05 | 0,002875578 | ENSMUSG00000027375 | Mal | L |
| ENSMUSG00000045091 | 83,67867019 | 3,246424195 | 0,976809864 | 3,323496531 | 0,000888965 | 0,031716765 | ENSMUSG00000045091 | Aqp12 | N |
| ENSMUSG00000084803 | 27,13973729 | 3,227414022 | 1,030642484 | 3,13145836 | 0,001739405 | 0,047554397 | ENSMUSG00000084803 | 5830444B04Rik | |
| ENSMUSG00000052477 | 196,095592 | 3,216595597 | 0,756426055 | 4,252359601 | 2,1153E-05 | 0,002844135 | ENSMUSG00000052477 | C130026I21Rik | U |
| ENSMUSG00000053886 | 112,2575096 | 3,192679975 | 0,758521497 | 4,20908305 | 2,56409E-05 | 0,003255512 | ENSMUSG00000053886 | Sh2d4a | M |
| ENSMUSG00000102425 | 190,7114927 | 3,147172378 | 0,549719078 | 5,725055767 | 1,034E-08 | 7,3215E-06 | ENSMUSG00000102425 | Gm26616 | |
| ENSMUSG00000021569 | 42,18905338 | 3,106946489 | 0,886426191 | 3,505025596 | 0,000456563 | 0,021118924 | ENSMUSG00000021569 | Trip13 | B |
| ENSMUSG00000001281 | 222,642467 | 3,08556019 | 0,780020183 | 3,955744042 | 7,62968E-05 | 0,006720691 | ENSMUSG00000001281 | Itgb7 | M |
| ENSMUSG00000070034 | 246,1829212 | 2,928708891 | 0,727386877 | 4,026342767 | 5,66511E-05 | 0,005490502 | ENSMUSG00000070034 | Sp110 | |
| ENSMUSG00000030157 | 130,9487314 | 2,80681151 | 0,61797305 | 4,541964262 | 5,57325E-06 | 0,001046975 | ENSMUSG00000030157 | Clec2d | B |
| ENSMUSG00000047415 | 88,44919483 | 2,75915412 | 0,711879464 | 3,875872613 | 0,000106243 | 0,00808239 | ENSMUSG00000047415 | Gpr68 | L |
| ENSMUSG00000027875 | 134,9049591 | 2,733283642 | 0,698002714 | 3,915863915 | 9,00811E-05 | 0,007363407 | ENSMUSG00000027875 | Hmgcs2 | O |
| ENSMUSG00000031957 | 105439,7715 | 2,730132413 | 0,851862125 | 3,2048994 | 0,001351097 | 0,040250873 | ENSMUSG00000031957 | Ctrb1 | O |
| ENSMUSG00000021508 | 159,4372799 | 2,716096425 | 0,84987867 | 3,195863739 | 0,001394129 | 0,041044436 | ENSMUSG00000021508 | Cxcl14 | M |
| ENSMUSG00000068341 | 141,0596367 | 2,707239055 | 0,811761369 | 3,335018341 | 0,000852938 | 0,031172947 | ENSMUSG00000068341 | Reg3d | P |
| ENSMUSG00000033590 | 223,2572187 | 2,700202667 | 0,534037533 | 5,05620392 | 4,27684E-07 | 0,000140601 | ENSMUSG00000033590 | Myo5c | M |
| ENSMUSG00000030219 | 168,6699977 | 2,688833081 | 0,645671914 | 4,164395298 | 3,12179E-05 | 0,003781061 | ENSMUSG00000030219 | Erp27 | O |
| ENSMUSG00000021903 | 59,92691244 | 2,66491067 | 0,683213484 | 3,900553386 | 9,59731E-05 | 0,007550701 | ENSMUSG00000021903 | Galnt15 | M |
| ENSMUSG00000000486 | 142,8768825 | 2,645501092 | 0,83360022 | 3,173584924 | 0,001505688 | 0,043177141 | ENSMUSG00000000486 | 37135 | |

| | | | | | | | | | |
|---|---|---|---|---|---|---|---|---|---|
| ENSMUSG00000039153 | 71,76150486 | 2,637301846 | 0,718959076 | 3,66822248 | 0,000244243 | 0,014333164 | ENSMUSG00000039153 | Runx2 | L |
| ENSMUSG00000002699 | 167,7242627 | 2,578187856 | 0,663859368 | 3,883635573 | 0,000102906 | 0,007861002 | ENSMUSG00000002699 | Lcp2 | N |
| ENSMUSG00000006777 | 140,9115596 | 2,56121887 | 0,760518049 | 3,36772924 | 0,0007579 | 0,029129305 | ENSMUSG00000006777 | Krt23 | N |
| ENSMUSG00000026786 | 120,4959889 | 2,559629641 | 0,818925231 | 3,125596261 | 0,00177445 | 0,048111372 | ENSMUSG00000026786 | Apbb1ip | M |
| ENSMUSG00000036106 | 101,8914186 | 2,55161992 | 0,759873527 | 3,357953433 | 0,000785218 | 0,029683513 | ENSMUSG00000036106 | Prr5 | L |
| ENSMUSG00000010080 | 124,5942993 | 2,544521252 | 0,767670077 | 3,314602625 | 0,000917735 | 0,032243318 | ENSMUSG00000010080 | Epn3 | M |
| ENSMUSG00000029727 | 160,2983861 | 2,518257003 | 0,440914164 | 5,711445008 | 1,12021E-08 | 7,38673E-06 | ENSMUSG00000029727 | Cyp3a13 | O |
| ENSMUSG00000037944 | 310,6562004 | 2,511224559 | 0,66524535 | 3,774884799 | 0,000160082 | 0,01048791 | ENSMUSG00000037944 | Ccr7 | N |
| ENSMUSG00000020911 | 1844,087497 | 2,500837121 | 0,711183169 | 3,516445876 | 0,000437366 | 0,020540463 | ENSMUSG00000020911 | Krt19 | K |
| ENSMUSG00000044309 | 185,2655361 | 2,49982064 | 0,643088439 | 3,887211286 | 0,000101402 | 0,007791756 | ENSMUSG00000044309 | Apol7c | O |
| ENSMUSG00000025813 | 137,1721412 | 2,456675242 | 0,606745632 | 4,048937666 | 5,14506E-05 | 0,005262258 | ENSMUSG00000025813 | Homer2 | N |
| ENSMUSG00000038599 | 199,0762937 | 2,455339588 | 0,715480737 | 3,431734024 | 0,000599735 | 0,025382909 | ENSMUSG00000038599 | Capn8 | I |
| ENSMUSG00000046352 | 250,6111806 | 2,452388121 | 0,680470315 | 3,603960479 | 0,000313405 | 0,016304531 | ENSMUSG00000046352 | Gjb2 | L |
| ENSMUSG00000027356 | 583,1602666 | 2,43779552 | 0,501895072 | 4,857181622 | 1,19068E-06 | 0,000321152 | ENSMUSG00000027356 | Fermt1 | M |
| ENSMUSG00000003882 | 158,2786188 | 2,410441078 | 0,562955747 | 4,281759428 | 1,85421E-05 | 0,002605807 | ENSMUSG00000003882 | Il7r | P |
| ENSMUSG00000033420 | 301,9275266 | 2,407794578 | 0,666394647 | 3,613166143 | 0,000302481 | 0,016124398 | ENSMUSG00000033420 | Antxr1 | L |
| ENSMUSG00000004707 | 197,2640214 | 2,404778127 | 0,691287933 | 3,478692471 | 0,000503866 | 0,022460488 | ENSMUSG00000004707 | Ly9 | P |
| ENSMUSG00000029193 | 164,404075 | 2,361001587 | 0,732854417 | 3,221651575 | 0,00127454 | 0,039023193 | ENSMUSG00000029193 | Cckar | K |
| ENSMUSG00000056306 | 196,8092342 | 2,348393738 | 0,52666089 | 4,459024356 | 8,23336E-06 | 0,001388257 | ENSMUSG00000056306 | Sertm1 | L |
| ENSMUSG00000022901 | 181,2045399 | 2,333819333 | 0,713355288 | 3,271608651 | 0,001069375 | 0,035341356 | ENSMUSG00000022901 | Cd86 | P |
| ENSMUSG00000037849 | 239,1532215 | 2,317181493 | 0,606612092 | 3,819873561 | 0,00013352 | 0,009293801 | ENSMUSG00000037849 | Ifi206 | |
| ENSMUSG00000062991 | 88,3431263 | 2,297519235 | 0,736796287 | 3,118255718 | 0,001819249 | 0,048893972 | ENSMUSG00000062991 | Nrg1 | L |
| ENSMUSG00000071553 | 6096,942173 | 2,289833446 | 0,601964319 | 3,803935506 | 0,000142415 | 0,009674779 | ENSMUSG00000071553 | Cpa2 | M |
| ENSMUSG00000025934 | 187,1042845 | 2,284316202 | 0,567350611 | 4,02628667 | 5,66646E-05 | 0,005490502 | ENSMUSG00000025934 | Gsta3 | N |
| ENSMUSG00000019899 | 247,3834648 | 2,281858863 | 0,545588654 | 4,182379613 | 2,88474E-05 | 0,00358838 | ENSMUSG00000019899 | Lama2 | M |
| ENSMUSG00000028927 | 104,4552052 | 2,27905463 | 0,625204939 | 3,64529211 | 0,000267088 | 0,015038657 | ENSMUSG00000028927 | Padi2 | L |
| ENSMUSG00000030124 | 153,7650045 | 2,270990795 | 0,725280311 | 3,131190468 | 0,001740992 | 0,047554397 | ENSMUSG00000030124 | Lag3 | X |
| ENSMUSG00000063354 | 188,8780169 | 2,266464311 | 0,612552585 | 3,700032236 | 0,000215572 | 0,013004205 | ENSMUSG00000063354 | Slc39a4 | O |
| ENSMUSG00000057969 | 87,80865349 | 2,238186498 | 0,553266083 | 4,045407024 | 5,22323E-05 | 0,005283497 | ENSMUSG00000057969 | Sema3b | O |
| ENSMUSG00000034266 | 121,3555448 | 2,198113965 | 0,703927672 | 3,122641789 | 0,001792357 | 0,048454187 | ENSMUSG00000034266 | Batf | N |

| | | | | | | | | | |
|---|---|---|---|---|---|---|---|---|---|
| ENSMUSG00000055612 | 86,04211026 | 2,190313662 | 0,578238342 | 3,787908034 | 0,000151921 | 0,010138148 | ENSMUSG00000055612 | Cdca7 | B |
| ENSMUSG00000055994 | 43,55919943 | 2,189940064 | 0,661066678 | 3,312737029 | 0,000923878 | 0,032397323 | ENSMUSG00000055994 | Nod2 | O |
| ENSMUSG00000038305 | 84,53369267 | 2,178887113 | 0,585755694 | 3,719788192 | 0,00019939 | 0,012276817 | ENSMUSG00000038305 | Spats2l | L |
| ENSMUSG00000099759 | 363,2053154 | 2,174594301 | 0,428682166 | 5,072742637 | 3,92123E-07 | 0,000133685 | ENSMUSG00000099759 | 1700030C10Rik | |
| ENSMUSG00000089929 | 178,016224 | 2,147772798 | 0,587569163 | 3,65535316 | 0,000256828 | 0,014775646 | ENSMUSG00000089929 | Bcl2a1b | P |
| ENSMUSG00000037946 | 278,5747105 | 2,133888379 | 0,623273501 | 3,423678974 | 0,000617796 | 0,025659629 | ENSMUSG00000037946 | Fgd3 | O |
| ENSMUSG00000035184 | 86,36109763 | 2,121553179 | 0,676446697 | 3,136319816 | 0,001710825 | 0,046939334 | ENSMUSG00000035184 | Fam124a | L |
| ENSMUSG00000074264 | 127,1957947 | 2,11232062 | 0,614417714 | 3,437922724 | 0,000586195 | 0,025097324 | ENSMUSG00000074264 | Amy1 | K |
| ENSMUSG00000037621 | 380,8351325 | 2,092459827 | 0,42176753 | 4,9611686 | 7,00703E-07 | 0,000208064 | ENSMUSG00000037621 | Atoh8 | L |
| ENSMUSG00000041119 | 67,01857038 | 2,088617716 | 0,535020088 | 3,903811769 | 9,46894E-05 | 0,007513933 | ENSMUSG00000041119 | Pde9a | L |
| ENSMUSG00000087259 | 158,3311445 | 2,048365315 | 0,353400704 | 5,796155158 | 6,78525E-09 | 5,38665E-06 | ENSMUSG00000087259 | 2610035D17Rik | L |
| ENSMUSG00000030144 | 330,2249677 | 2,044381428 | 0,512307414 | 3,990536484 | 6,5924E-05 | 0,006068305 | ENSMUSG00000030144 | Clec4d | O |
| ENSMUSG00000000058 | 220,9186651 | 2,036095107 | 0,370373151 | 5,49741552 | 3,85398E-08 | 2,02719E-05 | ENSMUSG00000000058 | Cav2 | L |
| ENSMUSG00000078920 | 347,6375101 | 2,031301269 | 0,509911024 | 3,983638657 | 6,7868E-05 | 0,006216172 | ENSMUSG00000078920 | Ifi47 | K |
| ENSMUSG00000070337 | 121,8999231 | 2,028076486 | 0,616990678 | 3,287045588 | 0,001012444 | 0,034516855 | ENSMUSG00000070337 | Gpr179 | D |
| ENSMUSG00000026994 | 250,9688539 | 2,018715418 | 0,640693466 | 3,15082879 | 0,001628079 | 0,045680808 | ENSMUSG00000026994 | Galnt3 | L |
| ENSMUSG00000073489 | 232,2905104 | 2,017279673 | 0,511262874 | 3,945679949 | 7,95738E-05 | 0,006942912 | ENSMUSG00000073489 | Ifi204 | P |
| ENSMUSG00000106634 | 94,3659941 | 2,011179764 | 0,543411295 | 3,701026794 | 0,000214729 | 0,013004205 | ENSMUSG00000106634 | Gm43042 | S |
| ENSMUSG00000026639 | 386,4440786 | 1,995023604 | 0,510347785 | 3,909145217 | 9,26233E-05 | 0,007426297 | ENSMUSG00000026639 | Lamb3 | O |
| ENSMUSG00000008090 | 103,9437039 | 1,994336308 | 0,536770855 | 3,715433296 | 0,000202856 | 0,012448583 | ENSMUSG00000008090 | Fgfrl1 | L |
| ENSMUSG00000071714 | 466,055316 | 1,993672862 | 0,526491311 | 3,786715605 | 0,000152652 | 0,010145548 | ENSMUSG00000071714 | Csf2rb2 | N |
| ENSMUSG00000029771 | 212,6141134 | 1,966998738 | 0,524311564 | 3,751583743 | 0,000175721 | 0,01107885 | ENSMUSG00000029771 | Irf5 | L |
| ENSMUSG00000015396 | 274,9966671 | 1,952864979 | 0,517774746 | 3,771649729 | 0,000162172 | 0,010522153 | ENSMUSG00000015396 | Cd83 | L |
| ENSMUSG00000024043 | 90,85308316 | 1,949157393 | 0,570372508 | 3,417341062 | 0,00063236 | 0,02604418 | ENSMUSG00000024043 | Arhgap28 | B |
| ENSMUSG00000061353 | 1415,013159 | 1,946329921 | 0,293918533 | 6,622004739 | 3,5436E-11 | 6,88589E-08 | ENSMUSG00000061353 | Cxcl12 | L |
| ENSMUSG00000024778 | 203,2718547 | 1,934088169 | 0,511146073 | 3,783826721 | 0,000154435 | 0,010227176 | ENSMUSG00000024778 | Fas | L |
| ENSMUSG00000032854 | 207,3849468 | 1,923336793 | 0,578194738 | 3,326451566 | 0,000879593 | 0,031526499 | ENSMUSG00000032854 | Ugt8a | O |
| ENSMUSG00000046056 | 111,716243 | 1,920500131 | 0,568969338 | 3,375401807 | 0,00073708 | 0,028627934 | ENSMUSG00000046056 | Sbsn | O |
| ENSMUSG00000037370 | 189,1189288 | 1,919581361 | 0,602440665 | 3,186340949 | 0,001440847 | 0,041904996 | ENSMUSG00000037370 | Enpp1 | B |
| ENSMUSG00000027864 | 650,9162172 | 1,857941736 | 0,336161526 | 5,526931528 | 3,2588E-08 | 1,81489E-05 | ENSMUSG00000027864 | Ptgfrn | B |

| | | | | | | | | | |
|---|---|---|---|---|---|---|---|---|---|
| ENSMUSG00000034463 | 725,9330609 | 1,84932063 | 0,41791366 | 4,425126067 | 9,6386E-06 | 0,001556549 | ENSMUSG00000034463 | Scara3 | L |
| ENSMUSG00000053559 | 86,91063235 | 1,847291883 | 0,477948776 | 3,865041564 | 0,00011107 | 0,008204797 | ENSMUSG00000053559 | Smagp | L |
| ENSMUSG00000006731 | 508,0715103 | 1,839221422 | 0,52163679 | 3,525865998 | 0,0004221 | 0,020072827 | ENSMUSG00000006731 | B4galnt1 | L |
| ENSMUSG00000045994 | 111,3807352 | 1,837926632 | 0,418217057 | 4,394671625 | 1,1094E-05 | 0,001760699 | ENSMUSG00000045994 | B3gat1 | N |
| ENSMUSG00000041734 | 373,7412006 | 1,824416959 | 0,483747114 | 3,771427065 | 0,000162317 | 0,010522153 | ENSMUSG00000041734 | Kirrel | L |
| ENSMUSG00000023348 | 136,6993142 | 1,821911185 | 0,532660387 | 3,420399245 | 0,000625293 | 0,025810859 | ENSMUSG00000023348 | Trip6 | M |
| ENSMUSG00000038128 | 130,6794909 | 1,81379695 | 0,508385226 | 3,567760937 | 0,000360045 | 0,018061101 | ENSMUSG00000038128 | Camk4 | L |
| ENSMUSG00000000628 | 218,9622236 | 1,801216937 | 0,563345328 | 3,197358438 | 0,001386925 | 0,040997983 | ENSMUSG00000000628 | Hk2 | L |
| ENSMUSG00000001761 | 227,4593755 | 1,792863736 | 0,421909192 | 4,249406674 | 2,14337E-05 | 0,002844135 | ENSMUSG00000001761 | Smo | L |
| ENSMUSG00000040026 | 1187,801305 | 1,778324649 | 0,335377948 | 5,302449547 | 1,14259E-07 | 4,7807E-05 | ENSMUSG00000040026 | Saa3 | M |
| ENSMUSG00000020282 | 296,0072115 | 1,767676451 | 0,27685528 | 6,384839212 | 1,71578E-10 | 2,10583E-07 | ENSMUSG00000020282 | Rhbdf1 | L |
| ENSMUSG00000048070 | 160,66973 | 1,735354612 | 0,554117544 | 3,131744575 | 0,00173771 | 0,047554397 | ENSMUSG00000048070 | Pirt | N |
| ENSMUSG00000032625 | 292,8382 | 1,724026072 | 0,394686112 | 4,368094087 | 1,25336E-05 | 0,001955446 | ENSMUSG00000032625 | Thsd7a | L |
| ENSMUSG00000046818 | 135,9375471 | 1,715615211 | 0,439778615 | 3,901088302 | 9,57612E-05 | 0,007550701 | ENSMUSG00000046818 | Ddit4l | L |
| ENSMUSG00000079363 | 1718,259507 | 1,711835335 | 0,265312885 | 6,452137948 | 1,10283E-10 | 1,45022E-07 | ENSMUSG00000079363 | Gbp4 | K |
| ENSMUSG00000029338 | 307,6649706 | 1,711186401 | 0,448242985 | 3,817541953 | 0,000134788 | 0,009293801 | ENSMUSG00000029338 | Antxr2 | L |
| ENSMUSG00000035314 | 138,283519 | 1,709195371 | 0,378993762 | 4,509824545 | 6,48813E-06 | 0,001159713 | ENSMUSG00000035314 | Gdpd5 | L |
| ENSMUSG00000021097 | 390,444565 | 1,707649706 | 0,255803368 | 6,675634173 | 2,46165E-11 | 5,66488E-08 | ENSMUSG00000021097 | Clmn | B |
| ENSMUSG00000028982 | 183,0787657 | 1,701846893 | 0,39014978 | 4,362034734 | 1,28858E-05 | 0,001972375 | ENSMUSG00000028982 | Slc25a33 | L |
| ENSMUSG00000061132 | 430,8863407 | 1,687688551 | 0,537800462 | 3,138131462 | 0,001700286 | 0,046789634 | ENSMUSG00000061132 | Blnk | K |
| ENSMUSG00000095115 | 1023,111809 | 1,680526932 | 0,314313315 | 5,346661594 | 8,95913E-08 | 4,02287E-05 | ENSMUSG00000095115 | Itpripl2 | L |
| ENSMUSG00000039981 | 307,3551595 | 1,674920834 | 0,442188774 | 3,787795919 | 0,00015199 | 0,010138148 | ENSMUSG00000039981 | Zc3h12d | L |
| ENSMUSG00000020303 | 237,8109374 | 1,674692458 | 0,445417454 | 3,759826745 | 0,000170031 | 0,010906871 | ENSMUSG00000020303 | Stc2 | L |
| ENSMUSG00000028937 | 456,5955846 | 1,672639767 | 0,406830924 | 4,111387974 | 3,93288E-05 | 0,004393852 | ENSMUSG00000028937 | Acot7 | L |
| ENSMUSG00000039997 | 277,1492292 | 1,660249919 | 0,488510832 | 3,398593872 | 0,000677332 | 0,027250507 | ENSMUSG00000039997 | Ifi203 | O |
| ENSMUSG00000002930 | 456,2295076 | 1,652470928 | 0,51492954 | 3,209120473 | 0,001331417 | 0,039945611 | ENSMUSG00000002930 | Ppp1r17 | O |
| ENSMUSG00000006519 | 554,3481545 | 1,65225171 | 0,453505714 | 3,643287527 | 0,000269178 | 0,015071561 | ENSMUSG00000006519 | Cyba | L |
| ENSMUSG00000026712 | 82,37929831 | 1,651273125 | 0,522494168 | 3,160366615 | 0,001575707 | 0,044491982 | ENSMUSG00000026712 | Mrc1 | N |
| ENSMUSG00000024511 | 162,7269012 | 1,649933936 | 0,409540453 | 4,028744717 | 5,60755E-05 | 0,005490502 | ENSMUSG00000024511 | Rab27b | L |
| ENSMUSG00000024349 | 250,5500285 | 1,649710312 | 0,529295408 | 3,116804505 | 0,001828228 | 0,04906366 | ENSMUSG00000024349 | Tmem173 | K |

| | | | | | | | | | |
|---|---|---|---|---|---|---|---|---|---|
| ENSMUSG00000047180 | 3056,954494 | 1,645261007 | 0,253329768 | 6,494542746 | 8,3286E-11 | 1,17946E-07 | ENSMUSG00000047180 | Neurl3 | K |
| ENSMUSG00000054200 | 524,416114 | 1,634165544 | 0,400933156 | 4,075905226 | 4,58357E-05 | 0,004934707 | ENSMUSG00000054200 | Ffar4 | L |
| ENSMUSG00000034528 | 315,7091757 | 1,632116681 | 0,506195164 | 3,224283436 | 0,001262883 | 0,038944192 | ENSMUSG00000034528 | Hsd17b13 | X |
| ENSMUSG00000027376 | 135,6548678 | 1,630977889 | 0,402033827 | 4,056817559 | 4,97459E-05 | 0,005116325 | ENSMUSG00000027376 | Prom2 | M |
| ENSMUSG00000039004 | 184,0497407 | 1,62822264 | 0,465044516 | 3,501218881 | 0,000463135 | 0,021209753 | ENSMUSG00000039004 | Bmp6 | M |
| ENSMUSG00000020592 | 1053,337664 | 1,625415163 | 0,385070686 | 4,22108257 | 2,43132E-05 | 0,00313011 | ENSMUSG00000020592 | Sdc1 | L |
| ENSMUSG00000072812 | 212,30577 | 1,61567904 | 0,479300666 | 3,370909231 | 0,000749205 | 0,028915873 | ENSMUSG00000072812 | Ahnak2 | N |
| ENSMUSG00000025017 | 553,7025633 | 1,611373271 | 0,460671372 | 3,497880201 | 0,000468972 | 0,021370717 | ENSMUSG00000025017 | Pik3ap1 | L |
| ENSMUSG00000041889 | 191,8267198 | 1,610626612 | 0,461545879 | 3,489634911 | 0,000483681 | 0,021771551 | ENSMUSG00000041889 | Shisa4 | L |
| ENSMUSG00000063873 | 141,5828868 | 1,606360782 | 0,427808874 | 3,754856153 | 0,000173441 | 0,011048616 | ENSMUSG00000063873 | Slc24a3 | L |
| ENSMUSG00000034957 | 172,3354614 | 1,589522388 | 0,442094298 | 3,595437429 | 0,000323847 | 0,016653694 | ENSMUSG00000034957 | Cebpa | L |
| ENSMUSG00000051855 | 1565,96506 | 1,576286516 | 0,309940952 | 5,08576394 | 3,66149E-07 | 0,000127185 | ENSMUSG00000051855 | Mest | M |
| ENSMUSG00000037679 | 317,8124647 | 1,569780183 | 0,471935937 | 3,326256936 | 0,000880208 | 0,031526499 | ENSMUSG00000037679 | Inf2 | L |
| ENSMUSG00000011256 | 214,8282829 | 1,560522105 | 0,432502404 | 3,608123543 | 0,00030842 | 0,016199394 | ENSMUSG00000011256 | Adam19 | R |
| ENSMUSG00000024538 | 237,5764176 | 1,556459688 | 0,439898984 | 3,538220694 | 0,000402833 | 0,019619469 | ENSMUSG00000024538 | Ppic | L |
| ENSMUSG00000027074 | 281,252518 | 1,552119908 | 0,388038001 | 3,999917284 | 6,33646E-05 | 0,005951749 | ENSMUSG00000027074 | Slc43a3 | M |
| ENSMUSG00000070530 | 190,295541 | 1,542637337 | 0,474930974 | 3,248129563 | 0,001161664 | 0,036872805 | ENSMUSG00000070530 | Wfdc16 | C |
| ENSMUSG00000012428 | 12022,2227 | 1,53881175 | 0,178604431 | 8,615753501 | 6,94831E-18 | 6,39592E-14 | ENSMUSG00000012428 | Steap4 | M |
| ENSMUSG00000005686 | 498,6185628 | 1,534914372 | 0,453279851 | 3,38624002 | 0,000708574 | 0,028140661 | ENSMUSG00000005686 | Ampd3 | M |
| ENSMUSG00000028776 | 362,4689967 | 1,527846361 | 0,426860562 | 3,579263337 | 0,000344564 | 0,017475002 | ENSMUSG00000028776 | Tinagl1 | L |
| ENSMUSG00000027878 | 973,0957829 | 1,525903649 | 0,418843806 | 3,643132896 | 0,00026934 | 0,015071561 | ENSMUSG00000027878 | Notch2 | L |
| ENSMUSG00000018849 | 961,0814833 | 1,520939927 | 0,229957075 | 6,614016663 | 3,7403E-11 | 6,88589E-08 | ENSMUSG00000018849 | Wwc1 | L |
| ENSMUSG00000029762 | 1447,397882 | 1,518326094 | 0,294885523 | 5,148866165 | 2,62066E-07 | 9,64926E-05 | ENSMUSG00000029762 | Akr1b8 | K |
| ENSMUSG00000044734 | 737,8229843 | 1,516803495 | 0,394290643 | 3,846917302 | 0,000119613 | 0,00860188 | ENSMUSG00000044734 | Serpinb1a | K |
| ENSMUSG00000031714 | 653,6121177 | 1,514922784 | 0,3312601 | 4,573212367 | 4,80303E-06 | 0,000940678 | ENSMUSG00000031714 | Gab1 | B |
| ENSMUSG00000028337 | 360,6760912 | 1,512302684 | 0,432540648 | 3,496325003 | 0,000471714 | 0,021442193 | ENSMUSG00000028337 | Coro2a | X |
| ENSMUSG00000001604 | 83,53673235 | 1,510807787 | 0,461232694 | 3,275586934 | 0,001054427 | 0,035230478 | ENSMUSG00000001604 | Tcea3 | L |
| ENSMUSG00000033174 | 157,9600037 | 1,508991985 | 0,332134406 | 4,543317281 | 5,53758E-06 | 0,001046975 | ENSMUSG00000033174 | Mgll | L |
| ENSMUSG00000049521 | 412,7446004 | 1,507142113 | 0,446385566 | 3,376323581 | 0,000734615 | 0,028592507 | ENSMUSG00000049521 | Cdc42ep1 | B |
| ENSMUSG00000004951 | 870,6102529 | 1,504242197 | 0,278257697 | 5,405932031 | 6,44722E-08 | 3,04342E-05 | ENSMUSG00000004951 | Hspb1 | L |

| | | | | | | | | | |
|---|---|---|---|---|---|---|---|---|---|
| ENSMUSG00000045763 | 1796,533013 | 1,496292453 | 0,296553663 | 5,045604346 | 4,5209E-07 | 0,000146017 | ENSMUSG00000045763 | Basp1 | L |
| ENSMUSG00000027800 | 618,0137188 | 1,493149299 | 0,383282449 | 3,895689208 | 9,79199E-05 | 0,007606351 | ENSMUSG00000027800 | Tm4sf1 | K |
| ENSMUSG00000001249 | 144,1553124 | 1,488800703 | 0,448056189 | 3,322799105 | 0,000891191 | 0,031734664 | ENSMUSG00000001249 | Hpn | O |
| ENSMUSG00000022126 | 751,1271323 | 1,481122167 | 0,351799409 | 4,210132623 | 2,55221E-05 | 0,003255512 | ENSMUSG00000022126 | Acod1 | |
| ENSMUSG00000075012 | 222,8796782 | 1,479180897 | 0,306742944 | 4,822216542 | 1,41972E-06 | 0,000363014 | ENSMUSG00000075012 | Fjx1 | M |
| ENSMUSG00000029484 | 892,5616786 | 1,470970312 | 0,432924188 | 3,39775497 | 0,000679412 | 0,027250507 | ENSMUSG00000029484 | Anxa3 | L |
| ENSMUSG00000014444 | 892,7862086 | 1,465468036 | 0,31449155 | 4,659800996 | 3,16515E-06 | 0,000702054 | ENSMUSG00000014444 | Piezo1 | B |
| ENSMUSG00000020010 | 1020,934892 | 1,461522214 | 0,32127666 | 4,549107964 | 5,38738E-06 | 0,001044018 | ENSMUSG00000020010 | Vnn3 | K |
| ENSMUSG00000024302 | 216,297008 | 1,460864218 | 0,301827204 | 4,840068087 | 1,29795E-06 | 0,00034136 | ENSMUSG00000024302 | Dtna | L |
| ENSMUSG00000026604 | 478,4227093 | 1,456027411 | 0,383124605 | 3,800401727 | 0,000144462 | 0,009741904 | ENSMUSG00000026604 | Ptpn14 | L |
| ENSMUSG00000029860 | 2041,707821 | 1,454645935 | 0,281325643 | 5,170683757 | 2,33239E-07 | 8,94569E-05 | ENSMUSG00000029860 | Zyx | A |
| ENSMUSG00000032068 | 1108,420223 | 1,442318874 | 0,356651534 | 4,044056277 | 5,25343E-05 | 0,005285008 | ENSMUSG00000032068 | Plet1 | D |
| ENSMUSG00000048905 | 1592,848067 | 1,430760829 | 0,29021665 | 4,929975001 | 8,22401E-07 | 0,000240324 | ENSMUSG00000048905 | 4930539E08Rik | K |
| ENSMUSG00000032776 | 651,1895891 | 1,418135323 | 0,428831425 | 3,306976218 | 0,000943089 | 0,032883084 | ENSMUSG00000032776 | Mctp2 | L |
| ENSMUSG00000017639 | 472,7423537 | 1,412877303 | 0,255863725 | 5,521991452 | 3,35179E-08 | 1,81489E-05 | ENSMUSG00000017639 | Rab11fip4 | L |
| ENSMUSG00000027401 | 911,3466028 | 1,406985336 | 0,183049657 | 7,686358764 | 1,51382E-14 | 5,57387E-11 | ENSMUSG00000027401 | Tgm3 | N |
| ENSMUSG00000047878 | 250,713182 | 1,403909271 | 0,430563501 | 3,260632332 | 0,001111641 | 0,036022203 | ENSMUSG00000047878 | A4galt | L |
| ENSMUSG00000055044 | 792,3899927 | 1,401136554 | 0,324714221 | 4,314983644 | 1,59615E-05 | 0,002313785 | ENSMUSG00000055044 | Pdlim1 | L |
| ENSMUSG00000002565 | 237,787529 | 1,399818374 | 0,344937374 | 4,058181226 | 4,94564E-05 | 0,005116325 | ENSMUSG00000002565 | Scin | L |
| ENSMUSG00000030616 | 495,5789006 | 1,386357344 | 0,400285839 | 3,463418412 | 0,000533358 | 0,023547066 | ENSMUSG00000030616 | Sytl2 | Q |
| ENSMUSG00000003617 | 1471,507295 | 1,383016705 | 0,323918606 | 4,269642683 | 1,95786E-05 | 0,002710991 | ENSMUSG00000003617 | Cp | K |
| ENSMUSG00000026249 | 735,433087 | 1,374013015 | 0,422040525 | 3,255642372 | 0,001131362 | 0,036413229 | ENSMUSG00000026249 | Serpine2 | L |
| ENSMUSG00000024197 | 659,4728485 | 1,370720611 | 0,321415986 | 4,264631105 | 2,00233E-05 | 0,002730581 | ENSMUSG00000024197 | Plin3 | L |
| ENSMUSG00000032038 | 375,9580472 | 1,36949221 | 0,373363227 | 3,667989 | 0,000244466 | 0,014333164 | ENSMUSG00000032038 | St3gal4 | B |
| ENSMUSG00000045103 | 257,7790424 | 1,364789615 | 0,383845217 | 3,555572805 | 0,000377156 | 0,018653657 | ENSMUSG00000045103 | Dmd | |
| ENSMUSG00000090175 | 470,3377357 | 1,363631058 | 0,345859317 | 3,942733334 | 8,05582E-05 | 0,006977316 | ENSMUSG00000090175 | Ugt1a9 | K |
| ENSMUSG00000023972 | 446,679564 | 1,360431446 | 0,347532745 | 3,914541764 | 9,05761E-05 | 0,007363407 | ENSMUSG00000023972 | Ptk7 | L |
| ENSMUSG00000020173 | 671,1241906 | 1,352535407 | 0,279619783 | 4,837051919 | 1,31779E-06 | 0,000341697 | ENSMUSG00000020173 | Cobl | X |
| ENSMUSG00000004631 | 905,4237463 | 1,351708519 | 0,346046933 | 3,906142171 | 9,37813E-05 | 0,007474089 | ENSMUSG00000004631 | Sgce | L |
| ENSMUSG00000042745 | 775,9098005 | 1,349359697 | 0,277930073 | 4,855033079 | 1,20367E-06 | 0,000321152 | ENSMUSG00000042745 | Id1 | L |

| | | | | | | | | | |
|---|---|---|---|---|---|---|---|---|---|
| ENSMUSG00000023249 | 232,1750687 | 1,343346772 | 0,311042007 | 4,318859645 | 1,56837E-05 | 0,00229157 | ENSMUSG00000023249 | Parp3 | K |
| ENSMUSG00000035277 | 1066,697241 | 1,338825965 | 0,381301527 | 3,511200117 | 0,000446088 | 0,020843879 | ENSMUSG00000035277 | Arx | |
| ENSMUSG00000002020 | 411,0252546 | 1,335567158 | 0,402920725 | 3,314714474 | 0,000917368 | 0,032243318 | ENSMUSG00000002020 | Ltbp2 | L |
| ENSMUSG00000050440 | 1702,251358 | 1,335246493 | 0,320879042 | 4,161214414 | 3,1656E-05 | 0,003784327 | ENSMUSG00000050440 | Hamp | O |
| ENSMUSG00000054855 | 366,8968503 | 1,328487473 | 0,282969395 | 4,694809744 | 2,66855E-06 | 0,0006141 | ENSMUSG00000054855 | Rnd1 | L |
| ENSMUSG00000004891 | 1833,338923 | 1,326427478 | 0,336657355 | 3,939992576 | 8,14841E-05 | 0,006977316 | ENSMUSG00000004891 | Nes | M |
| ENSMUSG00000038412 | 743,8694454 | 1,320564387 | 0,236028075 | 5,594946236 | 2,2069E-08 | 1,31061E-05 | ENSMUSG00000038412 | Higd1a | M |
| ENSMUSG00000024664 | 376,4399648 | 1,320081142 | 0,413509709 | 3,192382456 | 0,001411044 | 0,041365148 | ENSMUSG00000024664 | Fads3 | L |
| ENSMUSG00000018920 | 1723,998575 | 1,319552951 | 0,308160612 | 4,282029889 | 1,85196E-05 | 0,002605807 | ENSMUSG00000018920 | Cxcl16 | L |
| ENSMUSG00000024912 | 674,3718306 | 1,318870975 | 0,344984634 | 3,82298469 | 0,000131846 | 0,009229218 | ENSMUSG00000024912 | Fosl1 | L |
| ENSMUSG00000053110 | 411,6568618 | 1,318159957 | 0,395838737 | 3,330042858 | 0,000868326 | 0,031526499 | ENSMUSG00000053110 | Yap1 | L |
| ENSMUSG00000051439 | 792,3817648 | 1,315879779 | 0,401385316 | 3,278345584 | 0,001044175 | 0,035060717 | ENSMUSG00000051439 | Cd14 | M |
| ENSMUSG00000034271 | 276,4334162 | 1,314091206 | 0,364700654 | 3,603204961 | 0,000314317 | 0,016304531 | ENSMUSG00000034271 | Jdp2 | M |
| ENSMUSG00000030342 | 1680,569829 | 1,31038123 | 0,307159116 | 4,26613166 | 1,98891E-05 | 0,002730581 | ENSMUSG00000030342 | Cd9 | L |
| ENSMUSG00000042784 | 1126,254255 | 1,301580661 | 0,231990003 | 5,61050324 | 2,01739E-08 | 1,23801E-05 | ENSMUSG00000042784 | Muc1 | C |
| ENSMUSG00000042035 | 244,9746715 | 1,296383522 | 0,416818484 | 3,110187221 | 0,001869688 | 0,049770225 | ENSMUSG00000042035 | Igsf3 | A |
| ENSMUSG00000105096 | 851,6459807 | 1,293426048 | 0,229814818 | 5,628122927 | 1,82181E-08 | 1,15654E-05 | ENSMUSG00000105096 | Gbp10 | K |
| ENSMUSG00000015312 | 1425,252777 | 1,291326905 | 0,223675756 | 5,773209089 | 7,77758E-09 | 5,72741E-06 | ENSMUSG00000015312 | Gadd45b | A |
| ENSMUSG00000041859 | 403,9687236 | 1,287250447 | 0,384625372 | 3,346764257 | 0,000817607 | 0,030408377 | ENSMUSG00000041859 | Mcm3 | L |
| ENSMUSG00000034930 | 466,2419077 | 1,276359141 | 0,218396993 | 5,844215734 | 5,0896E-09 | 4,25907E-06 | ENSMUSG00000034930 | Rtkn | L |
| ENSMUSG00000042677 | 2381,683954 | 1,270069689 | 0,213688527 | 5,943555816 | 2,78905E-09 | 2,85258E-06 | ENSMUSG00000042677 | Zc3h12a | A |
| ENSMUSG00000040717 | 397,4977149 | 1,269128765 | 0,30908653 | 4,106062999 | 4,0246E-05 | 0,004463424 | ENSMUSG00000040717 | Il17rd | B |
| ENSMUSG00000105383 | 257,640768 | 1,266067273 | 0,38056545 | 3,326805612 | 0,000878476 | 0,031526499 | ENSMUSG00000105383 | NA | |
| ENSMUSG00000075010 | 525,3482214 | 1,257678087 | 0,374034716 | 3,362463519 | 0,000772503 | 0,029421596 | ENSMUSG00000075010 | AW112010 | K |
| ENSMUSG00000047281 | 857,5274312 | 1,248913694 | 0,390828991 | 3,195550288 | 0,001395645 | 0,041044436 | ENSMUSG00000047281 | Sfn | L |
| ENSMUSG00000031825 | 5496,929793 | 1,241791057 | 0,181255263 | 6,851062081 | 7,33037E-12 | 1,92789E-08 | ENSMUSG00000031825 | Crispld2 | M |
| ENSMUSG00000003153 | 325,5033009 | 1,235025745 | 0,289262394 | 4,269568984 | 1,95851E-05 | 0,002710991 | ENSMUSG00000003153 | Slc2a3 | L |
| ENSMUSG00000040249 | 2838,032494 | 1,225895171 | 0,242358749 | 5,058184103 | 4,23268E-07 | 0,000140601 | ENSMUSG00000040249 | Lrp1 | L |
| ENSMUSG00000066800 | 229,9969996 | 1,220097317 | 0,362561792 | 3,365212064 | 0,000764848 | 0,029213404 | ENSMUSG00000066800 | Rnasel | K |
| ENSMUSG00000028028 | 501,4109473 | 1,217365635 | 0,372506054 | 3,268042548 | 0,001082941 | 0,03560168 | ENSMUSG00000028028 | Alpk1 | N |

| | | | | | | | | | |
|---|---|---|---|---|---|---|---|---|---|
| ENSMUSG00000026980 | 819,513466 | 1,199311806 | 0,3096907 | 3,872611631 | 0,000107675 | 0,008124192 | ENSMUSG00000026980 | Ly75 | L |
| ENSMUSG00000037012 | 1333,903084 | 1,19678631 | 0,268483273 | 4,457582389 | 8,28892E-06 | 0,001388257 | ENSMUSG00000037012 | Hk1 | N |
| ENSMUSG00000021700 | 858,6387973 | 1,196041272 | 0,261086939 | 4,581007677 | 4,62741E-06 | 0,000916028 | ENSMUSG00000021700 | Rab3c | L |
| ENSMUSG00000001333 | 358,3233553 | 1,190380075 | 0,2872362 | 4,144255062 | 3,4092E-05 | 0,003997668 | ENSMUSG00000001333 | Sync | B |
| ENSMUSG00000050721 | 656,1617705 | 1,188890252 | 0,362690168 | 3,277977613 | 0,001045537 | 0,035060717 | ENSMUSG00000050721 | Plekho2 | A |
| ENSMUSG00000052920 | 191,0064624 | 1,18779709 | 0,363689447 | 3,265965234 | 0,001090916 | 0,035736248 | ENSMUSG00000052920 | Prkg1 | L |
| ENSMUSG00000038067 | 2307,697039 | 1,184221729 | 0,242076593 | 4,89192993 | 9,9852E-07 | 0,000282812 | ENSMUSG00000038067 | Csf3 | K |
| ENSMUSG00000037405 | 5789,658683 | 1,176574415 | 0,148144759 | 7,942058997 | 1,98852E-15 | 9,15218E-12 | ENSMUSG00000037405 | Icam1 | M |
| ENSMUSG00000036718 | 592,9082672 | 1,173622922 | 0,259387651 | 4,524590576 | 6,05125E-06 | 0,001103006 | ENSMUSG00000036718 | Micall2 | K |
| ENSMUSG00000018774 | 1081,605756 | 1,170254171 | 0,348130459 | 3,361539164 | 0,000775094 | 0,029421596 | ENSMUSG00000018774 | Cd68 | F |
| ENSMUSG00000027583 | 150,0648953 | 1,166897148 | 0,357730691 | 3,261943068 | 0,001106514 | 0,036022203 | ENSMUSG00000027583 | Zbtb46 | B |
| ENSMUSG00000074457 | 601,4570005 | 1,155686406 | 0,324021962 | 3,566691582 | 0,000361517 | 0,018085651 | ENSMUSG00000074457 | S100a16 | K |
| ENSMUSG00000044986 | 121,1118964 | 1,151520005 | 0,363194196 | 3,170535263 | 0,001521584 | 0,043363298 | ENSMUSG00000044986 | Tst | B |
| ENSMUSG00000001750 | 535,4453459 | 1,149877112 | 0,315855023 | 3,64052185 | 0,000272086 | 0,015087661 | ENSMUSG00000001750 | Tcirg1 | L |
| ENSMUSG00000047261 | 552,313033 | 1,137507267 | 0,294417883 | 3,863580761 | 0,000111737 | 0,008204797 | ENSMUSG00000047261 | Gap43 | M |
| ENSMUSG00000045348 | 409,5376867 | 1,126787098 | 0,312790334 | 3,60237186 | 0,000315327 | 0,01630665 | ENSMUSG00000045348 | Nyap1 | L |
| ENSMUSG00000056596 | 686,1646432 | 1,125873337 | 0,271153928 | 4,152155726 | 3,29358E-05 | 0,003886847 | ENSMUSG00000056596 | Trnp1 | L |
| ENSMUSG00000059401 | 993,2867895 | 1,120290026 | 0,275655868 | 4,064089171 | 4,82204E-05 | 0,005072785 | ENSMUSG00000059401 | Mamld1 | |
| ENSMUSG00000030257 | 450,6704845 | 1,115659789 | 0,346409289 | 3,220640508 | 0,001279045 | 0,039023193 | ENSMUSG00000030257 | Srgap3 | B |
| ENSMUSG00000020019 | 611,5342067 | 1,112924063 | 0,296644753 | 3,751706554 | 0,000175635 | 0,01107885 | ENSMUSG00000020019 | Ntn4 | L |
| ENSMUSG00000000409 | 267,8152783 | 1,108044409 | 0,333715069 | 3,320330759 | 0,000899109 | 0,031893234 | ENSMUSG00000000409 | Lck | O |
| ENSMUSG00000034127 | 3002,11689 | 1,097648554 | 0,281721258 | 3,89622197 | 9,77049E-05 | 0,007606351 | ENSMUSG00000034127 | Tspan8 | L |
| ENSMUSG00000035864 | 808,6587635 | 1,091321335 | 0,307023384 | 3,55452188 | 0,000378667 | 0,018653657 | ENSMUSG00000035864 | Syt1 | L |
| ENSMUSG00000028073 | 277,4641033 | 1,081652155 | 0,330859084 | 3,269223083 | 0,001078432 | 0,035516888 | ENSMUSG00000028073 | Pear1 | W |
| ENSMUSG00000055805 | 529,1171583 | 1,080220174 | 0,331222483 | 3,26131295 | 0,001108976 | 0,036022203 | ENSMUSG00000055805 | Fmnl1 | L |
| ENSMUSG00000012519 | 768,8170811 | 1,076756563 | 0,28659933 | 3,757010051 | 0,000171956 | 0,010992022 | ENSMUSG00000012519 | Mlkl | B |
| ENSMUSG00000090877 | 458,9004681 | 1,074853453 | 0,289606887 | 3,711422281 | 0,000206098 | 0,012605526 | ENSMUSG00000090877 | Hspa1b | A |
| ENSMUSG00000019851 | 977,2435441 | 1,06891275 | 0,31594974 | 3,383173378 | 0,000716534 | 0,028347727 | ENSMUSG00000019851 | Perp | A |
| ENSMUSG00000060550 | 1936,217918 | 1,060987897 | 0,334763195 | 3,169368421 | 0,001527706 | 0,043469968 | ENSMUSG00000060550 | H2-Q7 | L |
| ENSMUSG00000063531 | 1061,513588 | 1,059274541 | 0,289092538 | 3,664136578 | 0,000248174 | 0,014394851 | ENSMUSG00000063531 | Sema3e | L |

| | | | | | | | | | |
|---|---|---|---|---|---|---|---|---|---|
| ENSMUSG00000110386 | 2725,086453 | 1,056027181 | 0,291894546 | 3,617838001 | 0,000297074 | 0,015945005 | ENSMUSG00000110386 | Gm42031 | |
| ENSMUSG00000027533 | 1403,844985 | 1,046865199 | 0,316665008 | 3,305907413 | 0,000946694 | 0,032884207 | ENSMUSG00000027533 | Fabp5 | B |
| ENSMUSG00000027398 | 739,7715875 | 1,037783667 | 0,306833928 | 3,38223244 | 0,000718993 | 0,028347727 | ENSMUSG00000027398 | Il1b | O |
| ENSMUSG00000000078 | 2025,887264 | 1,036441029 | 0,229910326 | 4,508022958 | 6,54345E-06 | 0,001159713 | ENSMUSG00000000078 | Klf6 | A |
| ENSMUSG00000035385 | 3292,031895 | 1,035317613 | 0,304110411 | 3,404413577 | 0,000663063 | 0,026928231 | ENSMUSG00000035385 | Ccl2 | N |
| ENSMUSG00000003849 | 1067,644241 | 1,032739785 | 0,315503135 | 3,273310693 | 0,001062956 | 0,035292936 | ENSMUSG00000003849 | Nqo1 | D |
| ENSMUSG00000038463 | 255,1982929 | 1,014364486 | 0,262563935 | 3,863304701 | 0,000111863 | 0,008204797 | ENSMUSG00000038463 | Olfml2b | L |
| ENSMUSG00000022960 | 287,0301027 | 1,010574543 | 0,27310722 | 3,700284968 | 0,000215357 | 0,013004205 | ENSMUSG00000022960 | Donson | A |
| ENSMUSG00000070639 | 1421,174589 | 1,000593964 | 0,232903367 | 4,296176463 | 1,73769E-05 | 0,002489222 | ENSMUSG00000070639 | Lrrc8b | A |
| ENSMUSG00000045312 | 978,0733109 | 0,997353131 | 0,309916066 | 3,218139494 | 0,001290251 | 0,039132639 | ENSMUSG00000045312 | Lhfpl2 | L |
| ENSMUSG00000073409 | 2306,179091 | 0,997148411 | 0,317721331 | 3,138437092 | 0,001698514 | 0,046789634 | ENSMUSG00000073409 | H2-Q6 | L |
| ENSMUSG00000031453 | 659,6741476 | 0,996096158 | 0,299528088 | 3,325551749 | 0,000882437 | 0,031544987 | ENSMUSG00000031453 | Rasa3 | B |
| ENSMUSG00000066026 | 333,6487635 | 0,991101701 | 0,264864372 | 3,741921542 | 0,000182619 | 0,011396635 | ENSMUSG00000066026 | Dhrs3 | L |
| ENSMUSG00000031934 | 586,328822 | 0,990900931 | 0,316916942 | 3,12668968 | 0,001767864 | 0,048003511 | ENSMUSG00000031934 | Panx1 | L |
| ENSMUSG00000003032 | 709,7560929 | 0,988468846 | 0,197429037 | 5,006704495 | 5,53698E-07 | 0,000169893 | ENSMUSG00000003032 | Klf4 | A |
| ENSMUSG00000027347 | 583,7435932 | 0,98526962 | 0,27266628 | 3,613463388 | 0,000302134 | 0,016124398 | ENSMUSG00000027347 | Rasgrp1 | L |
| ENSMUSG00000091387 | 484,7289705 | 0,985184073 | 0,297984402 | 3,306159876 | 0,000945841 | 0,032884207 | ENSMUSG00000091387 | Gcnt4 | B |
| ENSMUSG00000030934 | 1176,886824 | 0,979759541 | 0,291140752 | 3,365243564 | 0,000764761 | 0,029213404 | ENSMUSG00000030934 | Oat | L |
| ENSMUSG00000060477 | 791,1432954 | 0,971025246 | 0,217614794 | 4,46212883 | 8,11494E-06 | 0,001388257 | ENSMUSG00000060477 | Irak2 | A |
| ENSMUSG00000019889 | 1943,976985 | 0,97085881 | 0,246316645 | 3,941507123 | 8,09712E-05 | 0,006977316 | ENSMUSG00000019889 | Ptprk | L |
| ENSMUSG00000031925 | 564,7736695 | 0,96750775 | 0,239339147 | 4,042413289 | 5,29039E-05 | 0,005293263 | ENSMUSG00000031925 | Maml2 | D |
| ENSMUSG00000004044 | 685,7837321 | 0,965029367 | 0,301041839 | 3,205632048 | 0,001347662 | 0,040250873 | ENSMUSG00000004044 | Cavin1 | |
| ENSMUSG00000019539 | 862,0184882 | 0,960857341 | 0,248704658 | 3,863447299 | 0,000111798 | 0,008204797 | ENSMUSG00000019539 | Rcn3 | M |
| ENSMUSG00000047963 | 1060,564858 | 0,957828186 | 0,274728438 | 3,486454452 | 0,000489469 | 0,021924864 | ENSMUSG00000047963 | Stbd1 | A |
| ENSMUSG00000039982 | 450,501155 | 0,956313486 | 0,272006771 | 3,515770886 | 0,000438479 | 0,020540463 | ENSMUSG00000039982 | Dtx4 | B |
| ENSMUSG00000049382 | 37058,38855 | 0,955222173 | 0,206771708 | 4,619694735 | 3,84305E-06 | 0,000786117 | ENSMUSG00000049382 | Krt8 | A |
| ENSMUSG00000104713 | 8600,034222 | 0,953524695 | 0,186039925 | 5,125376684 | 2,96943E-07 | 0,000107191 | ENSMUSG00000104713 | Gbp6 | L |
| ENSMUSG00000002489 | 525,1376559 | 0,950885087 | 0,275770991 | 3,448096855 | 0,000564552 | 0,024283631 | ENSMUSG00000002489 | Tiam1 | O |
| ENSMUSG00000039089 | 299,2746013 | 0,950112817 | 0,298612146 | 3,181762124 | 0,00146382 | 0,042372523 | ENSMUSG00000039089 | L3mbtl3 | L |
| ENSMUSG00000019850 | 3022,25271 | 0,946502636 | 0,234936096 | 4,028766332 | 5,60703E-05 | 0,005490502 | ENSMUSG00000019850 | Tnfaip3 | E |

| | | | | | | | | | |
|---|---|---|---|---|---|---|---|---|---|
| ENSMUSG00000037706 | 6076,760773 | 0,944096709 | 0,227330029 | 4,152978431 | 3,28176E-05 | 0,003886847 | ENSMUSG00000037706 | Cd81 | B |
| ENSMUSG00000085795 | 469,2844355 | 0,940963935 | 0,297461075 | 3,163317876 | 0,001559819 | 0,04424695 | ENSMUSG00000085795 | Zfp703 | B |
| ENSMUSG00000024074 | 765,4114084 | 0,938074783 | 0,273597815 | 3,428663288 | 0,000606562 | 0,025494972 | ENSMUSG00000024074 | Crim1 | B |
| ENSMUSG00000020674 | 1745,20397 | 0,937312657 | 0,259025721 | 3,61860843 | 0,000296191 | 0,015945005 | ENSMUSG00000020674 | Pxdn | L |
| ENSMUSG00000021959 | 805,5878052 | 0,935777159 | 0,257000587 | 3,641147943 | 0,000271425 | 0,015087661 | ENSMUSG00000021959 | Lats2 | B |
| ENSMUSG00000038387 | 722,6098899 | 0,924652125 | 0,26788493 | 3,451676527 | 0,000557115 | 0,024106381 | ENSMUSG00000038387 | Rras | B |
| ENSMUSG00000048489 | 2810,538616 | 0,924429087 | 0,153981683 | 6,003500351 | 1,93108E-09 | 2,09125E-06 | ENSMUSG00000048489 | Depp1 | |
| ENSMUSG00000058952 | 388,1179892 | 0,922627281 | 0,293644364 | 3,141988723 | 0,001678045 | 0,046455351 | ENSMUSG00000058952 | Cfi | L |
| ENSMUSG00000014599 | 6966,989842 | 0,919955566 | 0,174043946 | 5,285765957 | 1,2518E-07 | 5,12125E-05 | ENSMUSG00000014599 | Csf1 | A |
| ENSMUSG00000024816 | 958,0315116 | 0,91554084 | 0,255343985 | 3,585519517 | 0,000336408 | 0,017155868 | ENSMUSG00000024816 | Frmd8 | B |
| ENSMUSG00000032000 | 7737,926095 | 0,907808443 | 0,139767238 | 6,495144753 | 8,29536E-11 | 1,17946E-07 | ENSMUSG00000032000 | Birc3 | B |
| ENSMUSG00000035954 | 993,5170332 | 0,903835936 | 0,245299926 | 3,684615607 | 0,000229048 | 0,013602499 | ENSMUSG00000035954 | Dock4 | A |
| ENSMUSG00000029298 | 3064,196944 | 0,901207523 | 0,201362886 | 4,475539353 | 7,62187E-06 | 0,001336367 | ENSMUSG00000029298 | Gbp9 | K |
| ENSMUSG00000047501 | 14263,69912 | 0,899538843 | 0,175899656 | 5,113931793 | 3,15521E-07 | 0,000111707 | ENSMUSG00000047501 | Cldn4 | A |
| ENSMUSG00000038740 | 474,5688412 | 0,899167654 | 0,264620638 | 3,397949836 | 0,000678929 | 0,027250507 | ENSMUSG00000038740 | Mvb12b | M |
| ENSMUSG00000000861 | 389,9228408 | 0,894636459 | 0,24765992 | 3,612358669 | 0,000303424 | 0,016124398 | ENSMUSG00000000861 | Bcl11a | L |
| ENSMUSG00000016256 | 10454,48054 | 0,892977595 | 0,18582802 | 4,805397988 | 1,54444E-06 | 0,000389495 | ENSMUSG00000016256 | Ctsz | L |
| ENSMUSG00000029919 | 2581,469114 | 0,891955834 | 0,265302247 | 3,362036488 | 0,000773699 | 0,029421596 | ENSMUSG00000029919 | Hpgds | L |
| ENSMUSG00000030166 | 251,4153044 | 0,883604211 | 0,27793213 | 3,179208574 | 0,001476778 | 0,042613605 | ENSMUSG00000030166 | Rad52 | A |
| ENSMUSG00000062661 | 441,026324 | 0,881345912 | 0,269763495 | 3,267105917 | 0,00108653 | 0,035656009 | ENSMUSG00000062661 | Ncs1 | L |
| ENSMUSG00000017144 | 1328,438048 | 0,881066214 | 0,207359111 | 4,24898722 | 2,14739E-05 | 0,002844135 | ENSMUSG00000017144 | Rnd3 | A |
| ENSMUSG00000017652 | 1460,870384 | 0,88053622 | 0,277647193 | 3,171421299 | 0,00151695 | 0,043363298 | ENSMUSG00000017652 | Cd40 | L |
| ENSMUSG00000075415 | 1347,00511 | 0,879762044 | 0,251106067 | 3,503547538 | 0,000459105 | 0,021161273 | ENSMUSG00000075415 | Fnbp1 | A |
| ENSMUSG00000072115 | 5555,308144 | 0,877858657 | 0,202824578 | 4,328167047 | 1,50355E-05 | 0,002214435 | ENSMUSG00000072115 | Ang | C |
| ENSMUSG00000066363 | 1105,28223 | 0,875481561 | 0,220499518 | 3,970446602 | 7,1738E-05 | 0,006474006 | ENSMUSG00000066363 | Serpina3f | N |
| ENSMUSG00000030123 | 2192,987835 | 0,872630125 | 0,269295516 | 3,240418321 | 0,001193545 | 0,037625268 | ENSMUSG00000030123 | Plxnd1 | B |
| ENSMUSG00000079014 | 665,0806467 | 0,863315347 | 0,215437945 | 4,007257617 | 6,14278E-05 | 0,005859517 | ENSMUSG00000079014 | Serpina3i | P |
| ENSMUSG00000024070 | 571,8872553 | 0,862743908 | 0,257988823 | 3,344113505 | 0,00082546 | 0,03063853 | ENSMUSG00000024070 | Prkd3 | B |
| ENSMUSG00000055069 | 304,6786254 | 0,857618319 | 0,267122018 | 3,210586396 | 0,001324644 | 0,039912772 | ENSMUSG00000055069 | Rab39 | A |
| ENSMUSG00000002227 | 1364,184728 | 0,840595497 | 0,160241268 | 5,245811577 | 1,55596E-07 | 6,22722E-05 | ENSMUSG00000002227 | Mov10 | B |

| | | | | | | | | | |
|---|---|---|---|---|---|---|---|---|---|
| ENSMUSG00000027276 | 785,3908154 | 0,839048587 | 0,242935652 | 3,453789428 | 0,000552769 | 0,024106381 | ENSMUSG00000027276 | Jag1 | A |
| ENSMUSG00000035476 | 1266,050071 | 0,838570664 | 0,167250147 | 5,013871007 | 5,33458E-07 | 0,000166457 | ENSMUSG00000035476 | Tab3 | |
| ENSMUSG00000034438 | 1108,476757 | 0,829299122 | 0,155679726 | 5,326956459 | 9,98721E-08 | 4,27592E-05 | ENSMUSG00000034438 | Gbp8 | K |
| ENSMUSG00000060961 | 927,3418024 | 0,827296224 | 0,209334569 | 3,952028691 | 7,74914E-05 | 0,006793417 | ENSMUSG00000060961 | Slc4a4 | B |
| ENSMUSG00000017765 | 1063,015204 | 0,819984447 | 0,214199344 | 3,828137065 | 0,000129117 | 0,009107438 | ENSMUSG00000017765 | Slc12a4 | B |
| ENSMUSG00000024558 | 1064,796632 | 0,81456998 | 0,22320761 | 3,649382653 | 0,000262871 | 0,01498285 | ENSMUSG00000024558 | Mapk4 | A |
| ENSMUSG00000032366 | 5903,222455 | 0,810227522 | 0,181672668 | 4,45982068 | 8,20283E-06 | 0,001388257 | ENSMUSG00000032366 | Tpm1 | L |
| ENSMUSG00000023232 | 733,2553529 | 0,803934106 | 0,228374706 | 3,520241443 | 0,000431154 | 0,020352686 | ENSMUSG00000023232 | Serinc2 | B |
| ENSMUSG00000029761 | 3461,695347 | 0,801907708 | 0,224891532 | 3,565753244 | 0,000362813 | 0,018101302 | ENSMUSG00000029761 | Cald1 | L |
| ENSMUSG00000024885 | 733,7121322 | 0,801865959 | 0,20741039 | 3,866083849 | 0,000110597 | 0,008204797 | ENSMUSG00000024885 | Aldh3b1 | D |
| ENSMUSG00000031628 | 2384,973356 | 0,800504101 | 0,173705567 | 4,608396349 | 4,05787E-06 | 0,000820937 | ENSMUSG00000031628 | Casp3 | A |
| ENSMUSG00000031379 | 505,2731192 | 0,797570922 | 0,232136891 | 3,43577842 | 0,000590854 | 0,025121529 | ENSMUSG00000031379 | Pir | |
| ENSMUSG00000041827 | 1144,151315 | 0,790139422 | 0,251639868 | 3,139961201 | 0,001689702 | 0,046663634 | ENSMUSG00000041827 | Oasl1 | L |
| ENSMUSG00000052560 | 639,5012181 | 0,78826109 | 0,243917007 | 3,231677449 | 0,001230659 | 0,038270996 | ENSMUSG00000052560 | Cpne8 | B |
| ENSMUSG00000038608 | 1253,031885 | 0,786260872 | 0,211115908 | 3,724308986 | 0,000195851 | 0,012099382 | ENSMUSG00000038608 | Dock10 | I |
| ENSMUSG00000058325 | 1353,815459 | 0,784266829 | 0,214178369 | 3,661746201 | 0,000250502 | 0,014456869 | ENSMUSG00000058325 | Dock1 | L |
| ENSMUSG00000047139 | 6974,698281 | 0,782262565 | 0,194914455 | 4,013363534 | 5,98596E-05 | 0,005769051 | ENSMUSG00000047139 | Cd24a | L |
| ENSMUSG00000055172 | 2662,679604 | 0,780668003 | 0,207623989 | 3,760008694 | 0,000169907 | 0,010906871 | ENSMUSG00000055172 | C1ra | L |
| ENSMUSG00000039621 | 2233,421714 | 0,780154776 | 0,219602409 | 3,552578407 | 0,000381475 | 0,018653657 | ENSMUSG00000039621 | Prex1 | B |
| ENSMUSG00000022270 | 2420,329266 | 0,776088052 | 0,233100197 | 3,329418263 | 0,000870276 | 0,031526499 | ENSMUSG00000022270 | Retreg1 | |
| ENSMUSG00000060675 | 1524,414584 | 0,772245431 | 0,187446375 | 4,119820568 | 3,79168E-05 | 0,004362797 | ENSMUSG00000060675 | Pla2g16 | B |
| ENSMUSG00000007872 | 1265,443411 | 0,768232228 | 0,223553438 | 3,436459015 | 0,000589372 | 0,025116504 | ENSMUSG00000007872 | Id3 | L |
| ENSMUSG00000030691 | 1925,729873 | 0,768059579 | 0,19243493 | 3,991269042 | 6,57207E-05 | 0,006068305 | ENSMUSG00000030691 | Fchsd2 | A |
| ENSMUSG00000020400 | 4521,207477 | 0,765667733 | 0,143457702 | 5,337236861 | 9,43737E-08 | 4,13671E-05 | ENSMUSG00000020400 | Tnip1 | D |
| ENSMUSG00000020806 | 961,63189 | 0,76281832 | 0,217538624 | 3,506587956 | 0,000453891 | 0,021101365 | ENSMUSG00000020806 | Rhbdf2 | B |
| ENSMUSG00000040158 | 872,0324626 | 0,762812737 | 0,229965558 | 3,317073847 | 0,000909656 | 0,032081918 | ENSMUSG00000040158 | Tax1bp3 | A |
| ENSMUSG00000034312 | 1135,754338 | 0,756630024 | 0,225914795 | 3,349183154 | 0,000810502 | 0,03032793 | ENSMUSG00000034312 | Iqsec1 | L |
| ENSMUSG00000026421 | 3785,429203 | 0,753833175 | 0,228439352 | 3,29992696 | 0,0009671 | 0,033341405 | ENSMUSG00000026421 | Csrp1 | A |
| ENSMUSG00000006435 | 4419,975017 | 0,748139984 | 0,172254179 | 4,343232699 | 1,40401E-05 | 0,002118678 | ENSMUSG00000006435 | Neurl1a | C |
| ENSMUSG00000037411 | 13897,63324 | 0,7432801 | 0,229945088 | 3,232424338 | 0,001227446 | 0,038235681 | ENSMUSG00000037411 | Serpine1 | C |

| | | | | | | | | | |
|---|---|---|---|---|---|---|---|---|---|
| ENSMUSG00000048218 | 3840,742564 | 0,741983903 | 0,182772573 | 4,059602013 | 4,91564E-05 | 0,005116325 | ENSMUSG00000048218 | Amigo2 | A |
| ENSMUSG00000037434 | 1313,609049 | 0,739985085 | 0,228859186 | 3,23336414 | 0,001223415 | 0,038235681 | ENSMUSG00000037434 | Slc30a1 | A |
| ENSMUSG00000039115 | 452,473176 | 0,738978579 | 0,203927754 | 3,623727342 | 0,000290388 | 0,015816681 | ENSMUSG00000039115 | Itga9 | L |
| ENSMUSG00000028701 | 802,5607005 | 0,735835398 | 0,224255394 | 3,281238344 | 0,001033524 | 0,034848298 | ENSMUSG00000028701 | Lurap1 | L |
| ENSMUSG00000040093 | 672,5164557 | 0,727056235 | 0,219113023 | 3,318179021 | 0,000906064 | 0,032016577 | ENSMUSG00000040093 | Bmf | B |
| ENSMUSG00000086841 | 737,7874245 | 0,726155291 | 0,22452759 | 3,234147269 | 0,001220065 | 0,038229998 | ENSMUSG00000086841 | 2410006H16Rik | |
| ENSMUSG00000046711 | 4679,133652 | 0,723456773 | 0,187842358 | 3,851403806 | 0,000117443 | 0,0084789 | ENSMUSG00000046711 | Hmga1 | A |
| ENSMUSG00000020227 | 1023,534669 | 0,719653709 | 0,226563888 | 3,176383114 | 0,001491239 | 0,042896413 | ENSMUSG00000020227 | Irak3 | L |
| ENSMUSG00000042228 | 2396,393218 | 0,711794553 | 0,194285574 | 3,663651087 | 0,000248645 | 0,014394851 | ENSMUSG00000042228 | Lyn | B |
| ENSMUSG00000005973 | 2201,68985 | 0,711379706 | 0,206307139 | 3,448158448 | 0,000564423 | 0,024283631 | ENSMUSG00000005973 | Rcn1 | B |
| ENSMUSG00000039835 | 2022,292619 | 0,710167155 | 0,149812551 | 4,740371539 | 2,13327E-06 | 0,000510045 | ENSMUSG00000039835 | Nhsl1 | L |
| ENSMUSG00000042331 | 1133,545488 | 0,696245133 | 0,197142927 | 3,531676966 | 0,000412933 | 0,019797148 | ENSMUSG00000042331 | Specc1 | A |
| ENSMUSG00000039145 | 514,3061008 | 0,692561552 | 0,220245656 | 3,144495851 | 0,001663733 | 0,046197904 | ENSMUSG00000039145 | Camk1d | B |
| ENSMUSG00000024066 | 1400,05722 | 0,689022426 | 0,215589215 | 3,195996733 | 0,001393487 | 0,041044436 | ENSMUSG00000024066 | Xdh | K |
| ENSMUSG00000039960 | 2809,842054 | 0,688190542 | 0,124042939 | 5,548002531 | 2,88952E-08 | 1,66238E-05 | ENSMUSG00000039960 | Rhou | A |
| ENSMUSG00000028469 | 705,0401645 | 0,686806371 | 0,213559117 | 3,216001177 | 0,001299903 | 0,039295918 | ENSMUSG00000028469 | Npr2 | L |
| ENSMUSG00000042129 | 5558,487444 | 0,686475836 | 0,174892204 | 3,925136845 | 8,66804E-05 | 0,007253577 | ENSMUSG00000042129 | Rassf4 | A |
| ENSMUSG00000023905 | 2627,192605 | 0,681121165 | 0,167481826 | 4,066836283 | 4,76557E-05 | 0,005072785 | ENSMUSG00000023905 | Tnfrsf12a | A |
| ENSMUSG00000104346 | 1730,512126 | 0,676379475 | 0,203795531 | 3,3189122 | 0,000903688 | 0,031994044 | ENSMUSG00000104346 | Pcdhga3 | S |
| ENSMUSG00000038508 | 1353,992334 | 0,675482979 | 0,208286864 | 3,243041666 | 0,001182609 | 0,03740865 | ENSMUSG00000038508 | Gdf15 | X |
| ENSMUSG00000021087 | 2149,483762 | 0,672498599 | 0,182896376 | 3,676937808 | 0,000236051 | 0,013928496 | ENSMUSG00000021087 | Rtn1 | L |
| ENSMUSG00000021876 | 35026,24833 | 0,670112603 | 0,183452477 | 3,652785799 | 0,000259411 | 0,014877724 | ENSMUSG00000021876 | Rnase4 | B |
| ENSMUSG00000020407 | 5245,917214 | 0,669408444 | 0,181062499 | 3,697112579 | 0,000218066 | 0,013076836 | ENSMUSG00000020407 | Upp1 | B |
| ENSMUSG00000048087 | 1140,468676 | 0,666589408 | 0,197347547 | 3,377743576 | 0,000730832 | 0,028566065 | ENSMUSG00000048087 | Gm4737 | S |
| ENSMUSG00000022965 | 2645,598957 | 0,665460442 | 0,193080609 | 3,446542069 | 0,00056781 | 0,024366865 | ENSMUSG00000022965 | Ifngr2 | A |
| ENSMUSG00000018417 | 3078,222163 | 0,664284393 | 0,181979405 | 3,650327309 | 0,000261906 | 0,014974211 | ENSMUSG00000018417 | Myo1b | L |
| ENSMUSG00000009585 | 2945,936756 | 0,659122478 | 0,179880295 | 3,664228366 | 0,000248085 | 0,014394851 | ENSMUSG00000009585 | Apobec3 | B |
| ENSMUSG00000069805 | 2053,285029 | 0,649190827 | 0,185351112 | 3,502492222 | 0,000460927 | 0,021161273 | ENSMUSG00000069805 | Fbp1 | L |
| ENSMUSG00000047648 | 652,4183483 | 0,64368009 | 0,164481427 | 3,913390707 | 9,10091E-05 | 0,007363407 | ENSMUSG00000047648 | Fbxo30 | A |
| ENSMUSG00000042524 | 1094,119682 | 0,64182055 | 0,171280046 | 3,747199772 | 0,00017882 | 0,011197517 | ENSMUSG00000042524 | Sun2 | A |

| | | | | | | | | | |
|---|---|---|---|---|---|---|---|---|---|
| ENSMUSG00000090035 | 2214,641251 | 0,638548191 | 0,143509102 | 4,449530955 | 8,6058E-06 | 0,001427323 | ENSMUSG00000090035 | Galnt4 | B |
| ENSMUSG00000026425 | 900,4422523 | 0,638144521 | 0,202934827 | 3,14457863 | 0,001663262 | 0,046197904 | ENSMUSG00000026425 | Srgap2 | B |
| ENSMUSG00000040525 | 2765,403669 | 0,629244881 | 0,191402659 | 3,287545133 | 0,00101065 | 0,034516855 | ENSMUSG00000040525 | Cblc | B |
| ENSMUSG00000021025 | 9684,254012 | 0,622779741 | 0,100167779 | 6,21736598 | 5,0557E-10 | 5,81721E-07 | ENSMUSG00000021025 | Nfkbia | A |
| ENSMUSG00000021895 | 1695,0604 | 0,614760443 | 0,141636793 | 4,340400749 | 1,42223E-05 | 0,002128721 | ENSMUSG00000021895 | Arhgef3 | L |
| ENSMUSG00000028413 | 2250,364335 | 0,604623279 | 0,159033604 | 3,801858629 | 0,000143615 | 0,00972039 | ENSMUSG00000028413 | B4galt1 | A |
| ENSMUSG00000027994 | 1189,944027 | 0,603123131 | 0,15752557 | 3,82873161 | 0,000128805 | 0,009107438 | ENSMUSG00000027994 | Mcub | |
| ENSMUSG00000040659 | 3776,159532 | 0,600525019 | 0,150053092 | 4,002083614 | 6,27871E-05 | 0,005927746 | ENSMUSG00000040659 | Efhd2 | A |
| ENSMUSG00000024048 | 5727,080849 | 0,598705452 | 0,147975896 | 4,04596606 | 5,21078E-05 | 0,005283497 | ENSMUSG00000024048 | Myl12a | A |
| ENSMUSG00000030042 | 717,1635305 | 0,592103556 | 0,167562143 | 3,533635603 | 0,000409886 | 0,019702342 | ENSMUSG00000030042 | Pole4 | A |
| ENSMUSG00000036908 | 4884,226537 | 0,587448958 | 0,142007019 | 4,136760013 | 3,52244E-05 | 0,004104313 | ENSMUSG00000036908 | Unc93b1 | D |
| ENSMUSG00000021823 | 4760,383473 | 0,585343282 | 0,168810736 | 3,467452945 | 0,000525416 | 0,023308201 | ENSMUSG00000021823 | Vcl | A |
| ENSMUSG00000021996 | 11455,73734 | 0,58226953 | 0,152506089 | 3,818008399 | 0,000134533 | 0,009293801 | ENSMUSG00000021996 | Esd | F |
| ENSMUSG00000034917 | 2521,475146 | 0,579266385 | 0,160485742 | 3,609457006 | 0,000306839 | 0,016199394 | ENSMUSG00000034917 | Tjp3 | C |
| ENSMUSG00000023036 | 1424,739139 | 0,576445068 | 0,1811266 | 3,182553349 | 0,001459826 | 0,042323465 | ENSMUSG00000023036 | Pcdhgc4 | #N/A |
| ENSMUSG00000041135 | 2900,218972 | 0,573995744 | 0,140491135 | 4,085636759 | 4,39561E-05 | 0,004788354 | ENSMUSG00000041135 | Ripk2 | A |
| ENSMUSG00000024052 | 1959,961636 | 0,563175137 | 0,124163407 | 4,535757765 | 5,7397E-06 | 0,001067353 | ENSMUSG00000024052 | Lpin2 | B |
| ENSMUSG00000024661 | 72052,40933 | 0,557881099 | 0,174072175 | 3,204883836 | 0,00135117 | 0,040250873 | ENSMUSG00000024661 | Fth1 | A |
| ENSMUSG00000052684 | 5574,423264 | 0,556329774 | 0,141402357 | 3,934374118 | 8,34138E-05 | 0,007076717 | ENSMUSG00000052684 | Jun | A |
| ENSMUSG00000034422 | 6466,152334 | 0,550408307 | 0,164043964 | 3,355248763 | 0,000792936 | 0,029913853 | ENSMUSG00000034422 | Parp14 | B |
| ENSMUSG00000019947 | 2055,182047 | 0,539384864 | 0,137568954 | 3,920832783 | 8,82435E-05 | 0,007306461 | ENSMUSG00000019947 | Arid5b | L |
| ENSMUSG00000023030 | 7772,112677 | 0,537390533 | 0,113010018 | 4,755246879 | 1,98204E-06 | 0,000486525 | ENSMUSG00000023030 | Slc11a2 | A |
| ENSMUSG00000032786 | 2948,649781 | 0,537228519 | 0,156936643 | 3,423219131 | 0,000618842 | 0,025659629 | ENSMUSG00000032786 | Alas1 | A |
| ENSMUSG00000022500 | 4316,464458 | 0,532184753 | 0,155117758 | 3,430843504 | 0,000601708 | 0,025406966 | ENSMUSG00000022500 | Litaf | A |
| ENSMUSG00000040128 | 8074,875922 | 0,530226067 | 0,137512557 | 3,855837456 | 0,000115334 | 0,008392494 | ENSMUSG00000040128 | Pnrc1 | A |
| ENSMUSG00000006818 | 21722,58205 | 0,527614909 | 0,150991484 | 3,494335542 | 0,000475243 | 0,021496878 | ENSMUSG00000006818 | Sod2 | A |
| ENSMUSG00000018340 | 19504,63145 | 0,52511555 | 0,107628835 | 4,878948569 | 1,06653E-06 | 0,000297497 | ENSMUSG00000018340 | Anxa6 | A |
| ENSMUSG00000051043 | 2900,190086 | 0,521727379 | 0,167167964 | 3,120977042 | 0,001802521 | 0,048515218 | ENSMUSG00000051043 | Gprc5c | B |
| ENSMUSG00000044864 | 3048,367388 | 0,520882681 | 0,161468501 | 3,225908943 | 0,001255733 | 0,038866653 | ENSMUSG00000044864 | Ankrd50 | A |
| ENSMUSG00000060803 | 25611,63141 | 0,519355829 | 0,152005792 | 3,416684468 | 0,000633887 | 0,026048795 | ENSMUSG00000060803 | Gstp1 | A |

| | | | | | | | | | |
|---|---|---|---|---|---|---|---|---|---|
| ENSMUSG00000029455 | 2453,996351 | 0,512356358 | 0,151869251 | 3,373667498 | 0,000741739 | 0,028687848 | ENSMUSG00000029455 | Aldh2 | B |
| ENSMUSG00000069662 | 4918,73417 | 0,510735351 | 0,147854958 | 3,454299788 | 0,000551724 | 0,024106381 | ENSMUSG00000069662 | Marcks | A |
| ENSMUSG00000002983 | 2105,624371 | 0,50898455 | 0,145004716 | 3,510124104 | 0,000447898 | 0,02087543 | ENSMUSG00000002983 | Relb | A |
| ENSMUSG00000050912 | 4070,006174 | 0,50538249 | 0,126470033 | 3,996065137 | 6,4404E-05 | 0,005988273 | ENSMUSG00000050912 | Tmem123 | A |
| ENSMUSG00000016382 | 3883,820564 | 0,503241149 | 0,153572377 | 3,276898869 | 0,00104954 | 0,035130951 | ENSMUSG00000016382 | Pls3 | |
| ENSMUSG00000030203 | 3197,737328 | 0,502578095 | 0,160542684 | 3,130495148 | 0,001745119 | 0,0475965 | ENSMUSG00000030203 | Dusp16 | A |
| ENSMUSG00000035356 | 8865,468683 | 0,499378575 | 0,115368782 | 4,328541624 | 1,501E-05 | 0,002214435 | ENSMUSG00000035356 | Nfkbiz | A |
| ENSMUSG00000038235 | 10131,95788 | 0,498855077 | 0,145615718 | 3,425832629 | 0,000612918 | 0,025645044 | ENSMUSG00000038235 | F11r | A |
| ENSMUSG00000050708 | 136184,542 | 0,498418957 | 0,130554966 | 3,817694345 | 0,000134705 | 0,009293801 | ENSMUSG00000050708 | Ftl1 | A |
| ENSMUSG00000036620 | 2703,732063 | 0,495106858 | 0,148730432 | 3,328887378 | 0,000871937 | 0,031526499 | ENSMUSG00000036620 | Mgat4b | B |
| ENSMUSG00000033545 | 2516,805536 | 0,492743863 | 0,133458076 | 3,692124723 | 0,000222388 | 0,013292758 | ENSMUSG00000033545 | Znrf1 | B |
| ENSMUSG00000005087 | 9483,344542 | 0,486333346 | 0,151534435 | 3,209391612 | 0,001330162 | 0,039945611 | ENSMUSG00000005087 | Cd44 | L |
| ENSMUSG00000034573 | 2420,931349 | 0,480274155 | 0,137391083 | 3,495671963 | 0,00047287 | 0,021442193 | ENSMUSG00000034573 | Ptpn13 | B |
| ENSMUSG00000026479 | 5554,417345 | 0,477770834 | 0,117510232 | 4,065780698 | 4,78719E-05 | 0,005072785 | ENSMUSG00000026479 | Lamc2 | B |
| ENSMUSG00000039943 | 3178,787824 | 0,471311184 | 0,135647021 | 3,474541349 | 0,000511727 | 0,022755805 | ENSMUSG00000039943 | Plcb4 | I |
| ENSMUSG00000028961 | 9811,403023 | 0,464718297 | 0,141352318 | 3,287659537 | 0,001010239 | 0,034516855 | ENSMUSG00000028961 | Pgd | A |
| ENSMUSG00000068876 | 13346,75979 | 0,461662007 | 0,103571367 | 4,457428932 | 8,29485E-06 | 0,001388257 | ENSMUSG00000068876 | Cgn | A |
| ENSMUSG00000057406 | 3192,646452 | 0,461545374 | 0,119177923 | 3,872742216 | 0,000107618 | 0,008124192 | ENSMUSG00000057406 | Nsd2 | |
| ENSMUSG00000059456 | 2885,232509 | 0,454850739 | 0,143236847 | 3,175514888 | 0,001495708 | 0,042957867 | ENSMUSG00000059456 | Ptk2b | H |
| ENSMUSG00000023959 | 1442,839845 | 0,454837706 | 0,134491948 | 3,38189544 | 0,000719875 | 0,028347727 | ENSMUSG00000023959 | Clic5 | L |
| ENSMUSG00000032852 | 13558,39992 | 0,449523034 | 0,118071698 | 3,807203938 | 0,000140547 | 0,009618838 | ENSMUSG00000032852 | Rspo4 | B |
| ENSMUSG00000031897 | 4669,005952 | 0,446398157 | 0,112723617 | 3,96011208 | 7,49146E-05 | 0,006639341 | ENSMUSG00000031897 | Psmb10 | L |
| ENSMUSG00000017009 | 7289,876496 | 0,444407571 | 0,133136357 | 3,337988079 | 0,000843874 | 0,031133697 | ENSMUSG00000017009 | Sdc4 | A |
| ENSMUSG00000075254 | 11345,26872 | 0,437272009 | 0,110230198 | 3,966898509 | 7,2814E-05 | 0,006539051 | ENSMUSG00000075254 | Heg1 | A |
| ENSMUSG00000023224 | 25746,99976 | 0,433166655 | 0,112941244 | 3,835327465 | 0,000125397 | 0,008913362 | ENSMUSG00000023224 | Serping1 | D |
| ENSMUSG00000027366 | 7505,51624 | 0,432691792 | 0,108766549 | 3,978169714 | 6,94478E-05 | 0,006329376 | ENSMUSG00000027366 | Sppl2a | A |
| ENSMUSG00000009394 | 6071,168651 | 0,430132972 | 0,105825156 | 4,064562596 | 4,81226E-05 | 0,005072785 | ENSMUSG00000009394 | Syn2 | A |
| ENSMUSG00000071637 | 11719,96773 | 0,429391222 | 0,131633504 | 3,262020762 | 0,00110621 | 0,036022203 | ENSMUSG00000071637 | Cebpd | |
| ENSMUSG00000025068 | 2149,586888 | 0,418134567 | 0,116689806 | 3,583299861 | 0,000339281 | 0,017254581 | ENSMUSG00000025068 | Gsto1 | B |
| ENSMUSG00000042688 | 8847,900184 | 0,40989152 | 0,112390785 | 3,647020706 | 0,000265299 | 0,015038657 | ENSMUSG00000042688 | Mapk6 | B |

| | | | | | | | | | |
|---|---|---|---|---|---|---|---|---|---|
| ENSMUSG00000028788 | 9191,583247 | 0,407815364 | 0,103940129 | 3,923560315 | 8,72499E-05 | 0,007268194 | ENSMUSG00000028788 | Ptp4a2 | A |
| ENSMUSG00000035441 | 6016,627508 | 0,402964254 | 0,119325081 | 3,377028957 | 0,000732733 | 0,028579706 | ENSMUSG00000035441 | Myo1d | A |
| ENSMUSG00000057963 | 1924,320203 | 0,402890198 | 0,117689594 | 3,423328975 | 0,000618592 | 0,025659629 | ENSMUSG00000057963 | Itpk1 | A |
| ENSMUSG00000028980 | 6194,969729 | 0,401362863 | 0,111231276 | 3,608363388 | 0,000308135 | 0,016199394 | ENSMUSG00000028980 | H6pd | D |
| ENSMUSG00000027775 | 4219,987278 | 0,391098861 | 0,097700739 | 4,003028664 | 6,25367E-05 | 0,005927746 | ENSMUSG00000027775 | Mfsd1 | A |
| ENSMUSG00000001552 | 10633,8479 | 0,380787269 | 0,110233381 | 3,454373503 | 0,000551573 | 0,024106381 | ENSMUSG00000001552 | Jup | A |
| ENSMUSG00000022636 | 18960,92562 | 0,379240009 | 0,120050211 | 3,159011585 | 0,001583052 | 0,044630914 | ENSMUSG00000022636 | Alcam | B |
| ENSMUSG00000029723 | 3973,966824 | 0,369530534 | 0,111230208 | 3,322213822 | 0,000893062 | 0,031739916 | ENSMUSG00000029723 | Tsc22d4 | B |
| ENSMUSG00000008859 | 4581,832124 | 0,361316008 | 0,09979459 | 3,620597156 | 0,000293924 | 0,015915112 | ENSMUSG00000008859 | Rala | A |
| ENSMUSG00000034902 | 5935,918883 | 0,351538727 | 0,108129575 | 3,251087657 | 0,001149644 | 0,0367447 | ENSMUSG00000034902 | Pip5k1c | A |
| ENSMUSG00000022863 | 3250,867034 | 0,344318693 | 0,107299756 | 3,208941999 | 0,001332244 | 0,039945611 | ENSMUSG00000022863 | Btg3 | A |
| ENSMUSG00000056608 | 18013,98981 | 0,338337735 | 0,092899378 | 3,641980624 | 0,000270548 | 0,015087661 | ENSMUSG00000056608 | Chd9 | B |
| ENSMUSG00000021756 | 18257,68801 | 0,328541568 | 0,083963608 | 3,912904379 | 9,11927E-05 | 0,007363407 | ENSMUSG00000021756 | Il6st | A |
| ENSMUSG00000020372 | 32409,3549 | 0,300049722 | 0,096478536 | 3,110015302 | 0,001870777 | 0,049770225 | ENSMUSG00000020372 | Rack1 | |
| ENSMUSG00000038366 | 20468,77625 | 0,276526245 | 0,081408186 | 3,396786731 | 0,000681821 | 0,027287656 | ENSMUSG00000038366 | Lasp1 | A |
| ENSMUSG00000091655 | 24,15087796 | -7,935816534 | 1,617315601 | -4,9067829 | 9,25824E-07 | 0,000266319 | ENSMUSG00000091655 | Gm8741 | |
| ENSMUSG00000092541 | 16,26353522 | -7,364732978 | 2,258982953 | -3,260198563 | 0,001113342 | 0,036022203 | ENSMUSG00000092541 | Gm20537 | |
| ENSMUSG00000076128 | 13,81261739 | -7,129675485 | 1,955873807 | -3,645263544 | 0,000267118 | 0,015038657 | ENSMUSG00000076128 | Mir686 | |
| ENSMUSG00000104729 | 10,53475218 | -6,736162711 | 1,923008757 | -3,502928775 | 0,000460172 | 0,021161273 | ENSMUSG00000104729 | Gm44471 | |
| ENSMUSG00000049235 | 9,514200853 | -6,593144111 | 1,951104863 | -3,379184911 | 0,000727011 | 0,028537891 | ENSMUSG00000049235 | Gm7324 | |
| ENSMUSG00000031384 | 8,415538404 | -6,415790897 | 1,938946039 | -3,308906368 | 0,000936612 | 0,032781404 | ENSMUSG00000031384 | Asb9 | |
| ENSMUSG00000078497 | 11,50424377 | -5,416082965 | 1,721254994 | -3,146589544 | 0,001651867 | 0,04607707 | ENSMUSG00000078497 | Zfp978 | |
| ENSMUSG00000049414 | 25,98147166 | -5,109286589 | 1,329668605 | -3,842526302 | 0,000121774 | 0,008723214 | ENSMUSG00000049414 | Gm5417 | |
| ENSMUSG00000030737 | 20,82599611 | -4,448522289 | 1,28798069 | -3,453873436 | 0,000552597 | 0,024106381 | ENSMUSG00000030737 | Slco2b1 | N |
| ENSMUSG00000019464 | 179,1210842 | -3,839280685 | 0,763820427 | -5,026417927 | 4,99726E-07 | 0,00015862 | ENSMUSG00000019464 | Ptger1 | D |
| ENSMUSG00000092386 | 87,26207655 | -3,386509246 | 1,010602615 | -3,350980096 | 0,000805261 | 0,030225842 | ENSMUSG00000092386 | Gm20536 | |
| ENSMUSG00000020014 | 132,9775046 | -1,390769227 | 0,427027497 | -3,256861064 | 0,001126516 | 0,036320756 | ENSMUSG00000020014 | Cfap54 | L |
| ENSMUSG00000032221 | 1629,03335 | -1,327811291 | 0,286249111 | -4,638656468 | 3,50681E-06 | 0,000750599 | ENSMUSG00000032221 | Mns1 | B |
| ENSMUSG00000031837 | 507,9333177 | -1,266941701 | 0,315775589 | -4,012158454 | 6,01661E-05 | 0,005769051 | ENSMUSG00000031837 | Necab2 | B |
| ENSMUSG00000022490 | 4342,834035 | -1,092163349 | 0,250373861 | -4,36213007 | 1,28802E-05 | 0,001972375 | ENSMUSG00000022490 | Ppp1r1a | A |

| | | | | | | | | | |
|---|---|---|---|---|---|---|---|---|---|
| ENSMUSG00000037016 | 368,3314038 | -1,068646014 | 0,259487957 | -4,118287507 | 3,81698E-05 | 0,004364637 | ENSMUSG00000037016 | Frem2 | L |
| ENSMUSG00000098132 | 475,191995 | -1,043554533 | 0,317730227 | -3,284404326 | 0,001021982 | 0,034713444 | ENSMUSG00000098132 | Rassf10 | A |
| ENSMUSG00000020072 | 406,0246461 | -1,021439187 | 0,300070136 | -3,404001483 | 0,000664064 | 0,026928231 | ENSMUSG00000020072 | Pbld2 | A |
| ENSMUSG00000040543 | 2370,569337 | -1,006707836 | 0,221424859 | -4,546498707 | 5,45457E-06 | 0,001046028 | ENSMUSG00000040543 | Pitpnm3 | A |
| ENSMUSG00000042942 | 273,1204495 | -0,996588703 | 0,245655862 | -4,056848858 | 4,97392E-05 | 0,005116325 | ENSMUSG00000042942 | Greb1l | L |
| ENSMUSG00000005268 | 31472,86791 | -0,983320371 | 0,209405939 | -4,695761611 | 2,65615E-06 | 0,0006141 | ENSMUSG00000005268 | Prlr | B |
| ENSMUSG00000028926 | 2810,638068 | -0,970737149 | 0,188400603 | -5,152516148 | 2,57014E-07 | 9,64926E-05 | ENSMUSG00000028926 | Cdk14 | B |
| ENSMUSG00000105263 | 8287,220339 | -0,969909547 | 0,23541155 | -4,12005931 | 3,78775E-05 | 0,004362797 | ENSMUSG00000105263 | Gm42427 | |
| ENSMUSG00000039628 | 734,6382013 | -0,960140017 | 0,308360032 | -3,113698003 | 0,001847585 | 0,04943901 | ENSMUSG00000039628 | Hs3st6 | B |
| ENSMUSG00000073791 | 184,4739954 | -0,944319948 | 0,29974172 | -3,150445485 | 0,001630217 | 0,045680808 | ENSMUSG00000073791 | Efcab7 | E |
| ENSMUSG00000030785 | 1631,938161 | -0,937222907 | 0,291073052 | -3,219888963 | 0,001282403 | 0,039023193 | ENSMUSG00000030785 | Cox6a2 | H |
| ENSMUSG00000042797 | 652,9864155 | -0,936042471 | 0,220277644 | -4,249375712 | 2,14367E-05 | 0,002844135 | ENSMUSG00000042797 | Aqp11 | B |
| ENSMUSG00000051435 | 681,6915899 | -0,92676326 | 0,283150375 | -3,273042676 | 0,001063964 | 0,035292936 | ENSMUSG00000051435 | Fhad1 | A |
| ENSMUSG00000037843 | 5970,418924 | -0,916070892 | 0,253188746 | -3,618134326 | 0,000296734 | 0,015945005 | ENSMUSG00000037843 | Vstm2l | A |
| ENSMUSG00000059824 | 746,7541073 | -0,898189384 | 0,282568809 | -3,178657212 | 0,00147959 | 0,042627925 | ENSMUSG00000059824 | Dbp | A |
| ENSMUSG00000036800 | 772,4370102 | -0,894664734 | 0,277572813 | -3,223171334 | 0,001267797 | 0,039023193 | ENSMUSG00000036800 | Fam135b | L |
| ENSMUSG00000016179 | 272,8207984 | -0,875807177 | 0,237948306 | -3,68066153 | 0,00023263 | 0,013770777 | ENSMUSG00000016179 | Camk1g | L |
| ENSMUSG00000057069 | 81456,37061 | -0,874627724 | 0,219971593 | -3,976093968 | 7,00565E-05 | 0,006353396 | ENSMUSG00000057069 | Ero1lb | A |
| ENSMUSG00000027487 | 857,7664583 | -0,870466072 | 0,248711687 | -3,499900151 | 0,000465432 | 0,021262064 | ENSMUSG00000027487 | Cdk5rap1 | A |
| ENSMUSG00000035299 | 1310,349301 | -0,867340958 | 0,275350554 | -3,149951742 | 0,001632974 | 0,045685073 | ENSMUSG00000035299 | Mid1 | |
| ENSMUSG00000028825 | 561,6371392 | -0,86730947 | 0,260724505 | -3,326536071 | 0,000879326 | 0,031526499 | ENSMUSG00000028825 | Rhd | Q |
| ENSMUSG00000005373 | 1925,580749 | -0,865794203 | 0,26384639 | -3,281432819 | 0,001032811 | 0,034848298 | ENSMUSG00000005373 | Mlxipl | A |
| ENSMUSG00000021287 | 440,807743 | -0,863383062 | 0,241928775 | -3,568748956 | 0,00035869 | 0,018042299 | ENSMUSG00000021287 | Xrcc3 | B |
| ENSMUSG00000038058 | 1623,420677 | -0,862463612 | 0,205647953 | -4,193883775 | 2,74219E-05 | 0,003457783 | ENSMUSG00000038058 | Nod1 | D |
| ENSMUSG00000021071 | 1255,944353 | -0,849562613 | 0,210562082 | -4,034736959 | 5,46635E-05 | 0,005410511 | ENSMUSG00000021071 | Trim9 | B |
| ENSMUSG00000022235 | 845,4250531 | -0,848535698 | 0,260397512 | -3,258616764 | 0,001119568 | 0,03616008 | ENSMUSG00000022235 | Cmbl | L |
| ENSMUSG00000024553 | 1343,051801 | -0,815976106 | 0,255151585 | -3,198005245 | 0,001383818 | 0,040997983 | ENSMUSG00000024553 | Galr1 | B |
| ENSMUSG00000027984 | 18235,73686 | -0,811756169 | 0,225897828 | -3,593466019 | 0,000326308 | 0,016733522 | ENSMUSG00000027984 | Hadh | A |
| ENSMUSG00000097530 | 681,5678173 | -0,808122058 | 0,259489226 | -3,11427981 | 0,001843945 | 0,04941344 | ENSMUSG00000097530 | Kansl2-ps | |
| ENSMUSG00000024027 | 7143,335379 | -0,806171402 | 0,205649409 | -3,920125063 | 8,8503E-05 | 0,007306461 | ENSMUSG00000024027 | Glp1r | A |

| | | | | | | | | | |
|---|---|---|---|---|---|---|---|---|---|
| ENSMUSG00000031026 | 536,8382021 | -0,802702344 | 0,250508756 | -3,20428858 | 0,001353967 | 0,040269032 | ENSMUSG00000031026 | Trim66 | L |
| ENSMUSG00000034218 | 3095,615055 | -0,798771384 | 0,137947732 | -5,790391566 | 7,02225E-09 | 5,38665E-06 | ENSMUSG00000034218 | Atm | B |
| ENSMUSG00000024972 | 547,5412522 | -0,796767195 | 0,224253118 | -3,552981573 | 0,000380891 | 0,018653657 | ENSMUSG00000024972 | Lgals12 | L |
| ENSMUSG00000074207 | 4071,950756 | -0,794488671 | 0,211956101 | -3,748364254 | 0,000177992 | 0,011183706 | ENSMUSG00000074207 | Adh1 | A |
| ENSMUSG00000044139 | 57317,62381 | -0,787997278 | 0,204238214 | -3,858226452 | 0,000114213 | 0,00834388 | ENSMUSG00000044139 | Prss53 | A |
| ENSMUSG00000058248 | 4901,878824 | -0,77826955 | 0,166688709 | -4,668999818 | 3,0267E-06 | 0,00068792 | ENSMUSG00000058248 | Kcnh1 | L |
| ENSMUSG00000009246 | 1952,47086 | -0,773548737 | 0,239290734 | -3,232673181 | 0,001226378 | 0,038235681 | ENSMUSG00000009246 | Trpm5 | K |
| ENSMUSG00000030761 | 1294,919401 | -0,771882992 | 0,167689884 | -4,603038489 | 4,16371E-06 | 0,000833195 | ENSMUSG00000030761 | Myo7a | N |
| ENSMUSG00000036402 | 53072,5212 | -0,756331509 | 0,180570997 | -4,18855476 | 2,80737E-05 | 0,003515891 | ENSMUSG00000036402 | Gng12 | B |
| ENSMUSG00000037279 | 642,9904923 | -0,753935098 | 0,234477926 | -3,215377719 | 0,00130273 | 0,039316814 | ENSMUSG00000037279 | Ovol2 | L |
| ENSMUSG00000031255 | 12802,42819 | -0,75039097 | 0,172079571 | -4,360720836 | 1,29635E-05 | 0,001972375 | ENSMUSG00000031255 | Sytl4 | |
| ENSMUSG00000026185 | 7384,795711 | -0,746561707 | 0,14432811 | -5,172670148 | 2,30772E-07 | 8,94569E-05 | ENSMUSG00000026185 | Igfbp5 | L |
| ENSMUSG00000022702 | 2316,854056 | -0,743837372 | 0,15804947 | -4,706357894 | 2,52182E-06 | 0,000595213 | ENSMUSG00000022702 | Hira | A |
| ENSMUSG00000027314 | 861,6576648 | -0,743230849 | 0,169995629 | -4,372058584 | 1,2308E-05 | 0,001936677 | ENSMUSG00000027314 | Dll4 | L |
| ENSMUSG00000044645 | 2897,481789 | -0,73830317 | 0,227160141 | -3,250144004 | 0,001153466 | 0,036802957 | ENSMUSG00000044645 | Gm7334 | L |
| ENSMUSG00000056158 | 3569,286629 | -0,735376281 | 0,214929045 | -3,421483974 | 0,000622804 | 0,025765889 | ENSMUSG00000056158 | Car10 | A |
| ENSMUSG00000097908 | 688,7545295 | -0,729992781 | 0,224841828 | -3,246694742 | 0,001167535 | 0,036995399 | ENSMUSG00000097908 | 4933404O12Rik | B |
| ENSMUSG00000013089 | 3296,154732 | -0,726299039 | 0,123707852 | -5,871082775 | 4,32958E-09 | 3,7956E-06 | ENSMUSG00000013089 | Etv5 | I |
| ENSMUSG00000070880 | 3095,521111 | -0,721828946 | 0,207009326 | -3,486939264 | 0,000488582 | 0,021924864 | ENSMUSG00000070880 | Gad1 | A |
| ENSMUSG00000000215 | 9157806,966 | -0,709676093 | 0,193592675 | -3,665821001 | 0,000246546 | 0,014394851 | ENSMUSG00000000215 | Ins2 | A |
| ENSMUSG00000055409 | 11017,1338 | -0,708512055 | 0,153140531 | -4,626548259 | 3,7181E-06 | 0,000777844 | ENSMUSG00000055409 | Nell1 | B |
| ENSMUSG00000106133 | 434,734907 | -0,707928548 | 0,227574645 | -3,110753173 | 0,001866109 | 0,049770225 | ENSMUSG00000106133 | Gm3724 | |
| ENSMUSG00000049971 | 1242,083927 | -0,698021657 | 0,202246609 | -3,451339237 | 0,000557812 | 0,024106381 | ENSMUSG00000049971 | Glt1d1 | A |
| ENSMUSG00000079620 | 7582,19844 | -0,689708173 | 0,206760515 | -3,335782815 | 0,000850596 | 0,031172947 | ENSMUSG00000079620 | Muc4 | D |
| ENSMUSG00000022479 | 11056,3572 | -0,679308285 | 0,201076816 | -3,378352105 | 0,000729216 | 0,028563564 | ENSMUSG00000022479 | Vdr | A |
| ENSMUSG00000055254 | 19604,4819 | -0,67696089 | 0,194281334 | -3,484436088 | 0,000493175 | 0,022037272 | ENSMUSG00000055254 | Ntrk2 | L |
| ENSMUSG00000041842 | 4606,442382 | -0,675794096 | 0,209868741 | -3,220079817 | 0,001281549 | 0,039023193 | ENSMUSG00000041842 | Fhdc1 | A |
| ENSMUSG00000035804 | 3975391,361 | -0,663569766 | 0,207543073 | -3,197262888 | 0,001387384 | 0,040997983 | ENSMUSG00000035804 | Ins1 | C |
| ENSMUSG00000020169 | 1155,376083 | -0,662953385 | 0,197856329 | -3,350680714 | 0,000806132 | 0,030225842 | ENSMUSG00000020169 | Best3 | L |
| ENSMUSG00000041895 | 12913,85929 | -0,662362588 | 0,195619236 | -3,385978811 | 0,000709249 | 0,028140661 | ENSMUSG00000041895 | Wipi1 | A |

| | | | | | | | | | |
|---|---|---|---|---|---|---|---|---|---|
| ENSMUSG00000042638 | 3859,924277 | -0,657821798 | 0,186601606 | -3,525274033 | 0,000423045 | 0,020072827 | ENSMUSG00000042638 | Gucy2c | L |
| ENSMUSG00000070348 | 13490,28492 | -0,65616414 | 0,138218331 | -4,747301879 | 2,06148E-06 | 0,000499367 | ENSMUSG00000070348 | Ccnd1 | L |
| ENSMUSG00000025140 | 5783,033348 | -0,646346808 | 0,169441197 | -3,814578872 | 0,000136415 | 0,009370921 | ENSMUSG00000025140 | Pycr1 | A |
| ENSMUSG00000026443 | 786,2616553 | -0,641878012 | 0,196225103 | -3,271130978 | 0,001071183 | 0,035341356 | ENSMUSG00000026443 | Lrrn2 | L |
| ENSMUSG00000037106 | 2648,201143 | -0,641554878 | 0,188987614 | -3,394692713 | 0,000687057 | 0,027437556 | ENSMUSG00000037106 | Fer1l6 | O |
| ENSMUSG00000046593 | 15666,92085 | -0,638839765 | 0,165132665 | -3,868645647 | 0,000109442 | 0,008190321 | ENSMUSG00000046593 | Tmem215 | A |
| ENSMUSG00000026189 | 1603,542421 | -0,637566488 | 0,192726991 | -3,308132835 | 0,000939202 | 0,032809711 | ENSMUSG00000026189 | Pecr | B |
| ENSMUSG00000096956 | 531,3460911 | -0,634889846 | 0,179653064 | -3,533977268 | 0,000409356 | 0,019702342 | ENSMUSG00000096956 | Snhg18 | B |
| ENSMUSG00000044453 | 9859,490984 | -0,626983538 | 0,159682738 | -3,926432801 | 8,6215E-05 | 0,007247568 | ENSMUSG00000044453 | Ffar1 | B |
| ENSMUSG00000019876 | 576,2191923 | -0,621166351 | 0,196697355 | -3,157980192 | 0,001588664 | 0,044720635 | ENSMUSG00000019876 | Pkib | X |
| ENSMUSG00000026930 | 4126,487245 | -0,613585248 | 0,169877124 | -3,611935688 | 0,00030392 | 0,016124398 | ENSMUSG00000026930 | Gpsm1 | C |
| ENSMUSG00000028445 | 5679,972936 | -0,612633369 | 0,173272057 | -3,535673198 | 0,000406738 | 0,019702342 | ENSMUSG00000028445 | Enho | C |
| ENSMUSG00000025036 | 3747,972807 | -0,609408247 | 0,187974849 | -3,241966938 | 0,001187078 | 0,0374856 | ENSMUSG00000025036 | Sfxn2 | A |
| ENSMUSG00000034584 | 3733,346337 | -0,60767448 | 0,194650446 | -3,121875609 | 0,001797028 | 0,04850923 | ENSMUSG00000034584 | Exph5 | B |
| ENSMUSG00000020614 | 2338,469872 | -0,605115059 | 0,154810725 | -3,908741204 | 9,27783E-05 | 0,007426297 | ENSMUSG00000020614 | Fam20a | A |
| ENSMUSG00000033653 | 3549,399595 | -0,602843317 | 0,155100356 | -3,886795179 | 0,000101576 | 0,007791756 | ENSMUSG00000033653 | Vps8 | I |
| ENSMUSG00000020262 | 2064,695148 | -0,596011129 | 0,143077235 | -4,165660097 | 3,10453E-05 | 0,003781061 | ENSMUSG00000020262 | Adarb1 | A |
| ENSMUSG00000022420 | 1563,013973 | -0,59202939 | 0,169629515 | -3,490131954 | 0,000482782 | 0,021771551 | ENSMUSG00000022420 | Dnal4 | A |
| ENSMUSG00000003355 | 3355,526468 | -0,591533977 | 0,156846342 | -3,771423481 | 0,000162319 | 0,010522153 | ENSMUSG00000003355 | Fkbp11 | B |
| ENSMUSG00000039137 | 2778,40206 | -0,591169974 | 0,18802782 | -3,144055876 | 0,001666236 | 0,046197904 | ENSMUSG00000039137 | Whrn | A |
| ENSMUSG00000029644 | 10843,76514 | -0,575789139 | 0,149384994 | -3,854397451 | 0,000116015 | 0,008408804 | ENSMUSG00000029644 | Pdx1 | A |
| ENSMUSG00000039062 | 9107,218462 | -0,575425027 | 0,167998668 | -3,425176137 | 0,000614401 | 0,025648802 | ENSMUSG00000039062 | Anpep | L |
| ENSMUSG00000019066 | 19129,84062 | -0,575167971 | 0,139715618 | -4,116704913 | 3,84328E-05 | 0,004367575 | ENSMUSG00000019066 | Rab3d | A |
| ENSMUSG00000027004 | 6949,341352 | -0,573320663 | 0,161124403 | -3,558248489 | 0,000373336 | 0,018525921 | ENSMUSG00000027004 | Frzb | B |
| ENSMUSG00000062944 | 1782,05135 | -0,56707281 | 0,168787994 | -3,359675037 | 0,000780342 | 0,029559869 | ENSMUSG00000062944 | 9130023H24Rik | A |
| ENSMUSG00000020471 | 1914,033066 | -0,565341715 | 0,12193826 | -4,636294751 | 3,5471E-06 | 0,000750599 | ENSMUSG00000020471 | Pold2 | A |
| ENSMUSG00000070392 | 1318,754284 | -0,565180718 | 0,133144295 | -4,244873699 | 2,18717E-05 | 0,002855725 | ENSMUSG00000070392 | Gm20634 | |
| ENSMUSG00000039601 | 2368,258798 | -0,563974337 | 0,175301536 | -3,21716712 | 0,001294632 | 0,039200935 | ENSMUSG00000039601 | Rcan2 | A |
| ENSMUSG00000031626 | 13680,41829 | -0,56268756 | 0,179890272 | -3,127948791 | 0,001760308 | 0,047939762 | ENSMUSG00000031626 | Sorbs2 | D |
| ENSMUSG00000061911 | 1767,467763 | -0,561127428 | 0,180419508 | -3,110126131 | 0,001870075 | 0,049770225 | ENSMUSG00000061911 | Myt1l | B |

| | | | | | | | | | |
|---|---|---|---|---|---|---|---|---|---|
| ENSMUSG00000000881 | 1820,569354 | -0,559727955 | 0,163104046 | -3,431723307 | 0,000599759 | 0,025382909 | ENSMUSG00000000881 | Dlg3 | |
| ENSMUSG00000036599 | 2230,513745 | -0,558007361 | 0,136469956 | -4,088866 | 4,33487E-05 | 0,004778742 | ENSMUSG00000036599 | Chst12 | A |
| ENSMUSG00000034171 | 2105,461458 | -0,554514383 | 0,174053447 | -3,185885677 | 0,001443116 | 0,041904996 | ENSMUSG00000034171 | Faah | B |
| ENSMUSG00000030515 | 1292,840573 | -0,554350744 | 0,152011143 | -3,646777034 | 0,00026555 | 0,015038657 | ENSMUSG00000030515 | Tarsl2 | A |
| ENSMUSG00000032965 | 6160,397492 | -0,553471658 | 0,118649076 | -4,664778486 | 3,0895E-06 | 0,00069363 | ENSMUSG00000032965 | Ift57 | A |
| ENSMUSG00000026163 | 29676,50528 | -0,553351824 | 0,161344203 | -3,429635604 | 0,000604392 | 0,025461929 | ENSMUSG00000026163 | Sphkap | A |
| ENSMUSG00000022661 | 3376,254265 | -0,550400751 | 0,139997739 | -3,931497418 | 8,44184E-05 | 0,007129095 | ENSMUSG00000022661 | Cd200 | A |
| ENSMUSG00000028601 | 1216,579347 | -0,546880254 | 0,140267252 | -3,898844859 | 9,66527E-05 | 0,007571812 | ENSMUSG00000028601 | Echdc2 | B |
| ENSMUSG00000023393 | 3151,940646 | -0,546022419 | 0,154875215 | -3,525563589 | 0,000422583 | 0,020072827 | ENSMUSG00000023393 | Slc17a9 | A |
| ENSMUSG00000052271 | 8610,79896 | -0,542675467 | 0,145422073 | -3,731726939 | 0,000190172 | 0,011788077 | ENSMUSG00000052271 | Bhlha15 | A |
| ENSMUSG00000040943 | 1088,887326 | -0,538308858 | 0,14344798 | -3,75264161 | 0,000174981 | 0,01107885 | ENSMUSG00000040943 | Tet2 | A |
| ENSMUSG00000035187 | 21218,44073 | -0,537898852 | 0,159394646 | -3,374635628 | 0,000739135 | 0,02864731 | ENSMUSG00000035187 | Nkx6-1 | A |
| ENSMUSG00000061601 | 13759,86544 | -0,536110226 | 0,165772451 | -3,234012785 | 0,00122064 | 0,038229998 | ENSMUSG00000061601 | Pclo | A |
| ENSMUSG00000041556 | 1649,120403 | -0,528231951 | 0,149835208 | -3,525419415 | 0,000422813 | 0,020072827 | ENSMUSG00000041556 | Fbxo2 | B |
| ENSMUSG00000058070 | 2600,742924 | -0,525434557 | 0,139069164 | -3,778224735 | 0,00015795 | 0,010385233 | ENSMUSG00000058070 | Eml1 | O |
| ENSMUSG00000047747 | 2388,09083 | -0,522312087 | 0,141611475 | -3,688345779 | 0,000225717 | 0,013448045 | ENSMUSG00000047747 | Rnf150 | L |
| ENSMUSG00000031840 | 6713,324306 | -0,51584279 | 0,162699139 | -3,170531781 | 0,001521602 | 0,043363298 | ENSMUSG00000031840 | Rab3a | A |
| ENSMUSG00000026544 | 1023,179588 | -0,509706121 | 0,154454861 | -3,300032892 | 0,000966735 | 0,033341405 | ENSMUSG00000026544 | Dusp23 | A |
| ENSMUSG00000109946 | 947,0185955 | -0,509590275 | 0,162044458 | -3,14475597 | 0,001662254 | 0,046197904 | ENSMUSG00000109946 | Brd3os | |
| ENSMUSG00000044254 | 1867,674399 | -0,509015926 | 0,152939814 | -3,328210713 | 0,000874057 | 0,031526499 | ENSMUSG00000044254 | Pcsk9 | C |
| ENSMUSG00000066640 | 1386,504143 | -0,501133227 | 0,15222025 | -3,292158743 | 0,000994215 | 0,034148309 | ENSMUSG00000066640 | Fbxl18 | B |
| ENSMUSG00000035095 | 1265,26074 | -0,49740839 | 0,14601428 | -3,406573587 | 0,000657838 | 0,026793806 | ENSMUSG00000035095 | Fam167a | L |
| ENSMUSG00000033278 | 5823,687626 | -0,489370419 | 0,155997959 | -3,137030901 | 0,001706681 | 0,046895529 | ENSMUSG00000033278 | Ptprm | A |
| ENSMUSG00000055737 | 11275,69835 | -0,487660293 | 0,142301031 | -3,426962461 | 0,000610373 | 0,025596754 | ENSMUSG00000055737 | Ghr | A |
| ENSMUSG00000020917 | 248750,2656 | -0,485415237 | 0,144100875 | -3,368579388 | 0,000755566 | 0,029100369 | ENSMUSG00000020917 | Acly | A |
| ENSMUSG00000028127 | 11198,98031 | -0,484529093 | 0,142864456 | -3,391530038 | 0,000695035 | 0,027696109 | ENSMUSG00000028127 | Abcd3 | A |
| ENSMUSG00000027765 | 1940,042542 | -0,480473131 | 0,146740391 | -3,274307289 | 0,001059214 | 0,035292936 | ENSMUSG00000027765 | P2ry1 | A |
| ENSMUSG00000042303 | 3590,747377 | -0,480394093 | 0,133155816 | -3,607758981 | 0,000308853 | 0,016199394 | ENSMUSG00000042303 | Sgsm3 | A |
| ENSMUSG00000054423 | 7860,613419 | -0,480157903 | 0,147766305 | -3,249441081 | 0,00115632 | 0,036830202 | ENSMUSG00000054423 | Cadps | A |
| ENSMUSG00000026171 | 2003,287572 | -0,475908207 | 0,120953374 | -3,934641841 | 8,33209E-05 | 0,007076717 | ENSMUSG00000026171 | Rnf25 | A |

| | | | | | | | | | |
|---|---|---|---|---|---|---|---|---|---|
| ENSMUSG00000032890 | 4314,60039 | -0,470844714 | 0,137998387 | -3,411958087 | 0,00064498 | 0,026328353 | ENSMUSG00000032890 | Rims3 | F |
| ENSMUSG00000044550 | 4810,437094 | -0,462822862 | 0,137479788 | -3,36647932 | 0,000761343 | 0,029200673 | ENSMUSG00000044550 | Tceal3 | |
| ENSMUSG00000041650 | 1965,930976 | -0,458164374 | 0,144045479 | -3,180692489 | 0,001469235 | 0,042462505 | ENSMUSG00000041650 | Pcca | A |
| ENSMUSG00000013593 | 13479,31721 | -0,450102638 | 0,139394124 | -3,228992911 | 0,00124227 | 0,038566924 | ENSMUSG00000013593 | Ndufs2 | A |
| ENSMUSG00000044768 | 2471,310361 | -0,443752964 | 0,130573069 | -3,398502989 | 0,000677557 | 0,027250507 | ENSMUSG00000044768 | D1Ertd622e | A |
| ENSMUSG00000050640 | 2493,891491 | -0,443575349 | 0,132832378 | -3,339361653 | 0,000839712 | 0,031042353 | ENSMUSG00000050640 | Tmem150c | L |
| ENSMUSG00000062444 | 15990,24913 | -0,442435696 | 0,126213915 | -3,505443092 | 0,000455848 | 0,021118924 | ENSMUSG00000062444 | Ap3b2 | B |
| ENSMUSG00000060227 | 8122,33621 | -0,441073421 | 0,123343674 | -3,575971154 | 0,00034893 | 0,017599462 | ENSMUSG00000060227 | Casc4 | B |
| ENSMUSG00000031144 | 30610,87281 | -0,440005673 | 0,097610559 | -4,507767165 | 6,55134E-06 | 0,001159713 | ENSMUSG00000031144 | Syp | |
| ENSMUSG00000038871 | 2493,880401 | -0,437954344 | 0,139484835 | -3,139799 | 0,001690638 | 0,046663634 | ENSMUSG00000038871 | Bpgm | A |
| ENSMUSG00000044026 | 3861,957486 | -0,435872241 | 0,132789134 | -3,282439065 | 0,001029132 | 0,034837008 | ENSMUSG00000044026 | Slc35g1 | A |
| ENSMUSG00000030659 | 27855,75041 | -0,431009843 | 0,111327467 | -3,871549886 | 0,000108146 | 0,008126363 | ENSMUSG00000030659 | Nucb2 | A |
| ENSMUSG00000022098 | 4997,059971 | -0,429936215 | 0,133509581 | -3,22026489 | 0,001280722 | 0,039023193 | ENSMUSG00000022098 | Bmp1 | B |
| ENSMUSG00000024038 | 5328,994454 | -0,429349797 | 0,135793715 | -3,161779585 | 0,001568082 | 0,044412909 | ENSMUSG00000024038 | Ndufv3 | A |
| ENSMUSG00000027495 | 10974,93641 | -0,425472032 | 0,131565394 | -3,233920556 | 0,001221034 | 0,038229998 | ENSMUSG00000027495 | Fam210b | A |
| ENSMUSG00000042203 | 5472,431258 | -0,423803738 | 0,126612397 | -3,3472531 | 0,000816167 | 0,030408377 | ENSMUSG00000042203 | Tbc1d22b | A |
| ENSMUSG00000029467 | 91654,93969 | -0,420004883 | 0,131631278 | -3,190768091 | 0,001418951 | 0,041399196 | ENSMUSG00000029467 | Atp2a2 | A |
| ENSMUSG00000032557 | 7881,072312 | -0,416455263 | 0,12993674 | -3,205061654 | 0,001350336 | 0,040250873 | ENSMUSG00000032557 | Uba5 | A |
| ENSMUSG00000017764 | 1951,666245 | -0,415361445 | 0,12765493 | -3,253783024 | 0,001138792 | 0,036588421 | ENSMUSG00000017764 | Zswim1 | C |
| ENSMUSG00000020260 | 14551,26937 | -0,413337849 | 0,113697289 | -3,635423955 | 0,000277524 | 0,015205994 | ENSMUSG00000020260 | Pofut2 | A |
| ENSMUSG00000066900 | 5646,041715 | -0,412707629 | 0,130861997 | -3,15376226 | 0,001611803 | 0,045302747 | ENSMUSG00000066900 | Suds3 | A |
| ENSMUSG00000039263 | 8166,600461 | -0,41124954 | 0,10065138 | -4,085880782 | 4,39099E-05 | 0,004788354 | ENSMUSG00000039263 | Npepl1 | A |
| ENSMUSG00000029629 | 2563,479544 | -0,410790686 | 0,121477882 | -3,381608896 | 0,000720627 | 0,028347727 | ENSMUSG00000029629 | Phf14 | A |
| ENSMUSG00000023915 | 4774,617527 | -0,403332537 | 0,12400434 | -3,252567903 | 0,001143673 | 0,036617415 | ENSMUSG00000023915 | Tnfrsf21 | A |
| ENSMUSG00000042178 | 4184,20181 | -0,401835106 | 0,122298582 | -3,285688999 | 0,001017333 | 0,034619398 | ENSMUSG00000042178 | Armc5 | B |
| ENSMUSG00000052942 | 5713,177341 | -0,400983075 | 0,1202595 | -3,334315154 | 0,000855097 | 0,031172947 | ENSMUSG00000052942 | Glis3 | A |
| ENSMUSG00000067924 | 4911,838402 | -0,398071725 | 0,099601952 | -3,996625726 | 6,42518E-05 | 0,005988273 | ENSMUSG00000067924 | Rtl8b | |
| ENSMUSG00000032120 | 7883,8877 | -0,397379363 | 0,107184228 | -3,707442519 | 0,000209363 | 0,012762819 | ENSMUSG00000032120 | C2cd2l | A |
| ENSMUSG00000022769 | 10645,89718 | -0,395391387 | 0,111291307 | -3,552760736 | 0,000381211 | 0,018653657 | ENSMUSG00000022769 | Sdf2l1 | A |
| ENSMUSG00000032839 | 2979,579928 | -0,39196468 | 0,115255284 | -3,400839116 | 0,000671794 | 0,0271818 | ENSMUSG00000032839 | Trpc1 | A |

| | | | | | | | | | |
|---|---|---|---|---|---|---|---|---|---|
| ENSMUSG00000023921 | 2998,880766 | -0,388505658 | 0,099949534 | -3,887018198 | 0,000101483 | 0,007791756 | ENSMUSG00000023921 | Mut | A |
| ENSMUSG00000030780 | 2842,428676 | -0,388439033 | 0,106827124 | -3,636146121 | 0,000276747 | 0,015205994 | ENSMUSG00000030780 | BC017158 | B |
| ENSMUSG00000034850 | 12587,35693 | -0,38443241 | 0,112557331 | -3,415436453 | 0,000636799 | 0,026110172 | ENSMUSG00000034850 | Tmem127 | A |
| ENSMUSG00000041891 | 50559,83839 | -0,382473545 | 0,10077414 | -3,795354102 | 0,000147433 | 0,009905983 | ENSMUSG00000041891 | Lman1 | A |
| ENSMUSG00000029086 | 18314,63456 | -0,380380144 | 0,116190428 | -3,273764879 | 0,001061249 | 0,035292936 | ENSMUSG00000029086 | Prom1 | B |
| ENSMUSG00000028976 | 19224,79516 | -0,379554251 | 0,09891521 | -3,83716772 | 0,000124461 | 0,008881146 | ENSMUSG00000028976 | Slc2a5 | D |
| ENSMUSG00000057530 | 46918,95336 | -0,375862178 | 0,095376939 | -3,940807748 | 8,12077E-05 | 0,006977316 | ENSMUSG00000057530 | Ece1 | A |
| ENSMUSG00000039474 | 21446,12834 | -0,375383165 | 0,101472599 | -3,699354974 | 0,000216148 | 0,013004205 | ENSMUSG00000039474 | Wfs1 | B |
| ENSMUSG00000020256 | 20384,87856 | -0,373202713 | 0,116849154 | -3,1938846 | 0,001403722 | 0,041216149 | ENSMUSG00000020256 | Aldh1l2 | A |
| ENSMUSG00000031634 | 6575,896164 | -0,372847598 | 0,114776193 | -3,248475026 | 0,001160254 | 0,036872805 | ENSMUSG00000031634 | Ufsp2 | A |
| ENSMUSG00000007594 | 23405,25567 | -0,370032438 | 0,104712255 | -3,533802597 | 0,000409627 | 0,019702342 | ENSMUSG00000007594 | Hapln4 | L |
| ENSMUSG00000061306 | 38242,00441 | -0,367326377 | 0,087903887 | -4,178727359 | 2,93145E-05 | 0,003622011 | ENSMUSG00000061306 | Slc38a10 | A |
| ENSMUSG00000041997 | 7979,776404 | -0,367221429 | 0,113850617 | -3,225467187 | 0,001257673 | 0,038866653 | ENSMUSG00000041997 | Tlk1 | A |
| ENSMUSG00000018102 | 7365,912236 | -0,360880958 | 0,113084029 | -3,191263717 | 0,001416519 | 0,041393839 | ENSMUSG00000018102 | Hist1h2bc | A |
| ENSMUSG00000028032 | 7794,246647 | -0,35198861 | 0,101549823 | -3,46616664 | 0,000527936 | 0,023363704 | ENSMUSG00000028032 | Papss1 | A |
| ENSMUSG00000032575 | 34023,93709 | -0,349883098 | 0,09193823 | -3,805632307 | 0,000141442 | 0,009644275 | ENSMUSG00000032575 | Manf | A |
| ENSMUSG00000022685 | 4281,025535 | -0,348980733 | 0,101110197 | -3,451488997 | 0,000557502 | 0,024106381 | ENSMUSG00000022685 | Parn | A |
| ENSMUSG00000003500 | 10383,75467 | -0,347431309 | 0,101088555 | -3,436900551 | 0,000588412 | 0,025116504 | ENSMUSG00000003500 | Impdh1 | A |
| ENSMUSG00000032743 | 2602,916463 | -0,34623877 | 0,106075012 | -3,264093609 | 0,001098149 | 0,03590927 | ENSMUSG00000032743 | D430042O09Rik | A |
| ENSMUSG00000042298 | 2422,136369 | -0,344725972 | 0,108936694 | -3,164461474 | 0,001553702 | 0,04414145 | ENSMUSG00000042298 | Ttc19 | A |
| ENSMUSG00000027230 | 8366,127433 | -0,341784841 | 0,104481234 | -3,271255796 | 0,00107071 | 0,035341356 | ENSMUSG00000027230 | Creb3l1 | A |
| ENSMUSG00000043733 | 20117,70888 | -0,341021098 | 0,105865773 | -3,221259222 | 0,001276287 | 0,039023193 | ENSMUSG00000043733 | Ptpn11 | A |
| ENSMUSG00000035227 | 26177,87793 | -0,340455569 | 0,094488667 | -3,603136555 | 0,0003144 | 0,016304531 | ENSMUSG00000035227 | Spcs2 | A |
| ENSMUSG00000025935 | 29617,13461 | -0,33999618 | 0,093878307 | -3,621669287 | 0,000292708 | 0,015896039 | ENSMUSG00000025935 | Tram1 | A |
| ENSMUSG00000002845 | 5684,630611 | -0,332542156 | 0,093423815 | -3,559500921 | 0,00037156 | 0,018487632 | ENSMUSG00000002845 | Tmem39a | N |
| ENSMUSG00000045752 | 7764,694737 | -0,332314013 | 0,103202827 | -3,220008823 | 0,001281867 | 0,039023193 | ENSMUSG00000045752 | Tssc4 | A |
| ENSMUSG00000032042 | 69776,49007 | -0,329953801 | 0,087325253 | -3,778446539 | 0,00015781 | 0,010385233 | ENSMUSG00000032042 | Srpr | A |
| ENSMUSG00000029472 | 14016,66516 | -0,319355737 | 0,095216426 | -3,353998376 | 0,000796528 | 0,029987906 | ENSMUSG00000029472 | Anapc5 | A |
| ENSMUSG00000026307 | 12454,39985 | -0,308271842 | 0,084767921 | -3,636656861 | 0,000276199 | 0,015205994 | ENSMUSG00000026307 | Scly | A |
| ENSMUSG00000074102 | 4761,144515 | -0,307006184 | 0,096195484 | -3,191482305 | 0,001415448 | 0,041393839 | ENSMUSG00000074102 | Rbm15b | A |

| | | | | | | | | | |
|---|---|---|---|---|---|---|---|---|---|
| ENSMUSG00000079487 | 7484,02724 | -0,297935911 | 0,091362903 | -3,261016241 | 0,001110137 | 0,036022203 | ENSMUSG00000079487 | Med12 | |
| ENSMUSG00000028173 | 13310,53768 | -0,295379867 | 0,093785374 | -3,149530198 | 0,001635332 | 0,045685073 | ENSMUSG00000028173 | Wls | A |
| ENSMUSG00000034586 | 27057,80754 | -0,293478881 | 0,090718227 | -3,235059719 | 0,001216173 | 0,038229998 | ENSMUSG00000034586 | Hid1 | A |
| ENSMUSG00000030282 | 11812,57743 | -0,276735176 | 0,082958168 | -3,335840021 | 0,000850421 | 0,031172947 | ENSMUSG00000030282 | Cmas | A |
